# Stimulation of Gαq Promotes Stress Granule Formation

**DOI:** 10.1101/521369

**Authors:** Androniqi Qifti, Lela Jackson, Ashima Singla, Osama Garwain, Suzanne Scarlata

## Abstract

During adverse conditions, mammalian cells regulate protein production by carefully sequestering the translation machinery in membraneless organelles referred to as stress granules. Here, we show that activation of Gαq promotes the formation of particles that contain stress granule proteins through a mechanism linked to the presence of phospholipase Cβ1 (PLCβ1). In cells, PLCβ1, the most prominent isoform of PLCβ in neuronal cells, localizes to both the cytoplasm and plasma membrane. We show that a major population of cytosolic PLCβ1 binds to stress granule proteins, such as PABPC1, eIF5A and Ago2. PLCβ1 is activated by Gαq in response to hormones and neurotransmitters and we find that activation of Gαq shifts the cytosolic population of PLCβ1 to the plasma membrane, reducing its association to stress granule proteins. The loss of cytosolic PLCβ1 is accompanied by an increase in the size and number of particles containing PABPC1, G3BP1 or Ago2, and a shift of cytosolic RNAs to larger sizes consistent with cessation of translation. Particles containing stress granule proteins are seen when the cytosolic level of PLCβ1 is lowered by siRNA or by osmotic stress but not cold, heat, oxidative or arsenite stress suggesting that their composition is distinct from those formed from other stresses. Our results fit a simple thermodynamic model in which cytosolic PLCβ1 solubilizes stress granule proteins and its movement to Gαq upon stimulation releases these particles to allow the formation of stress granules. Taken together, our studies show a link between Gαq-coupled signals and translation through stress granule formation.

## INTRODUCTION

When cells are subjected to environmental stress, they halt the production of many housekeeping proteins to preserve resources for the synthesis of proteins that will help the cell alleviate the particular stress. These stalled translation complexes, called stress granules, are thought to act as triage sites that protect non-translated mRNAs from degradation until the stress is removed while allowing the synthesis of other proteins (for review see (1, 2)). Stress granules are distinct from processing bodies or P-bodies that store and process mRNA, although they have also been observed under non-stress conditions. Depending on the cell conditions, the mRNA held in these stalled complexes may be degraded, translated or stored until needed. Additionally, studies in yeast subjected to hypo-osmotic stress found that P-bodies and stress granules may form hybrid structures (3) although this behavior has not been observed in higher eukaryotes. Physically, stress granules are phase-separated domains composed of non-translating mRNAs, translation initiation complexes, poly (A)-binding protein, and many additional mRNA-related proteins (4). They consist of a packed core with loosely associated peripheral proteins (5). Stress granules appear when cells are subjected to environmental conditions such as cold or heat shock, exposure to toxic molecules, oxidative stress, hypo- or hyper-osmolarity, UV irradiation and nutrient deprivation. The molecular mechanisms that transmit these different stresses into the cell interior remain largely unresolved.

Although stress granules appear in many types of cells, we focus here on those that form in mammalian cells. Stress granules have been implicated in the pathogenesis of various diseases such as cancer, neurodegeneration and viral infections (1, 6). Many stress granule proteins contain disordered domains and these regions play important roles in the liquid-like nature of stress granules. Neuronal cells, in particular, contain many proteins with disordered domains and so it is not surprising that some neurological diseases (e.g. ALS) have been attributed to abnormal stability of stress granules (see (7)). Thus, it is important that cells have mechanisms to prevent premature formation of stress granules, and to insure their reversible assembly and disassembly.

While stress granules primarily contain proteins associated with translation, it is notable that argonaute 2 (Ago2) can be found in these domains (see (8)). Ago2 is the main nuclease component of the RNA-induced silencing complex (9). Ago2 binds small, silencing RNAs in their double-stranded form, and holds the guide strand after the passenger strand is degraded to allow hybridization with a target mRNA. If pairing between the passenger strand and the mRNA is perfect, as is the case in exogenous siRNAs, then Ago2 will undergo conformational changes that result in mRNA degradation. Alternately, if pairing is imperfect, as is frequently the case for endogenous microRNAs, the conformational changes that allow Ago2 nuclease activity do not occur resulting in a stalled complex (see (10)). Thus, the formation and stability of these stalled complexes and their incorporation into stress granules will alter protein populations, which may alter the down-stream proteins interactions that may ultimately alter the properties of the cell.

The mechanisms through which environmental changes are communicated into the cell to promote stress granule formation are unclear and likely to differ with different types of stress. Here, we show that extracellular signals can impact stress granule formation via G proteins to stall translation. Signaling through G proteins is initiated when external ligands bind to their target G protein coupled receptor that activates intracellular Gα subunits. The Gαq family of G proteins is activated by agents such as acetylcholine, dopamine, bradykinin, serotonin, histamine, etc. (11, 12). Activated Gαq in turn activates phospholipase Cβ (PLCβ) which catalyzes the hydrolysis of the signaling lipid phosphatidylinositol 4, 5 bisphosphate resulting in an increase in intracellular calcium. There are four known family members of PLCβ that differ in their tissue distribution and their ability to be activated by G proteins (12), and these studies focus on PLCβ1 which is the most highly activated by Gαq and is prominent in neuronal tissue. While the major population of PLCβ1 lies on the plasma membrane where it binds Gαq and accesses its substrate, PLCβ is also found in the cytosol in every cell type examined and under different conditions (13, 14).

Several years ago we found that a cytoplasmic PLCβ1 population binds to C3PO, the promoter of RNA-induced silencing, and this binding can reverse RNA-induced silencing of specific genes (14, 15). Association between these two proteins occur in the same region that Gαq binds to PLCβ1 and is upstream of the active site. Thus, while PLCβ1’s catalytic activity is not effected by C3PO binding, its activation by Gαq is eliminated. Subsequent studies showed that the association of PLCβ and C3PO is critical for PC12 differentiation (16, 17), but little or no association is seen in non-differentiating cells leading to the question of whether cytosolic PLCβ1 has other binding partners. Recently, we found that PLCβ1 will bind to and inhibit a neuronal-specific enzyme required for proliferation, CDK16 (18, 19), and this association allows cells to cease proliferation and transition into a differentiated state (17). Again, this association is confined to a specific event that drives neuronal cells out of stemness, and suggests that under non-proliferating, non-differentiating conditions cytosolic PLCβ serves some other function. In this study, we show that a major population of cytosolic PLCβ is bound to stress granules proteins, and that this binding prevents premature stress granule formation. Removal of PLCβ1 from the cytoplasm by stress or by Gαq stimulation promotes particle assembly. The interaction between PLCβ1 and stress granule proteins shows a novel feedback mechanism between the external environment and the protein translation machinery.

## RESULTS

### PLCβ1 binds to stress granule-associated proteins

We began this work by carrying out experiments to determine novel binding partners to cytosolic PLCβ1 in PC12 cells under non-differentiating conditions. Our approach was to isolate the cytosolic fractions of unsynchronized, undifferentiated PC12 cells, and pull down proteins bound to PLCβ1 using a monoclonal antibody. We collected the PLCβ1-bound proteins and identified them by mass spectrometry. Unexpectedly, we found that ∼30% of the total proteins associated with cytosolic PLCβ1 are markers for stress granules (1, 20) (Fig. S1A). The most prevalent ones are listed in Fig.1A in terms of the percent of their contribution to the total intensity. In contrast, control studies using cytosolic fractions of cells with reduced PLCβ1 levels only and identified non-specific proteins such as tubulin, actin and mitochondrial proteins (Fig. S1B).

**Figure 1.**
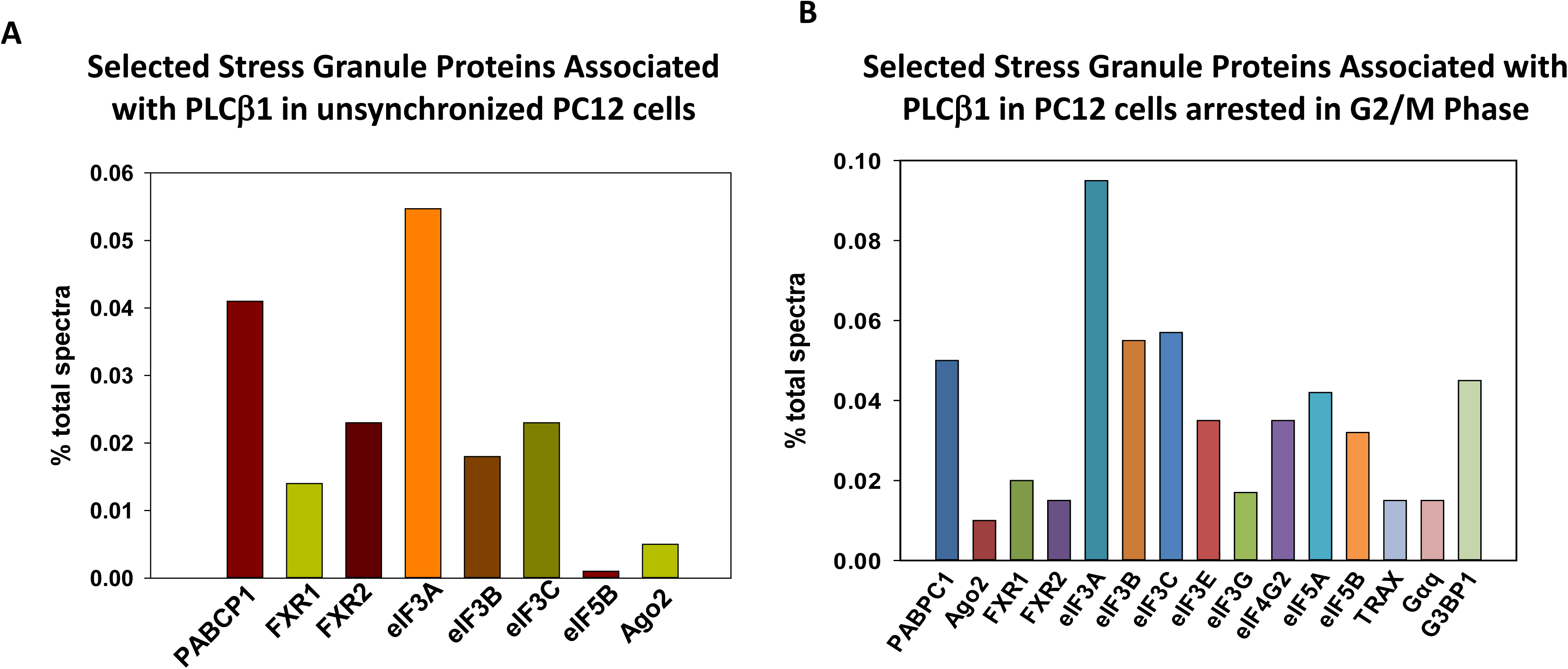
Major proteins complexed to PLCβ1 in PC12 cells as identified by mass spectrometry. Proteins associated with PLCβ1 in cytosolic fractions of PC12 cells were pulled down using a monoclonal antibody in **A-** unsynchronized cells primarily in the G1 phase, or **B-** PC12 cells arrested in the G2/M phase. Levels of proteins were calcuated as described in Methods. Underlined proteins were found to be phosphorylated. All proteins identified in panel (**A**) were also found in an identical study in which proteins associated with Ago2 were identified except for eIF5B. Proteins that were found to be phosphorylated include FXR2 (**A**) and eIF3A, 3B and 3C (**B**).

The studies in Fig.1A were carried out in unsynchronized cells where ∼90% in the G1 phase (17). However, our previous studies suggested that PLCβ1 may change binding partners as cells move through the cell cycle (21). Additionally, PLCβ1 has a nuclear population that may exchange with the cytosolic one (see (22)). Therefore, we repeated the studies in cells where the nuclear population is also available for binding by arresting cells in the G2/M phase where the nuclear matrix has broken down and these results are presented in Fig.1B and Fig. S1C. Again, we find that 32% of the proteins bound to PLCβ1 are markers for stress granules, with the most prominent being eukaryotic initiation factor 5A (eIF5A) and polyadenylate binding protein C1 (PABPC1) (Fig.1B). Additionally, other stress-granule and translation proteins appear in both groups such as FXR1/2, G3BP1 and other eukaryotic initiation factors. It is also notable that Ago2, which is associated with both RNA-induced silencing complexes and stress granules (8), appears in these cells.

The proteomics studies described above are only indicative of potential protein partners of PLCβ1 since they are identified under non-physiological conditions. Therefore, we verified the binding of PLCβ1 to stress granule proteins by several methods. First, we again carried out pull-down studies using the same monoclonal antibody as used above and monitored the association of two stress granule proteins by western blotting. The first, PABPC1, is an established marker for stress granules (20). The second, Ago2 moves into granules under stress conditions (8). Using unsynchronized, undifferentiated PC12 cells, we verified that PABPC1 and Ago2 bind to PLCβ1 (Fig. 2A-B). We then determined whether the levels of these proteins changed when two of PLCβ1’s established binding partners, Gαq and C3PO, are over-expressed (Fig. 2A, and Fig. S7A-B). We find that the level of Ago2 is reduced when the levels of either of these partners is raised, suggesting that Ago2 binds to similar regions of PLCβ1. However, the amount of PABPC1 pulled down with PLCβ1 does not significantly change with over-expression of either Gαq or C3PO (p=0.81 and p=0.54, respectively) but is lowered with Ago2 over-expression. The simplest interpretation of these data is that interactions between PLCβ1 and PABPC1 differ from those of Ago2, and that the reduction in PABPC1 levels with Ago2 over-expression is simply due to a redistribution of the PLCβ1 pool (see *Discussion*). No bands were detected blotting with anti-Actin.

**Figure 2.**
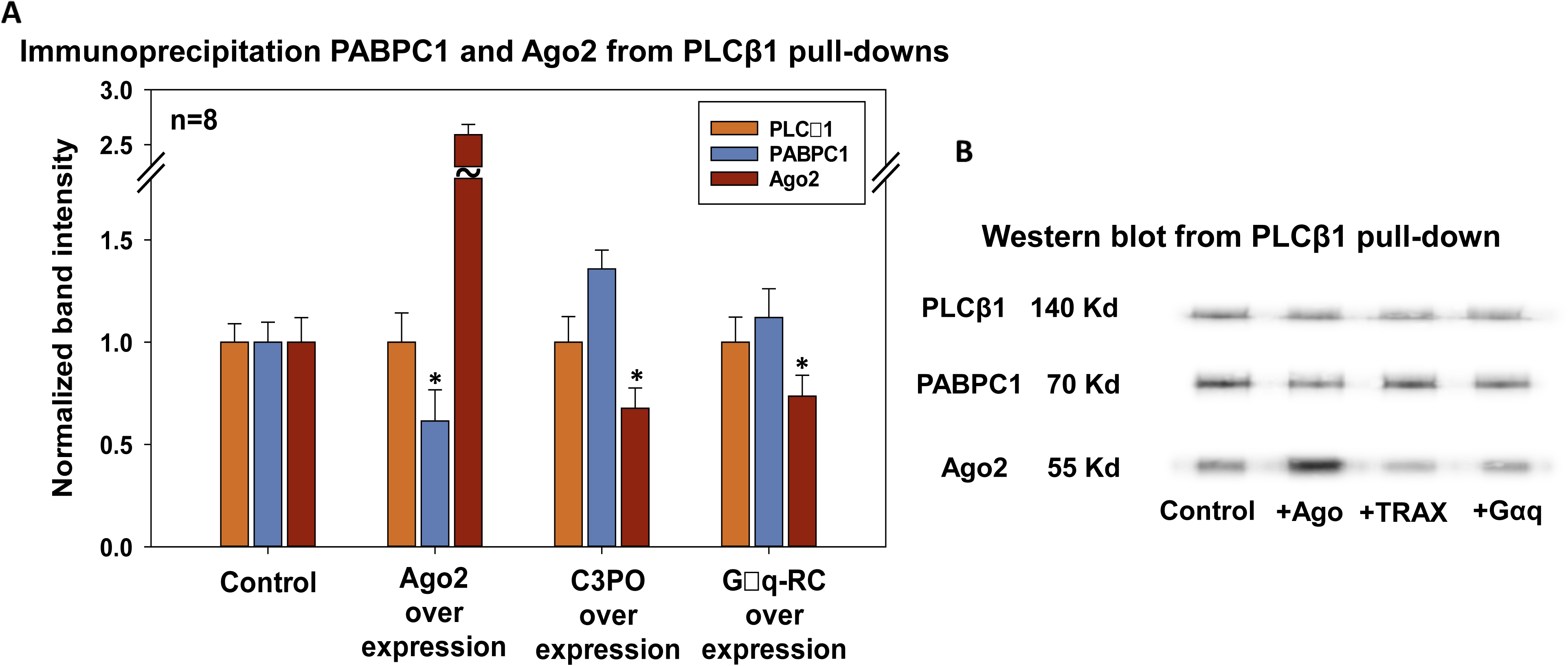
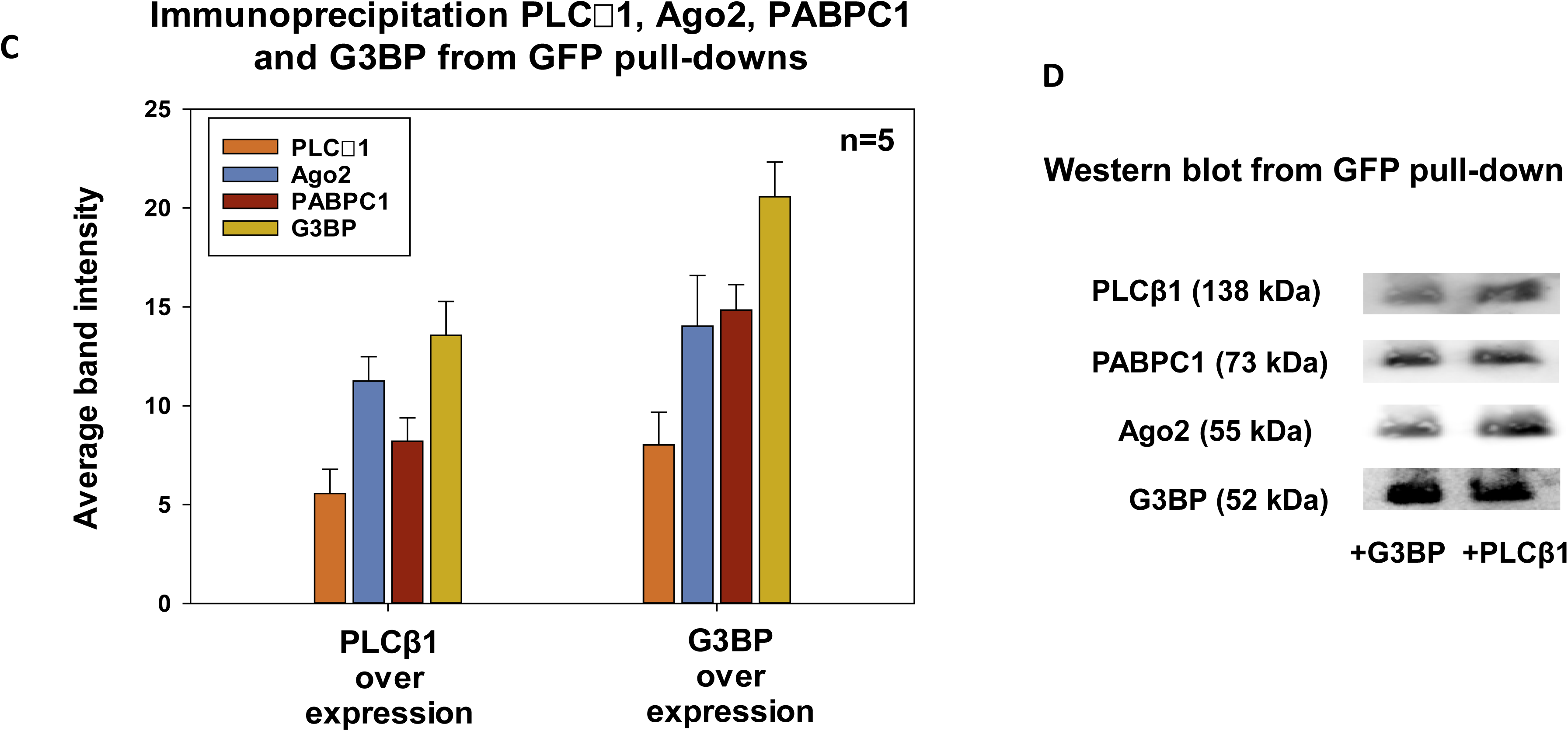
Competitive binding of protein partners to PLCβ1 in PC12 cells. Using a monoclonal anti-PLCβ1 to isolate PLCβ1-associated proteins from the cytsolic fractions of undifferentiated PC12 cells, we monitored two stress-granule associated proteins, PABPC and Ago2. Figures **A, B** presents a compilation of band intensities in mock transfected cells (control) and in cells over-expressing Ago2, C3PO or constituatively active Gαq, where * denotes p<0.001. Differences between C3PO and Gaq pulled down with PLCβ1/PABPC1 were not significant, p=0.81 and 0.54, respectively. Data are from n=8 independent experiments and S.D. is shown. A similar study was repeated using monoclonal anti-GFP antibody to isolate the eGFP tagged on either PLCβ1 or G3BP1 and the associated proteins from whole transfected PC12 cells. A non-binding conrol, actin, did not show bands. Figures **C, D** presents a compilation of band intensities in mock transfected cells (control) and in cells over-expressing PLCβ1 or G3BP1. Data are from n=5 experiments for PLCβ1, PABPC1 and Ago2 and n=2 for G3BP1. A non-binding control, IQGAP2, did not show bands. Full blots for representative samples can be found in the supplemental section, Fig. S7-8.

We wanted to verify that similar results are obtained when a different PLCβ1 antibody is used. Noting that differs from other PLCβ family members mainly in the long 400 amino acid C-terminal domain where Gαq and C3PO bind, and where some stress granule proteins may bind (*see below*), we repeated the pull down studies where the antibody epitope would not occlude any potential binding partners. Specifically, we transfected undifferentiated, unsynchronized PC12 cells with eGFP-PLCβ1 where the eGFP tag is tethered to the N-terminus, and carried out the pull down studies using anti-eGFP or eGFP tagged to another stress granule marker,G3BP1. The results (Fig. 2C-D **and** Fig. S8) show association between PLCβ1 and PABPC1, Ago2 and G3BP1. No bands were seen blotting for a non-binding protein. Taken together, these studies support the idea that cytosolic PLCβ1 associates with stress granules proteins.

### PLCβ1 associates to Ago2 but not G3BP1 in living cells

The above studies suggest that PLCβ1 and Ago2 interactions may be modulated by G protein stimulation and cellular events associated with C3PO. Keeping in mind that C3PO promotes RNA-induced silencing, we set-out to characterize the factors that regulate PLCβ-Ago2 association. We first isolated the cytosolic fractions of unsynchronized, undifferentiated PC12 cells, pulled-down proteins associated with Ago2 and identified them by mass spectrometry. We find that PLCβ is included in the data set. Interestingly, all of the proteins listed in Fig. 1A were found in both PLCβ1 and Ago2 pull-downs (see Fig. S1D).

We measured the association between PLCβ1 and Ago2 in living PC12 cells by Förster resonance energy transfer (FRET) as monitored by fluorescence lifetime imaging microscopy (FLIM). In this method, the FRET efficiency is determined by the reduction in the time that the donor spends in the excited state (i.e., the fluorescence lifetime) before transferring energy to an acceptor fluorophore (see (23)). If we excite the donor with light that has modulated intensity, the lifetime can be determined by the reduction in modulated intensity (M) as well as the shift in phase (φ) of the emitted light. If FRET occurs when the donor is in the excited state, the fluorescence lifetime will be reduced as indicated by a reduced change in modulation and phase shift. The amount of FRET can then be directly determined from the raw data by plotting the lifetimes in each pixel in the image on a phasor plot (i.e. S versus G where S = M*sin (φ) and G = M*cos (φ) (see (24)). In these plots, the lifetimes in each pixel in a FLIM image will fall on the phasor arc for a single population. However, when two or more lifetimes are present, the points will be a linear combination of the fractions with the points inside the phase arc that move towards the right due to shortened lifetimes (i.e. FRET). We note that phasor representation is simply the Fourier transform of the lifetime decay curves but readily displays lifetimes directly from raw data without the need for model-dependent fitting of the lifetimes or other corrections.

In Fig. 3 we show the phasor plot and the corresponding image of a PC12 cell expressing eGFP-PLCβ1 where the phase and modulation lifetime of each pixel in the image is presented. As expected for a single lifetime species, all points fall on the phasor arc (Fig. 3A). When we repeat this study in cells expressing both eGFP-PLCβ1 and mCherry-Ago2 where mCherry is a FRET acceptor, the average donor lifetime drops from 2.5 to 1.7ns, and the points move inside the phasor (Fig. 3B). This reduction in lifetime and movement of the points into the phasor show the occurrence of FRET between the probes. Because the amount of FRET depends on the distance between the fluorophores to the sixth power, and since the distance at which 50% transfers for the eGFP / mCherry pair is ∼30Å (25), our results indicate direct interaction between PLCβ1 and Ago2 in the cytosol.

**Figure 3.**
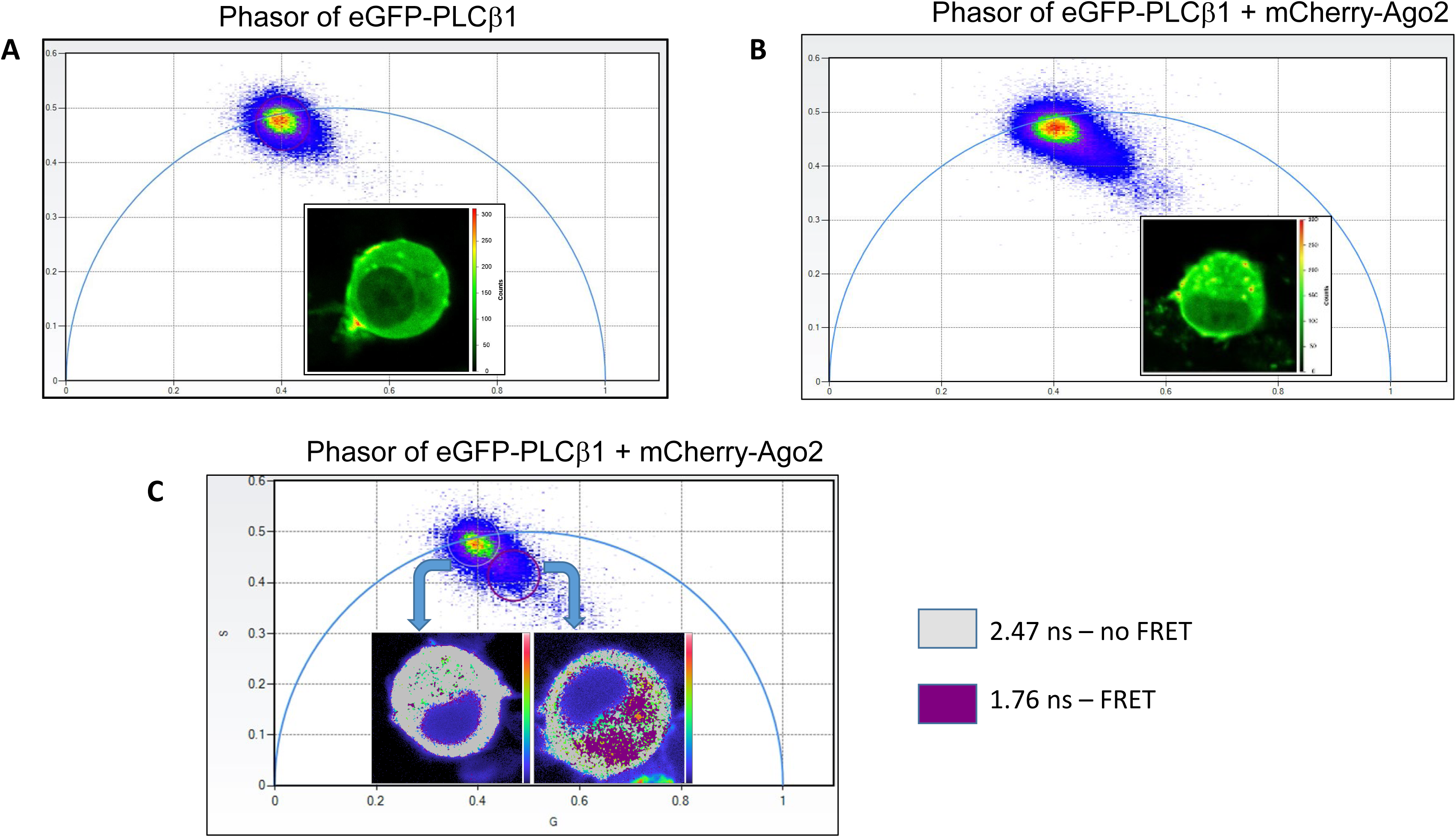
PLCβ1 binds to Ago2 in PC12 cells. Examples of phasor plots in which PC12 cells were transfected with (**A)** eGFP-PLCβ1 or (**B)** eGFP-PLCβ1 and mCherry-Ago2 where the raw lifetimes are plotted as S versus G (see text). Each point in the phasor plots corresponds to the lifetime from the eGFP-PLCβ1 emission measured in each pixel from the corresponding cell image shown in the graph. Images are representative of 5-8 cells over 3 independent experiments. **C –** a phasor diagram where the non-FRET and FRET points are selected and the pixels are shown in the cell images. We note that no significant FRET was detected in control studies of eGFP-C3PO and mCherry-Ago2 (Fig. S6).

We can select points in the phasor plots and visualize their localization in the cell image. We find that the points corresponding to FRET, and thus eGFP-PLCβ1 / mCherry-Ago2 complexes, are localized in the cytoplasm (Fig. 3C). In contrast, points corresponding eGFP-PLCβ1 alone are found both on the plasma membrane and the cytoplasm.

Additionally, we tested the extent of association between eGFP-PLCβ1 and eGFP-G3BP1 using homo-FRET (i.e. FRET between identical species). No changes in lifetime and thus no FRET, could be detected. This absence of FRET means that the two fluorophores are separated by more than ∼45 Angstroms. This result suggests that association between the two proteins is indirect.

### eIF5A binds to PLCβ1 competitively with C3PO and Gαq

To remain cytosolic, stress granule proteins must bind to PLCβ1 in a manner competitive with Gαq or it would localize to the plasma membrane. We previously have shown that PLCβ1 binds to C3PO in the same C-terminal region as Gαq (15) and that competition between C3PO/Gαq regulates PLCβ1’s ability to generate calcium signals through Gαq activation, or its ability to reverse siRNA, respectively (26). With this in mind, we searched the proteins identified in the mass spectrometry for stress granule proteins that could draw PLCβ1 away from Gαq. We noted that eIF5A, which is a GTP-activating protein (27), has homologous regions to the GTPase region of Gαq and appears in the mass spectrometry screen, (see *Discussion*) and so we chose it for further testing.

In an initial study, we purified PLCβ1 and covalently labeled it with a FRET donor, Alexa488. We then purified eIF5A, labeled it with a FRET acceptor, Alexa594, and measured the association between the two proteins in solution by fluorescence titrations similar to previous studies (28). We find that the two proteins bind with strong affinity (K_d_ = 27 ± 5nM). However, when we formed the Alexa488-PLCβ1-C3PO complex and titrated Alexa594-eIF5A into the solution, we could not detect FRET (Fig. 4A). This result suggests that eIF5A binds to the same region as C3PO, and similarly, to Gαq (see (15)).

**Figure 4.**
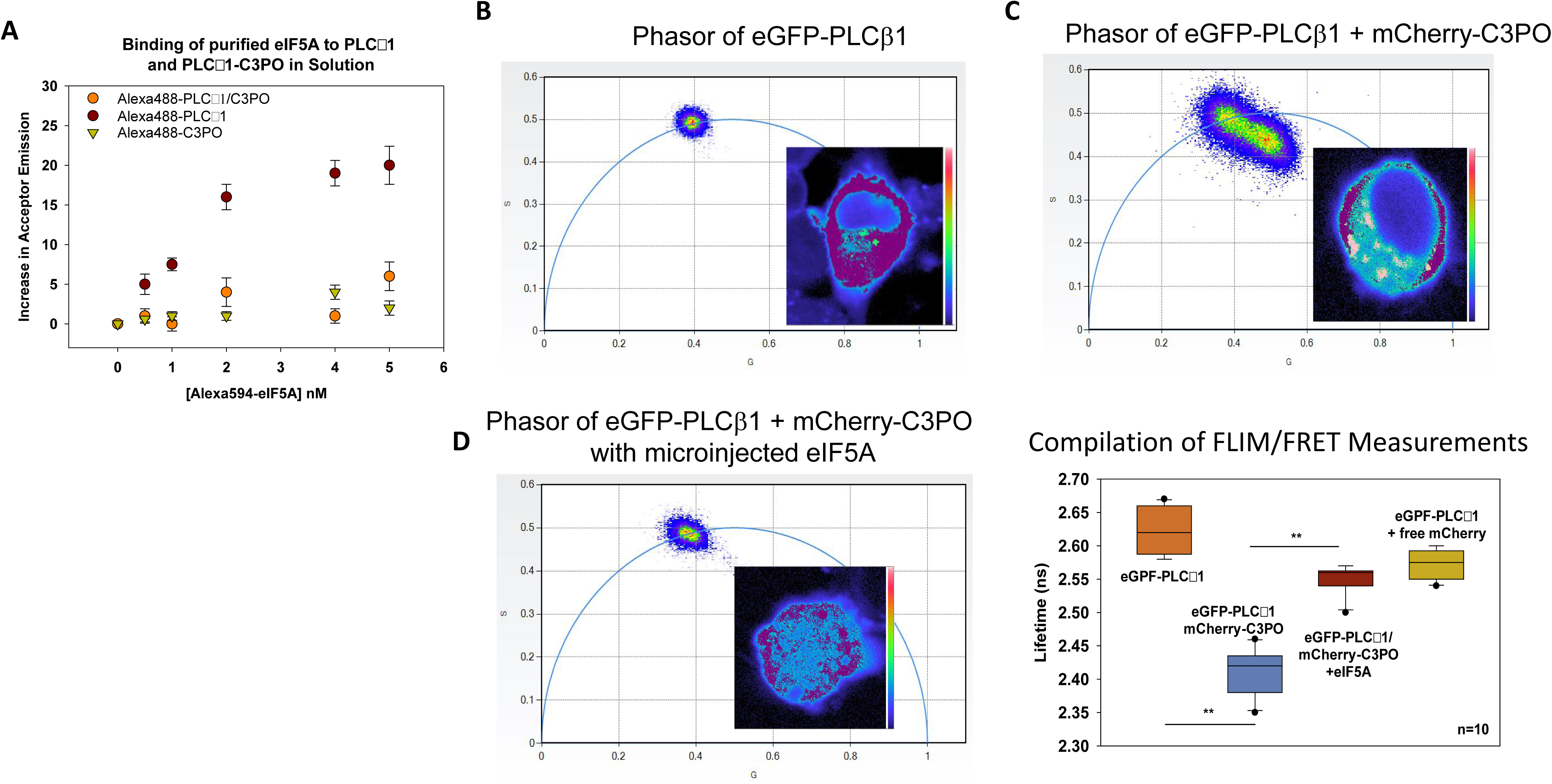
eIF5A competes with C3PO for PLCβ1 in PC12 cells. **A –** Fluorescence binding studies showing changes in FRET bewteen purified Alexa594-eIF5A (Fig. S2A) titrated into solutions of Alexa488-PLCβ1(brown circles), the Alexa488-PLCβ1/C3PO complex (orange circles), or Alexa488-C3PO (green triangles) where FRET is determined by subtracting the fluorescence intensity of Alexa594-eIF5A alone to its intensity in the presence of the different labeled proteins (sensitized emission). Data were corrected for background, where n=3 and SD is shown. **B** – Example of a phasor plot and the corresponding image from a FLIM measurement of eGFP-PLCβ1 expressed in a PC12 cells. The heat map indicates eGFP signal intensity. **C-** A similar study as in (**B)** except the PC12 cell is cotransfeted with eGFP-PLCβ1 and mCherry-C3PO (TRAX). Note the movement of pixels into the interior of the phasor arc due to FRET. Purple dots indicate pixels represented in the phasor plot. **D-** The same study as in (**C**) except this cell was microinjected with purifed, unlabeled eIF5A. Note the movement of points back to the phasor arc showing a loss of FRET due to displacement of C3PO from PLCβ1. **E** - Compilation of eGFP-PLCβ1 lifetime results for 2 independent studies where n=10 cells for each study and where a negative control using free mCherry is included. Comparison of data before and after eIF5A microinjection is statistically significant p=<0.001 t-10.665 and f value for Anova test is 96.606.

To determine whether eIF5A competes with C3PO for PLCβ1 in cells, we transfected PC12 cells with eGFP-PLCβ1 and mCherry-TRAX, to produce fluorescent C3PO (see (29)). Cellular association between these proteins is easily detected by FLIM/FRET (Fig. 4B-C). We then microinjected purified eIF5A to increase its intracellular amount by ∼10nM and find that FRET is completely eliminated (Fig. 4D). This result shows that eIF5A displaces C3PO from PLCβ1 and suggests that both proteins bind to a similar region in PLCβ1’s C-terminus (Fig. 4E).

We confirmed the idea that eIF5A binds to the C-terminal region of PLCβ1 using purified proteins in solution. In these studies, we formed the eIF5A-PLCβ1 complex, chemically cross-linked proteins, digested the samples, separated the fragments by electrophoresis and sequenced the peptides by mass spectrometry (Fig. S2A, S3A-B). We find several interaction sites between the proteins but one of the most prominent is between residues 1085-1095 of PLCβ1and 97-103 of eIF5A which are expected to be close to the Gαq activation region. Taken together, these studies suggest that eIF5A competes with both Gαq and C3PO for PLCβ1 binding and directs PLCβ1 to complexes containing stress granule proteins.

### Effect of osmotic stress on PLCβ1 isoforms

To determine whether PLCβ1 can impact stress granule formation, we started with mild, hypo-osmotic stress (300 to 150 mOsm where we have found has reversible effects on the Gαq/PLCβ signaling pathway in muscle cells (30, 31). We first determined whether hypo-osmotic stress affected the association between PLCβ1 and stress granule proteins within 5 minutes before adaptive changes in the cell occur.

PLCβ1 has two major subtypes (1a and 1b) where the 1a form is the best characterized, and most prevalent subtype, and has additional residues in its C terminus (1142-1216) (32). Although both forms are similarly activated by Gαq, some studies found differences in localization although these appear to be cell type specific (33–35). Surprisingly, we find that a 5 minute exposure of PC12 cells to osmotic stress caused a dramatic reduction in PLCβ1a while the amount of PLCβ1b is unchanged (Fig. 5A). Gαq stimulation did not affect the level of either isoform. Tracking the total amount PLCβ1 using an antibody that recognizes both isotypes, we found that the cytosolic population is preferentially reduced (Fig. 5B). Considering the half-life of total PLCβ1 in PC12 cells is 20 min (17), the data in Fig. 5 suggests that osmotic stress enhances PLCβ1a degradation. After 30 min of stress, the level of PLCβ1b increases as does the level of PABPC1, but PLCβ1a remains low as the cells adapt (Fig. 5B). Because the majority of our studies cannot distinguish between the a and b PLCβ1 isoforms, we will refer to only PLCβ1.

**Figure 5.**
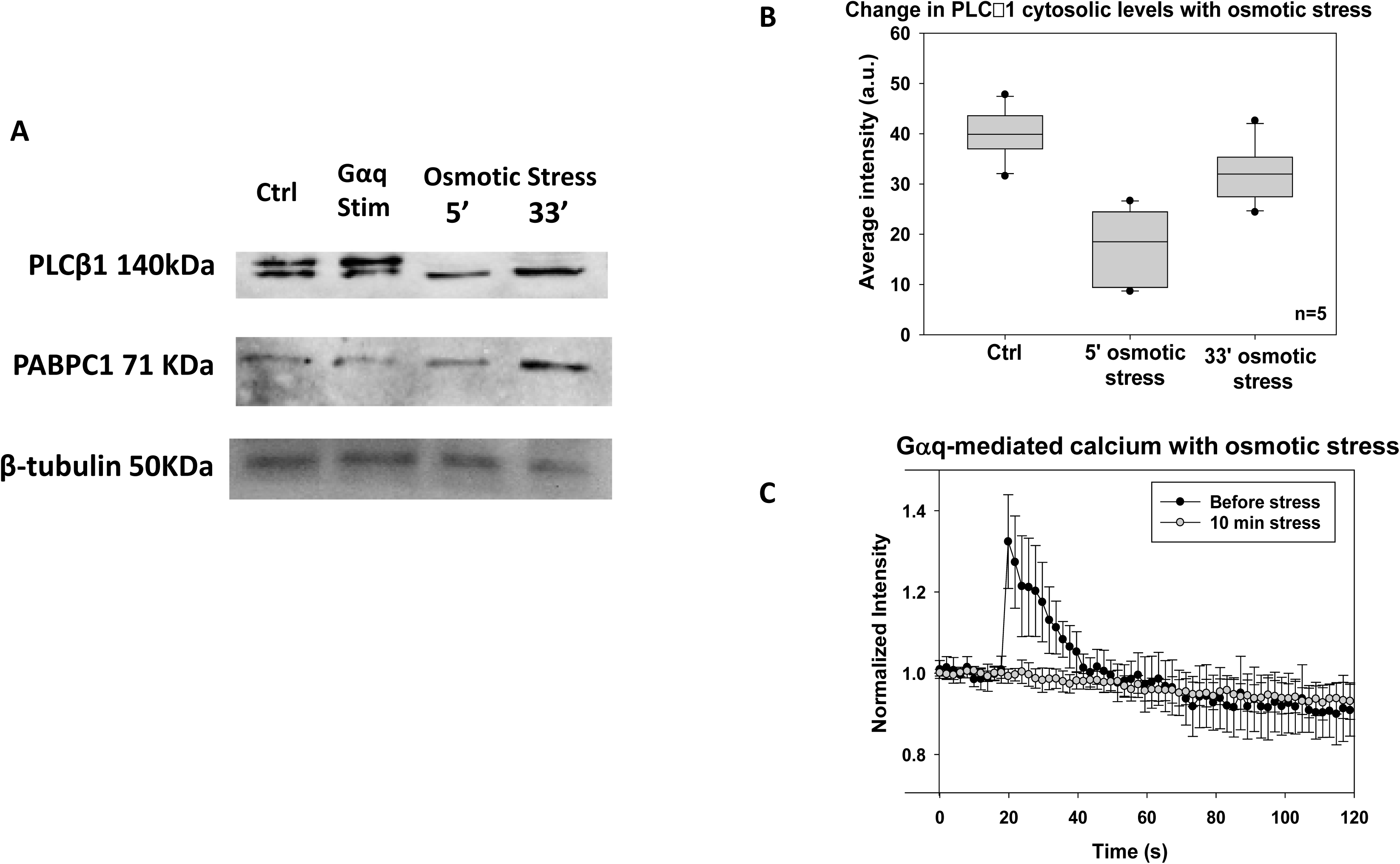
The effect of osmotic stress on cytosolic PLCβ1 in PC12 cells. **A-** A western blot of the cytosolic fractions of PLCβ1 at 300 mOsm under control conditions where PLCβ1a is the upper band of the doublet and PLCβ1b is the lower band. While the intensities of these band are unchanged 10 minutes after Gαq stimulation with 5µM carbachol, PLCβ1a is not detected at 150 mOsm for 5 and 30 minutes. PABPC1 and β-tubulin bands are shown for comparison and n=3. **B** – Results of a study in which eGFP-PLCβ1 was transfected into undifferentiated PC12 cells and changes in cytosolic fluorescence intensity in the a slice in middle of the cell was quantified. Identical behavior was seen over 9 independent experiments, and we note that the plasma membrane population showed similar changes in intensity. **C-** A study showing the change in calcium release when PC12 cells labeled with Calcium Green are stimulated with 5 µM carbachol under basal conditions (300 mOsm) and hypo-osmotic stress (150 mOsm) for 10 minutes where n=12 and SD is shown, and where that basal points for basal and osmotic stress completely overlap.

We determined the ability of hypo-osmotic stress to affect calcium signals generated by Gαq in response to 1µM carbachol. Using the fluorescent calcium indicator, Calcium Green (see *Methods*), we first verified that lowering the osmolarity from 300 to 150 mOsm does not affect the intracellular calcium levels. However, osmotic stress quenches increased calcium in response to Gαq / PLCβ1 activation (Fig. 5C). This loss is consistent with reduced cellular PLCβ1 levels.

### Cytosolic levels of PLCβ1 impact stress granule assembly

It is possible that PLCβ1 binds stress granule associated proteins to prevent premature assembly of stress particles. To test this idea, we followed stress granule formation in PC12 cells under hypo-osmotic stress by counting the number of particles microscopically in undifferentiated, unsynchronized PC12 cells using a 100x objective under normal (300 mOsm) and hypo-osmotic conditions (150 mOsm). We note that this resolution may not capture the formation of small particles (see (5)) and we might be viewing the assembly of primary particles as well as the fusion of small pre-formed ones. In the data shown, we analyzed particle number and sizes in 1.0 µ slices though several cells and report particle sizes in area as seen for each slice. We also note that converting the particles into three dimensions and analyzing the particles gave identical results but with reduced resolution.

We fixed PC12 cells under normal and hypo-osmotic conditions and stained them with monoclonal antibodies to the stress granule marker, PABPC1. In control cells, PABPC1 antibody staining shows ∼750 particles below 25 µm^2^ (Fig. 6A). When PLCβ1 is down-regulated, we find a large increase in PABPC1 particles from 25 to 100 µm^2^ suggesting that loss of PLCβ1 promotes the formation of larger particles. When we apply osmotic stress we find an increase in the number of particles between 25 and 50 µm^2^ (Fig. 6B) However, osmotic stress does not change the size or number of particles in cells where PLCβ1a has been down-regulated suggesting that osmotic stress and loss of PLCβ1 are not additive effects. In another series of studies, we stimulated cells with carbachol to activate Gαq (Fig. 6C). We find that stimulation produces a high number of particles up to ∼150 µm^2^ and down-regulating PLCβ1 does not greatly impact the size or number of PABPC1 particles. Taken together, these studies suggest that loss of PLCβ1 either from down-regulation, osmotic stress or binding to activated Gαq promotes the incorporation of PABPC1 into larger aggregates.

**Figure 6.**
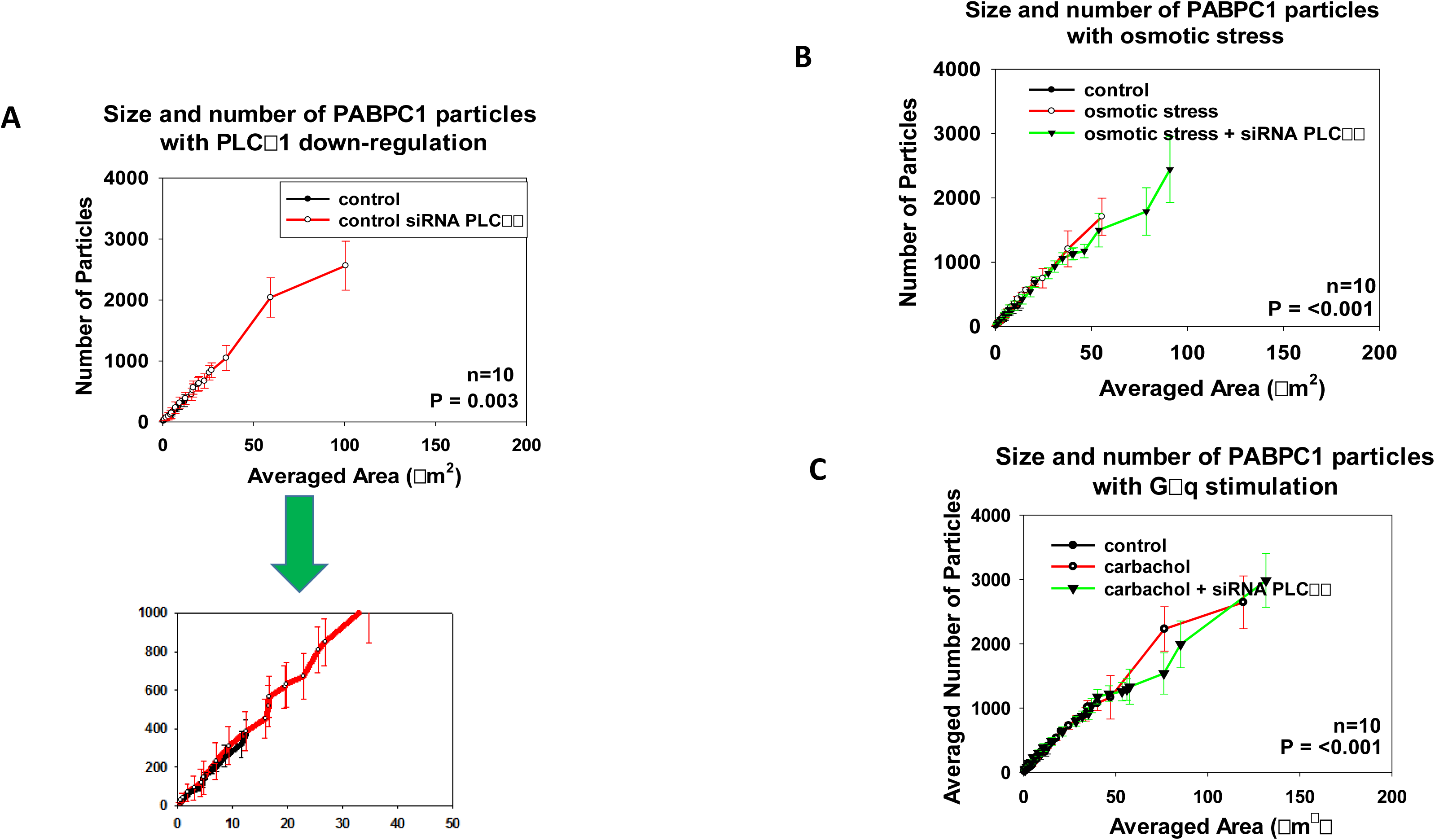
The effect of PLCβ1 on the formation of PABPC1-associated particles in PC12 cells. The size and number of particles associated with monoclonal anti-PABPC1 in the cytosol of fixed and immunostained PC12 cells was measured on a 100x objective and analyzed using Image J (see methods). **A –** Treatment of cells with siRNA(PLCβ1) results in the formation of larger aggregates relative to mock-transfected controls. An enlarged version of the plot is shown directly below to allow better comparison. **B-** While osmotic stress (150 mOsm, 5 minutes) does not affect the size or number of PABPC1-associated particles, PLCβ1 down-regulation causes a significant increase in particle size and number. **C-** Stimulation of Gαq by treatment with 5 µM carbachol also impacts particle size. All measurements are an average of 3 independent experiments that sampled 10 cells, where SD shown and where the p values was determined using ANOVA.

We then tested the effect of PLCβ1 on the size and number of particles associated with Ago2 by immunofluorescence. For Ago2, the number of smaller particles substantially increased when PLCβ1 was down-regulated (Fig. 7A). Unlike PABPC1, the size and number of Ago2-associated particles were not affected by osmotic stress, although an increase in the number of small particles with PLCβ1 down-regulation was still seen (Fig. 7B). Additionally, carbachol stimulation of Gαq resulted in an increase in the number of small particles (Fig. 7C).

**Figure 7.**
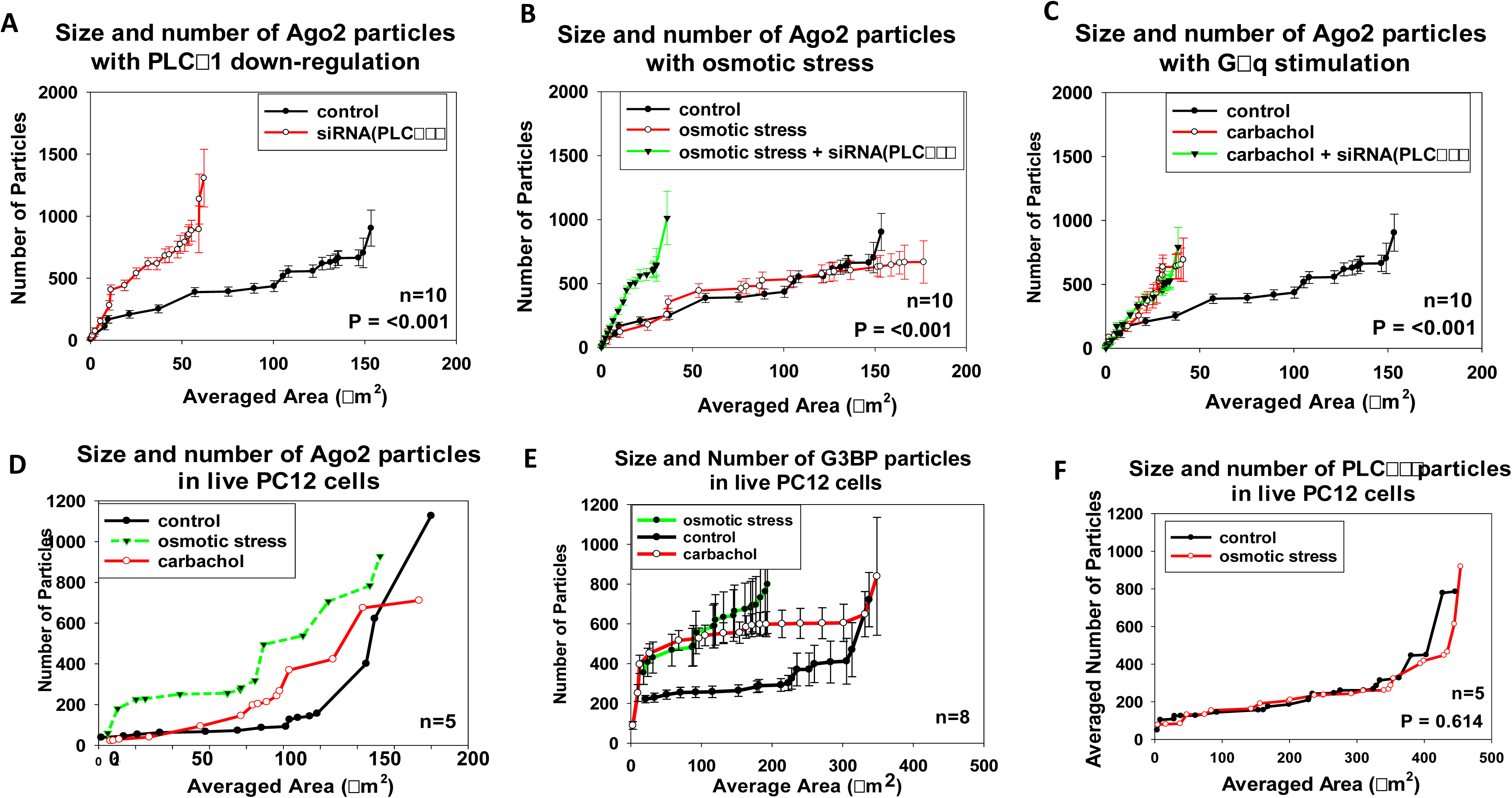
The effect of PLCβ1 on the formation of Ago2 and G3BP1 associated particles in PC12 cells. In similar study in Fig. 6, the size and number of particles associated with anti-Ago2 in the cytosol of PC12 cells was measured on a 100x objective and analyzed using Image J (see methods). **A –** Treatment of cells with siRNA(PLCβ1) results in the formation of numerous small aggregates relative to mock-transfected controls. **B-** Subjecting cells to 5 minutes of osmotic stress (150 mOsm) does not affect the size or number of Ago2-associated particles while siRNA(PLCβ1) causes a significant increase in the number of small aggregates. **C-** Stimulation of Gαq by treatment with 5 µM carbachol results in the formation of a large number of small aggregates. Similar behavior is observed when PLCβ1 is down-regulated. **D-** Analysis of particles associated with mCherry-Ago2 in live PC12 cells under control, 150 mOsm osmotic stress for 5 minutes and 5µ carbachol stimulation. **E-** Analysis of particles associated with eGFP-G3BP1 in live PC12 cells under control, 150 mOsm osmotic stress for 5 minutes and 5µ carbachol stimulation. **F-** Particles associated with eGFP-PLCβ1 in live cells under basal (300 mOsm) and stress (150 mOsm, 5 min) conditions. Measurements in **A-C** are an average of 3 independent experiments that sampled 10 cells, while measurements in **D-F** are an average of 3 independent experiments that sampled 5 cells. For **A-F,** SD is shown and for **A, C-F** the p values compare control versus osmotic stress while **B** compares cotrol versus osmotic stress + siRNA(PLCβ1). P values were determined using ANOVA.

The studies above were carried out in fixed, stained cells. We also followed particle formation in live cells by transfecting PC12 cells with mCherry-Ago2 or eGFP-G3BP1. While the number and areas varied somewhat with the level of transfection, the results show the same trend as the immunostained samples (Fig. 7D-E); that Gαq stimulation or osmotic stress increases the number of associated particles. These studies indicate that reduction of cellular PLCβ1 increases the number of particles of proteins associated with stress granules.

While the increase in particle assemblies could simply be due to the loss of cellular PLCβ1, it may also be due to removal of PLCβ1 from pre-formed particles. To address this question, we transfected PC12 cells with eGFP-PLCβ1 and analyzed the particles (Fig.7F). We could not detect particles below 400 µm^2^ after which the number climbed to ∼1000. No differences were found in cells subjected to osmotic stress. These data suggest that PLCβ1 does not associate with large particles in the cell. In support of this idea, we used fluorescence correlation spectroscopy to monitor changes in the diffusion of eYFP-PLCβ1 when osmotic stress is applied. We find the diffusion coefficient is unchanged at 4.2 +/-0.4 µm^2^/s during the first 10 minutes of exposure to osmotic stress.

The differences in the size and number of PABPC1 versus Ago2 or G3BP1 particles suggest they partition into different types of granules. We tested this idea by monitoring the effect of PLCβ1 and osmotic stress on the colocalization between Ago2 and PABPC1 (Fig. 8). Under normal osmolarity, we find little colocalization between the proteins both at endogenous and knocked-down levels of PLCβ1. However, when the cells are subjected to osmotic stress, colocalization between the species increases, and this increase is more pronounced when PLCβ1 is down-regulated. These results suggest that PABPC1 and Ago2 form distinct particles that may begin to fuse or associate under high stress conditions, such as loss of cytosolic PLCβ1 and osmotic stress.

**Figure 8.**
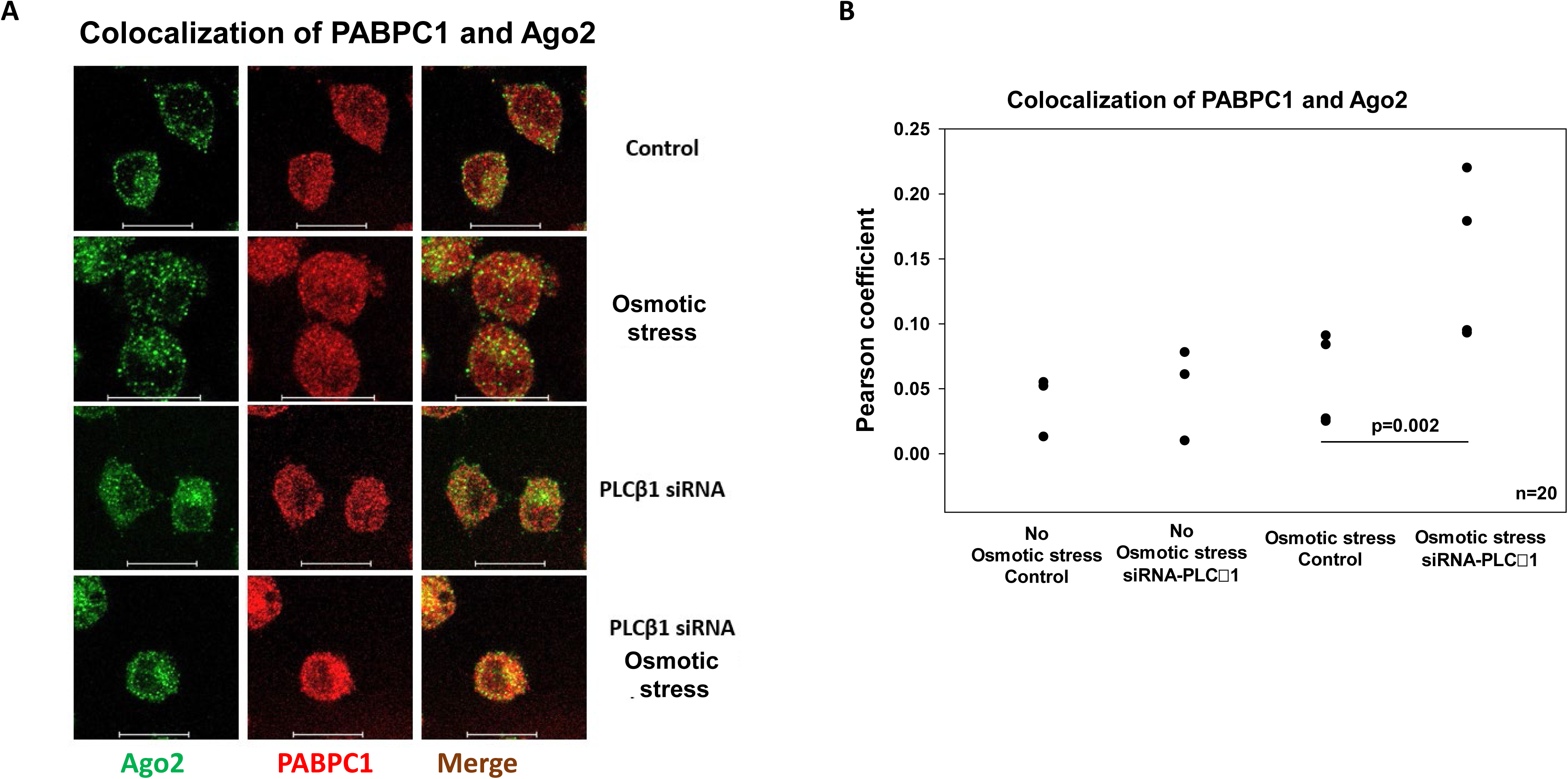
Change in colocalization of Ago2 and PABPC1 particles with osmotic stress. **A-** Images of PC12 cells immunstained for Ago-2 (*green*) and PABPC1 (*red*) under (*top to bottom*) basal conditions, when the osmolarity is lowered from 300 to 150 mOsm, when cells are treated with siRNA(PLCβ1) under normal conditions and under osmotic stress. **B-** Graph of the resulting colocalization data were a significant change between 50% osmotic stress sample with wt-PLCβ1 versus siRNA-PLCβ1 samples is noted p=0.002 n=20. Scale bars are 20 μm long.

### Assembly of Ago2 and G3BP1 stress granules depends on the type of environmental stress

It is probable that osmotic stress produces granules that are different in size, number and composition than those produced by other stresses. We compared the formation of Ago2 and G3BP1 particles under different types of stress by Number & Brightness (N&B) analysis (see *Methods*). This method measures the number of fluorescent molecules associated with a diffusing particle in living cells (36) (see *Methods*). Thus, N&B measurements of cells expressing eGFP-Ago2 will indicate the conditions that promote the formation of aggregates.

In Fig. 9 we present the N&B histograms for some example images of cells expressing eGFP-Ago2 (*top panels*), visualization of aggregates in the cells (*middle panels*), and the corresponding fluorescence images (*bottom panels*) where the purple areas correspond to higher intensities. To analyze these images, we selected the regions of N&B values that correspond to free eGFP (Fig. 9A) as outlined in red in the upper panels, and appear in red in the images in the middle panels. We then determined the number of pixels in each of the N&B images that had values outside these regions that correspond to small Ago2 aggregates (n=2-5) as shown in green, or large Ago2 aggregates shown in purple. The number of pixels corresponding to Ago2 aggregates were tabulated for 10-14 images and are given in Fig. 9.

**Figure 9.**
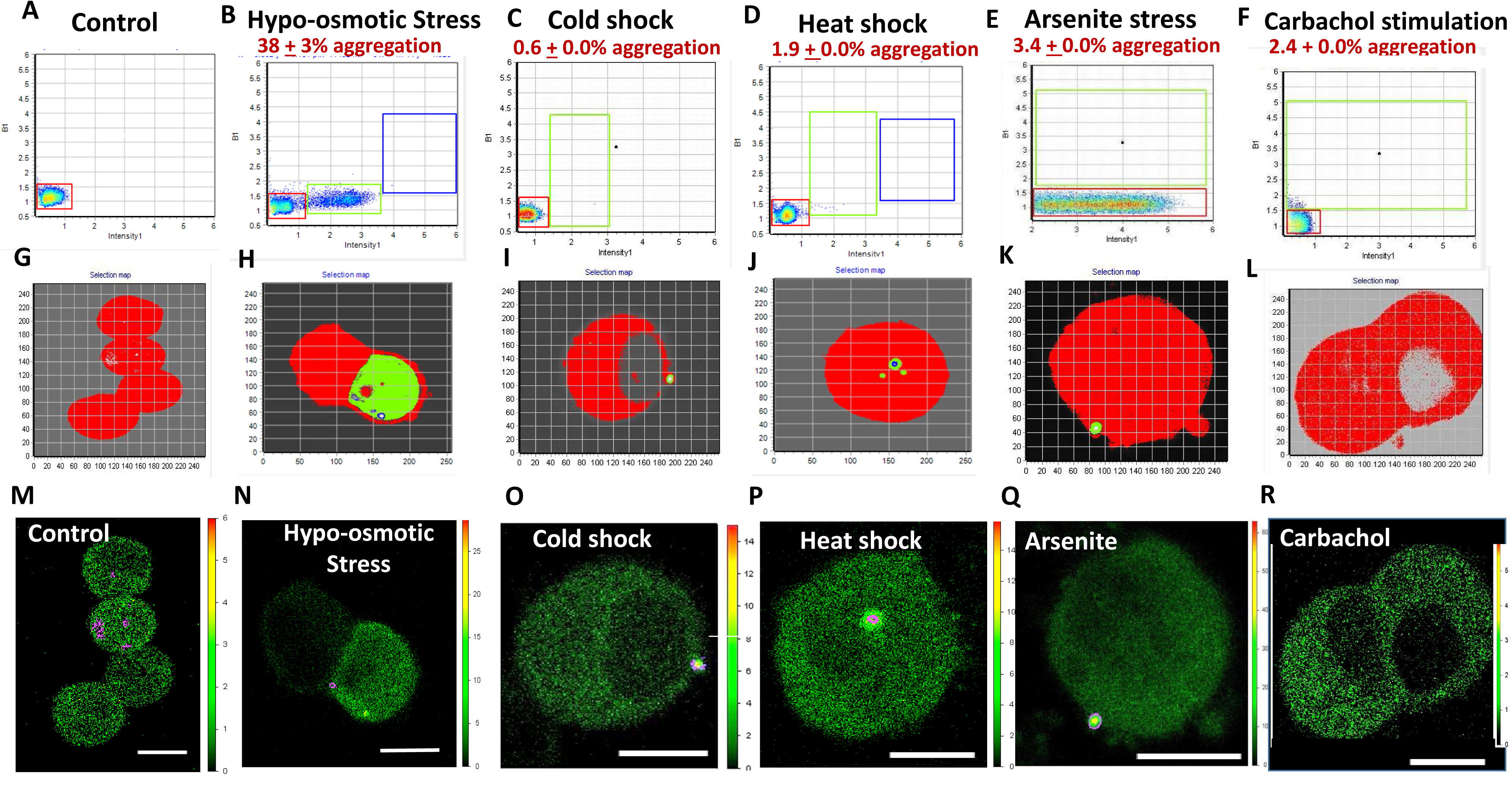
Ago2 aggregation in PC12 cells as monitored by N&B. *The top panels (A-F)* are the N&B data that display the pixels that correspond to different brightness versus intensity values. The pixels contained in the red boxes correspond to the same values found for free eGFP and correspond to monomeric eGFP-Ago2. Points outside the red boxes, shown in green and blue, correspond to higher order species. The pixels corresponding to monomeic (red) and higher order mEFGP-Ago2 aggregates (green and blue) can be seen in the cell images directly below (*middle panels*, **G-L**). The *bottom panels* (**M-R**) show the corresponding fluorescence microscopy images where the purple pixels denote Ago2 aggregates. Panels **A, G, M** are control cells (n=8); Panels **B, H, N** are cells subjected to hypo-osmotic stress (150 mOsm, 5 min) (n=8); Panels **C, I, O** are cells subjected at cold shock 12°C for 1 hr (n=6); Panels **D, J, P** are cells subjected to heat shock at 40°C for 1 hr (n=6); Panels **E, K, Q** are cells subjected to arsenite stress (0.5mM, 10 min) (n=5); Panels **F, L, R** are cells subjected to carbachol stimulation (5µm, 10 min) (n=5). Scale bars are 10 μm long.

We followed Ago2 aggregation in PC12 cells subjected to a variety of stress conditions (Fig. 9). Subjecting cells to osmotic stress resulted shift the distribution of eGFP-Ago2 particles to a point where ∼60% of the eGFP-Ago2 were significantly larger than a monomer (Fig. 9B). It is notable that these Ago2 stress granules formed throughout the cytoplasm, and is notable that only 75-80% of cells showed aggregation. We compared the aggregation of eGFP-Ago2 in cells subjected to other stresses: cold shock (12°C for 1 hour), heat shock (40°C for 1 hour), arsenite treatment (0.5mM for 10 min) (Fig. 9C,I,O **and** 9E,J,P) and oxidative stress (1mM CoCl_2_ 8hrs, Fig. S4). Unlike osmotic stress, these other stresses produced changes in all cells observed but eGFP-Ago2 aggregates were seen as a few large particles rather than evenly distributed through the cell. In a final series of studies, we stimulated cells under normal conditions with carbachol to activate Gαq and promote relocalization of cytosolic PLCβ1 to the plasma membrane (Fig. 9F,L,R). Unlike other stresses, we find the formation of small eGFP-Ago2 aggregates distributed throughout the cytosol. This behavior was seen in every cell tested. These data show that different stresses, including Gαq activation, result in patterns of formation of stress granules containing multiple Ago2 molecules.

We also viewed the aggregation of eGFP-G3BP1 expressed in undifferentiated, unsynchronized PC12 cells. Unlike eGFP-Ago2, PC12 cells expressing eGFP-G3BP1 showed aggregation in untreated conditions, n=4 (Fig. 10A,E,I), and this aggregation increased with hypo-osmotic stress, n=4 (Fig. 10B,F,J), PLCβ1 down-regulation, n=5 (Fig. 10 C,G,K), and carbachol treatment, n=4 (Fig. 10 D,H,L). Unlike the punctate pattern seen for Ago2, G3BP1 aggregation were more diffuse and occurred close to the plasma membrane showing its presence in a wide distribution of stress granules.

**Figure 10.**
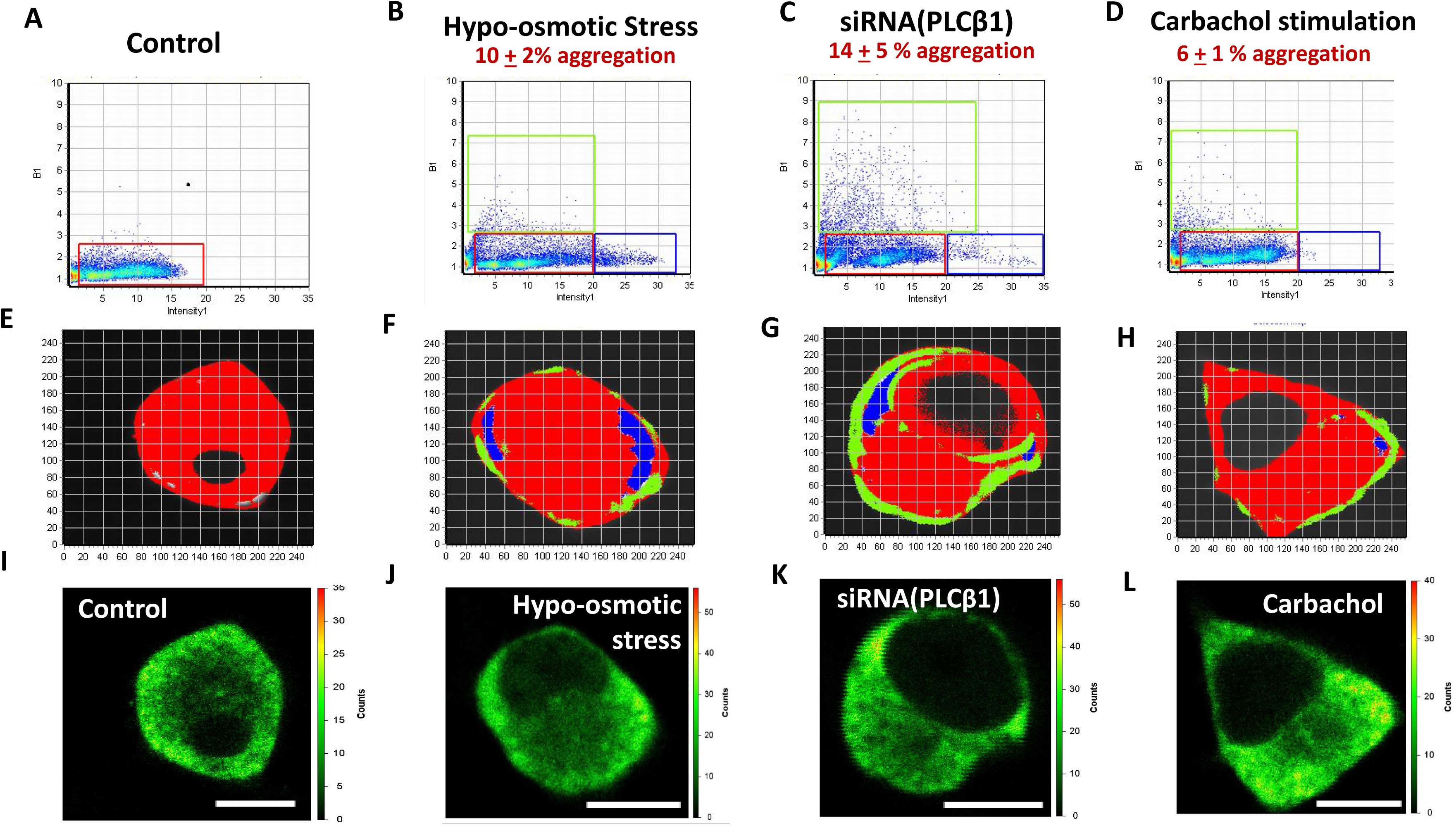
G3BP1 aggregation in PC12 cells as monitored by N&B. *The top panels (A-D)* are the N&B data that display the pixels that correspond to different brightness versus intensity values. The pixels contained in the red boxes correspond to the same values found for untreated PC12 cells transfected with eGFP-G3BP1. Note that untreated samples (**A**, **E** and **I**) contains both monomeric and higher order species (n=4). The green and blue boxes contain pixels corresponding to enhanced aggregation upon hypo-osmotic stress (150 mOsm, 5 min) (**B, F, J**) (n=4), PLCβ1 down-regulation (**C, G, K**) (n=4) and carbachol stimulation (5µm, 10 min) (**D,H, L**) (n=4). Scale bars are 10 μm long.

### Cytosolic PLCβ1 levels impact the size of cytosolic RNAs

The formation stress granules is expected to be accompanied by an increase in the size distribution of cytosolic RNA as mRNA accumulates due to the arrest of translation. We measured the sizes of cytosolic RNA by dynamic light scattering (DLS) (Fig. 11A). Subjecting cells to osmotic stress caused a significant shift to larger sizes. Down-regulating PLCβ1 resulted in a small peak at low molecular weights followed by a broad peak at larger sizes that shifts to the right when compared to control. The small peak is consistent with enhanced C3PO activity due to the relief of inhibition by PLCβ1 (28). Over-expressing Gαq resulted in a similar behavior as treatment with siRNA(PLCβ1). For reference, we show DLS spectra of cytosolic RNA from control and the antibiotic puromycin (*inset*), which halts translation, rendering mRNA in stress granules (see (37)). The RNAs from puromycin-treated cells were almost two-fold higher in molecular weight and show a small peak that corresponds to the sizes seen in control cells. Surprisingly, neither cold, heat, oxidative stress or arsenite increased the average size of cytosolic RNA (Fig. 11B). These data are consistent with the translation arrest and accumulation of higher molecular weight RNAs when the cellular level of PLCβ1 is reduced.

**Figure 11.**
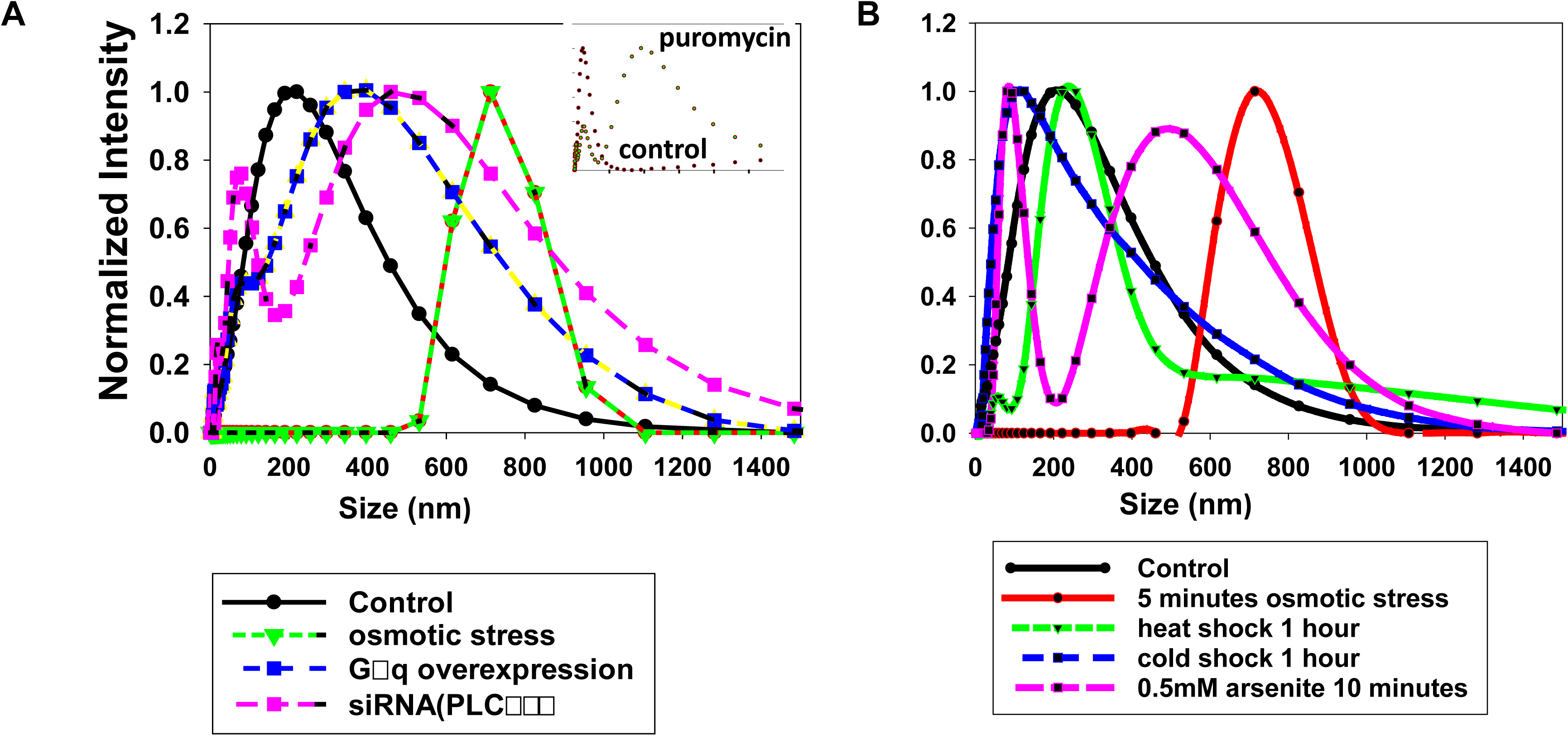
PLCβ1 levels impacts the size of cytosolic RNAs in PC12 cells. **A-** The size distribution of cytosolic RNA isolated from PC12 cells was measured by dynamic light scattering for control conditions (*black*) and was found to shift to higher sizes with hypo-osmotic stress (150 mOsm 5 min, *green*), over-expression of constituatively active Gαq (*blue*), and down-regulation of PLCβ1 (*pink*). Note that a peak at small RNA sizes is seen for these latter two samples due to reversal of inhibition of C3PO activity by PLCβ1. *Inset*– DLS spectra of cytosolic RNA extracted from control cells and cells treated with puromyocin to irreversibly form stress granules where the x-axis is from 0-3000 nm. **B-** An identical study study showing the size distribtion of control cells (*black*) as shown in (**A**), cells subjected to hypo-osmotic stress (150 mOsm 5min, *red*) also shown in (**A**), heat shock (40°C for 1 hour, green), cold shock (12°C for 1 hour, *blue*), arsenite treatment (0.5mM 10 min, *pink*). We note that no changes were observed in cells subjected to oxidatiave stress (12 mM CoCl_2_ 8 hrs) (Fig. S4). Normalized data are shown. Each sample was scanned 3 times with 10 minutes per run. The number of independent samples were control samples 6, PLCβ1 knockdown and Gαq over-expression, 2.

### Cytosolic PLCβ1 levels affect stress granules in smooth muscle cells

Myocytes and other cell types may experience changes in osmotic conditions during their lifetime. With this in mind, we extended these studies to two different smooth muscle cell types, rat aortic smooth muscle (A10) and Wystar Kyoto rat 3M22 (WKO-3M22) cells. We first focused on A10 cells where we identified PLCβ1-associated proteins from a monoclonal antibody pull-down by mass spectrometry under basal conditions and 5 minutes after hypo-osmotic stress (Fig. S2B). Interestingly, a large fraction of proteins pulled down with PLCβ1are associated with transcription which is most likely due to the nuclear population of PLCβ (see (38)). Stress granule proteins appear at lower levels. Many of the stress granule proteins were also found in PC12 cells, e.g. PABPC1 and eIF5A (Fig. 12A), while others, such as Ago2 and FXR, are not seen. Repeating these studies using A10 cells subjected to hypo-osmotic stress for 5 minutes before lysing, resulted in a loss in almost all of these transcription-associated proteins and most stress granule proteins (Fig. 12B). These results show that PLCβ1 binds to stress granule-associated proteins in A10 as well as PC12 cells, and that osmotic stress releases from PLCβ1. While the cellular amount of PLCβ1 in A10 cells is reduced with osmotic stress, the effect is much lower than in PC12 cells (Fig. S2B).

**Figure 12.**
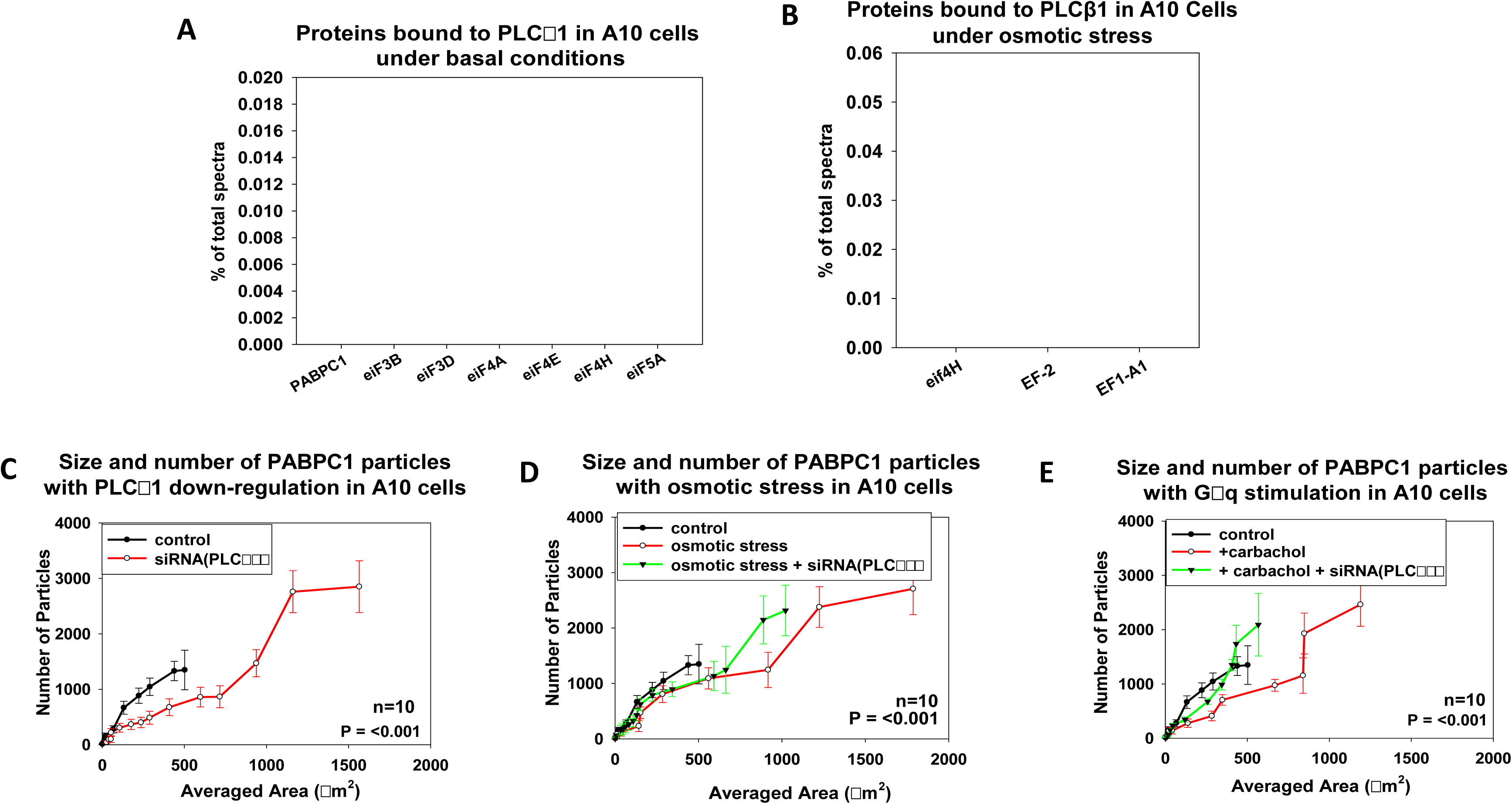
The effect of PLCβ1 on the stress granule formation in A10 cells. **A-** Mass spectrometry analysis for proteins associated with PLCβ1 in A10 cells under normal osmotic conditions and **B-** after 5 minutes of hypo-osmotic stress (150 mOsm). Shown below (**C-E)** are results for The size and number of particles associated with anti-PABPC1 in the cytosol of A10 cells was measured on a 100x objective and analyzed using Image J (*see methods*). **C**-Treatment of cells with siRNA(PLCβ1) results in the formation of larger aggregates relative to mock-transfected controls. **D-** Osmotic stress (150 mOsm) enhances the formation of PABPC1-associated particles and this behavior is promoted by PLCβ1 down-regulation. **E-** Stimulation of Gαq by treatment with 5 µM carbachol also impacts particle size. All measurements are an average of 3 independent experiments that sampled 10 cells, where S.D. is shown and p values were determined using ANOVA.

We measured the formation of PABPC1 particles in A10 cells when the cytosolic levels of PLCβ1 are perturbed, i.e. osmotic stress, siRNA down-regulation and Gαq stimulation (Fig. 12C-E). These cells showed particles that are ∼10 fold larger than in PC12 cells. Like PC12 cells, down-regulating PLCβ1 results in larger particles, and osmotic stress increases the formation of large particles. Also, carbachol stimulation results in a significant increase in particles. In summary, these studies show that the size and number of PABPC1 particles depend on the cytosolic level of PLCβ1.

Both osmotic stress and carbachol also promoted Ago2-related stress granules in WKO-3M22 cells. In Fig. 13A-L we show N&B results of these cells subjected to osmotic and arsenite stress, and carbachol stimulation. Similar to PC12 cells, both osmotic stress and carbachol stimulation promoted Ago2 aggregation while aresenite stress had only a minor effect. Down-regulating PLCβ1 promoted Ago2 aggregation in both control and arsenite stressed cells. Interestingly, down-regulating PLCβ1 did not result in higher levels of Ago2 in cells subjected to osmotic stress or carbachol stimulation suggesting that these conditions lower the level of PLCβ in untransfected cells to give results similar to the down-regulated ones. In a final series of studies, we monitored shifts in the sizes of cytosolic RNA in WKO-3M22 cells with osmotic stress and carbachol stimulation and found significant increases in the RNA sizes under both conditions (Fig. 13M). It is noteworthy that the particles and cytosolic RNAs in WKO-3M22 cells appear much larger than those found in PC12 cells.

**Figure 13.**
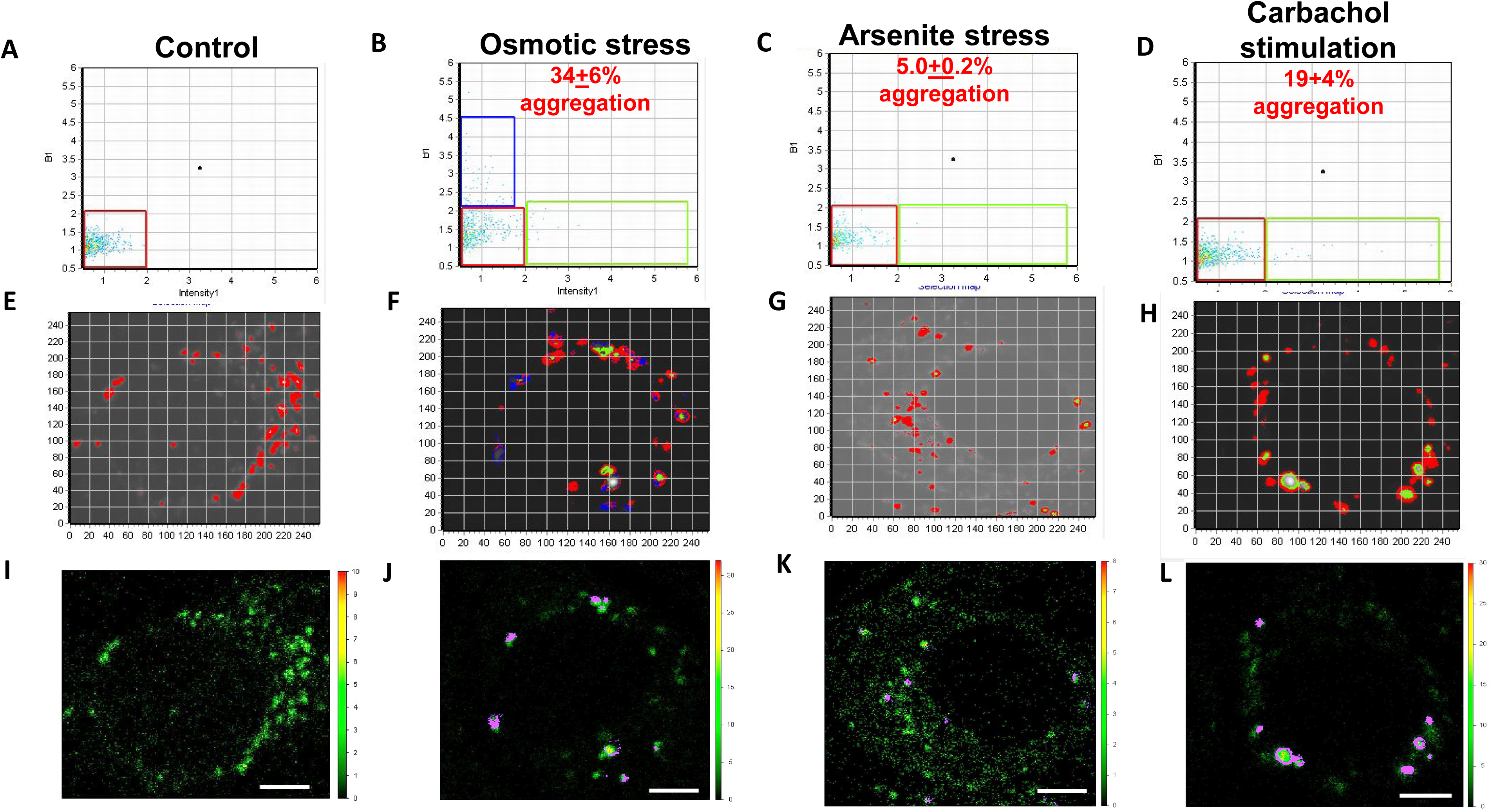
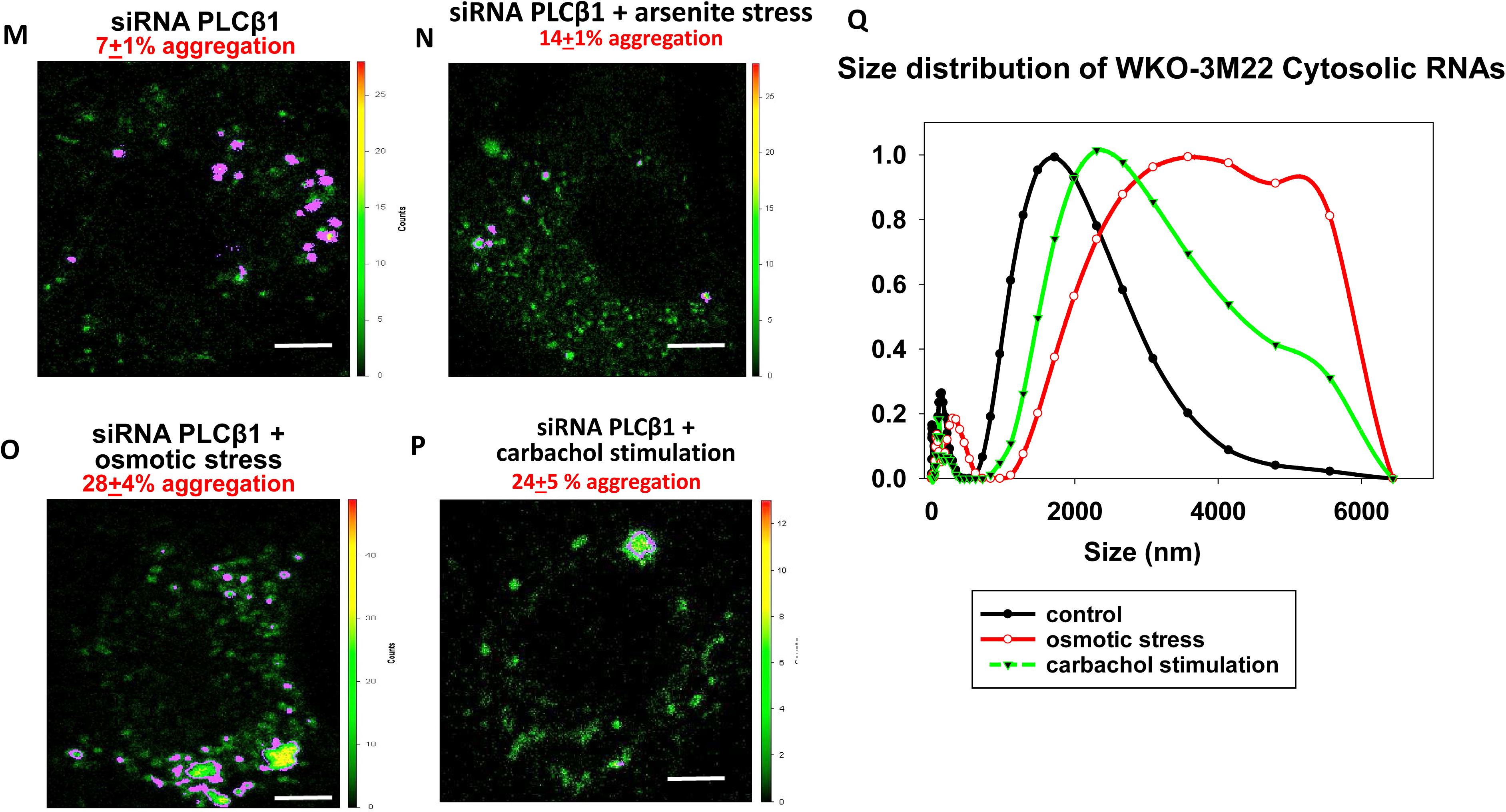
N&B analysis of eGFP-Ago2 aggregation in WKO-3M22 cells. The *top panels* (**A-D**) show graphs of the brightness versus intensity with the pixels of the colored boxes corresponding to the specific regions in the cells (**E-H**) using SIM-FCS 4 software. The *bottom panels* show the corresponding fluorescence microscopy images in ISS (**I-L**). The red box corresponds to monomeric eGFP-Ago2 while points outside this box and in the green and blue boxes correspond to higher order species. Panels **A, E, I are** control cells (n=4); Panels **B, F, J** are cells subjected to hypo-osmotic stress (150 mOsm, 5 min) (n=5); Panels **C, G, K** are cells subjected to arsenite stress (0.5mM, 10 min) (n=6); Panels **D, H, L** are cells subjected to carbachol stimulation (5µm, 10 min)(n=5); Panel **M** are cells treated with siRNA PLCβ1 (n=4); Panel **N** are cells treated with siRNA PLCβ1 and arsenite (0.5mM 10 min) (n=5); Panel **O** are cells treated with siRNA PLCβ1 and osmotic stress (150 mOsm) (n=5); Panel **P** are cells treated with siRNA PLCβ1 and carbachol stimulation (5µM 10 min)(n=3); Panel **Q** is a plot of the size distribution of cytosolic RNA of WKO-3M22 cells as measured by dynamic light scattering for control conditions (*black*), osmotic conditions (150 mOsm 5min, *red*) and stimulation of Gαq by treatment with carbachol (5 µM 10 min, *green*) where n =3. Scale bars are 10 μm long.

### Dependence of stress granule formation on PLCβ1 levels

To understand the dependence of stress granule formation on the concentration of PLCβ, we assume that eIF5A is the primary contact between PLCβ1 and stress granule proteins, although eIF5B or another could be invovled. We can then express the partitioning of eIF5A from the cytosol (c) to the particulate phase (p) as:

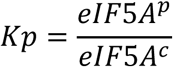

where eIF5A^p^ is the stress granule phase also termed ‘*G’*. The total amount of eIF5A is:

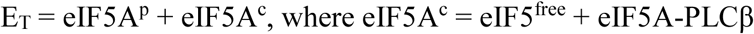

We can express the association between PLCβ and eIF5A in terms of a bimolecular dissociation constant.

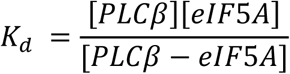

In this equation, PLCβ refers to cytosolic PLCβ. We only consider the cytosolic population and not the membrane-bound one in accord with our results showing that loss of cytosolic of PLCβ promotes stress granule formation. Thus, the total cytosolic amount of PLCβ is:

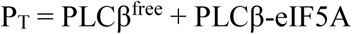

If we combine these equations to determine the relationship between and [PLCβ], we obtain an equation that is quadratic in *G* (i.e. eIF5A^p^.

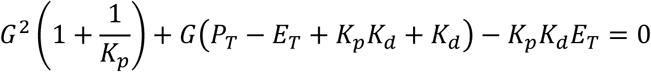

To give:

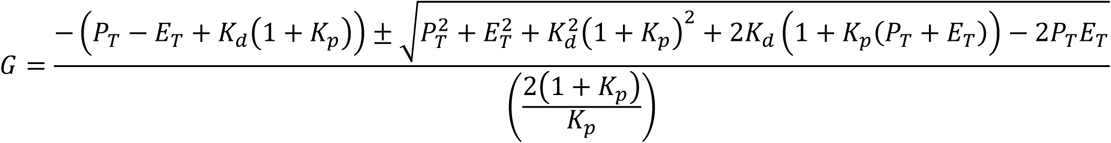

To determine the applicability of this model, we first need to estimate values for *G*. We find a linear dependence between the number of particles and the average area of the particles for PABPC1 (Fig. 14A-B) in PC12 and possibly in A10 cells. This linearity allows us to estimate *G* using either of these measurements. We note that this linearity does not occur for Ago2 particles where stress primarily increases the number of particles rather than the size (Figs. 6-7).

**Figure 14.**
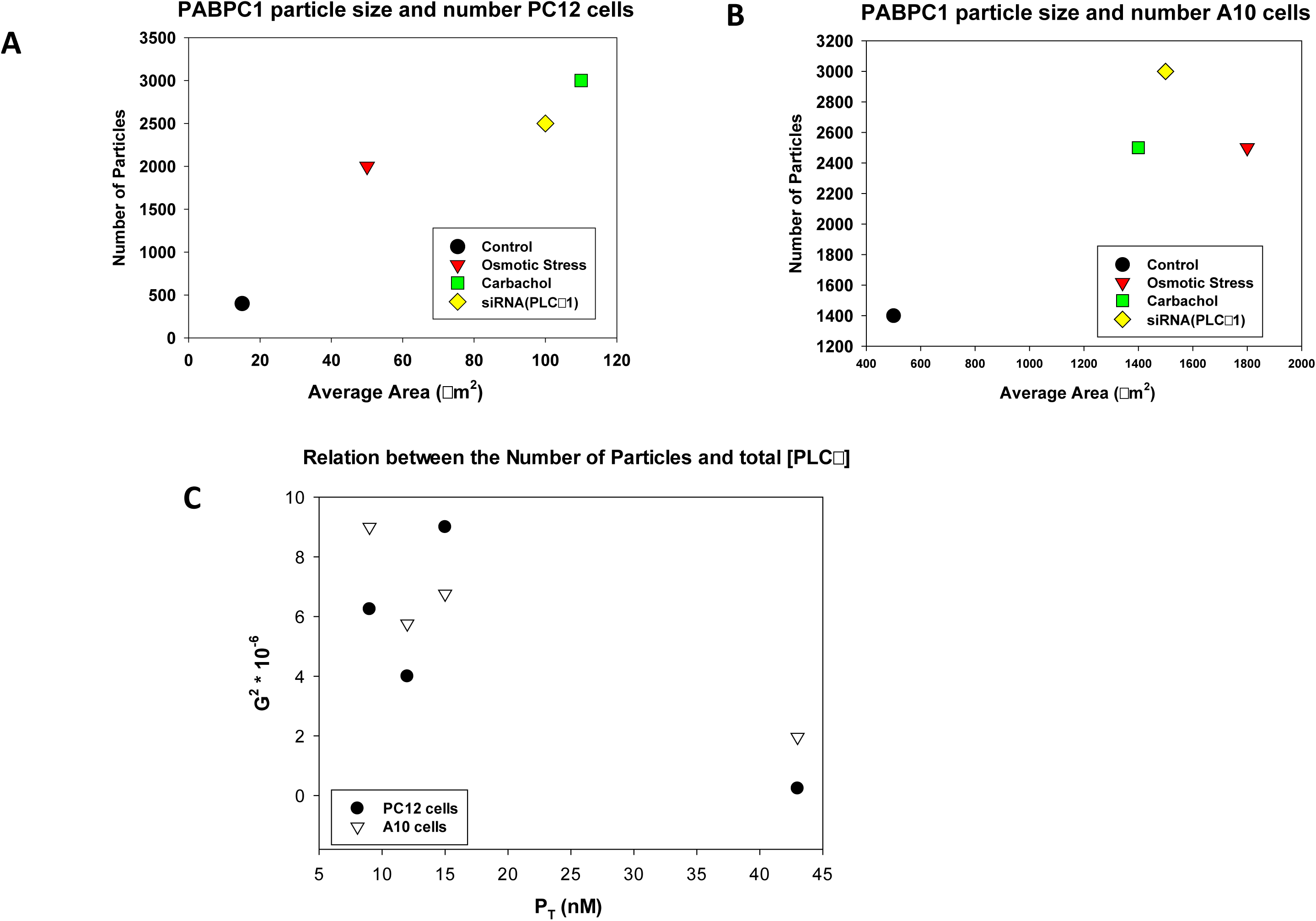
Particle formation is inversely related to PLCβ1 levels. Relationship between the number of particles and average area of PABPC1 particles found in **A-** PC12 and **B-** A10 cells. **C**-shows the inverse relationship of the number of particles and the estimated concentraiotn of total cytosolic PLCβ1.

We can estimate the total amount of cytosolic PLCβ using single molecule fluorescence measurements. Briefly, we measure the number of eYFP-PLCβ1 molecules diffusing in a specific confocal volume after calibration, and we note that we typically over-express ∼2 fold more protein as indicated by western blotting. Our measurements give a cytosolic eYFP-PLCβ1 concentration of ∼43 nM in PC12 cells and ∼49nM in A10 cells which is reduced to 10-15 nM under hypo-osmotic conditions. This decline can be compared with the ∼2.5 fold reduction in cytosolic PLCβ1 upon carbachol stimulation (e.g. Fig. 5B) and the 80-90% reduction in PLCβ1 levels with siRNA treatment. Although these values for PLCβ1 are approximate, we can use them to determine its dependence on stress granules as expressed as *G^2^*. The data in Fig. 14C show the sensitivity of stress granules in this concentration range of PLCβ1 (see *Discussion*)

## DISCUSSION

The studies presented here support the intriguing idea that cells direct signals through the reversible sequestration of proteins in membraneless organelles. In some cases these structures promote protein interactions by reducing the local concentration and mobility of the proteins, whereas in other cases they effectively halt a functional pathway (39, 40). In this work, we show that the atypical cytosolic population of PLCβ1 helps to organize particles containing stress granules proteins in response Gαq signals. The specific types of stress granules that result from Gαq/PLCβ1 signals appear similar to those that appear under osmotic stress, and distinct from those resulting from cold or heat shock, oxidative stress or arsenite treatment. As detailed below, our studies suggest that PLCβ1 regulates the entry of Ago2 and other stress granule proteins into particulates through a simple thermodynamic binding mechanism that is competitive with Gαq, and that is absolutely dependent on the cytosolic level of PLCβ.

The traditional function of PLCβ is to generate calcium signals through molecules that activate Gαq such as acetylcholine, dopamine, serotonin, melatonin, histamine and angiotensin II. It is notable that without Gαq stimulation, the enzymatic activity of PLCβ1 is very low, and Gαq only appears on the plasma membrane suggesting that cytosolic PLCβ1 does not function to modify inositol phospholipids in internal membranes. Thus, any effect of these novel binding partners on PLCβ1’s enzymatic function would immaterial; however, their ability to regulate PLCβ1’s access to Gαq may be important in calcium generation.

PLCβ1 plays multiple roles in cell function that are usually attributed to its enzymatic function. Cocco and others have shown that PLCβ1 can localize to the nucleus to regulate cell growth and differentiation possibly through modulation of PIP_2_ levels in the nuclear envelope (see (41, 42)). Our lab has found that PLCβ1 has a stable cytosolic population that impacts various cell functions such as RNA silencing, and neuronal cell differentiation and proliferation independent of its activity (43, 44). These alternate cytosolic functions of PLCβ only occur at specific and limited times in the cell cycle. For this reason, we set out to determine whether cytosolic PLCβ1 binds to other proteins in non-differentiating cells under non-stimulated conditions. With these parameters in mind, we used a proteomics approach to uncover potential interacting proteins and validated some of these interactions in living cells. Our studies show that under basal conditions cytosolic PLCβ1 associates with stress granule proteins in intact cultured cells.

Stress granules are RNA/protein aggregates that allow cells to halt the translation of non-essential proteins when they are subjected to environmental stress. We find that many of the proteins that associate with PLCβ1 complexes directly contribute to RNA processing and ribosome assembly; and these proteins are found in PLCβ1 complexes isolated from PC12 cells as well as A10 cells. During the initiation stage of protein translation, polyadenylate binding protein PABP binds to the tail of mRNA which then associates to eIF4 allowing the mRNA-protein complex to bind the 40S subunit (45). A cytosolic form of PABP, i.e. PABPC1, was found in our PLCβ1 pull-downs, along with eIF4 subtypes, which binds to PABPC1 (see https://www.uniprot.org/uniprot/P11940). Several other eukaryotic initiation factors were also identified in our analysis.

An important step in the progression of translation is the hydrolysis of GTP on eIF2 catalyzed by the GTPase activity of eIF5. Our results indicate that eIF5A may be a primary binding partner for PLCβ1’s association with stress granule proteins. EIF5A is found at very high levels in cells arrested at the G2/M checkpoint where protein translation is expected to be low. EIF5A has several regions in its sequence that are homologous to Gαq including a region where Gαq directly contacts PLCβ (aa 147-162) as indicated by homologous sequence alignment and chemical cross-linking (see (46)). We directly tested the association between PLCβ1 and eIF5A using purified proteins. Not only was the PLCβ1-eIF5A affinity in the same range as the affinity between PLCβ1 and C3PO (15) but binding was competitive both in solution and in cells. Because the binding site of C3PO overlaps the binding site of Gαq, PLCβ1 association with eIF5A will depend on the level of Gαq activation, and this behavior is observed in our studies. Thus, in the absence of Gαq stimulation, a population of cytosolic PLCβ1 may associate to eIF5A until specific events such as differentiation, causes PLCβ1 binding to shift to C3PO and inhibit of RNA-induced silencing.

We find that cytosolic PLCβ1 also binds to Ago2 as seen in pull-down studies, co-immunoprecipitation and FRET/FLIM. Our ability to disrupt their interactions by the addition of purified eIF5A suggest that it is a primary association. Ago2 is the key nuclease component of the RNA-induced silencing complex (RISC) (see (47)) and our previous work hinted an association between these enzymes (14). Sequence alignment of Ago2 and the TRAX subunits of C3PO shows four homologous regions ranging from ∼20-40 aa in length and from 21-30% identity and 40-56% homology (2-54, 87-119, 202-228, 109-136 on C3PO and 788-826, 555-598, 188-202, 831-858 Ago2). It is notable that at least three of the C3PO regions are potential interaction sites for PLCβ1 binding and at least one of these may be available for PLCβ1-Ago2 binding (28). By this argument, it is possible that PLCβ1 directly binds Ago2 through similar interactions as C3PO.

We determined whether PLCβ1 impacts stress granule formation by monitoring the behavior of two established stress granule markers, Ago2 and PABPC1. We initially used mild osmotic stress that may occur physiologically. Hypo-osmotic stress initiates a series of cellular events to reduce the number of osmolytes in the cell, such as the synthesis of glycogen from glucose, and as well as ion flow (48). While we expected osmotic strength to change the ability of PLCβ1 to interact with stress granules proteins by changes in tertiary or quaternary structure, we were surprised to find a large reduction in PLCβ1a in PC12 cells when osmotic stress is initially applied, although this effect is far less pronounced in A10 cells. PLCβ1a and 1b isoforms differ by ∼20 amino acids in the C-terminal region but are similarly activated by Gαq. Cocco and colleagues have found that PLCβ1b, but not PLCβ1a, prevents cell death under oxidative conditions by impacting levels of key signaling proteins (49). Additionally, these two PLCβ1s may localize differently depending on the cell types (32-35, 50, 51). While our studies cannot adequately distinguish between these isozymes, it would be interesting to see the separate roles they may play in stress granule formation. We note that in conjunction with changes in PLCβ1 levels or properties that occur with osmotic stress, we also varied cytosolic PLCb1 levels by stimulating Gαq to drive PLCβ1 to the plasma membrane, and we down-regulated the enzyme using siRNA(PLCβ1). All these methods showed a connection between cytosolic PLCβ1 and the formation of particles.

Our studies showed that the stress granules formed by osmotic stress differ from ones formed from other stresses. While we find a dramatic assembly of Ago2 under osmotic stress, arsenite, oxidation and temperature shock produce particles that contain monomeric Ago2. Additionally, osmotic stress results in a large increase in the size distribution of cytosolic RNA whereas arsenite, oxidation and temperature does not. Studies in *S.cerevisiae* (3) indicate that hypo-osmotic stress promotes particles composed of markers of both P-bodies and stress granules supporting our findings that subjecting mammalian cells to hypo-osmotic stress forms particles with compositions that differ from other types of stress. Our results also suggest that these latter granules, which have low Ago2 content and are rich in proteins associated with RNA processing, such as G3BP, are poised to prevent the translation of genes whose protein products would not survive arsensite stress or oxidation, such as those involved in phosphorylation (52). In contrast, Ago2-rich granules, which are mediated by the Gαq/PLCβ1 pathway, may shift translation to genes whose protein products allow cells to better respond to external signals. Thus, unlike arsenite or other stresses, Gαq activation may give rise to more physiologically relevant particles.

We monitored the appearance of stress granules under hypo-osmotic conditions structurally by fluorescence imaging and functionally by the accumulation of large cytosolic RNAs. Parker and colleagues have shown that initially, stress granules are small and grow in size in a time-dependent manner (5). Here, we resolved particles over 10 µ^2^ that form in the cytoplasm and the size and number of particles did not vary between 5 and 10 minutes after induction of stress. Additionally, while a very small population of eGFP-PLCβ1 incorporated into particles ∼400 µ^2^ these were unchanged with osmotic stress suggesting that PLCβ1 delivers proteins into particles but does not incorporate into them. PABPC1 was associated with a high number of aggregates whose numbers were affected by the level of cytosolic PLCβ1 as determined by immunofluorescence. Formation of Ago2-associated particles, as monitored by both immunofluorescence and live cell imaging, was sensitive to Gαq stimulation but not to other stresses. Interestingly, the formation of stress granules associated with G3BP1, which does not appear to bind directly to PLCβ1, appears very sensitive to cytosolic PLCβ1 levels and shows extensive and diffuse aggregation. These data suggest that cells respond to Gαq activation by sequestering Ago2, G3BP1 and others into stress granules to halt the production of specific proteins.

Our results also show that the stress granules generated by Gαq activation are more closely related to the ones formed by PLCβ1 down-regulation and osmotic stress in terms of Ago2 aggregates and distinct from those produced by thermal or oxidative stress. Specifically, cold, heat, oxidative or arsenite stress does not result in Ago2 aggregation and does not significantly affect the sizes of cytosolic RNAs even though oxidative stress greatly reduces the cellular level of PLCβ1 along with many cellular proteins (53). Our results correlate well with the variability of stress granule composition with different types of stress (37, 54, 55).

A loss in cytosolic PLCβ1 may arrest the translation of specific proteins by promoting the formation of mRNA-Ago2-associated stress granules. In a preliminary study, we monitored the levels of the heat-inducible chaperone Hsp90A (56) with PLCβ1 down-regulation since Hsp90 is not strongly regulated by PLCβ1-C3PO (14) and because its transcripts may be associated with stress granules formed by arsenite exposure stress (57). Western blotting shows a reduction in Hsp90 levels with siRNA(PLCβ1) consistent with increased formation of Ago2-associated stress granules (Fig. S2C). The idea that sustained Gαq activation can regulate the production of specific proteins is intriguing and a comprehensive study of all of the transcripts affected by PLCβ1 is now underway.

Neurons and cardiomyocytes are long-lived cells and their health depends on reversible assembly of stress granules. We used PC12 cells as model for the role of stress granules in neurological diseases and A10 cells as a model for muscle cells regularly handle changes in osmolarity. We also used WYK-3M22 rat aortic smooth muscle cells as another model of muscle cells that is used as the nonmotensive control for spontaneously hypertensive rats, which are a common models of hypertension (58). We were pleased to find a similar set of stress related proteins in PLCβ1 complexes in the two cell lines with the exception of neural specific proteins and RNA-induced silencing proteins (i.e. Ago2 and C3PO). Thus, PLCβ1 may serve a similar role in many cell types by mediating stress granule formation but not in regulating RNA processing.

We constructed a simple thermodynamic model in which the partitioning of eIF5A into particulates is regulated by its association with PLCβ1 but note that eIF5A can easily be replaced with Ago2. The expression derived from this model shows the scope that PLCβ1 impacts stress granule formation. Specifically, if the total amount of eIF5A is much higher than PLCβ1, then stress granule formation will be independent of PLCβ1. Considering the high concentration of ribosomes in cells, it is difficult to estimate the amount of eIF5A available to bind PLCβ1. We know that microinjection studies that delivered ∼10nM of eIF5A into cells can displace C3PO from PLCβ1 which can give us a quantitative handle for future studies. Regardless of the specific nature of eIF5A and its associated proteins, our data show that there is a concentration range of PLCβ1 that is sensitizes cells to stress granule formation and that this range is under the control of Gαq activation. Additionally, endogenous levels of PLCβ1 help to control premature stress responses.

In summary, our studies suggest a model in which cytosolic PLCβ1 binds to stress granules protein complexes, to keep these proteins disperse under basal conditions. Activation of Gαq shifts the cytosolic population of PLCβ1 to the plasma membrane displacing stress granules proteins and promoting the formation of particles. This dynamic nature of PLCβ1 fits in well with FCS studies showing rapid movement of the enzyme between the cytosol and plasma membrane (28). We propose a model (Fig. 15) in which cells use cytosolic PLCβ1 levels to regulate the formation and timing of protein synthesis, and to prevent the formation of irreversible aggregates. We note that the quenching of Gαq/PLCβ1 calcium signals in cells under osmotic stress suggests that the stress granules may effectively block this signaling pathway.

**Figure 15.**
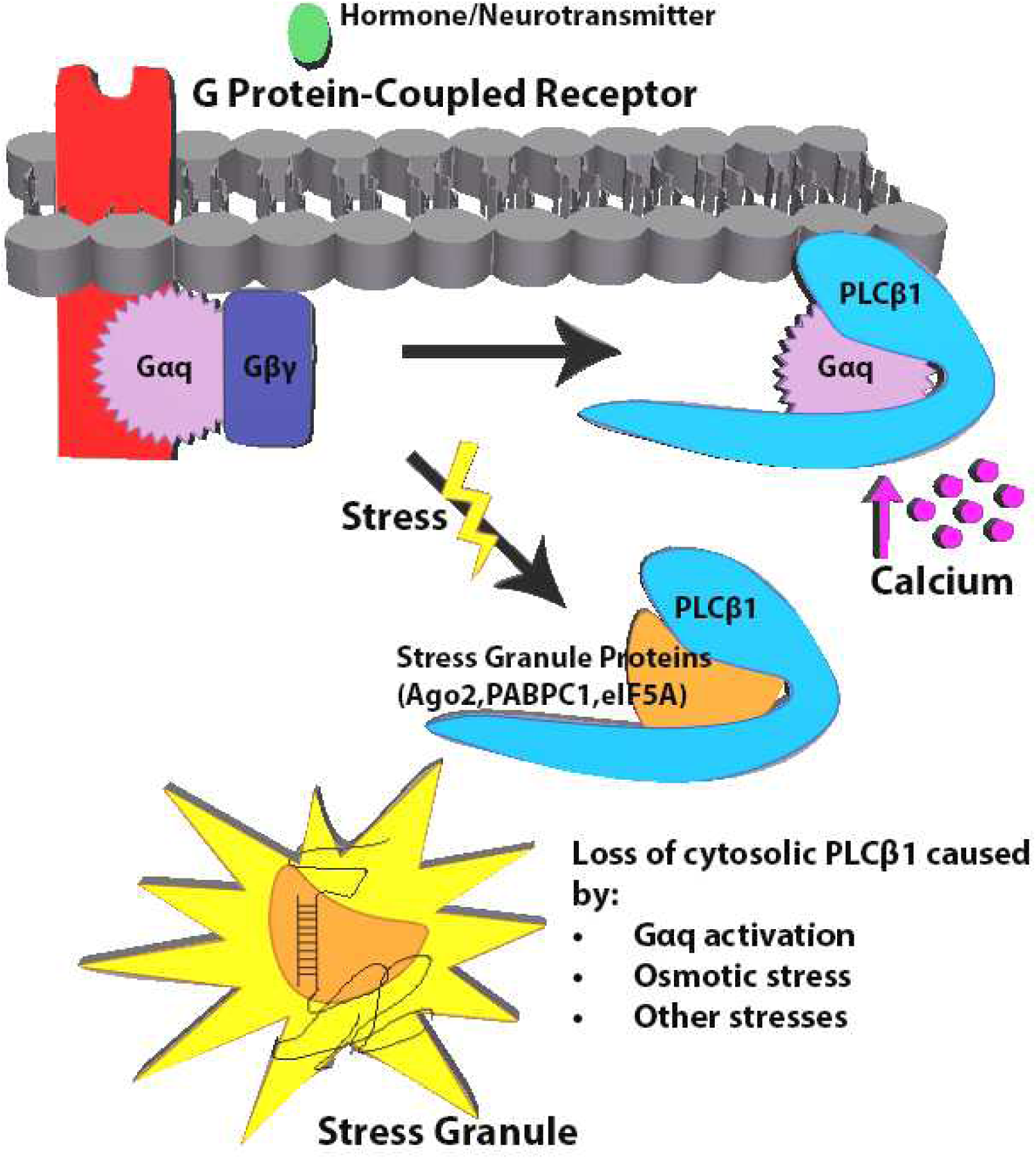
Model of the multiple interactions of PLCβ1 in cells. Under basal conditions, PLCβ1 is distibuted both on the plasma membrane and in the cytosol where it may interact with stress granules proteins. Activation of Gαq promotes movement of PLCβ1 to the plasma membrane releasing stress granule proteins and promoting particle formation.

We have previously found that PLCβ1 plays an important but unknown role in neuronal development. Specifically, the expression of PLCβ1a increases dramatically within the first 24 hrs of PC12 cells differentiation and the slowly decreases (16) leading to the question of why its expression is so variable. PLCβ1 is highly expressed in neuronal tissue where stress granule dysfunction has been implicated in disease (see (59, 60)). Our data suggest that PLCβ1 may act as a chaperone to keep stress granule proteins disperse under basal conditions. It is notable that reduced PLCβ1 levels are associated with a host of neurological disorders that may result from disruptions in calcium signaling, neuronal cells proliferation and differentiation (e.g. (61–64). Schizophrenia and suicide has been shown to specifically involve varying levels of PLCβ1a and 1b in the prefrontal cortex (63). It is also notable that the PLCβ1-associated proteins identified here play important roles in neuronal function. FXR proteins are associated with the most common form of hereditary mental retardation (see (65, 66)) while eIF5A is associated with neuronal growth and brain development (67). It is interesting to speculate connections between PLCβ1 neurological disorders and those associated with FXR and eIF5Awhich may involve dysfunction in stress granule assembly / disassembly.

## MATERIALS AND METHODS

### Cell culture

Rat phenochromocytoma (PC12) and rat aortic smooth muscle (A10) cells were obtained from ATCC. Wystar Kyoto rat 3M22 (WKO-3M22) cells, originally obtained from ATCC, were a generous gift from Dr. Marsh Rolle. PC12 cells were cultured in high glucose DMEM (GIBCO) with 10% heat-inactivated horse serum (GIBCO) and 5% fetal bovine serum (Atlanta Biologicals). Rat aortic smooth muscle (A10) cell lines were cultured in high glucose DMEM with 10 % fetal bovine serum and 1% sodium pyruvate. WKY-3M22 cell lines were cultured in high glucose DMEM (Corning) without L-glutamine with 10% fetal bovine serum, 1% sodium pyruvate, 1% non-essential amino acids (VWR) and 1% L-glutamine (VWR). All cells were incubated at 37°C in 5% CO_2._ Cells were synchronized in the G2/M phase as described (17). Briefly, 2mM thymidine was added to cells for 24 hours, the medium was removed and replaced by fresh complete culture medium for 8 hours after, 40 ng/ml nocodazole was added.

### Plasmids

EGFP-hArgonaute-2 (eGFP-Ago2) was purchased from (Addgene plasmid # 21981) and was prepared in the laboratory of Philip Sharp (MIT). MCherry-Ago2 was a gift from Alissa Weaver (Vanderbilt University). EGFP-G3BP1 was purchased from Addgene (plasmid # 119950) and was prepared in the laboratory of Jeffrey Chao (Friedrich Miescher Institute). MCherry-TRAX-C1 plasmid was constructed by inserting TRAX gene between BamHI and EcoRI restriction sites in mCherry-C1 backbone using T4 DNA ligase (NEB). Plasmid transfections and siRNA knock-downs were done using Lipofectamine 3000 (Invitrogen) in antibiotic-free media. Medium was changed to one containing antibiotic (1% Penicillin/Streptomycin) 12-18 hours post-transfection. For every FLIM experiment, two separate samples were prepared: donor alone, donor in presence of acceptor.

### Co-immunoprecipitation

PC12 cells were lysed in buffer containing 1% Triton X-100, 0.1% SDS, 1x protease inhibitor cocktail and 10 mM Tris, pH 7.4. After 200 μg of soluble protein was incubated with 2 μl of PLCβ1/Ago2 antibody overnight at 4 °C. After addition of 20 mg of protein A-Sepharose 4B beads (Invitrogen), the mixture was gently rotated for 4 h at 4 °C. Beads were washed three times with lysis buffer, and bound proteins were eluted with sample buffer for 5 min at 95 °C. Precipitated proteins were loaded onto two 10% polyacrylamide gel. After SDS-PAGE one gel was transferred to nitrocllulose membranes, proteins were detected by immunoblotting with anti-PLCβ1 (D-8, Santa Cruz) and anti-Ago2 (Abcam ab32381 Lot# GR3195666-1)) antibody.

### Application of stress conditions

For the hypo-osmotic stress conditions, the medium was diluted with 50% water for 5 minutes before it was removed and replaced with Hank’s Balanced Salt Solution (HBSS) for imaging. For the arsenite treatment, a stock solution of 100mM arsenite in water was prepared. Cells were exposed to a final concentration of 0.5mM arsenite for 10 minutes before the medium was removed and replaced by HBSS for imaging. For the heat shock, cells were incubated at 40°C for 1 hour whereas for the cold shock cells were incubated at 12°C for 1 hour. For the oxidative stress treatment, a stock solution of 1M CoCl_2_ was prepared and cells were exposed to a final concentration of 1mM CoCl_2_ for 12-16 hours (overnight) before the medium was removed. Puromycin treatment was carried out at a concentration of 20 μg/mL for 30 minutes in both PC12 and WKO-3M22 cell lines.

### FRET studies between purified PLCβ1 and eIF5A

PLCβ1 was purified by over-expression in HEK293 cells as previously described, see (29). Purification of eIF5A was based on the method described in (27). Purified eIF5A in a pET28-mhl vector was expressed in bacteria (Rosetta 2 DE3 plysS) by inoculating 100 mL of overnight culture grown in Luria-Bertani medium into a 2L of Terrific Broth medium in the presence of 50 μg/mL kanamycin and 25 μg/mL chloramphenicol at 37°C. When OD 600 reached ∼3.0, the temperature of the medium was lowered to 15°C and the culture was induced with 0.5 mM IPTG. The cells were allowed to grow overnight before harvesting and stored at −80°C. Frozen cells from 1.8L TB culture were thawed and resuspended in 150 mL lysis Buffer (20 mM HEPES pH 7.5, 300 mM NaCl, 5% glycerol, 2mM BME, 5mM imidazole, 0.5% CHAPS, protease inhibitor cocktail, 5 μL DNAase) and lysed using a panda homogenizer. The lysate was centrifuged at 15,000 rpm for 45 minutes, added to a cobalt column in binding (20 mM HEPES pH 7.5, 300 mM NaCl, 5% glycerol, 2mM BME, 5mM imidazole), equilibrated in a 4 x 1 mL 50% flurry of cobalt resin, passed the 150 ml of supernatant through each cobalt column at approx.. 0.5ml/min, washed with (1) 20 mM HEPES pH 7.5, 300 mM Knack, 5% glycerol, 2mM BME, 30 mm imidazole, and then (2) 20 mM HEPES pH 7.5, 300 mM NaCl, 5% glycerol, 2mM BME, 75 mM imidazole and eluted with 20 mM HEPES pH 7.5, 300 mM NaCl, 5% glycerol, 2mM BME, 300mM imidazole.

Protein associations were assessed by FRET using sensitized emission. Briefly, PLCβ1a and eIF5A were covalently labeled on their N-terminus with Alexa488 and Alexa594 (Invitrogen), respectively, and the increase in acceptor emission when exiting at the donor wavelength in the presence of Alexa488-PLCβ1 was noted. Studies were repeated using by pre-binding catalytically inactive C3PO with Alexa488-PLCβ1.

### Mass Spectrometry

Mass spectrometry measurements were carried out as previously described at the University of Massachusetts Medical School (68). Cytosolic fractions were isolated from cells, and proteins bound to monoclonal anti-PLCβ1a (Santa Cruz, D-8) were separated by electrophoresis. Protein bands were isolated by cutting the gels into 1×1 mm pieces, placed in 1.5 mL eppendorf tubes with 1mL of water for 30 min. The water was removed and 200µl of 250 mM ammonium bicarbonate was added. Disulfide bonds were reduced by incubating with DTT at 50°C for 30 min followed by addition of 20 µl of a 100 mM iodoacetamide 30 min at room temperature. The gel slices were washed 2 X with 1 mL water aliquots. The water was removed and 1mL of 50:50 (50 mM ammonium bicarbonate: acetonitrile) was placed in each tube and samples were incubated at room temperature for 1hr. The solution was then removed and 200 µl of acetonitrile was added to each tube at which point the gels slices turned opaque white. The acetonitrile was removed and gel slices were further dried in a Speed Vac (Savant Instruments, Inc.). Gel slices were rehydrated in 100 µl of 4ng/µl of sequencing grade trypsin (Sigma) in 0.01% ProteaseMAX Surfactant (Promega): 50 mM ammonium bicarbonate. Additional bicarbonate buffer was added to ensure complete submersion of the gel slices. Samples were incubated at 37 C for 18 hrs. The supernatant of each sample was then removed and placed in a separate 1.5 mL eppendorf tube. Gel slices were further extracted with 200 µl of 80:20 (acetonitrile: 1% formic acid). The extracts were combined with the supernatants of each sample. The samples were then completely dried down in a Speed Vac.

Tryptic peptide digests were reconstituted in 25 µL 5% acetonitrile containing 0.1% (v/v) trifluoroacetic acid and separated on a NanoAcquity (Waters) UPLC. In brief, a 2.5 µL injection was loaded in 5% acetonitrile containing 0.1% formic acid at 4.0 µL/min for 4.0 min onto a 100 µm I.D. fused-silica pre-column packed with 2 cm of 5 µm (200Å) Magic C18AQ (Bruker-Michrom) and eluted using a gradient at 300 nL/min onto a 75 µm I.D. analytical column packed with 25 cm of 3 µm (100Å) Magic C18AQ particles to a gravity-pulled tip. The solvents were A, water (0.1% formic acid); and B, acetonitrile (0.1% formic acid). A linear gradient was developed from 5% solvent A to 35% solvent B in 90 minutes. Ions were introduced by positive electrospray ionization via liquid junction into a Q Exactive hybrid mass spectrometer (Thermo). Mass spectra were acquired over m/z 300-1750 at 70,000 resolution (m/z 200) and data-dependent acquisition selected the top 10 most abundant precursor ions for tandem mass spectrometry by HCD fragmentation using an isolation width of 1.6 Da, collision energy of 27, and a resolution of 17,500.

Raw data files were peak processed with Proteome Discoverer (version 2.1, Thermo) prior to database searching with Mascot Server (version 2.5) against the Uniprot_Rat database Search parameters included trypsin specificity with two missed cleavages or no enzymatic specificity. The variable modifications of oxidized methionine, pyroglutamic acid for N-terminal glutamine, phosphorylation of serine and threonine, N-terminal acetylation of the protein, and a fixed modification for carbamidomethyl cysteine were considered. The mass tolerances were 10 ppm for the precursor and 0.05Da for the fragments. Search results were then loaded into the Scaffold Viewer (Proteome Software, Inc.) for peptide/ protein validation and label free quantitation. These data were analyzed using percentage of total spectra in Scaffold4 software before plotting with Sigma Plot 14.

*Fluorescence correlation (FCS) and*—FCS measurements were performed on the dual-channel confocal fluorescence correlation spectrometer (Alba version 5, ISS Inc.) equipped with avalanche photodiodes and a Nikon Eclipse Ti-U inverted microscope as previously described (31).

### Number and Brightness (N&B) measurements

N&B theory and measurement has been fully described (see (36). Experimentally, we collected ∼100 cell images viewing either free eGFP (control) or eGFP-Ago2, at a 66nm/pixel resolution and at a rate of 4 µs/pixel. Regions of interest (256×256 box) were analyzed from a 320×320 pixel image. Offset and noise were determined from the histograms of the dark counts performed every two measurements. Number and Brightness (N&B) data was analyzed using SimFC (www.lfd.uci.edu).

### N&B analysis

N&B defines the number of photons associated with a diffusing species by analyzing the variation of the fluorescence intensity in each pixel in the cell image. In this analysis, the apparent brightness, B, in each pixel is defined as the ratio of the variance, σ, over the average fluorescence intensity <k>:

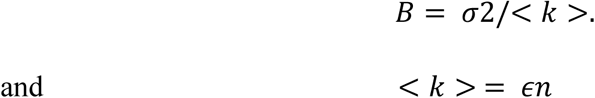

where n is the number of fluorophores. The determination of the variance in each pixel is obtained by rescanning the cell image for ∼100 times as described above. The average fluorescence intensity, <k> is directly related to the molecular brightness, €, in units of photons per second per molecule, and n. B can also be expressed as

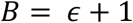

And the apparent number of molecules, N, as

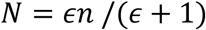

*Fluorescence lifetime imaging measurements (FLIM)—*FLIM measurements were performed on the dual-channel confocal fast FLIM (Alba version 5, ISS Inc.) equipped with photomultipliers and a Nikon Eclipse Ti-U inverted microscope. A x60 Plan Apo (1.2 NA, water immersion) objective and a mode-locked two-photon titanium-sapphire laser (Tsunami; Spectra-Physics) was used in this study. The lifetime of the laser was calibrated each time before experiments by measuring the lifetime of Atto 435 in water with a lifetime of 3.61 ns (Ref) at 80MHz, 160MHz and 240MHz. The samples were excited at 800/850 nm, and emission spectra were collected through a 525/50 bandpass filter. For each measurement, the data was acquired until the photon count was greater than 300. Fluorescence lifetimes were calculated by allowing ω = 80 MHz:

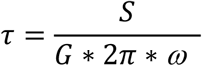

### Statistical analysis

Data was analyzed using Sigma Plot 13 statistical packages that included student’s t-test and one way analysis of variance (ANOVA).

### Dynamic light scattering (DLS)

DLS measurements were carried out on a Malvern Panalytical Zetasizer Nano ZS instrument. For these experiments, mRNA from PC12 cells was extracted following the instructions from the Qiagen Mini Kit (Cat #: 74104). Cells were exposed to stress, treated with siRNA(PLCβ1a), or transfected constitutively-active GαqRC (69), before the mRNA was extracted. For these measurements, approximately 50µL of extracted mRNA in RNase free water was added in a Hellma Fluorescence Quartz Cuvette (QS-3.00mm). Each sample was run 3 times, 10 minutes per run. Control samples were repeated 6 times, PLCβ knockdown twice, and Gαq over-expression once.

### Particle analysis

Samples were imaged using a 100X/1.49 oil TIRF objective to microscopically count the number of particles formed under different conditions per µm^2^. For each condition, 10-20 cells were randomly selected and z-stack measurements were taken (1.0 µ/frame). Analysis was performed using ImageJ and Fiji ImageJ in two ways. First, each measurement was thresholded before analyzing and the number of particles per frame per measurement was averaged. Second, all z-stack measurements were combined to create a 3D image per sample before analyzing the number of particles per and averaging the results. Both methods produced identical results.

### Microinjection studies

Microinjection of a solution of 100 nM eIF5A into PC12 cells was carried out using an Eppendorf Femtojet Microinjector mounted on Nikon Eclipse Ti-U inverted confocal microscope under 0.35 PSI pressure and 0.5 Seconds injection time per injection.

### Immunofluorescence

Cells were samples were fixed with 3.7% formaldehyde and permeabilized using 0.1% triton X-100 in PBS then blocked using 10% goat serum, 5% BSA, 50mM glycine in PBS. Cells were then stained with primary antibodies from (Abcam), incubated for 1 hour, washed and treated with a secondary antibody for 1 hour. After another wash, the cells were viewed on either a Zeiss Meta 510 laser confocal microscope. Data were analyzed using Zeiss LSM software and Image J. The secondary antibodies used were Anti-rabbit Alexa-488 for Ago-2 and Anti-mouse Alexa 647 for PABPC1.

### Calcium signal imaging

Single cell calcium measurements were carried out by labeling the cells with Calcium Green (Thermofisher), incubating in HBSS for 45 minutes and washing twice with HBSS. Release of intracellular calcium in live PC12 cells was initiated by the addition of 2 µM carbachol before imaging the time seirs on a Zeiss LSM 510 confocal microscope excitation at 488 using time series mode as previously described, see (70).

### Western blotting

Samples were placed in 6 well plates and collected in 250μL of lysis buffer that included NP-40 and protease inhibitors as mentioned before, sample buffer is added at 20% of the total volume. After SDS-PAGE electrophoresis, protein bands were transfer to nitrocellulose membrane (Bio-Rad, California USA). Primary antibodies include anti-PLCβ1 (Santa Cruz sc-5291), anti-Ago2 (abcam ab32381), anti-PABPC1 (Santa Cruz sc-32318), anti-G3BP1 (Santa Cruz sc-81940), anti-actin (abcam ab8226) and anti-eGFP (Santa Cruz sc-8334). Membranes were treated with antibodies diluted 1:1000 in 0.5% milk, washed 3 times for 5 minutes, before applying secondary antibiotic (anti-mouse or anti-rabbit from Santa Cruz) at a concentration of 1:2000. Membranes were washed 3 times for 10 minutes before imaging on a BioRad chemi-doc imager to determine the band intensities. Bands were measured at several sensitivities and exposure times to insure the intensities were in a linear range. Data were analyzed using Image-J.

## ACKNOWLEDGEMENTS

The authors are grateful for the support of NIH GM116187. Dr. Garwain was supported by a Richard Whitcomb Fellowship. The authors also would like to thank Dr. Elizabeth Bafaro for her help in cloning mCherry-TRAX and to Dr. Siddartha Yerramilli for help with N&B and for his helpful comments throughout this work, Gabriella Fiorentino for her help gathering data and to an anonymous reviewer for his/her constructive comments.

## Supplemental Data

**Figure 1A.**
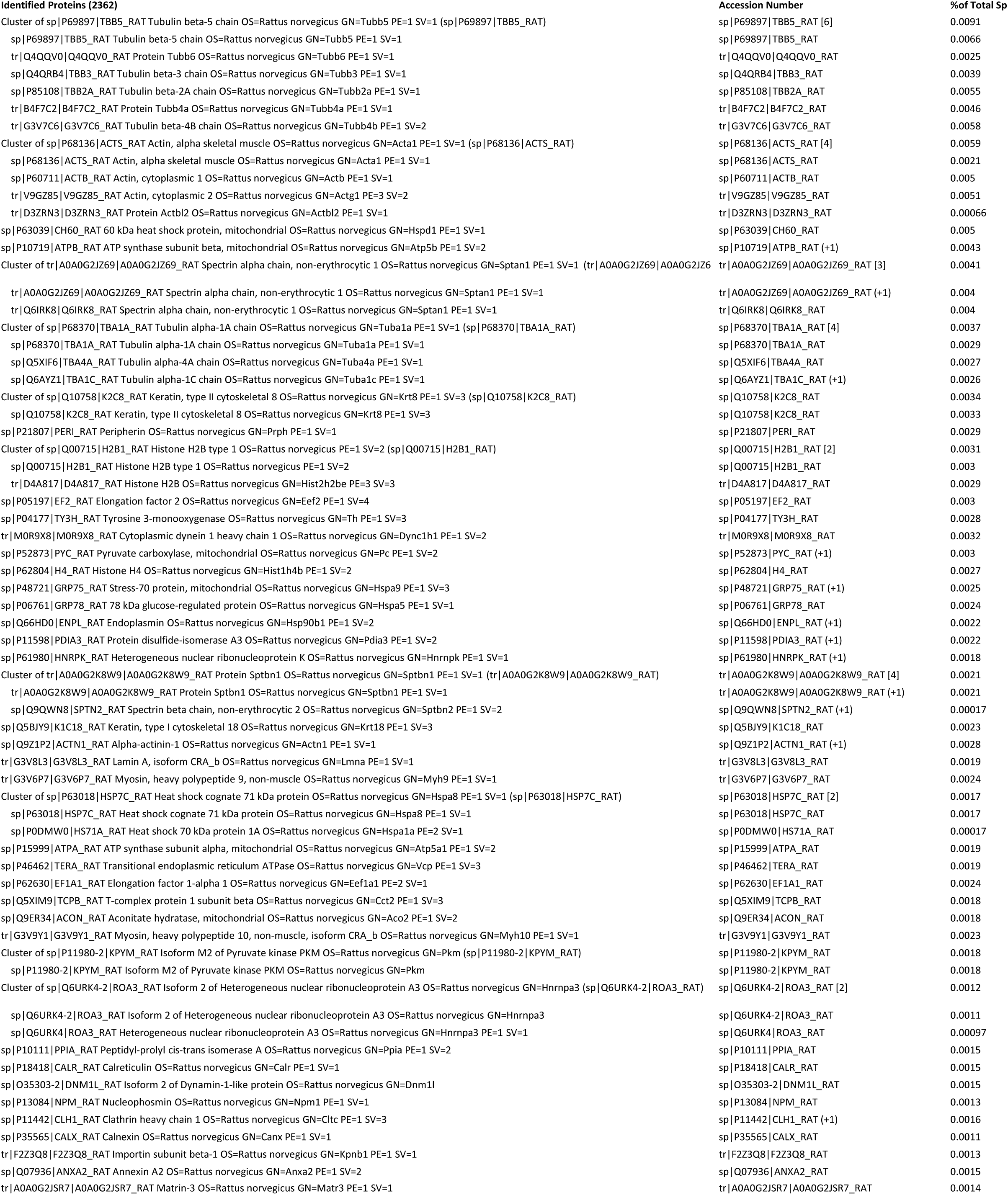
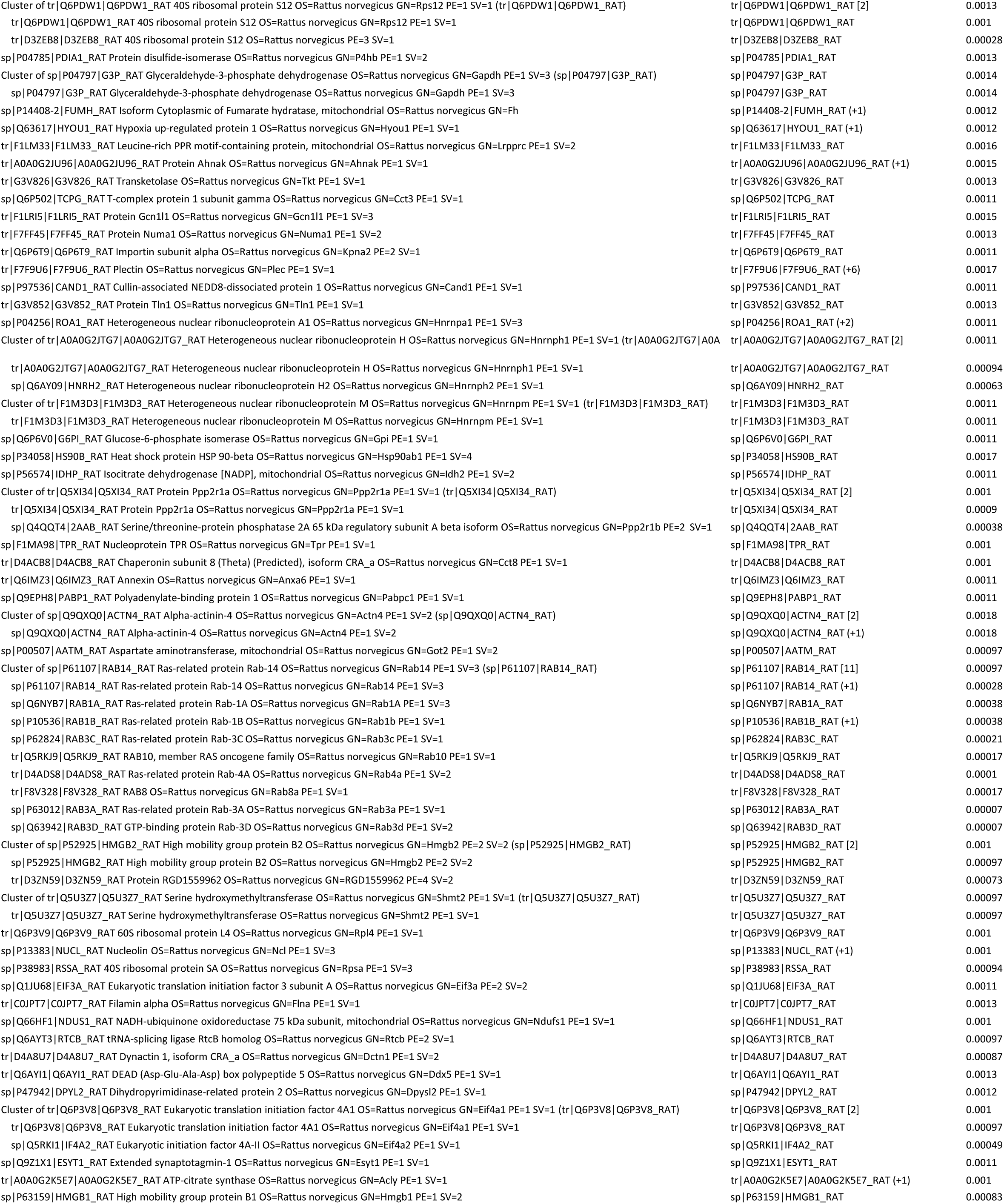
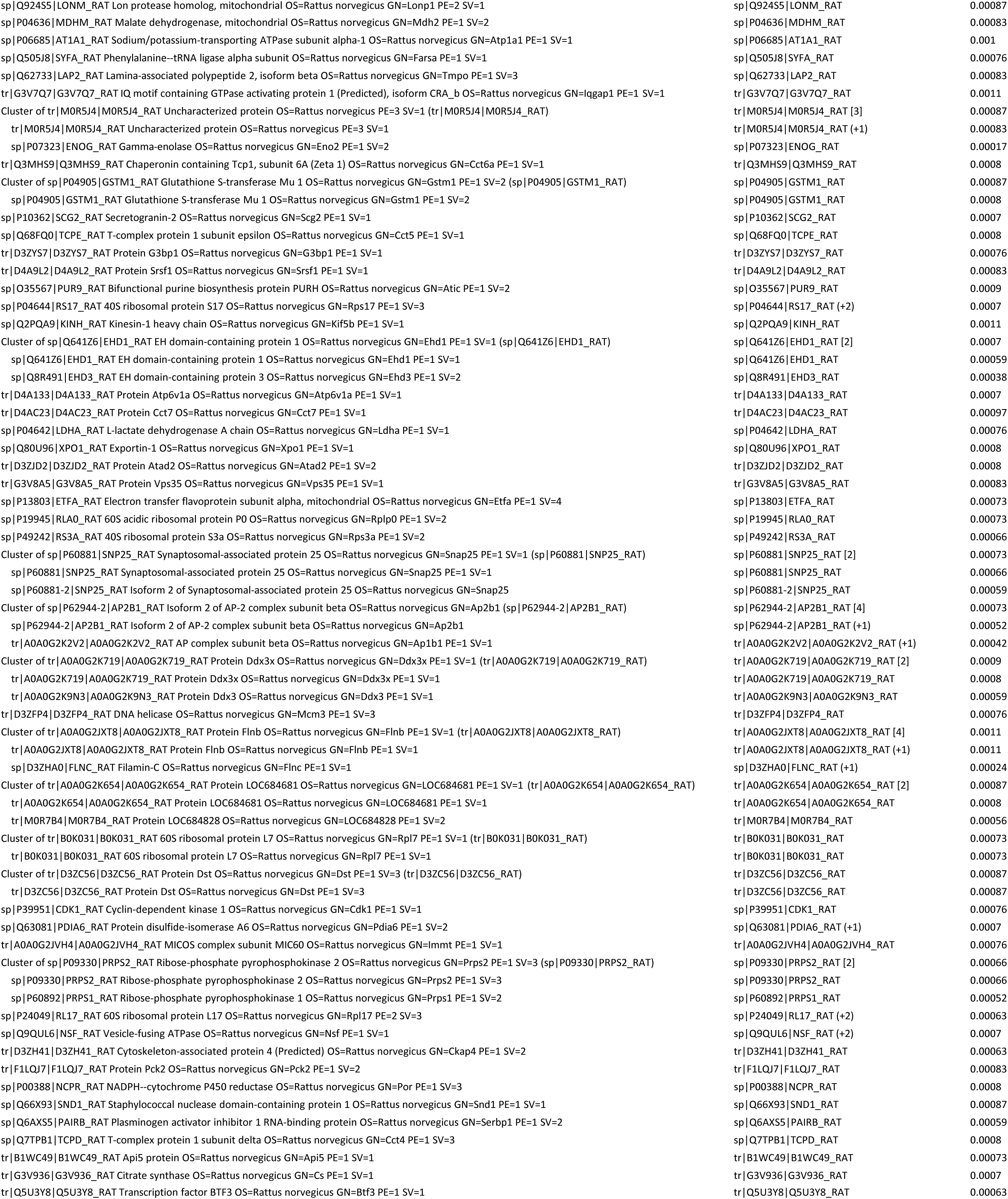
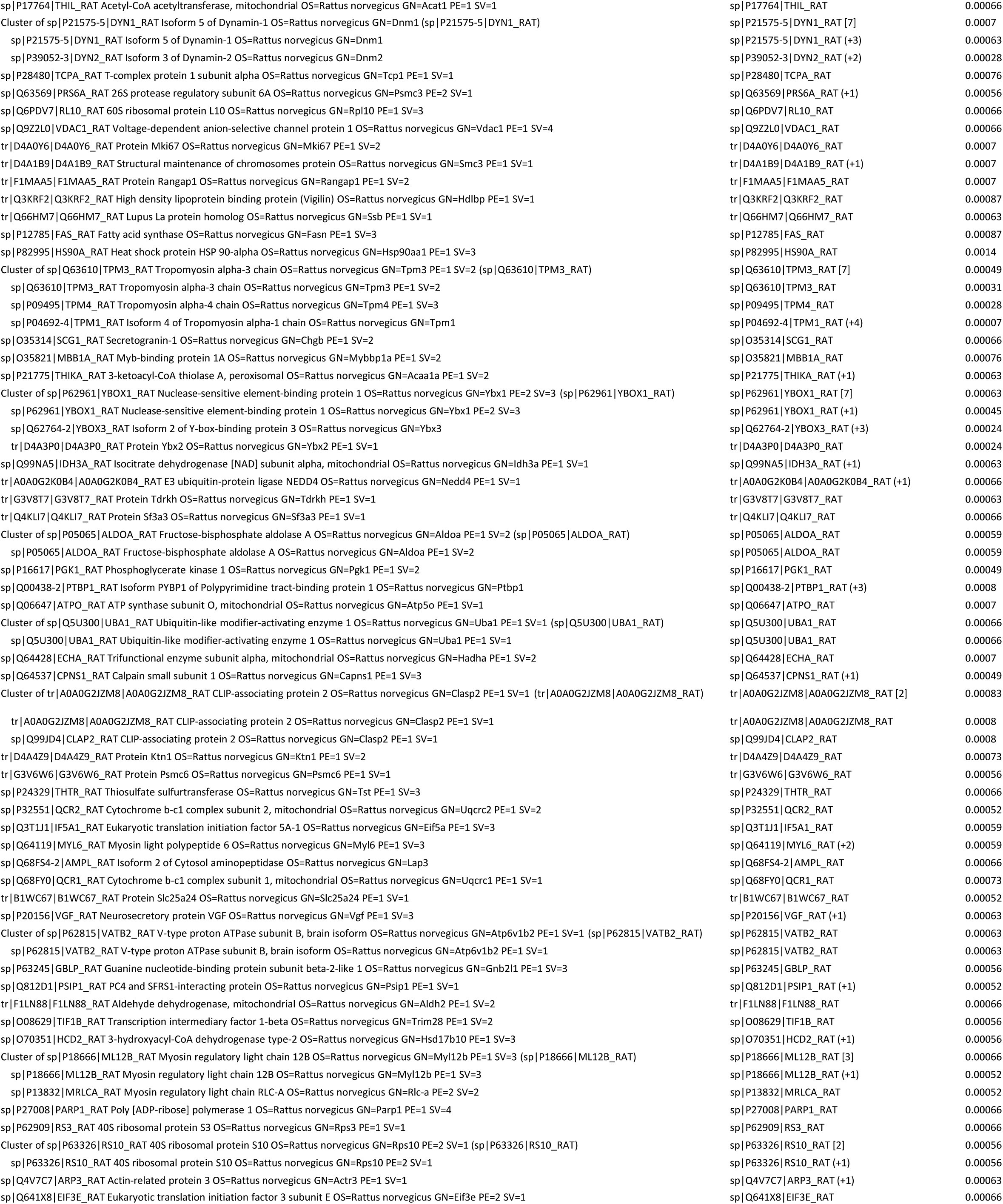
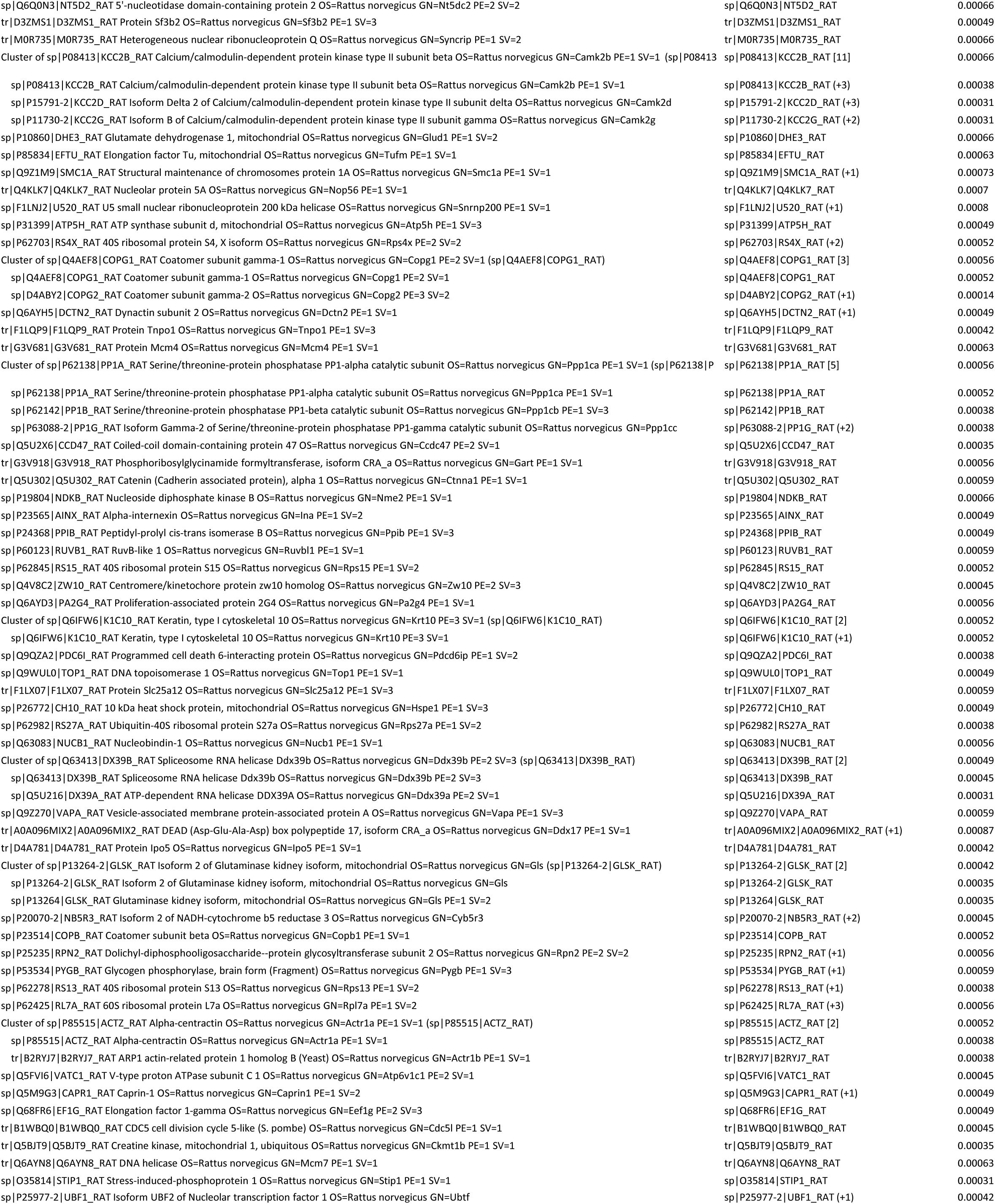
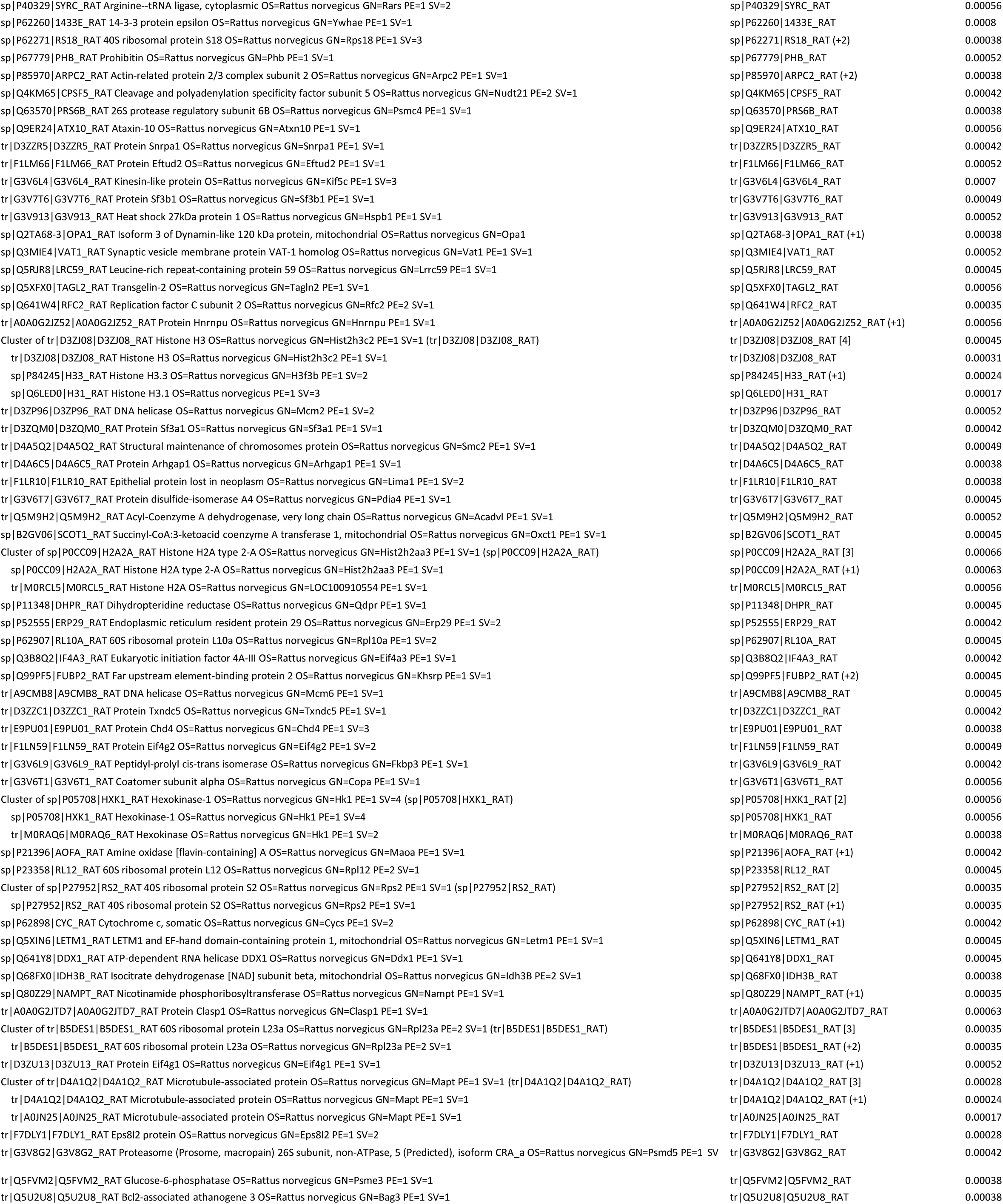
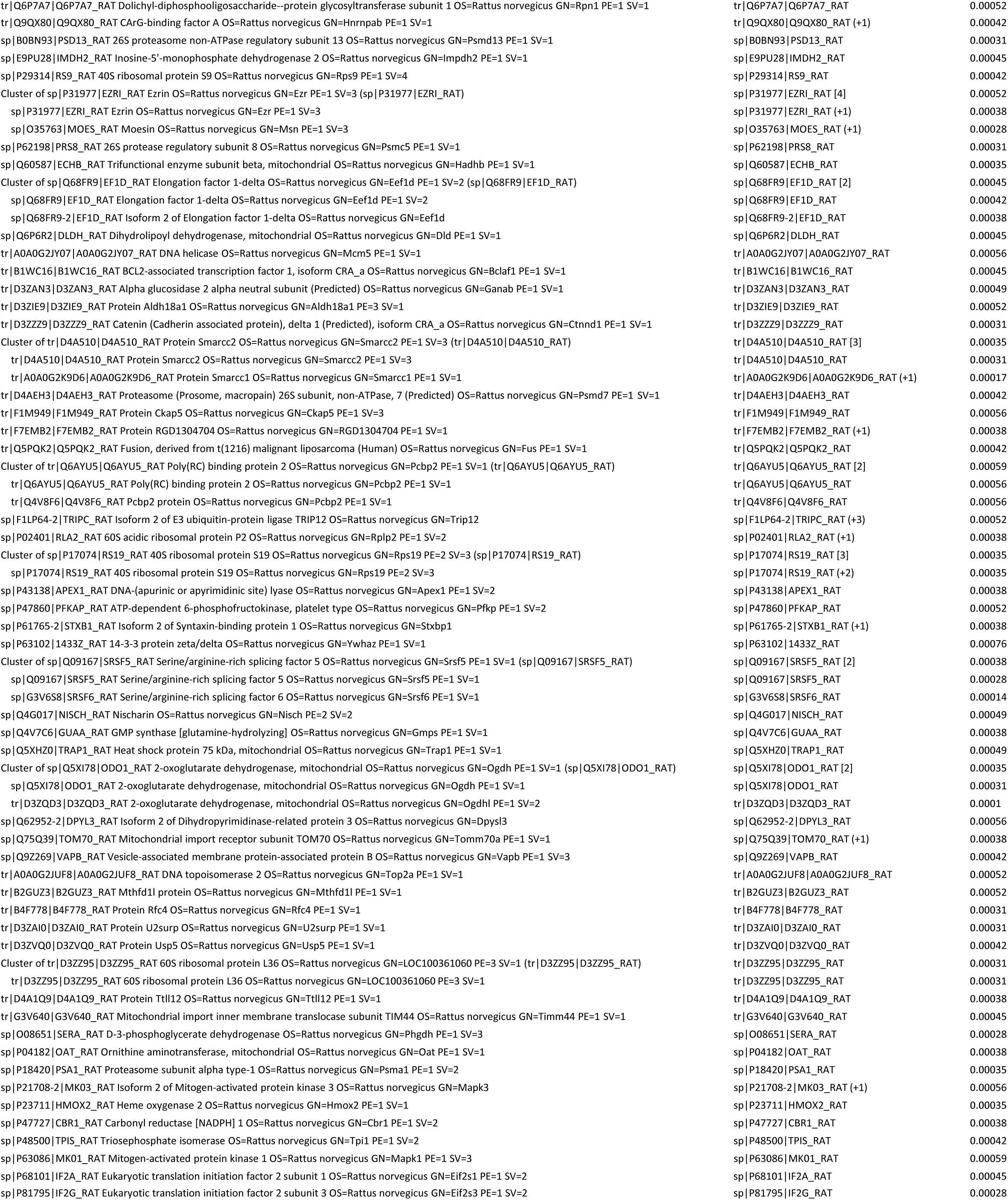
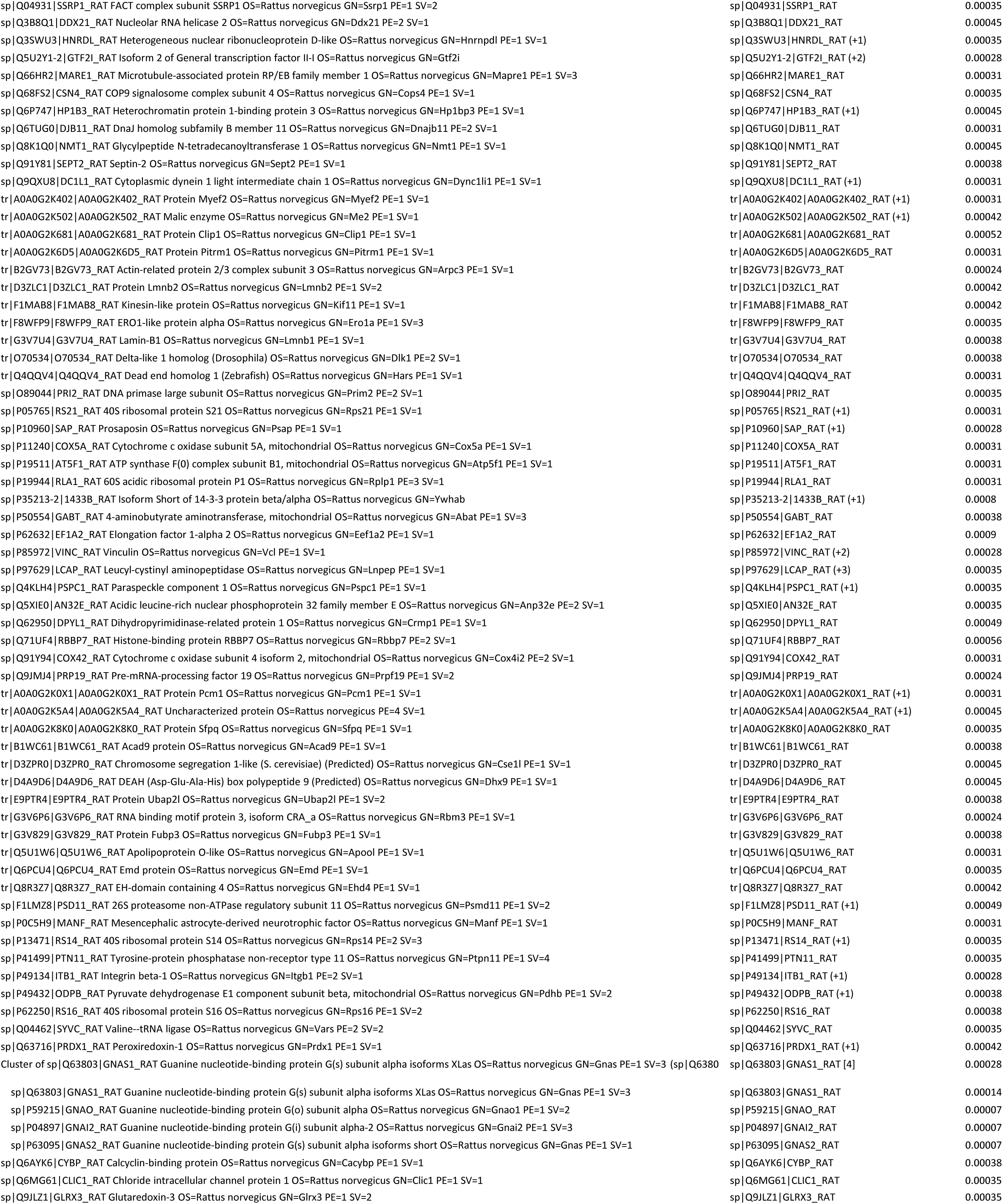
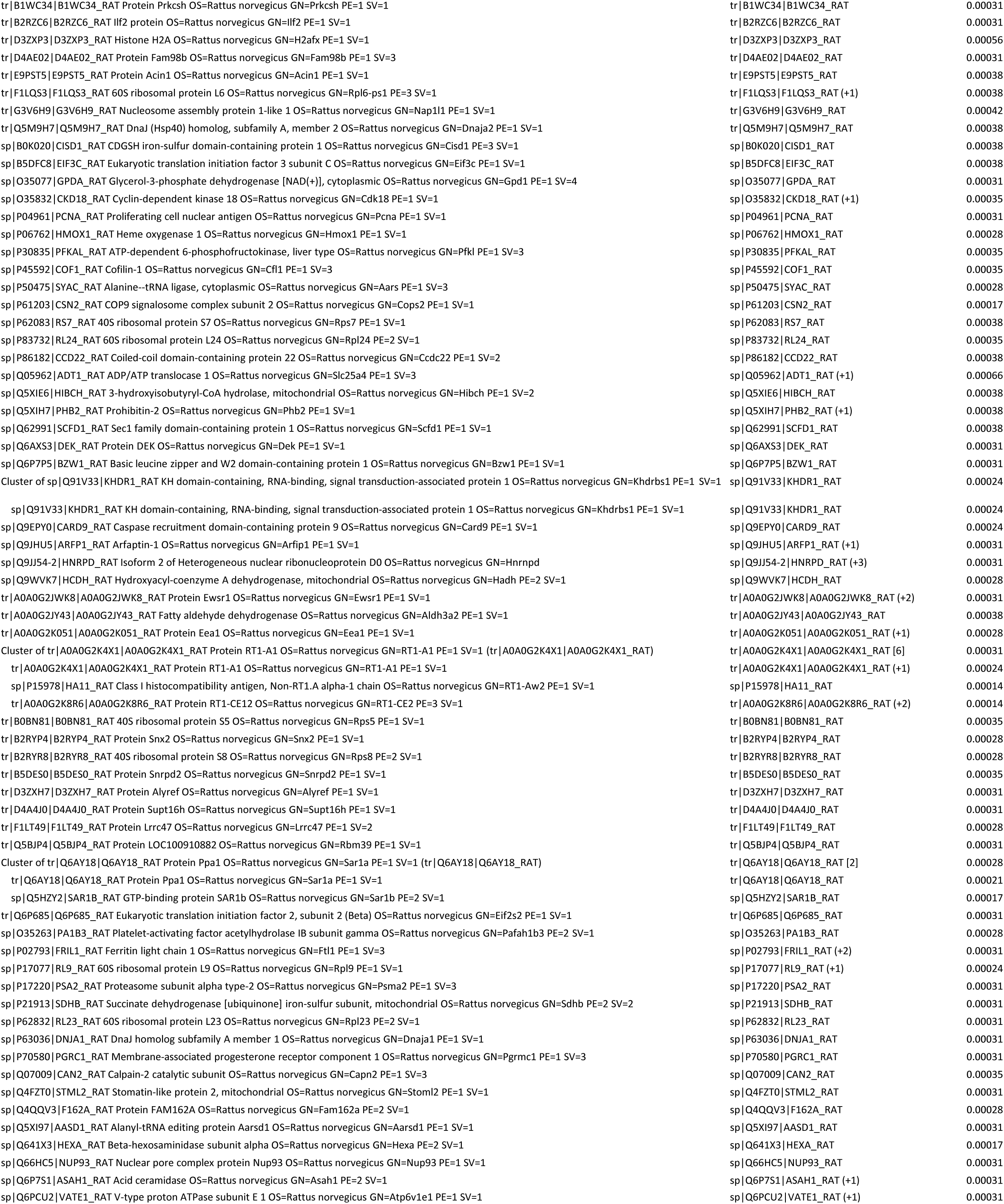
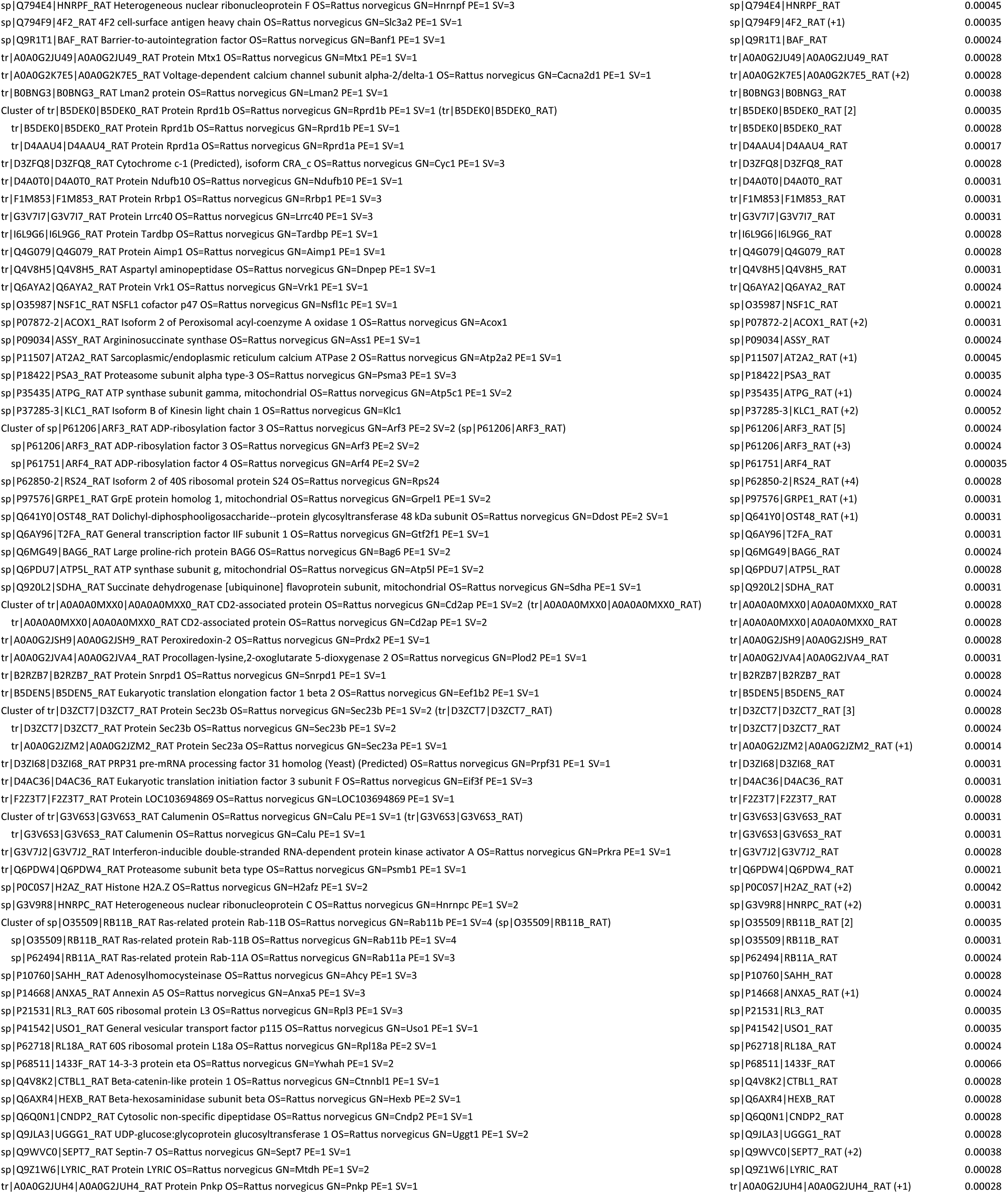
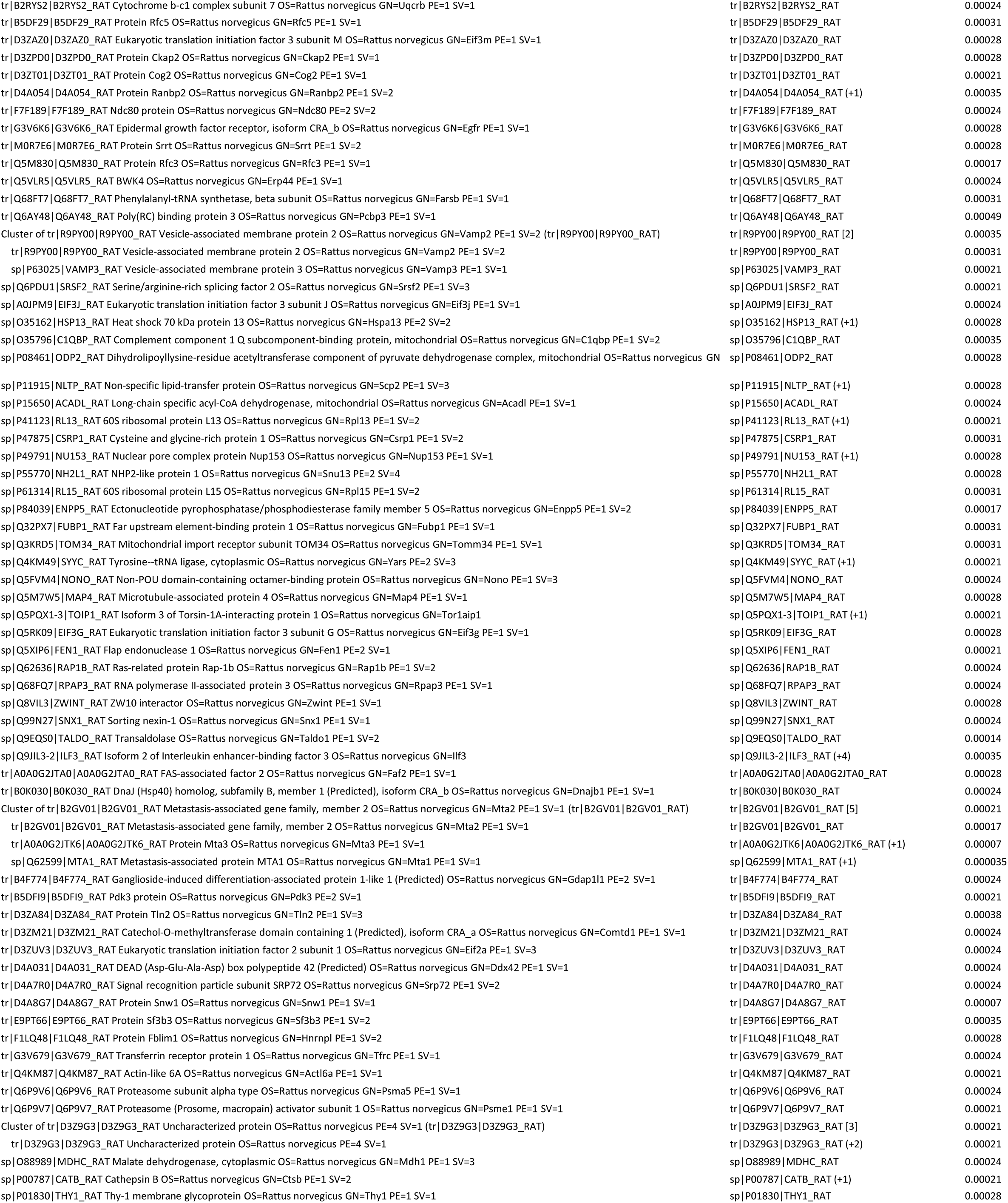
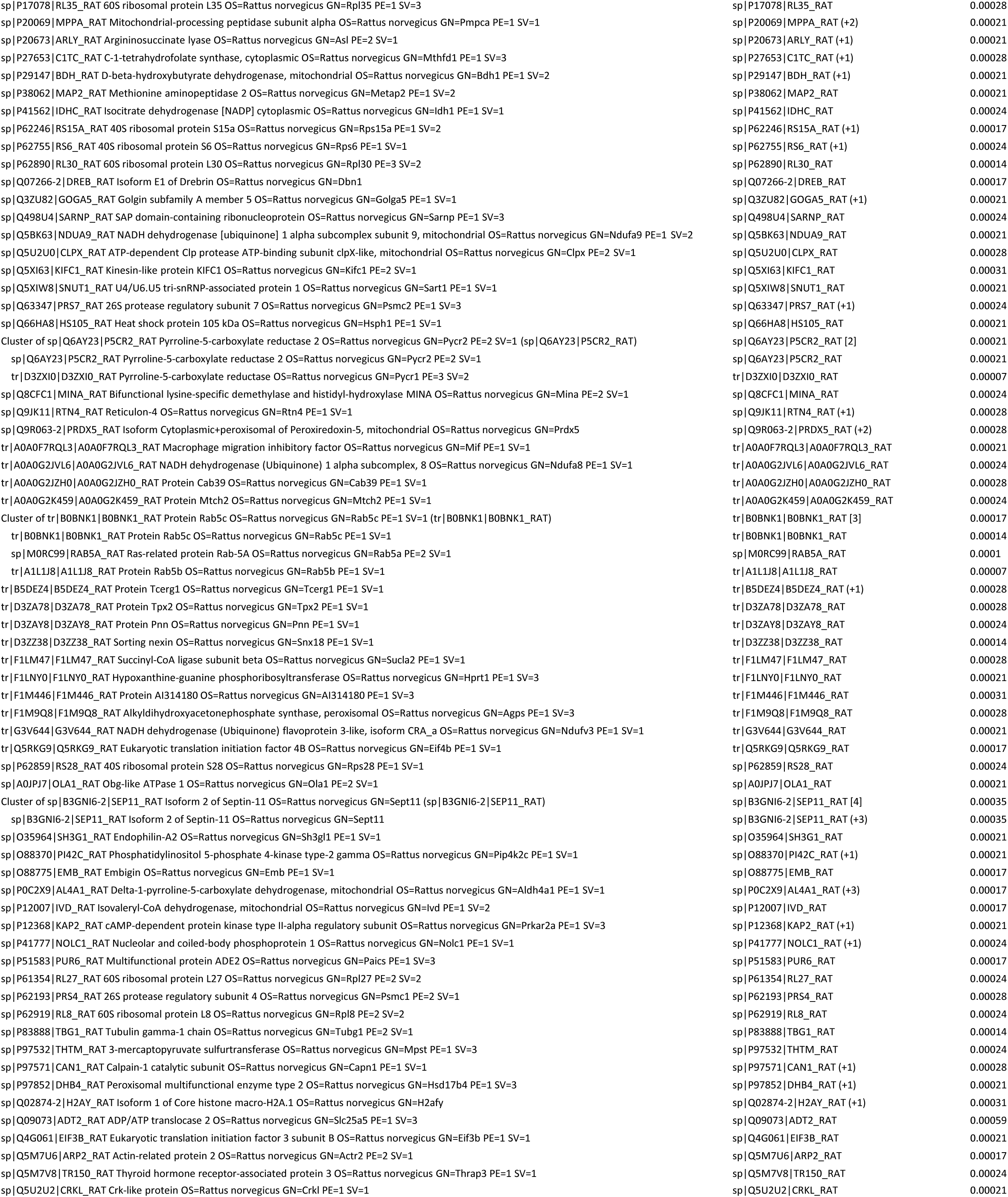
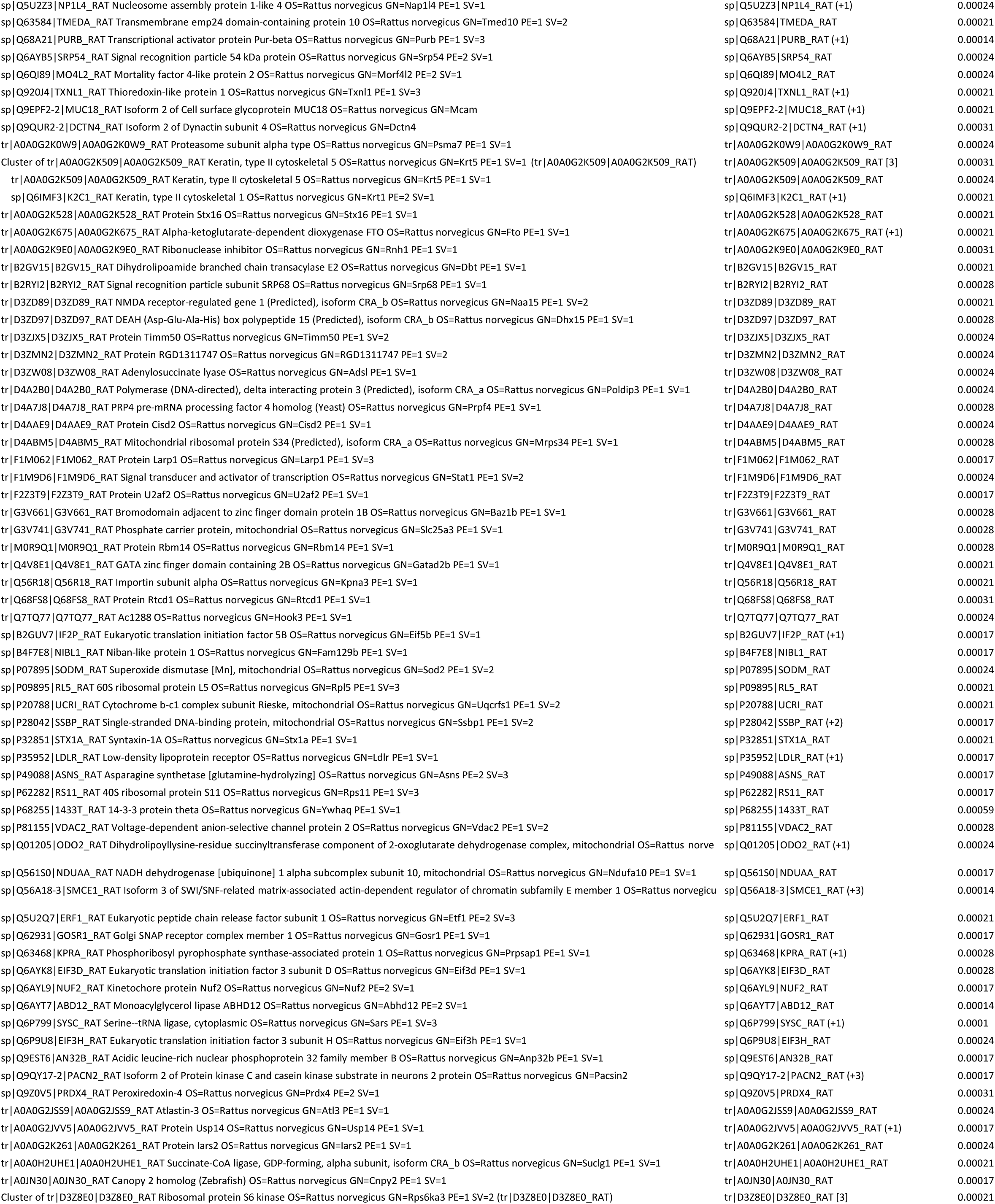
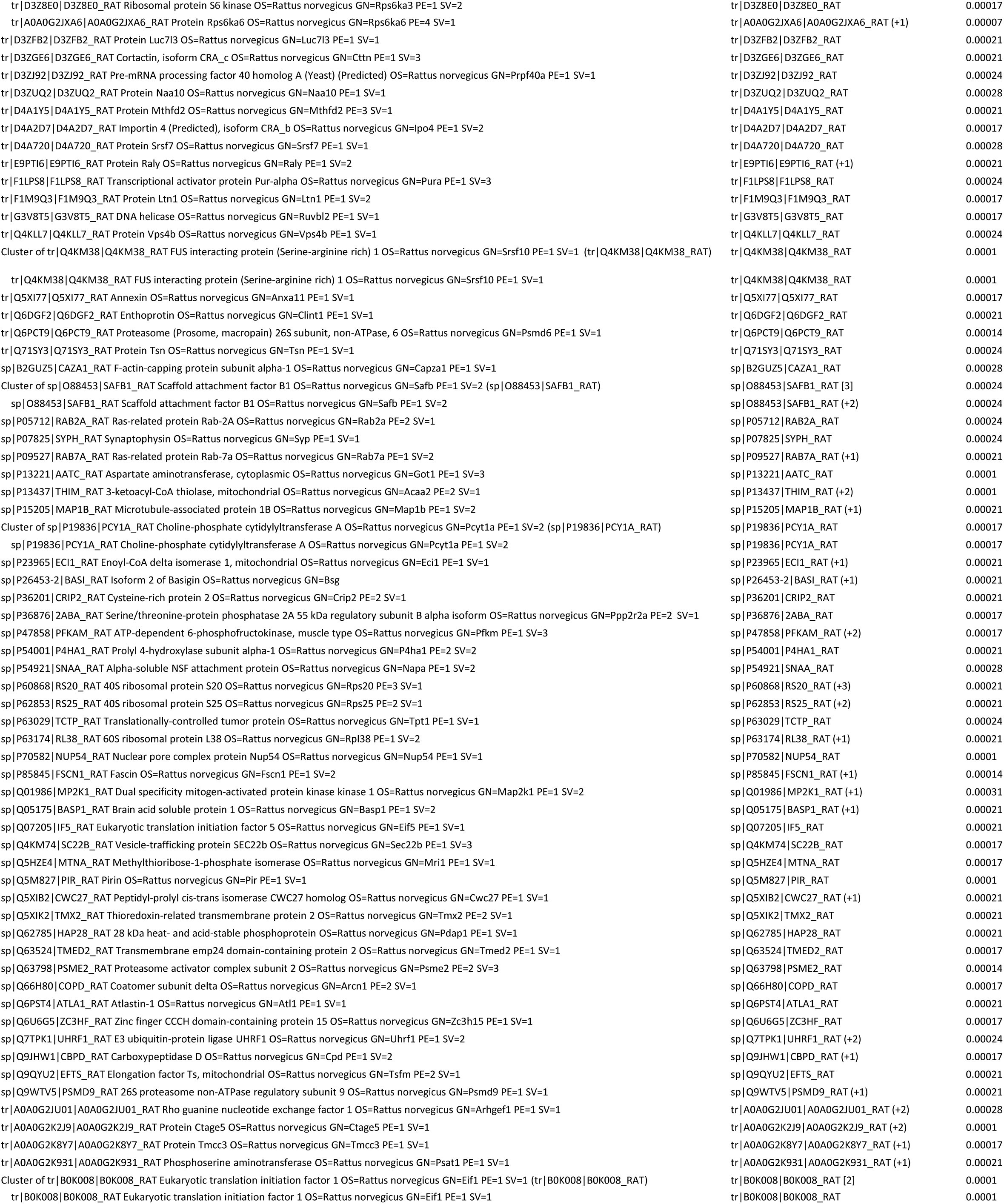
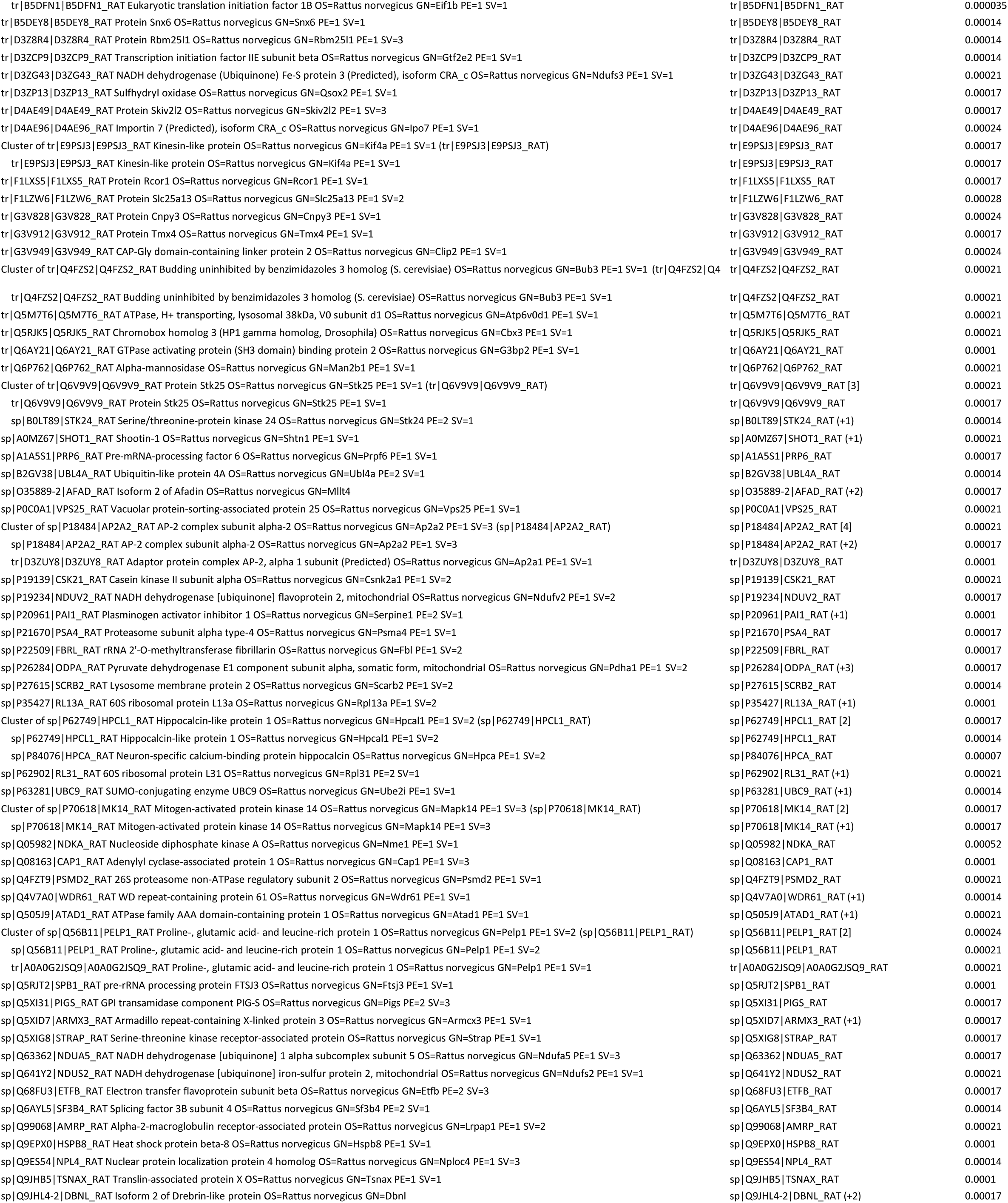
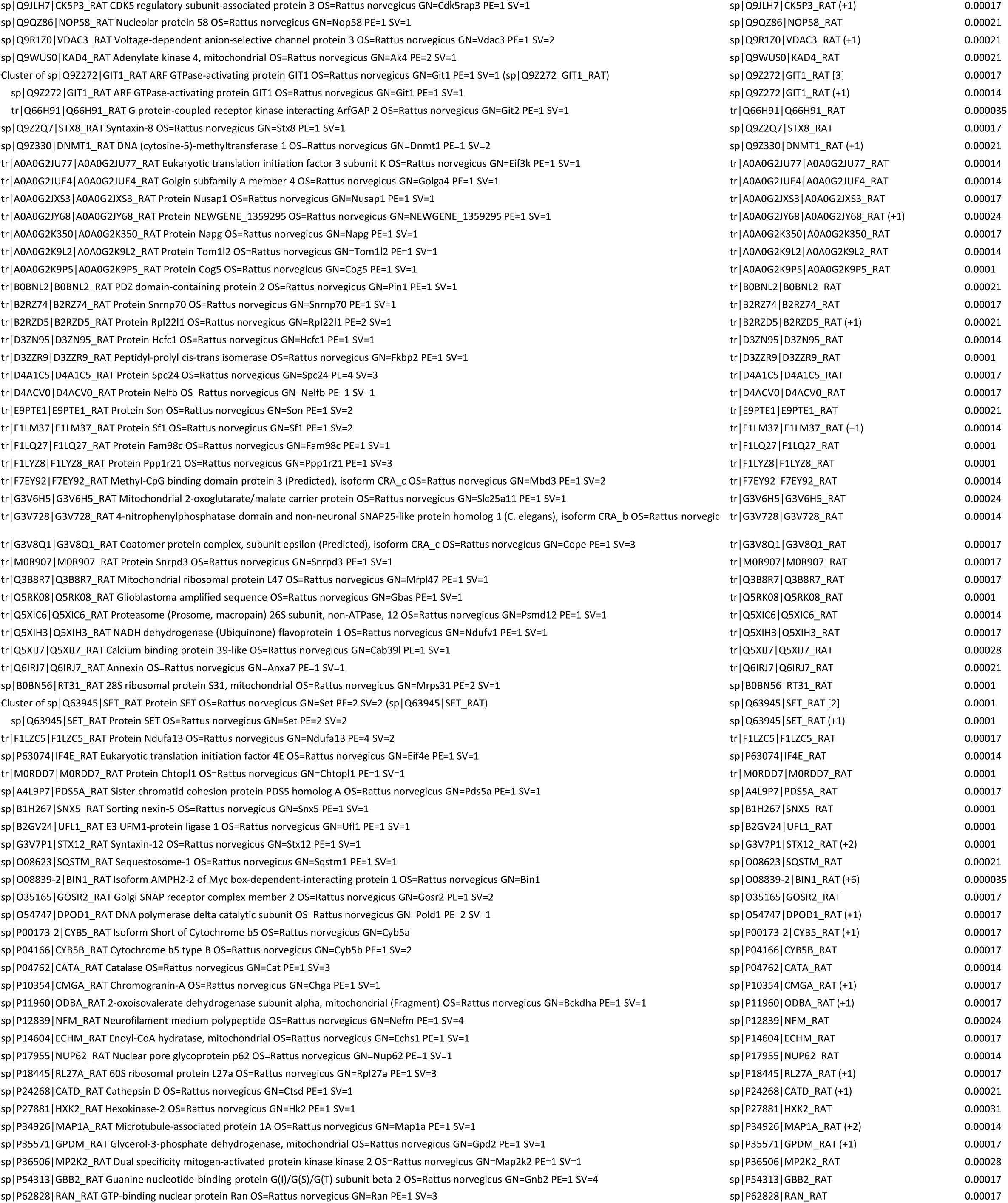
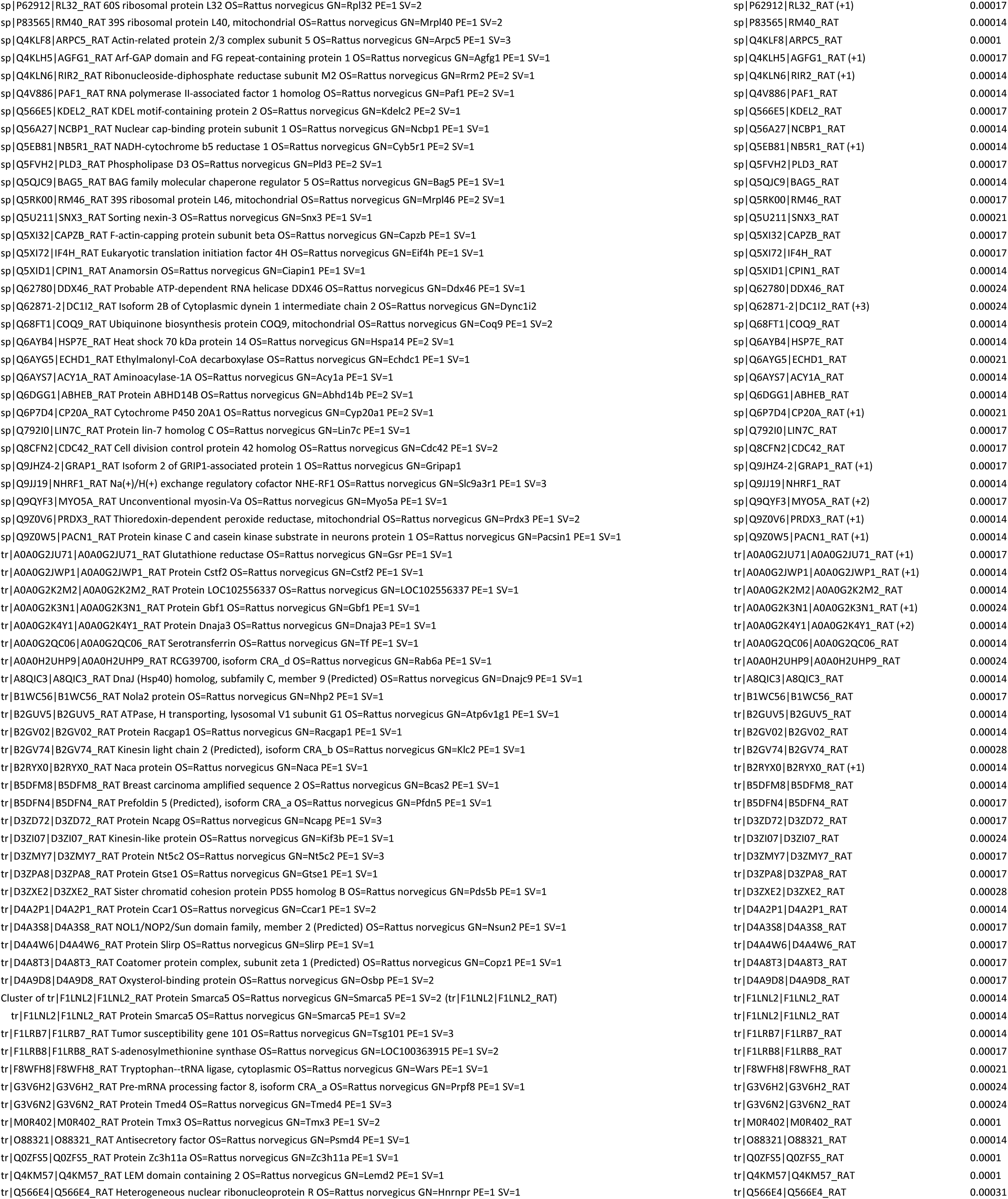
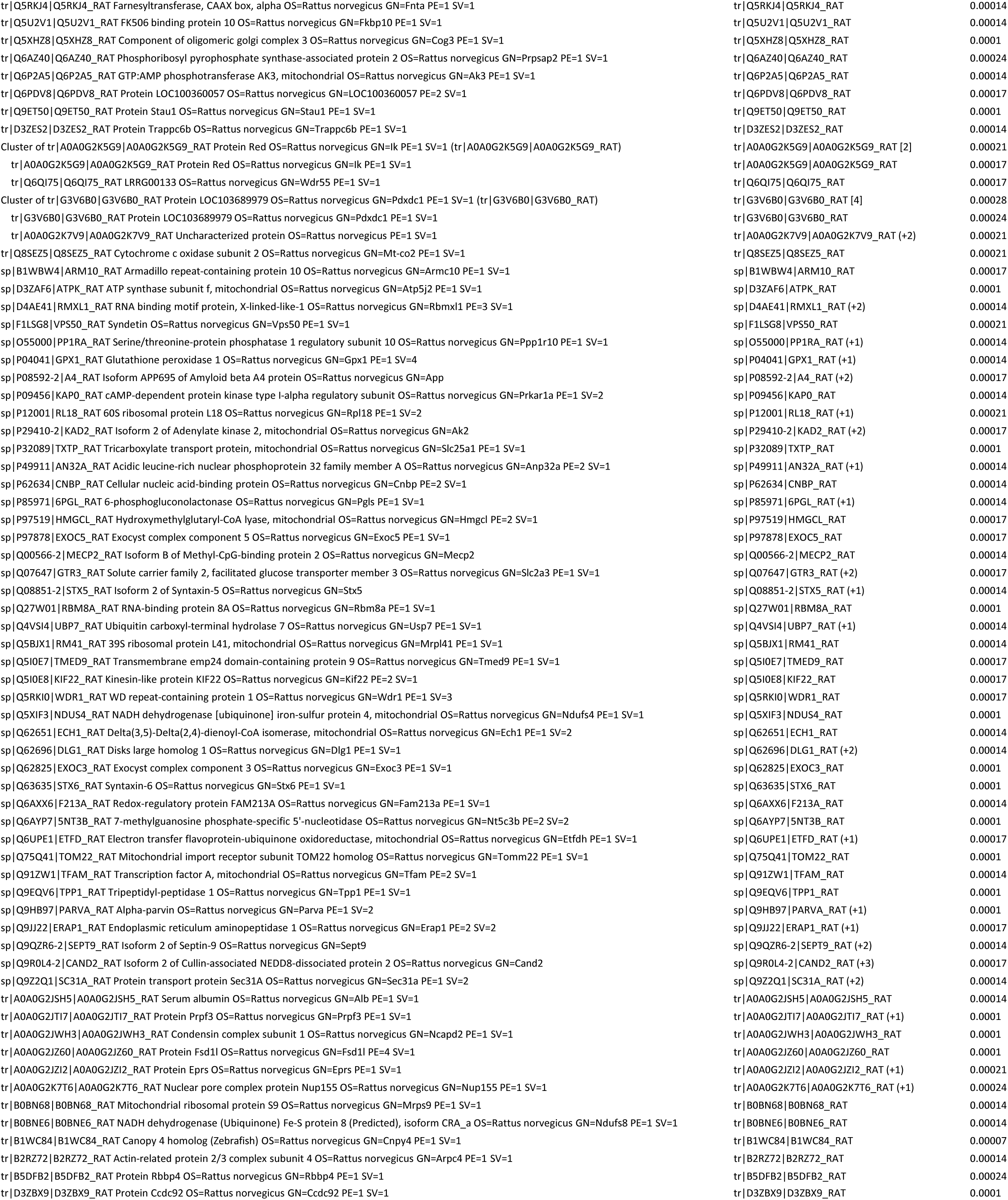
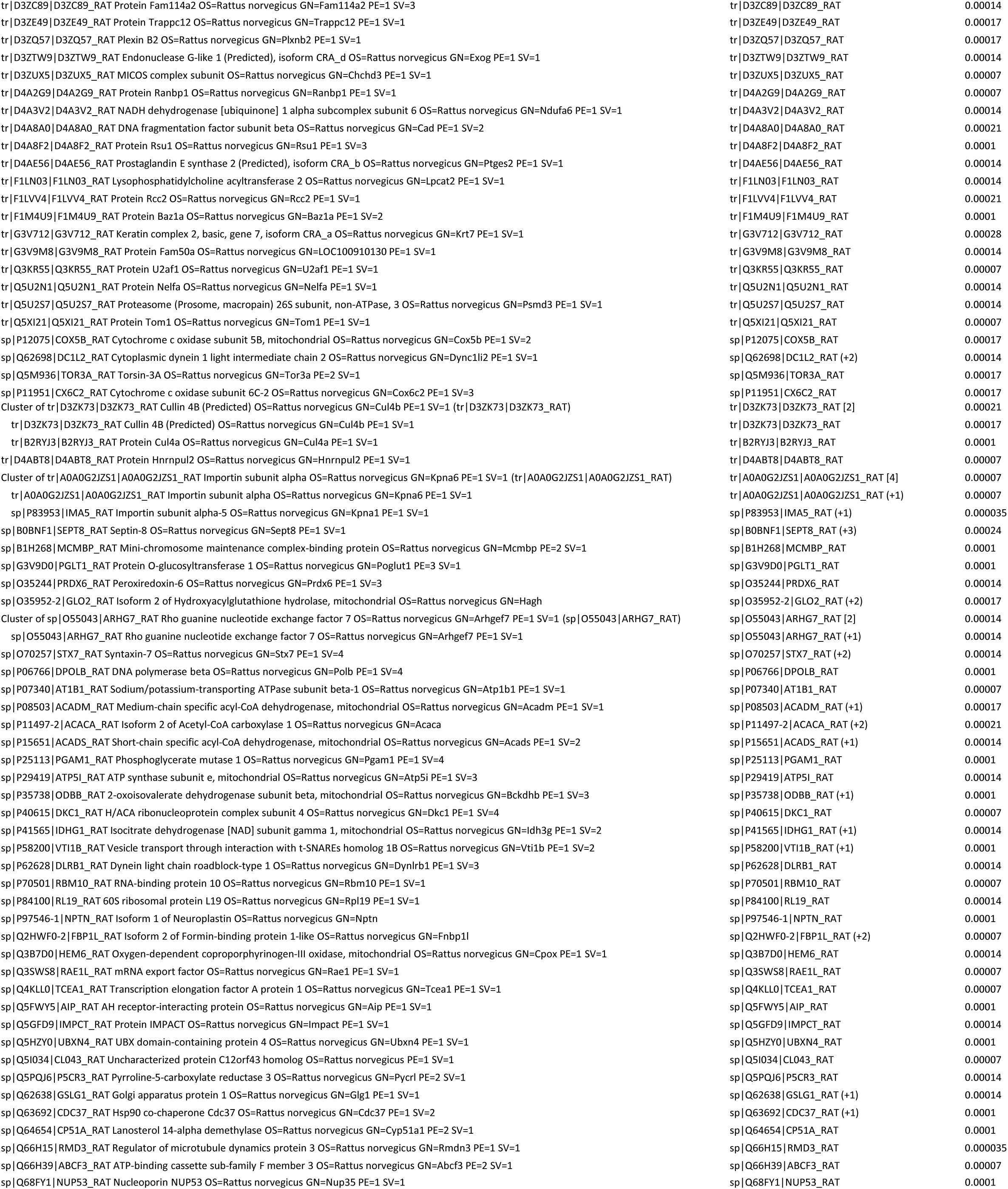
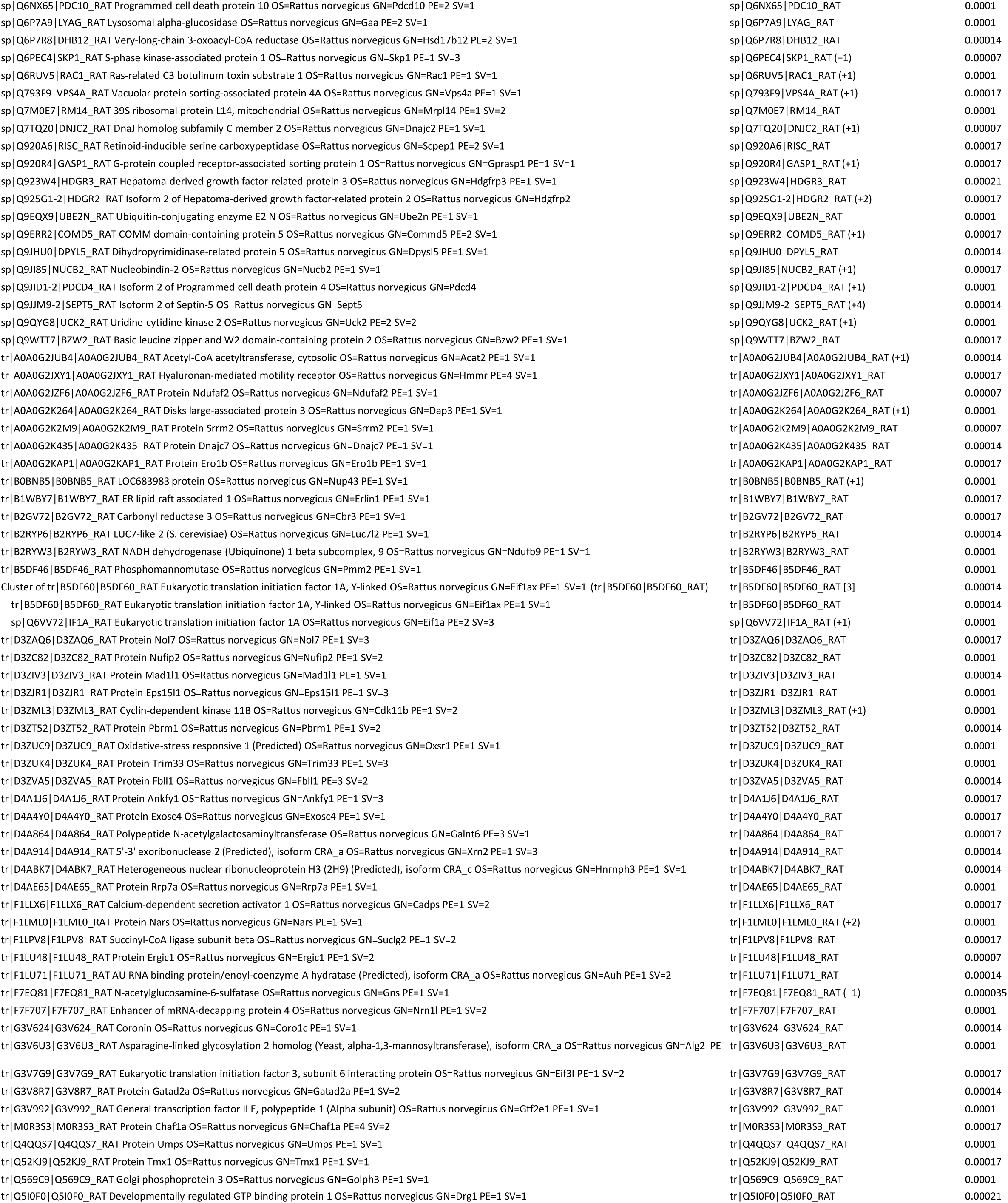
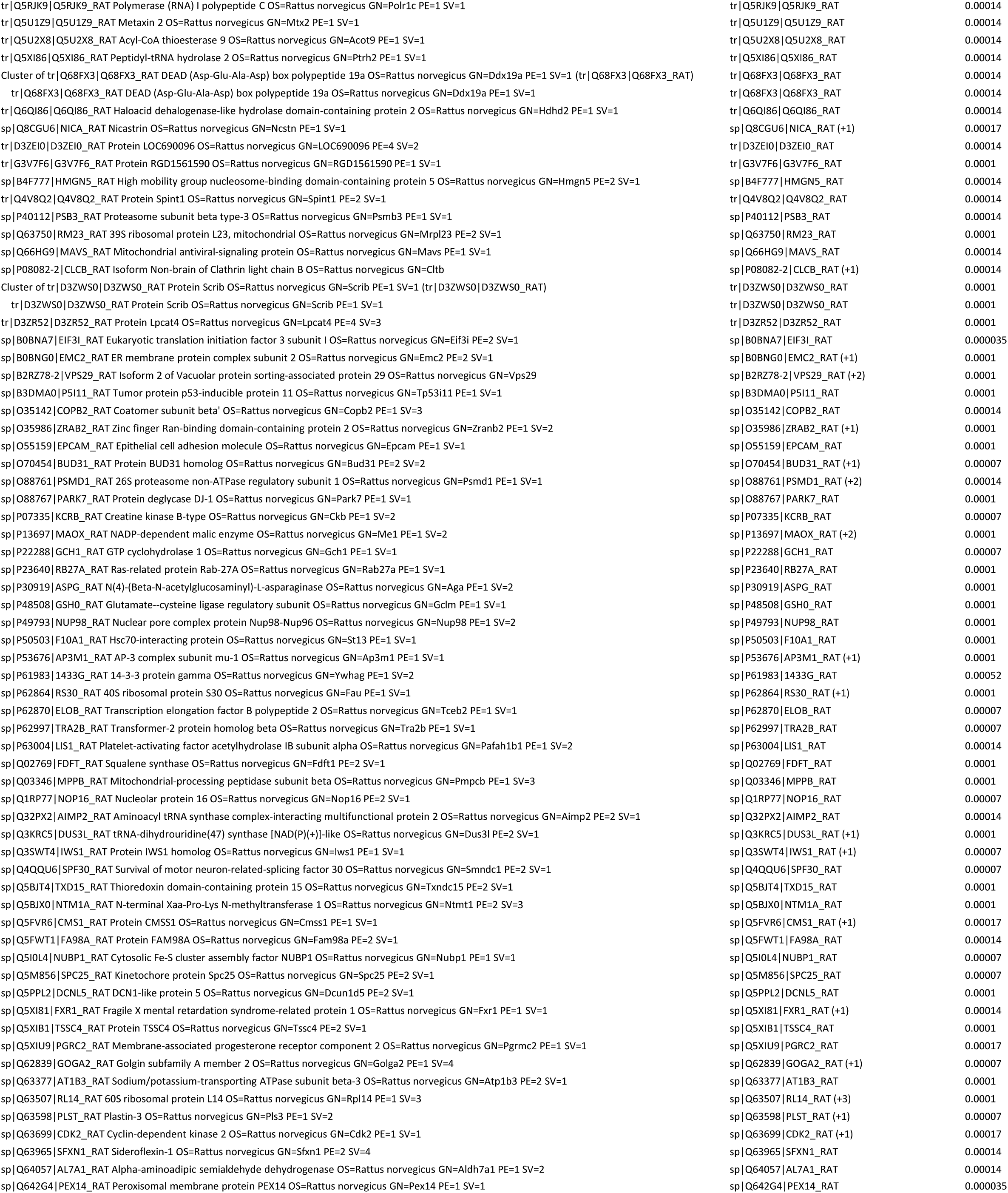
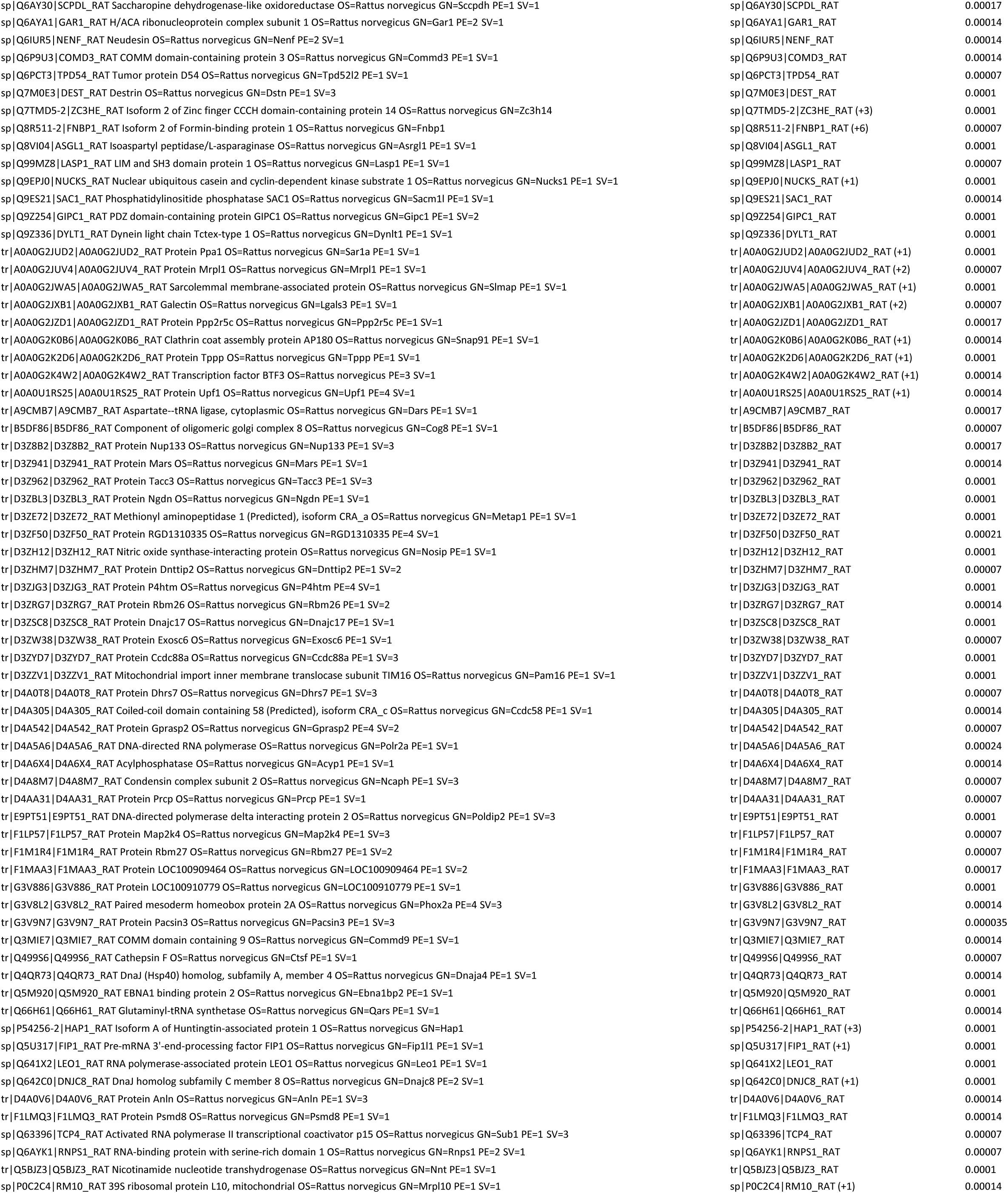
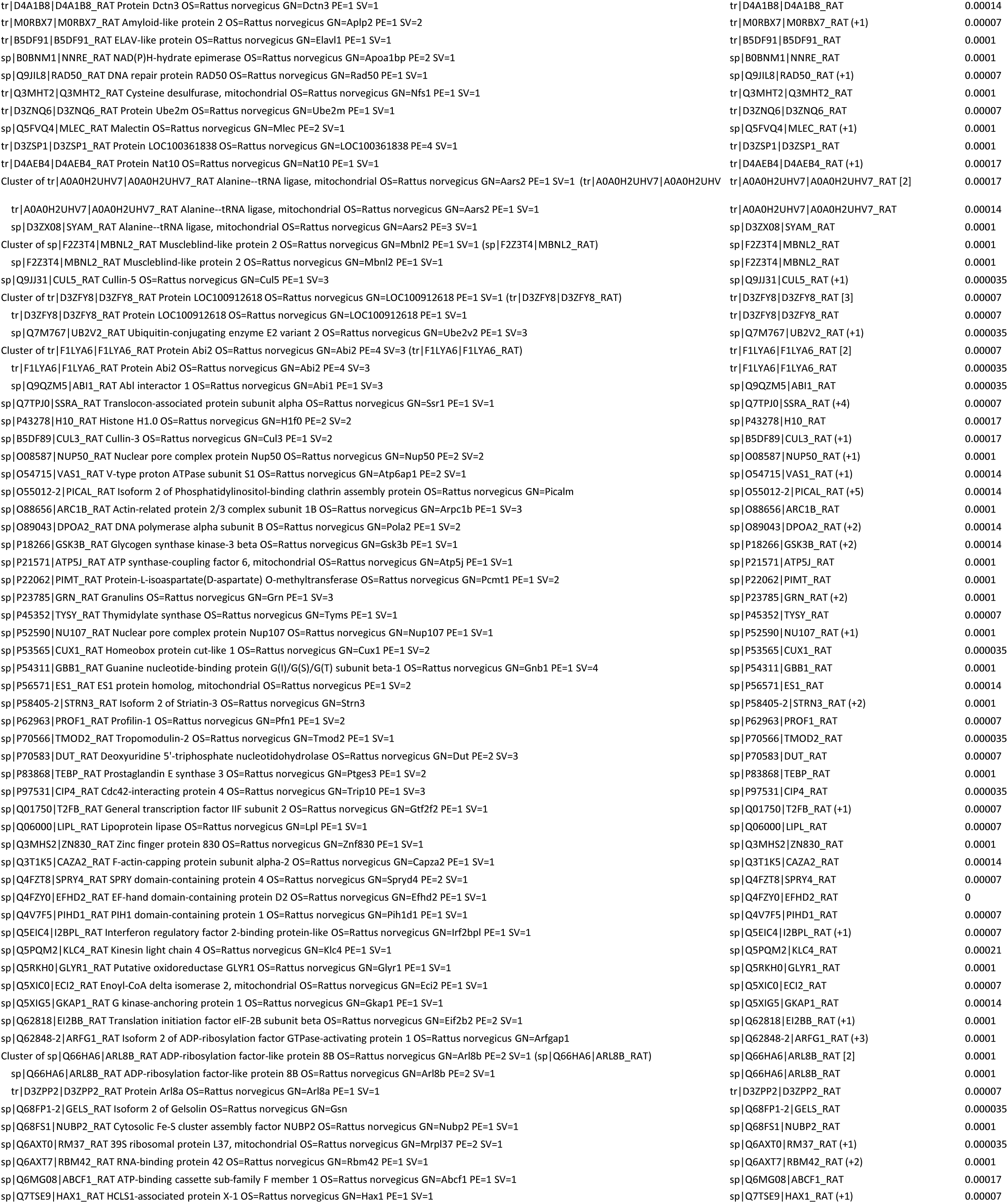
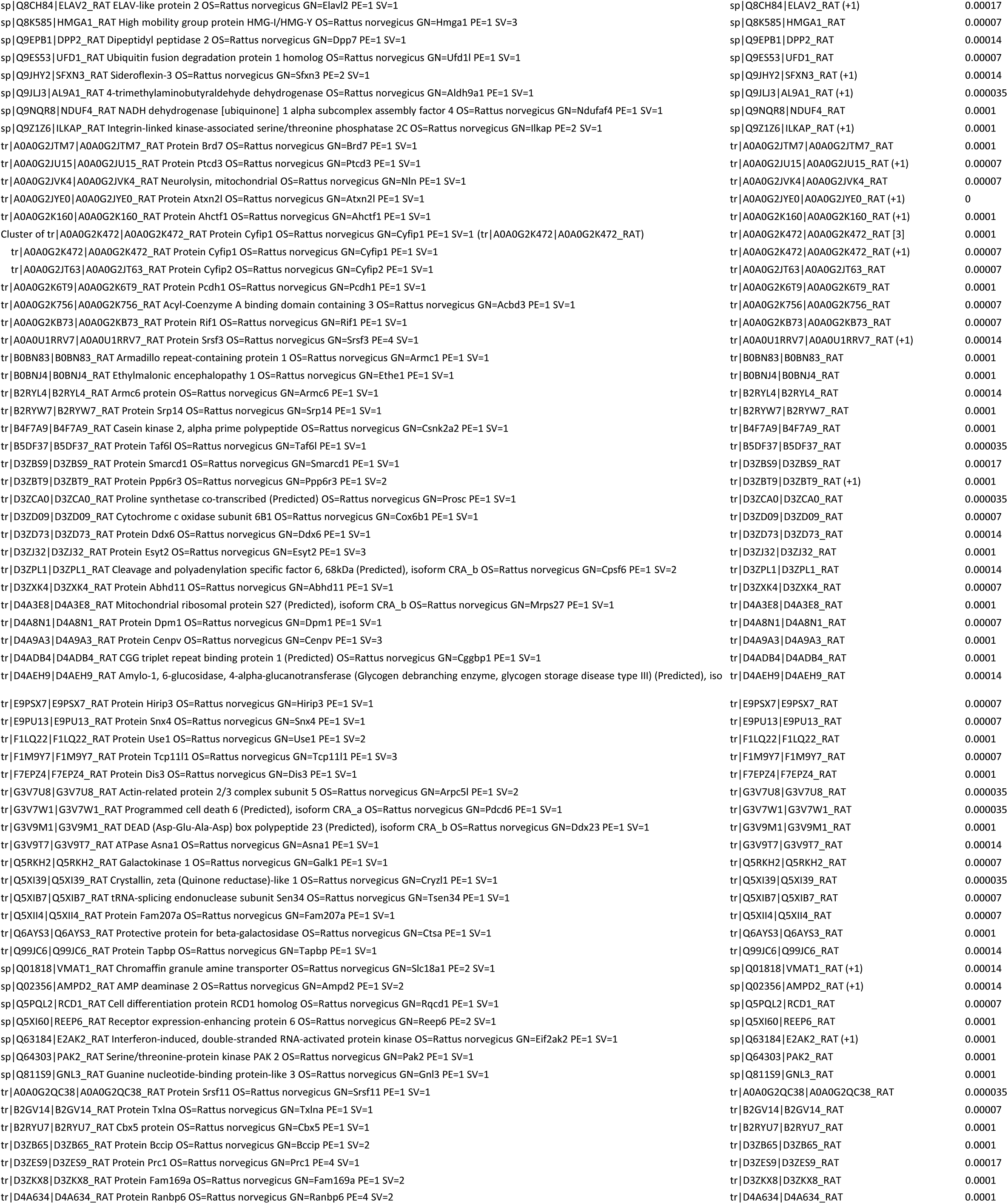
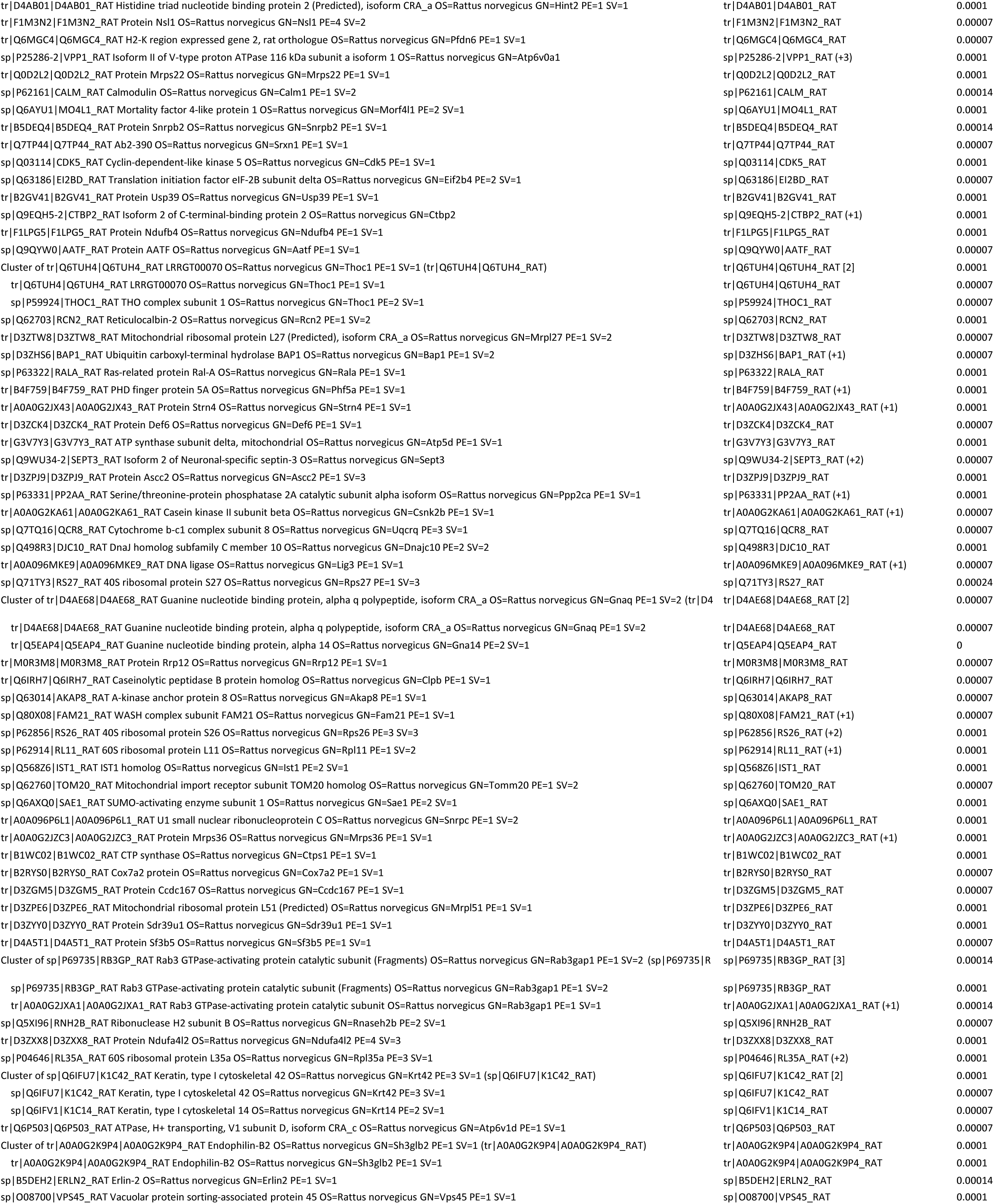
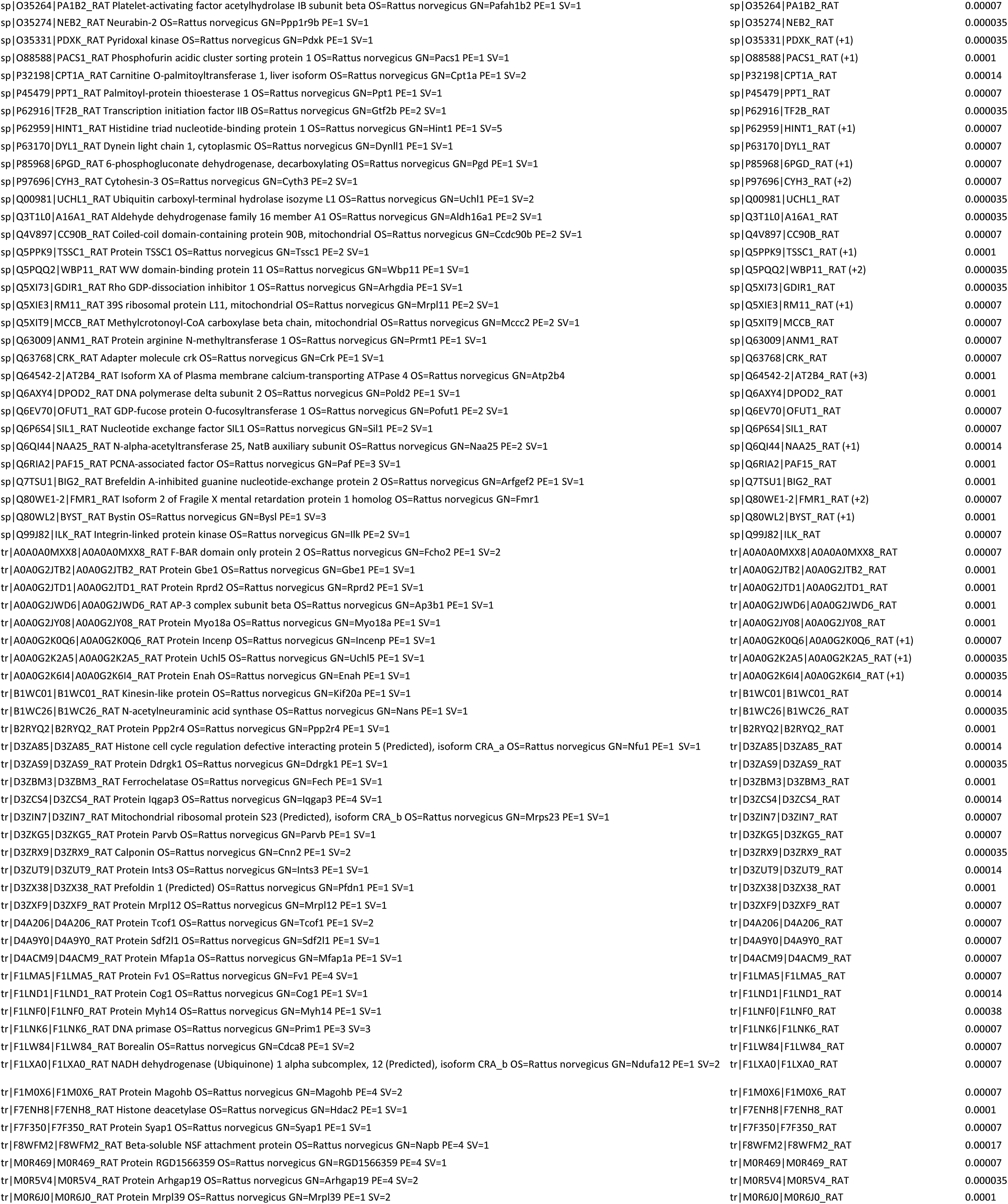
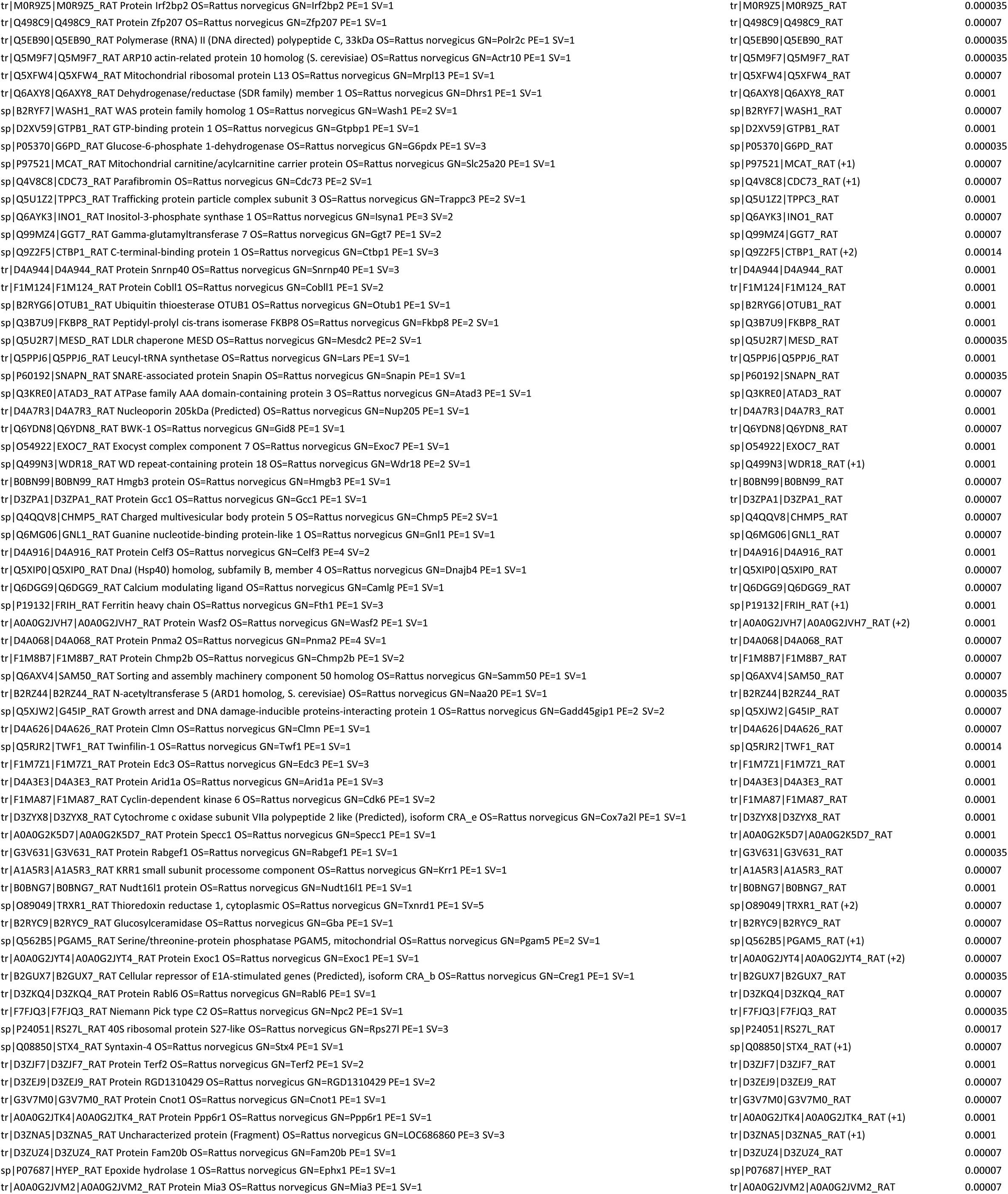
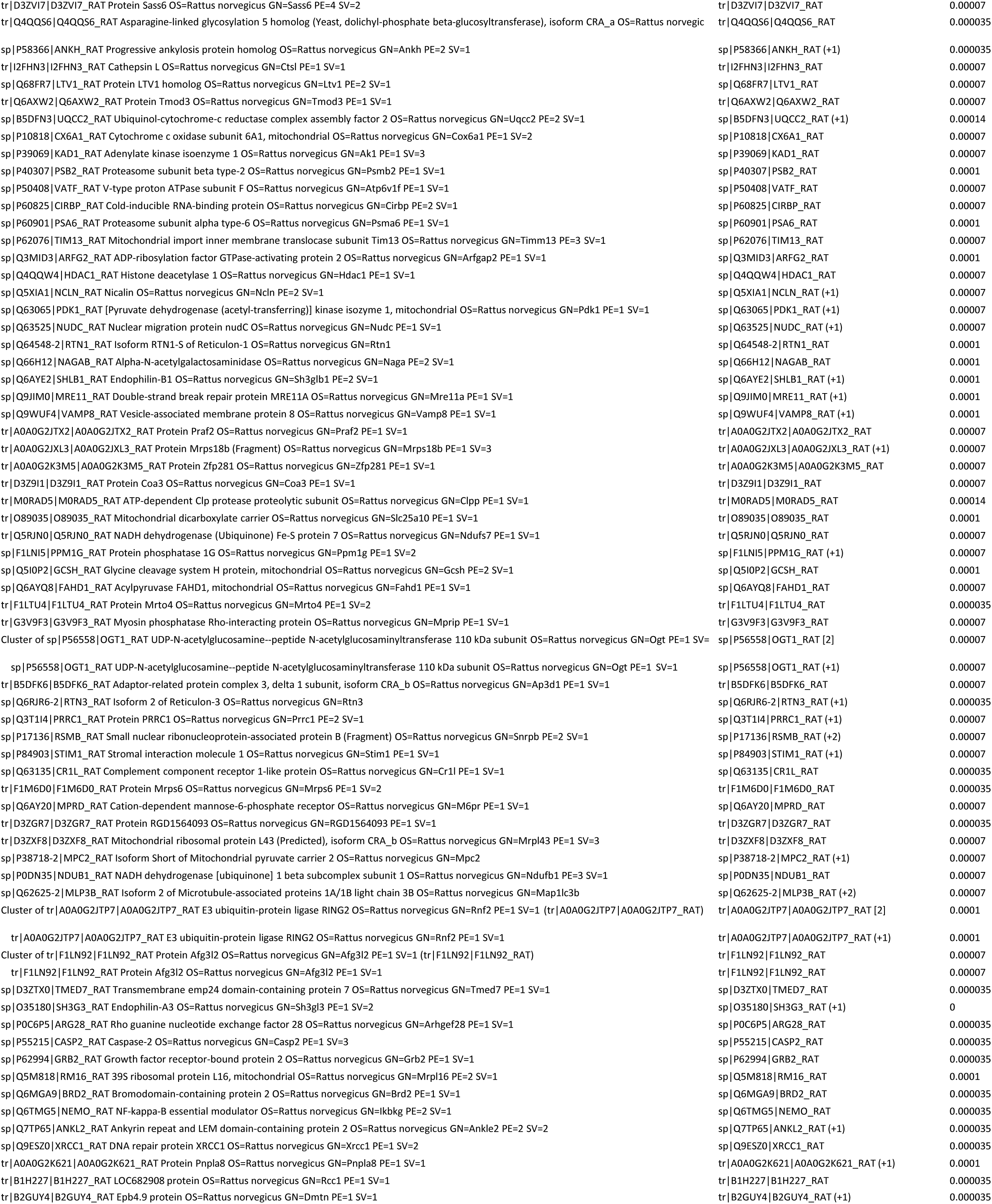
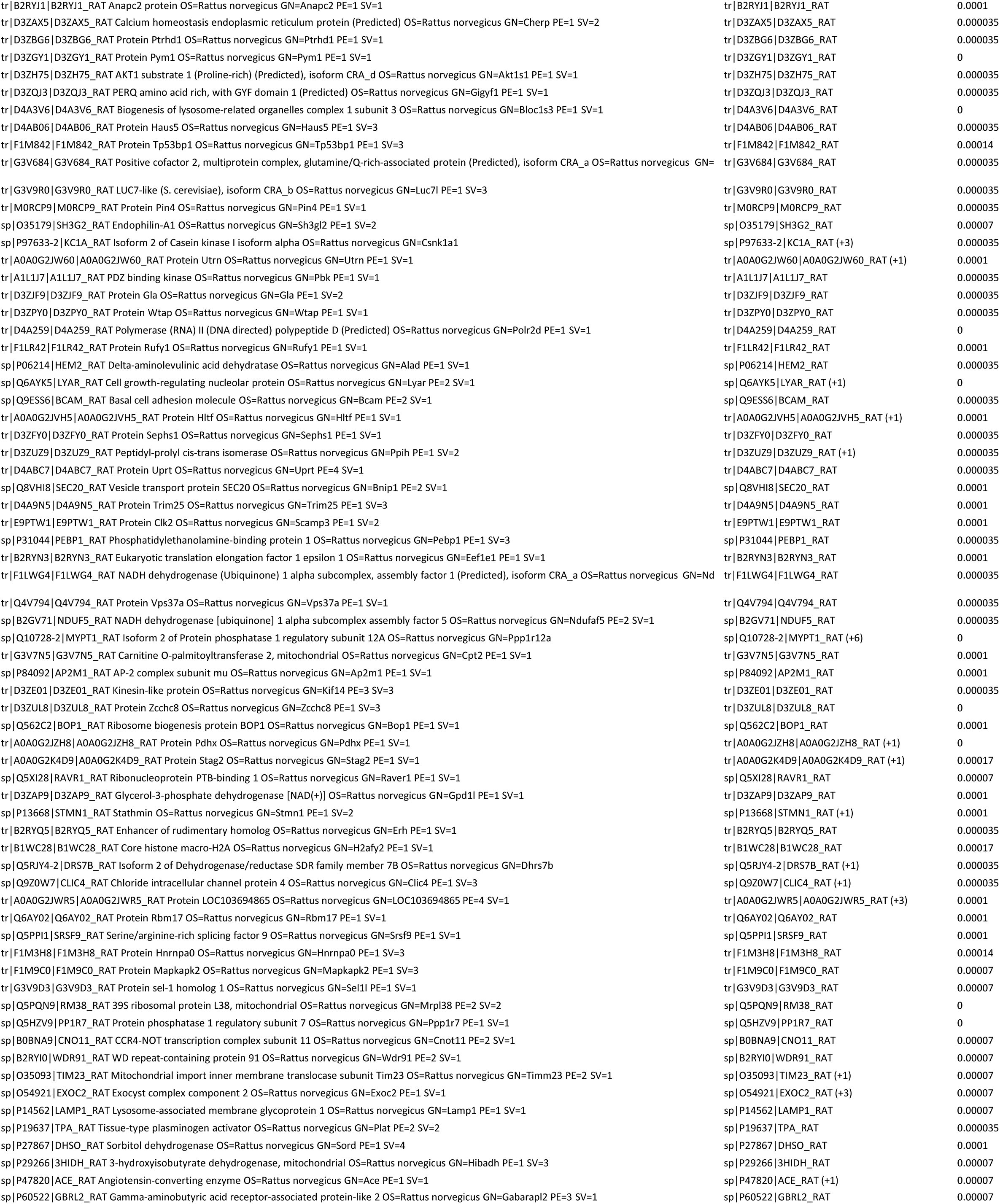
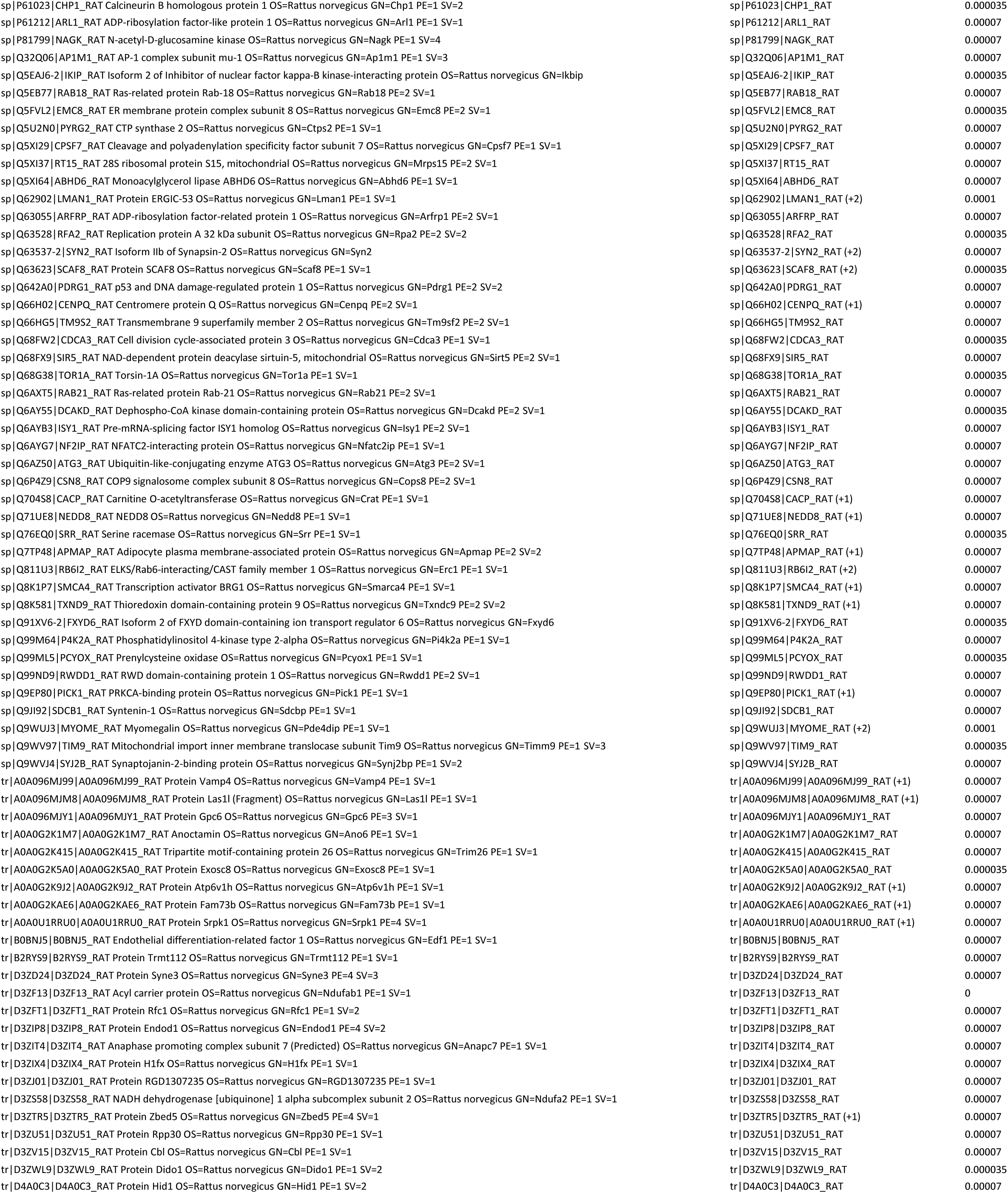
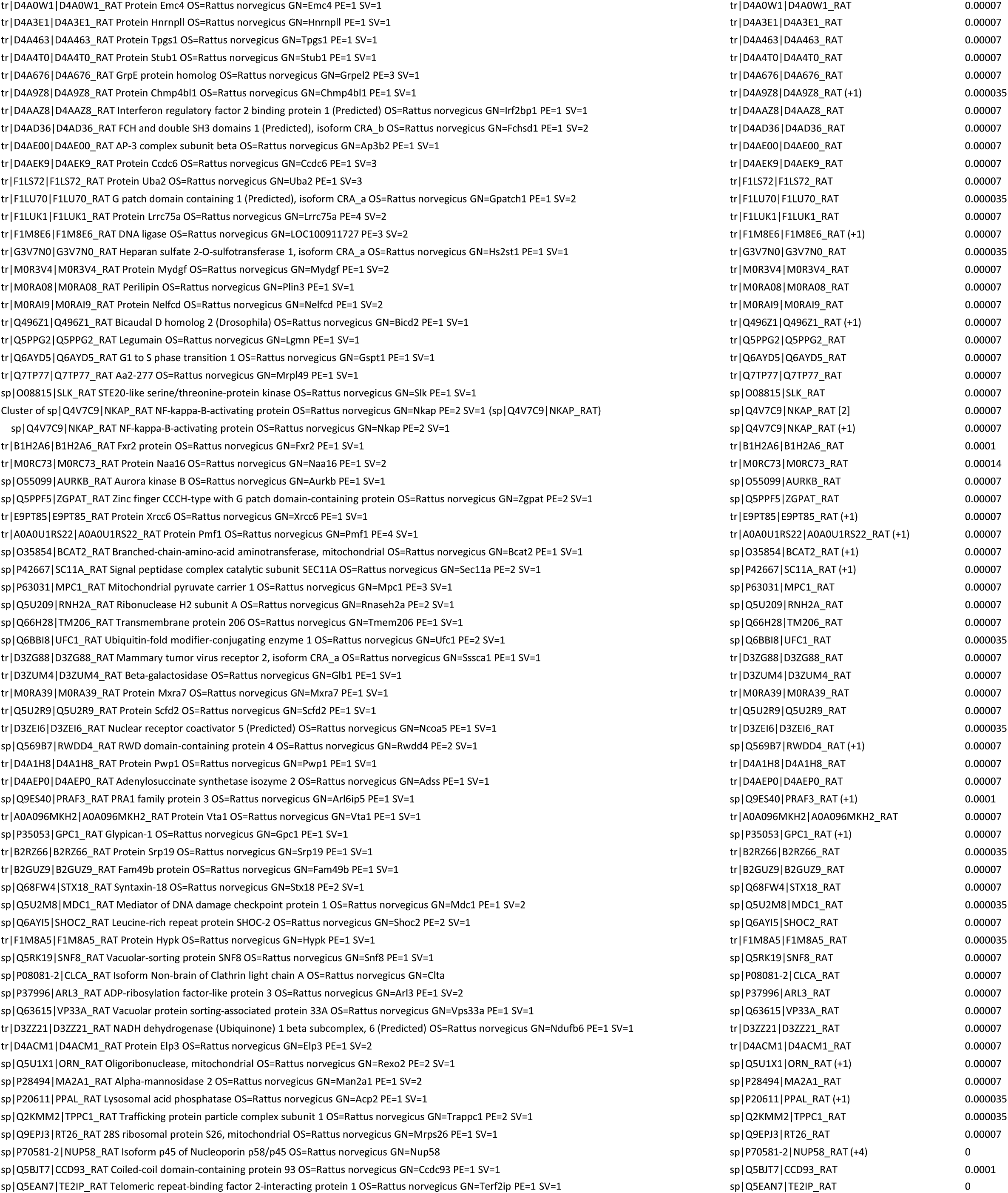
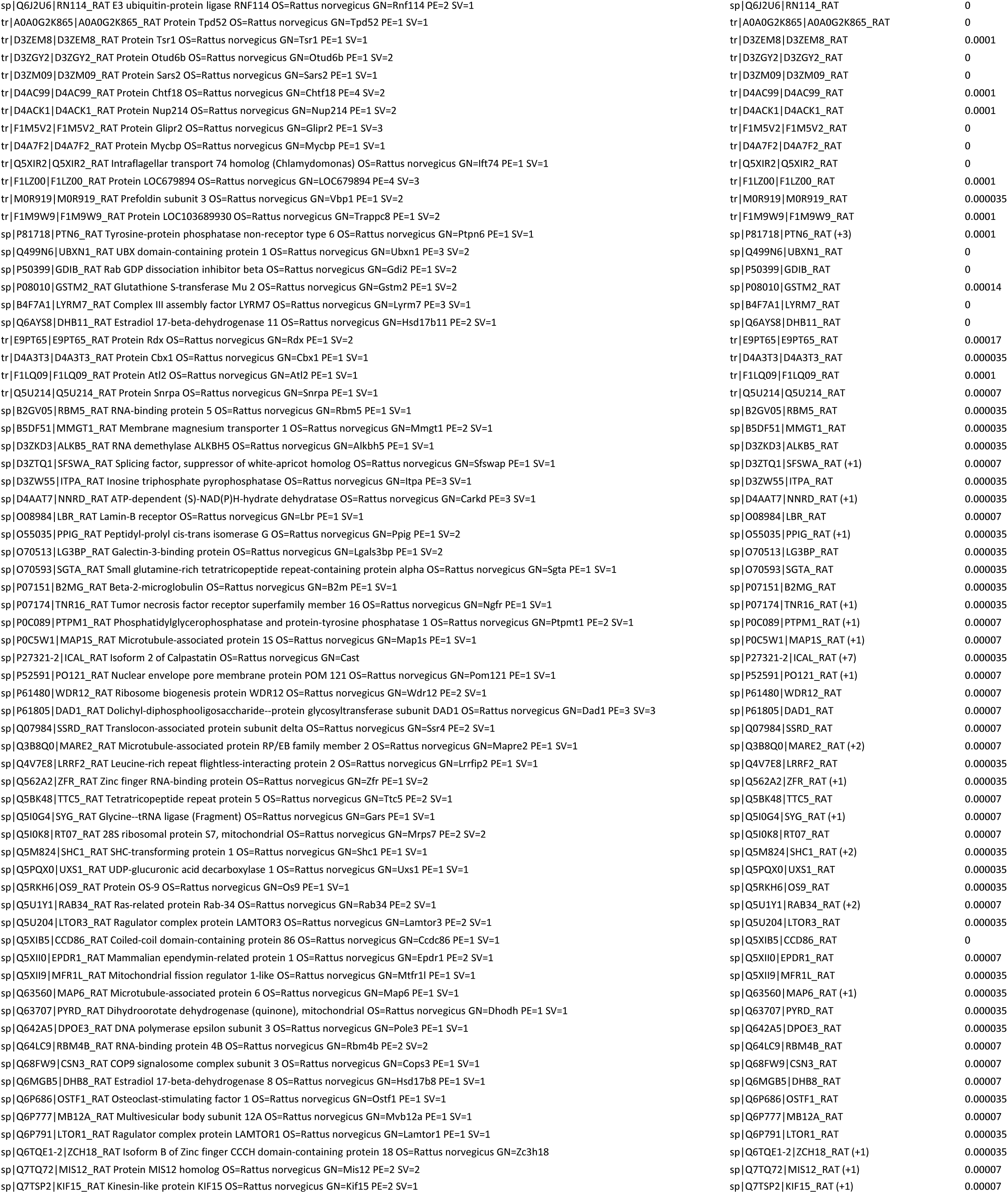
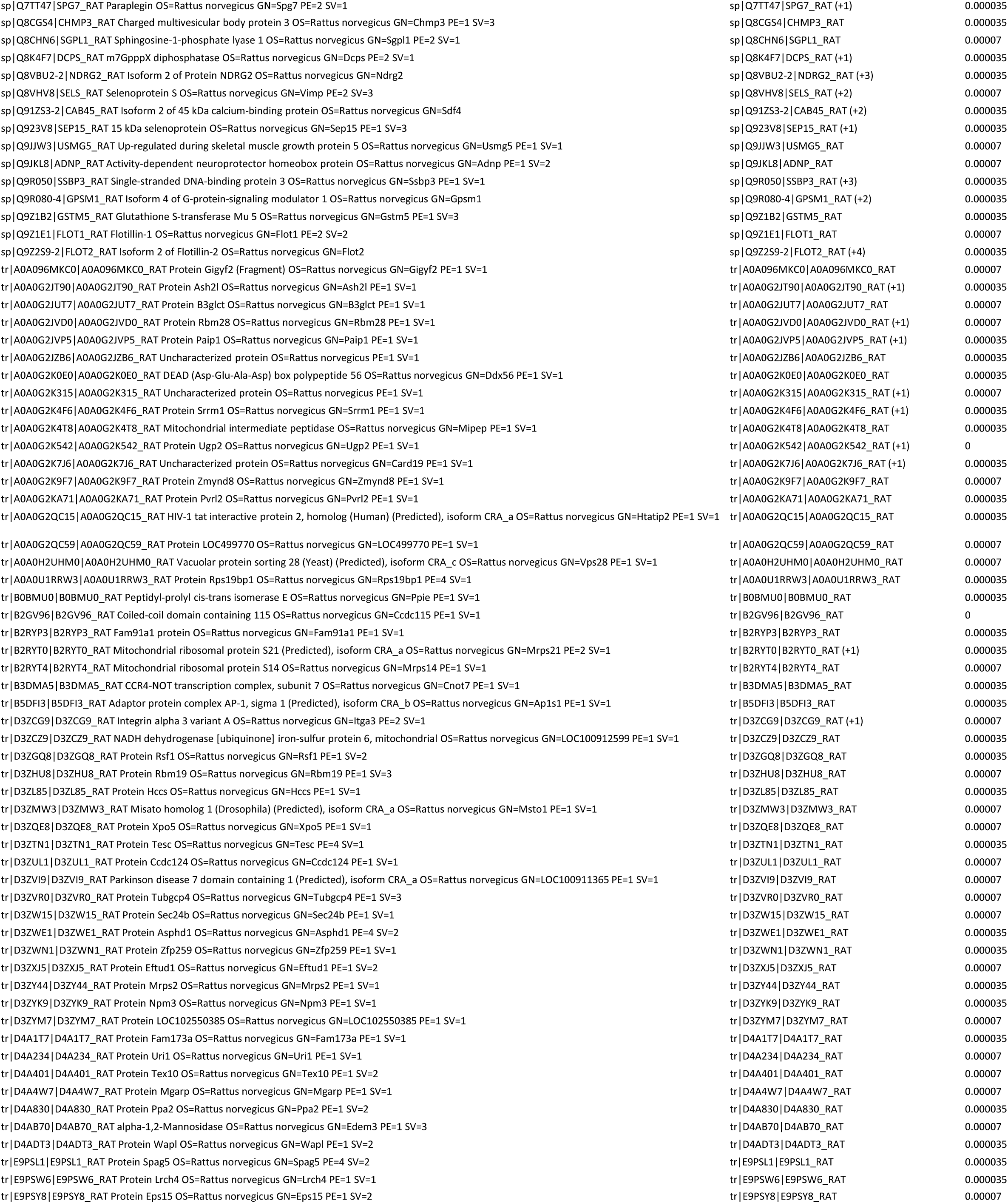
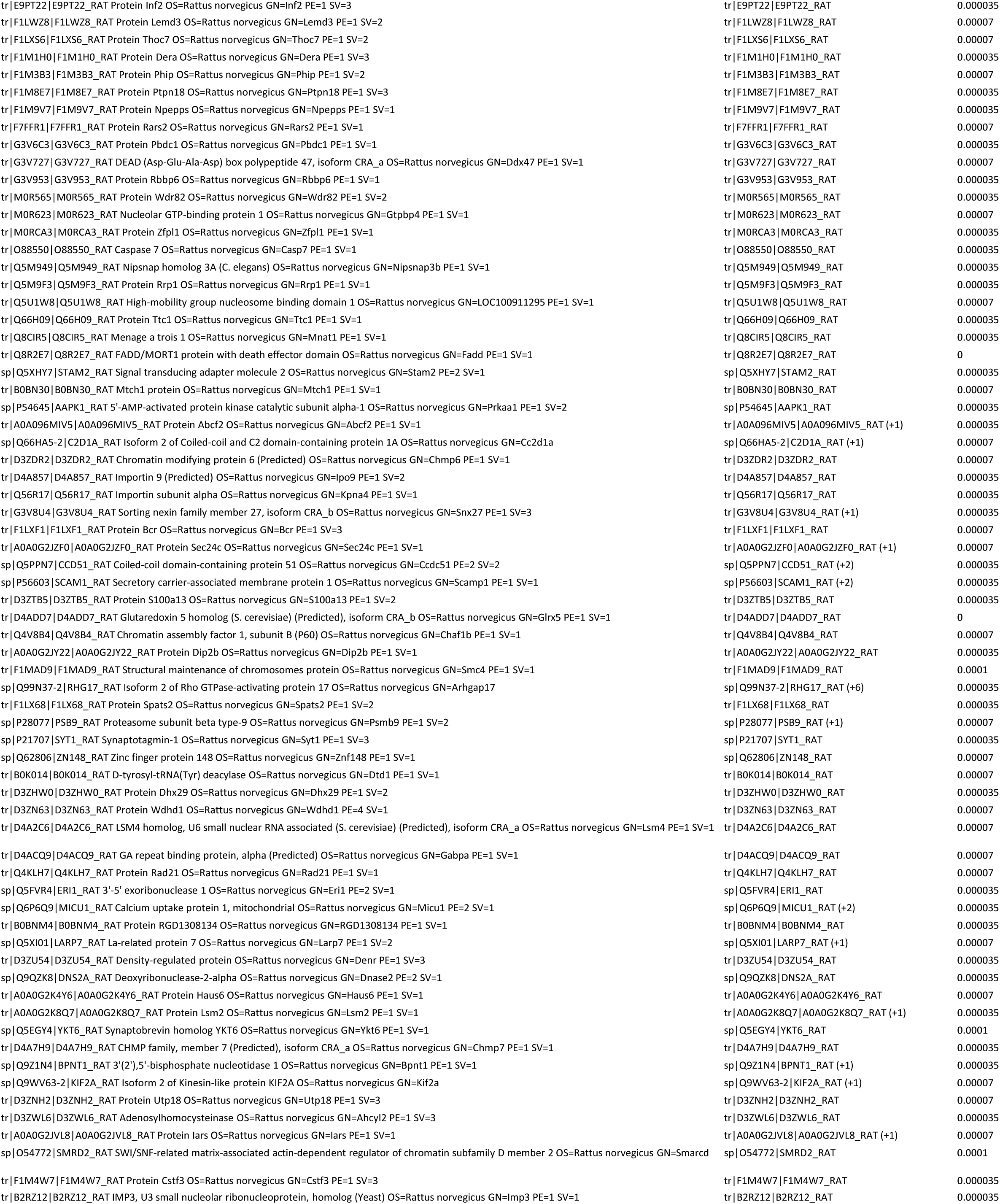
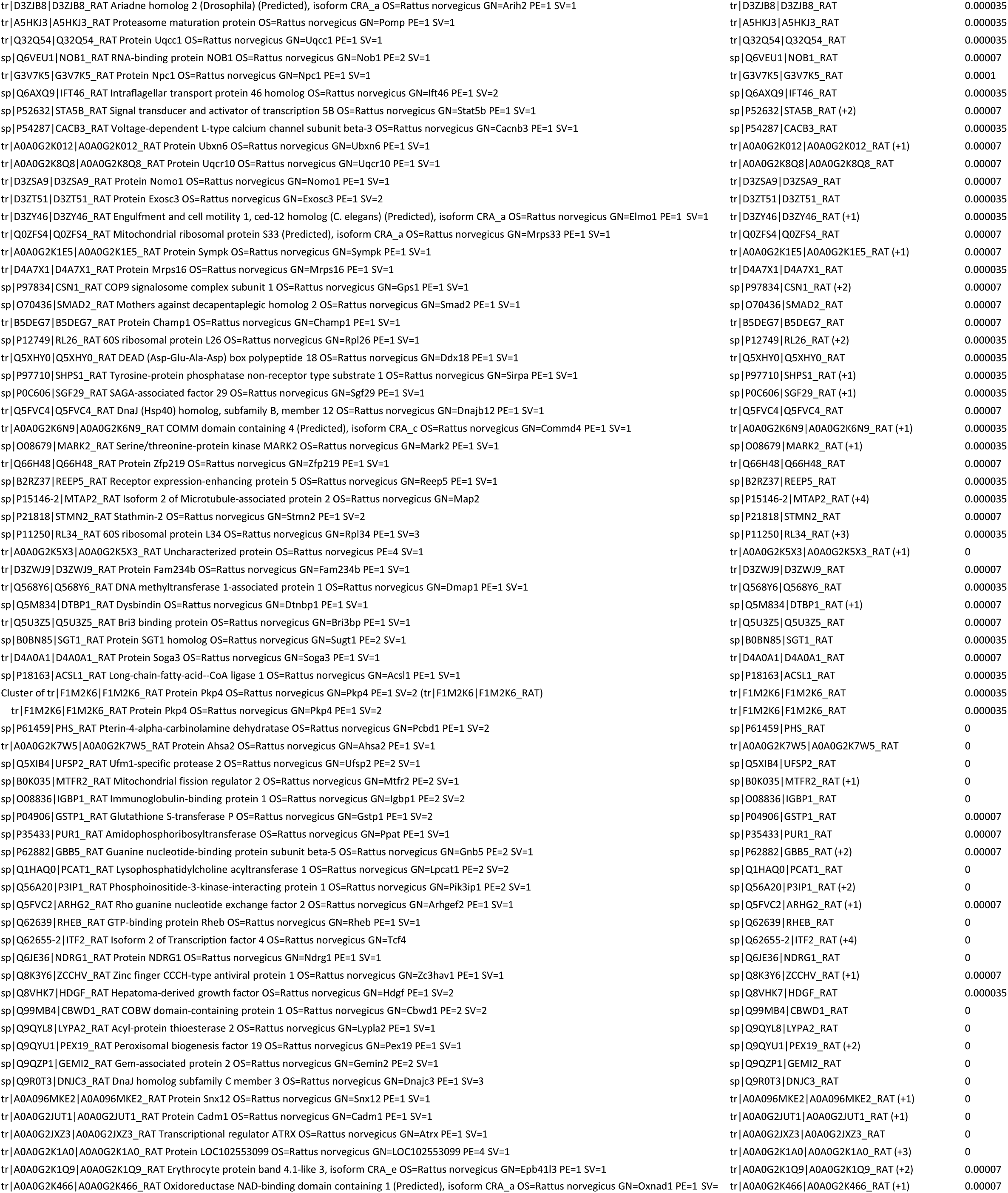
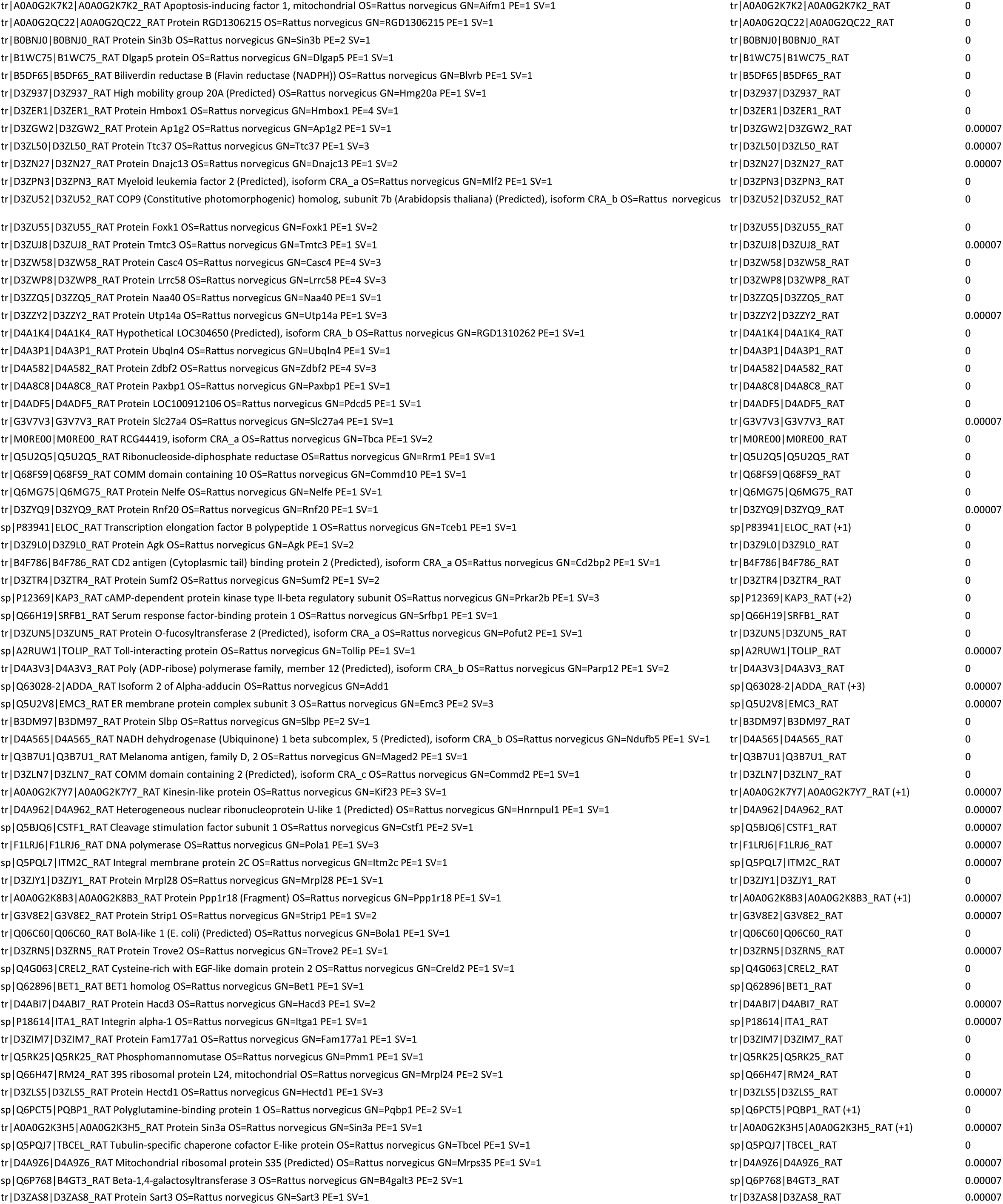
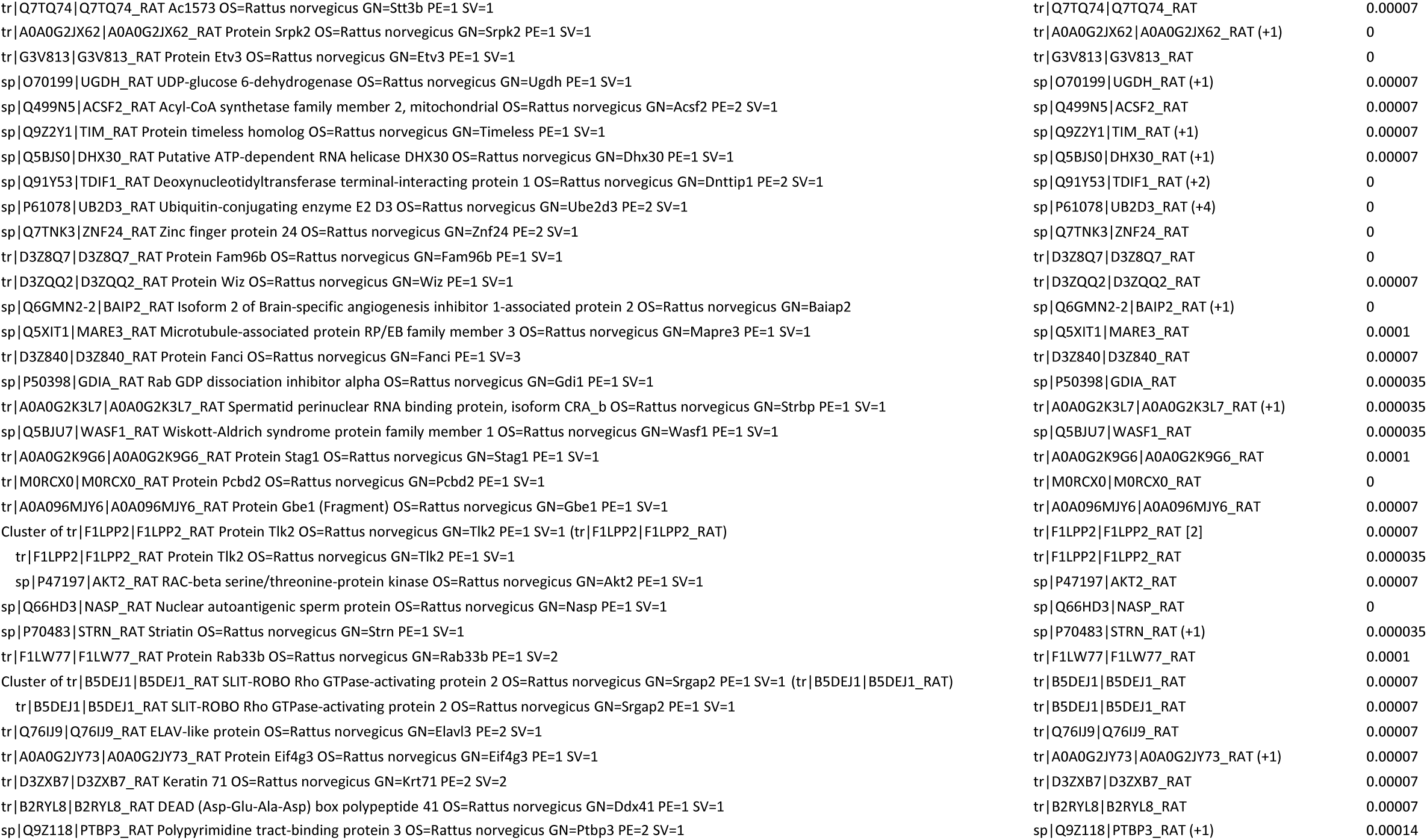
PLCβ1 associated proteins from cytosolic fractions from unsynchronized PC12 cells with reduced PLCb1 expression.

**Figure 1B.**
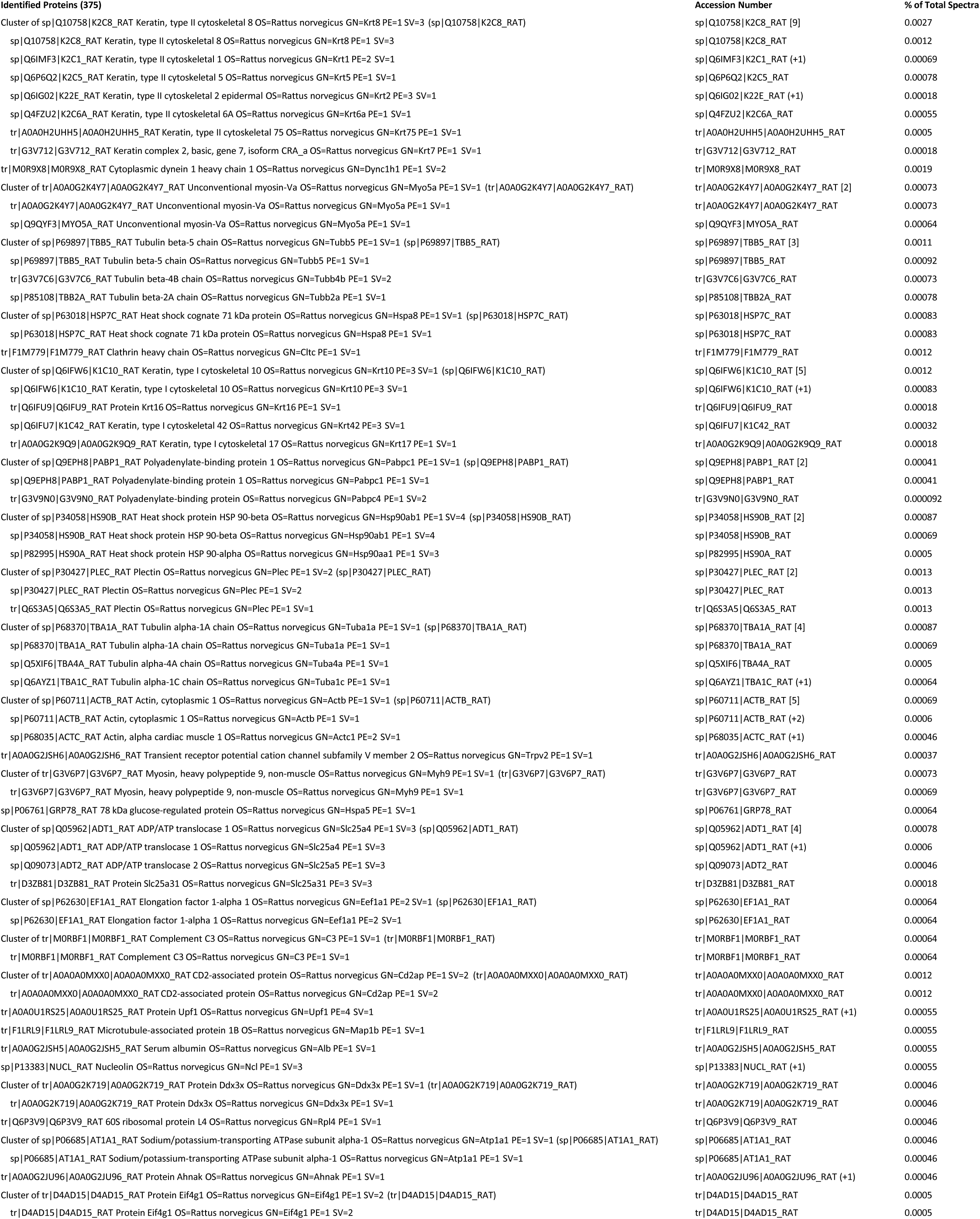
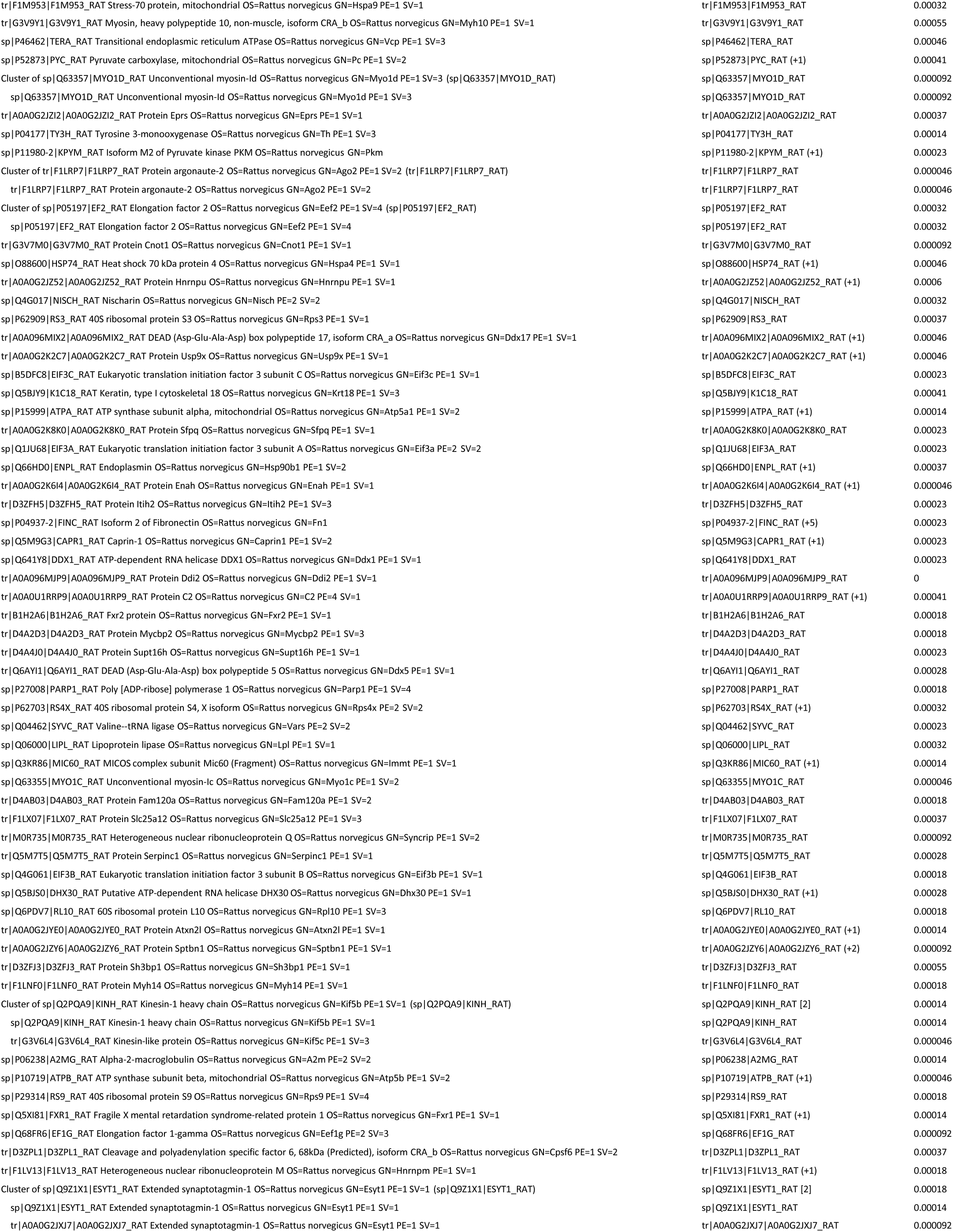
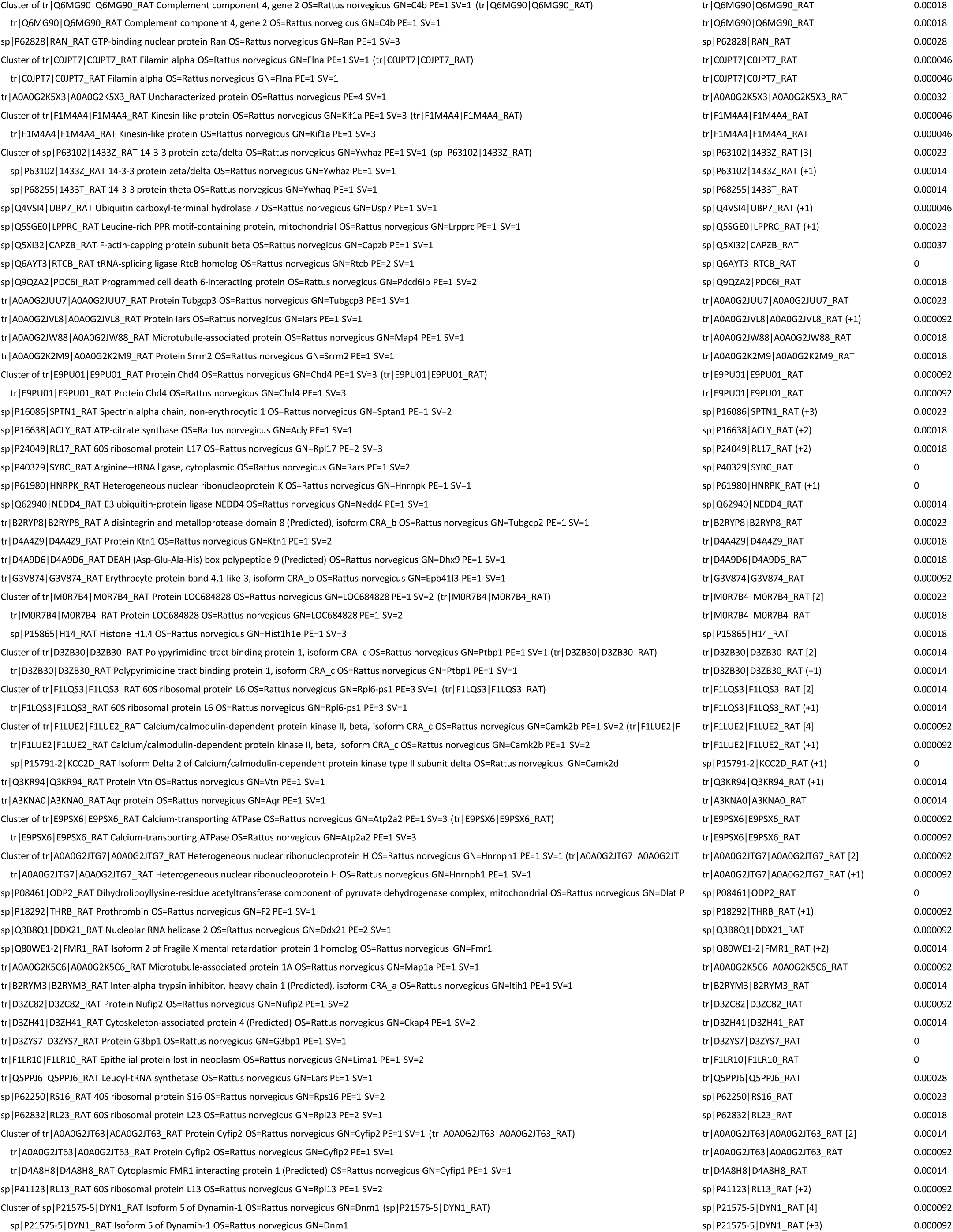
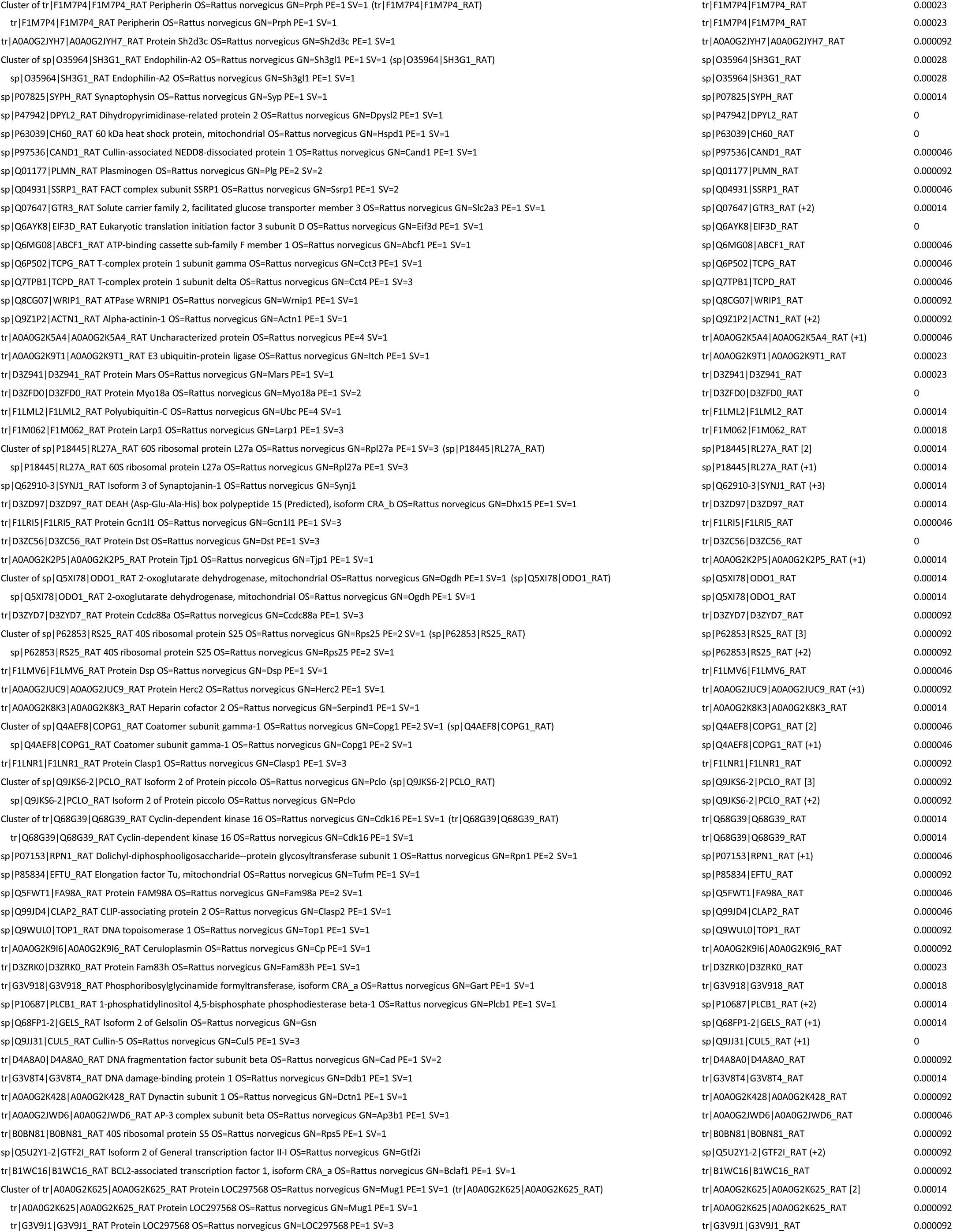
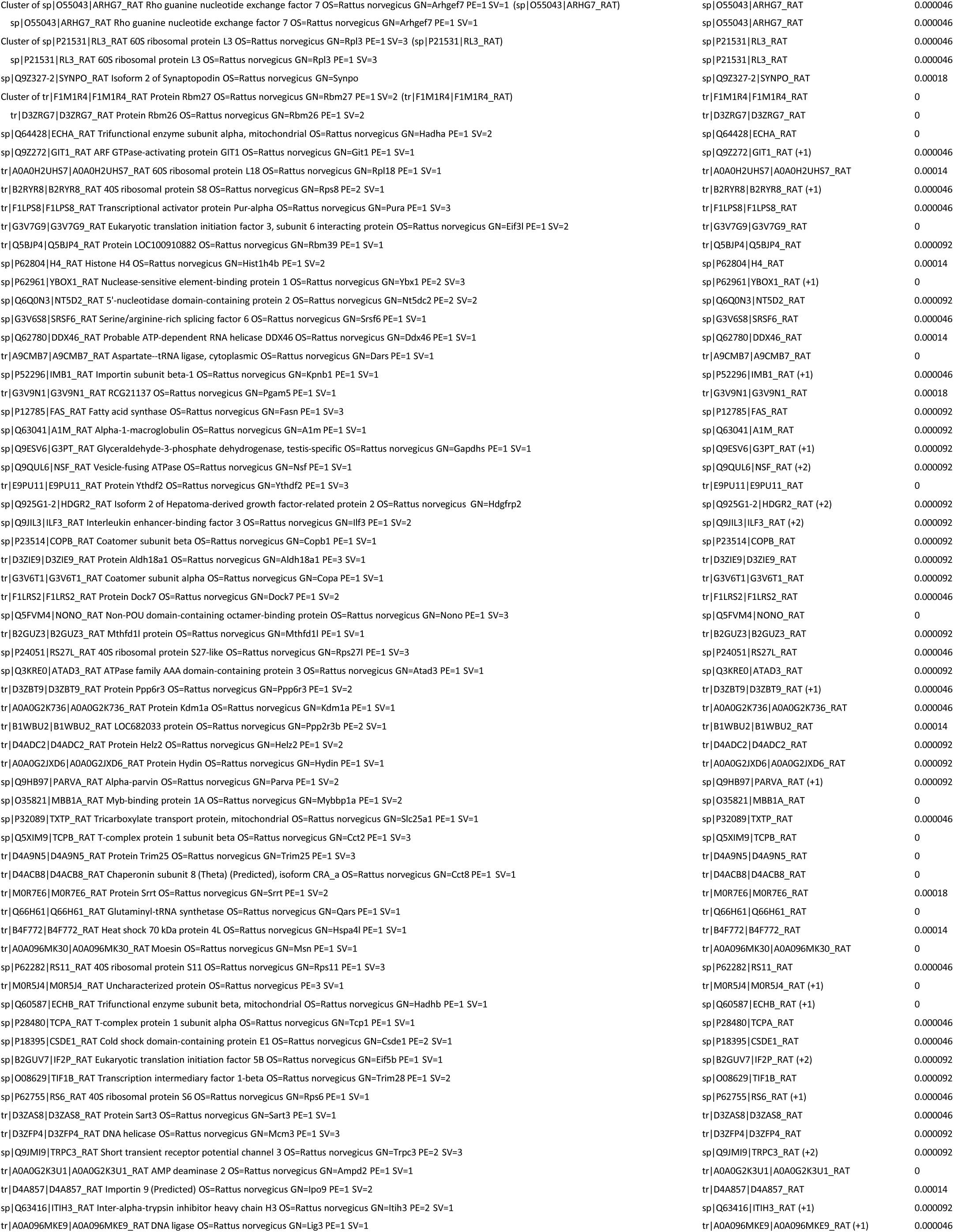
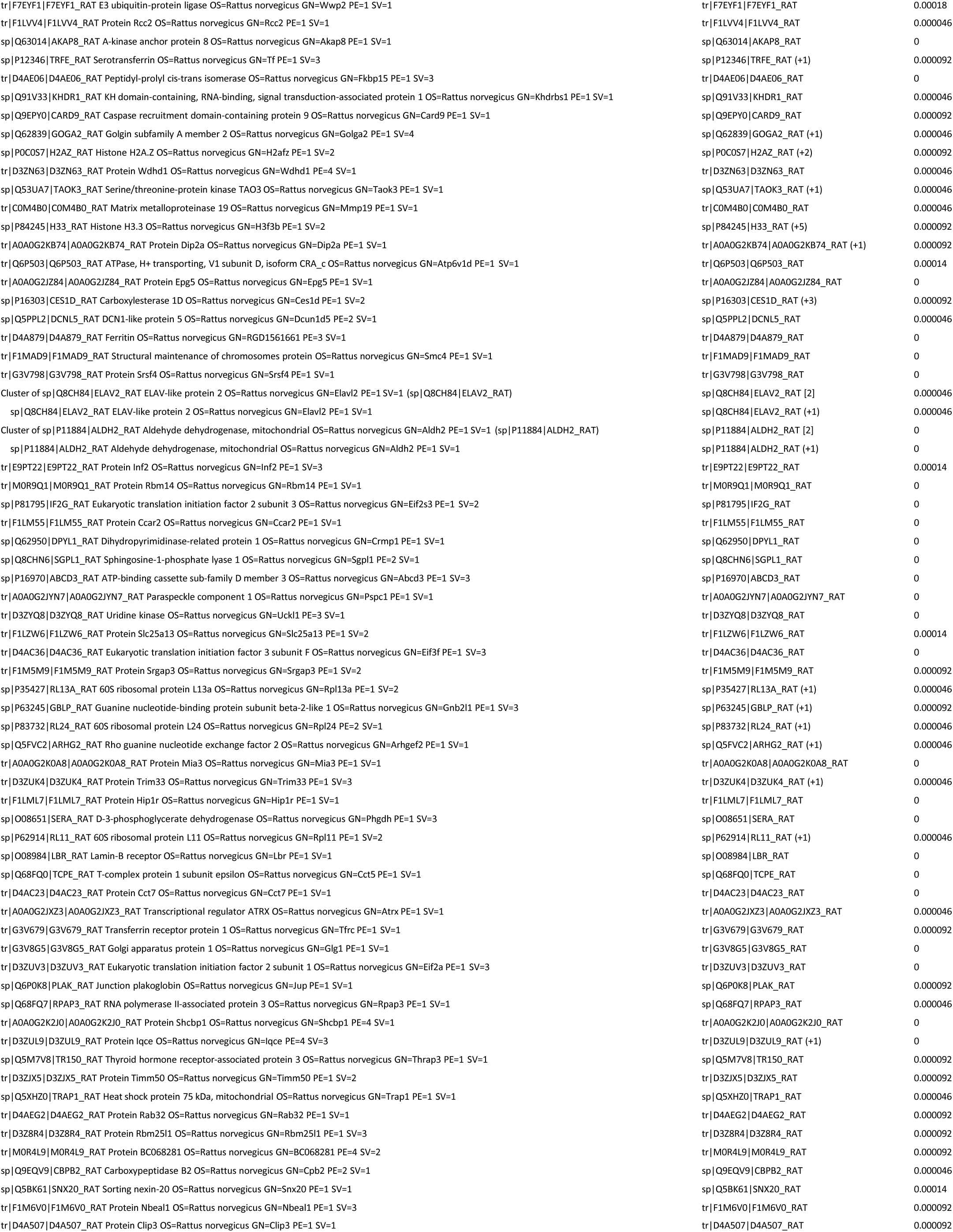
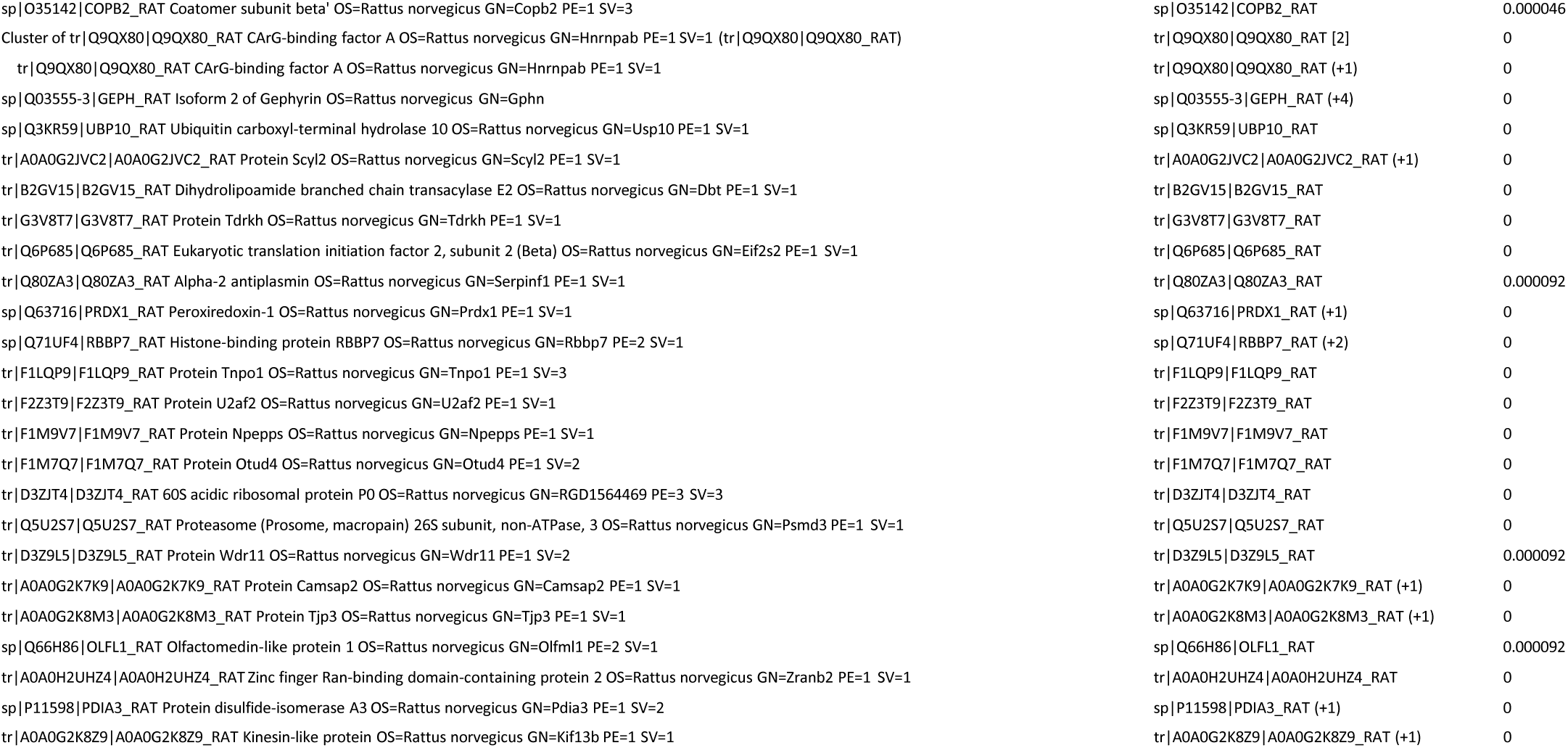
PLCβ1 associated proteins from cytosolic fractions from unsynchronized PC12 cells with reduced PLCb1 expression.

**Figure 1C.**
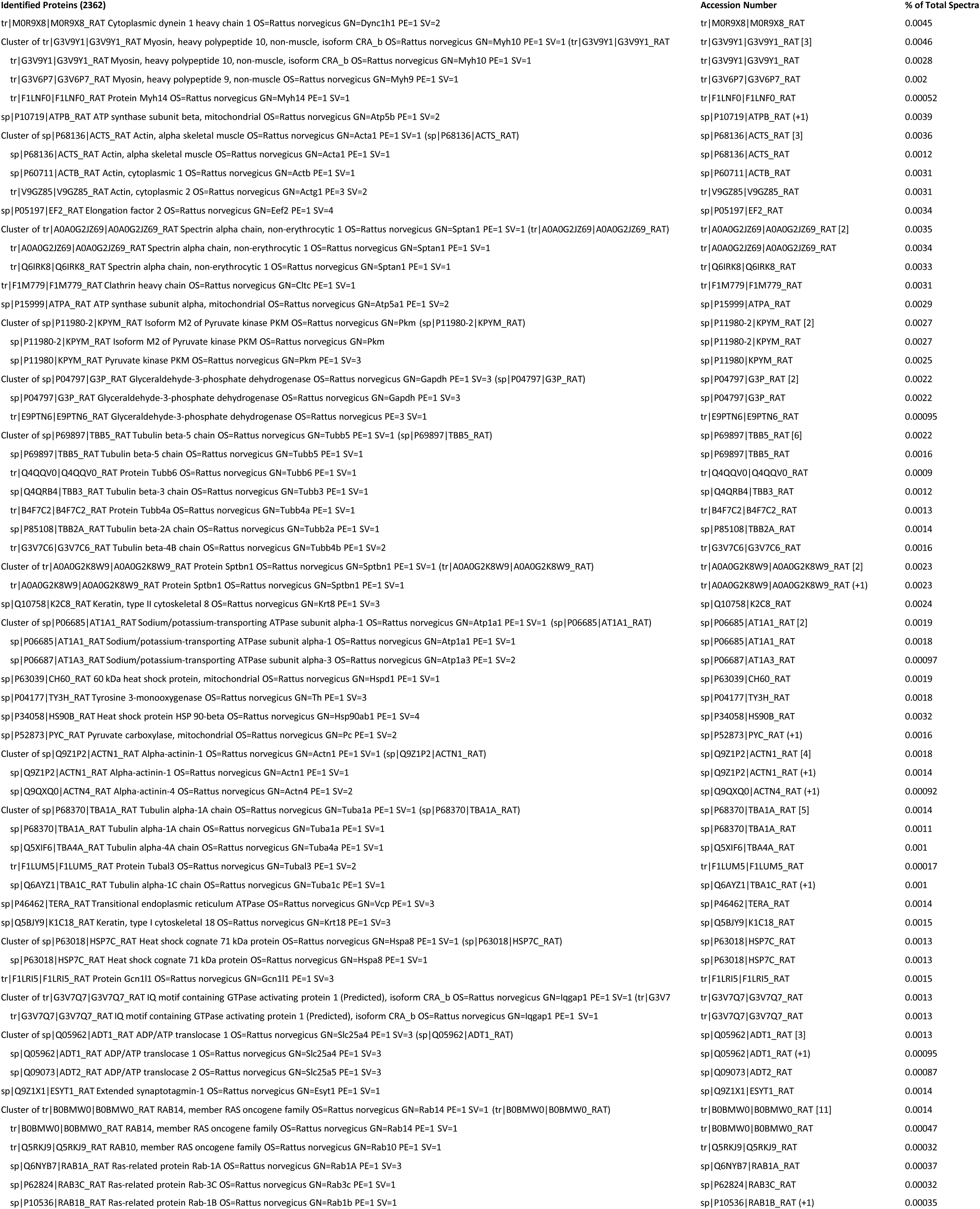
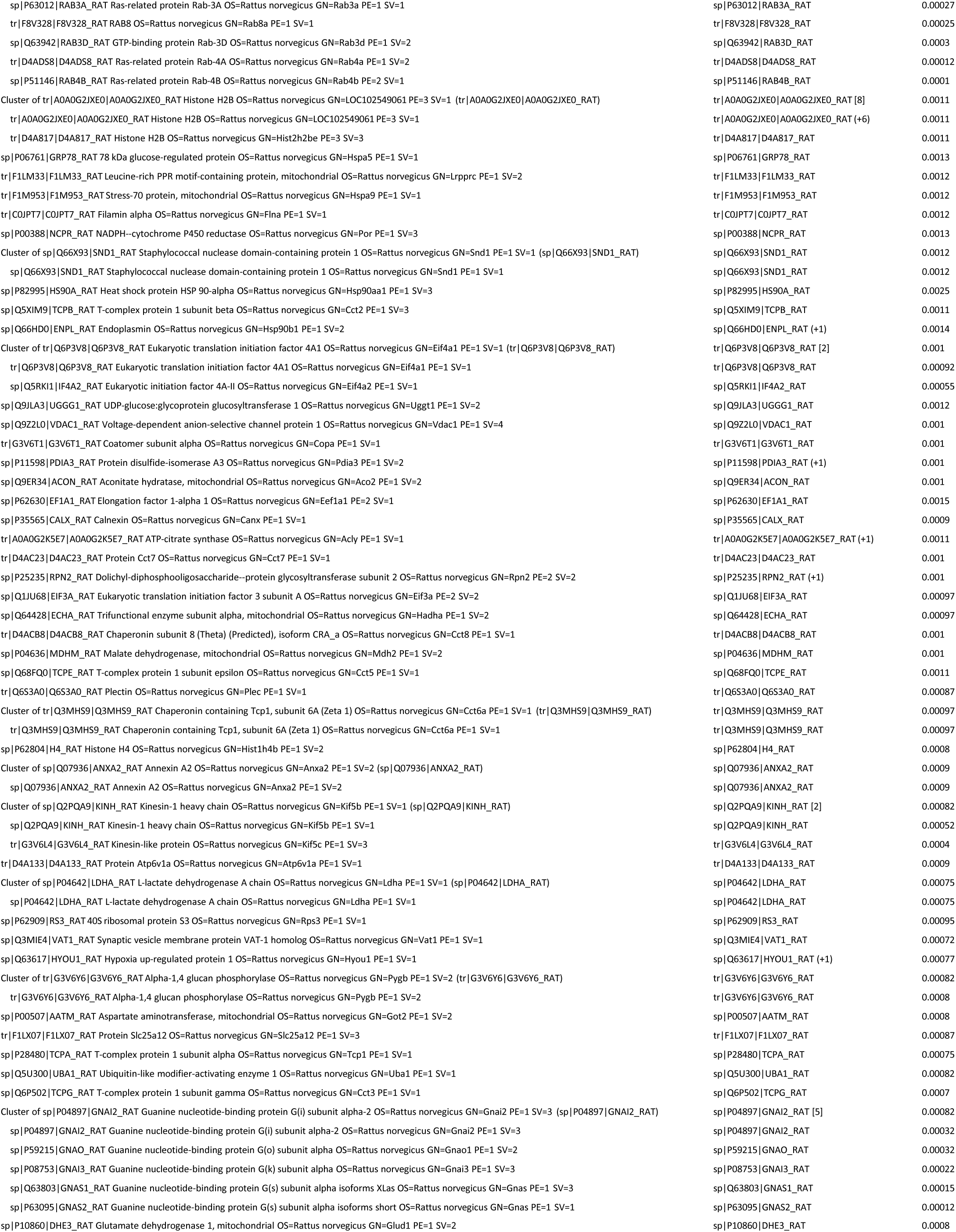
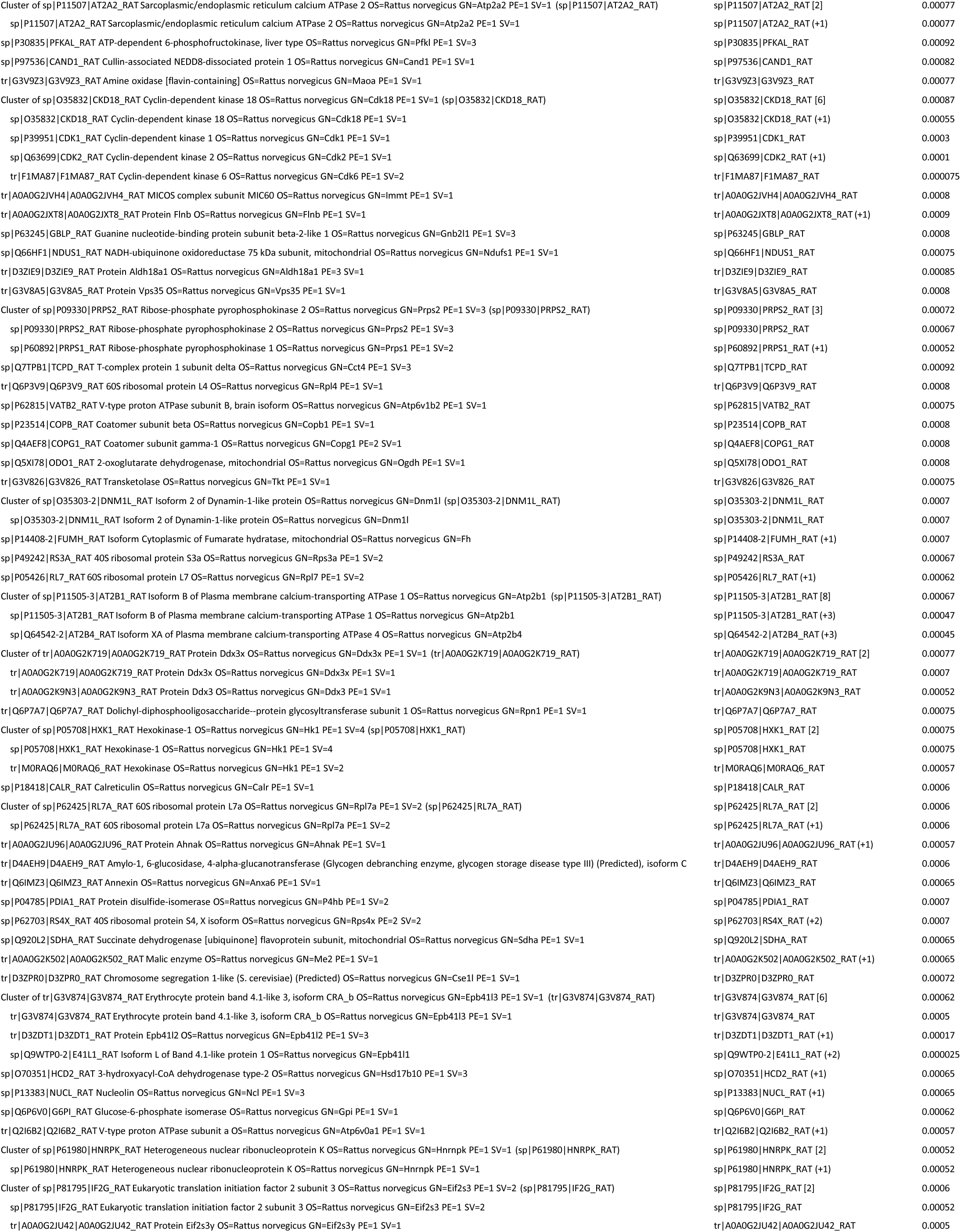
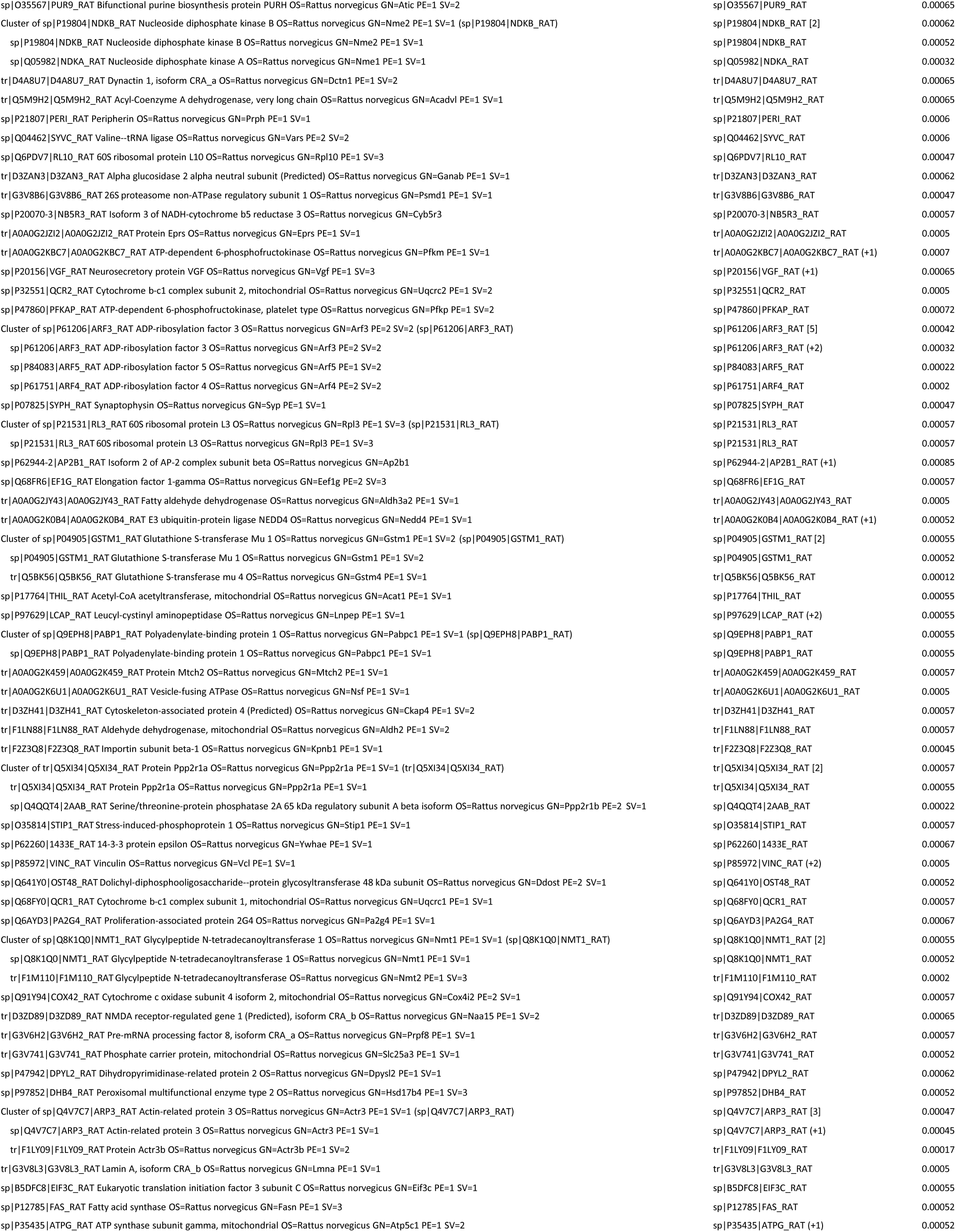
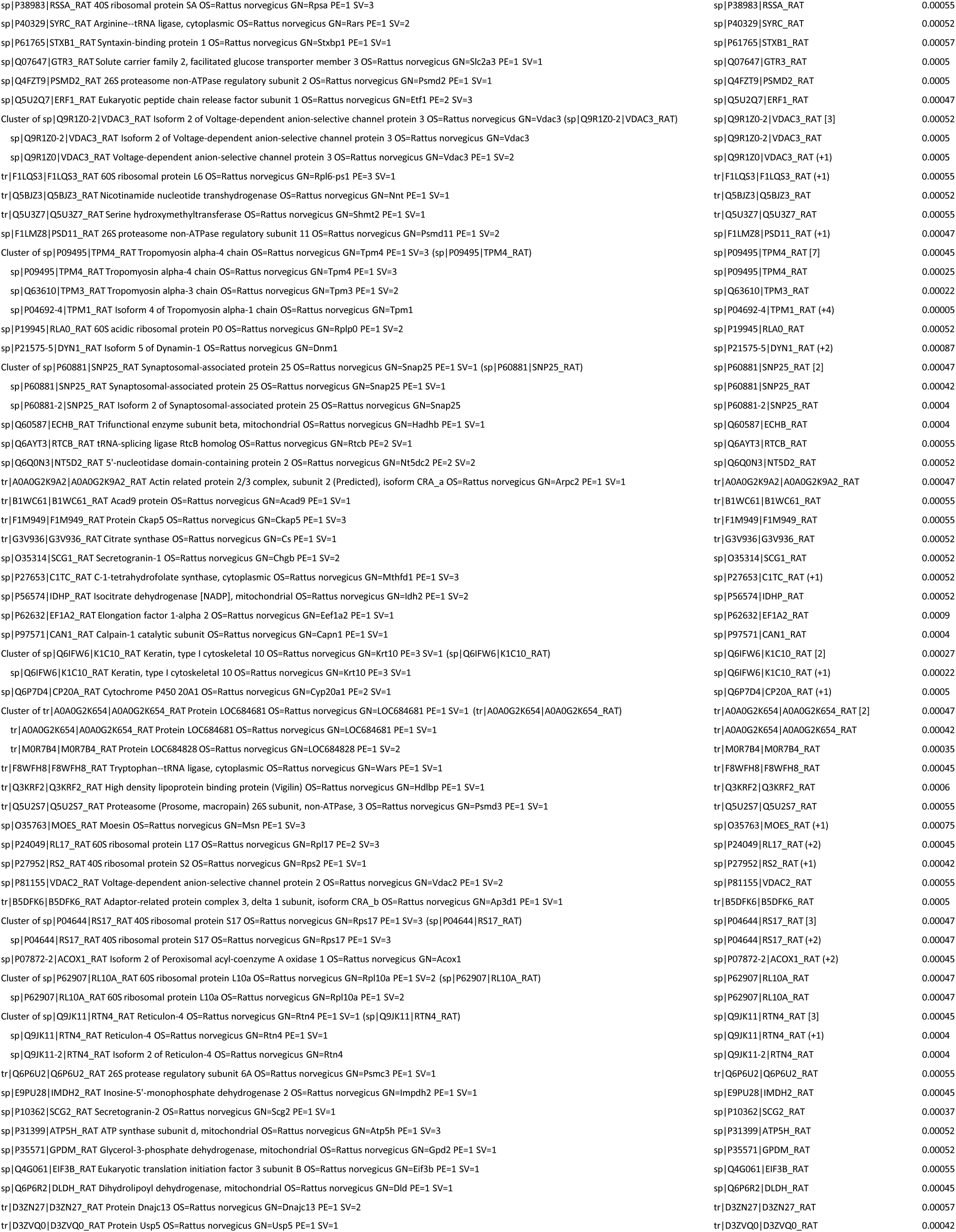
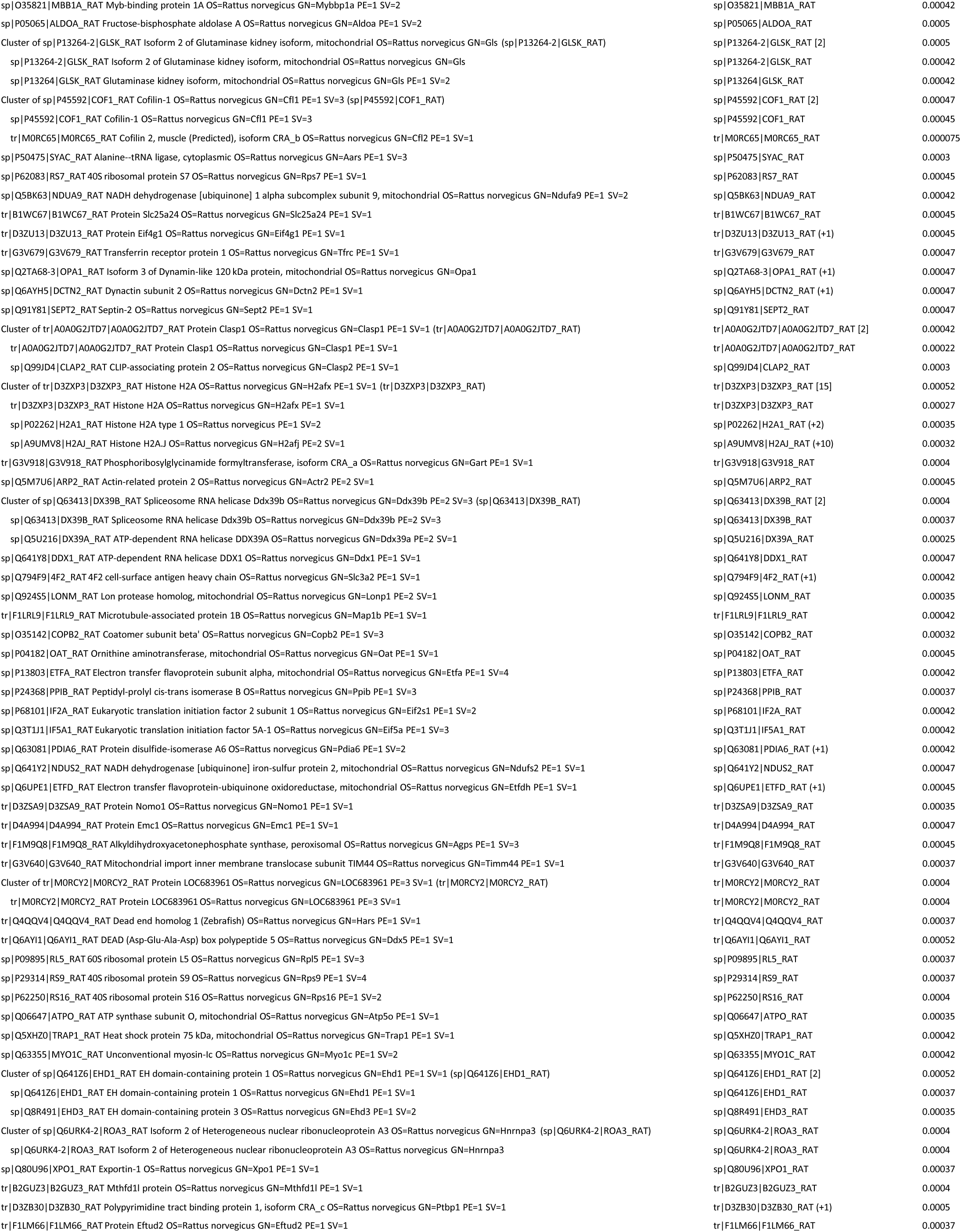
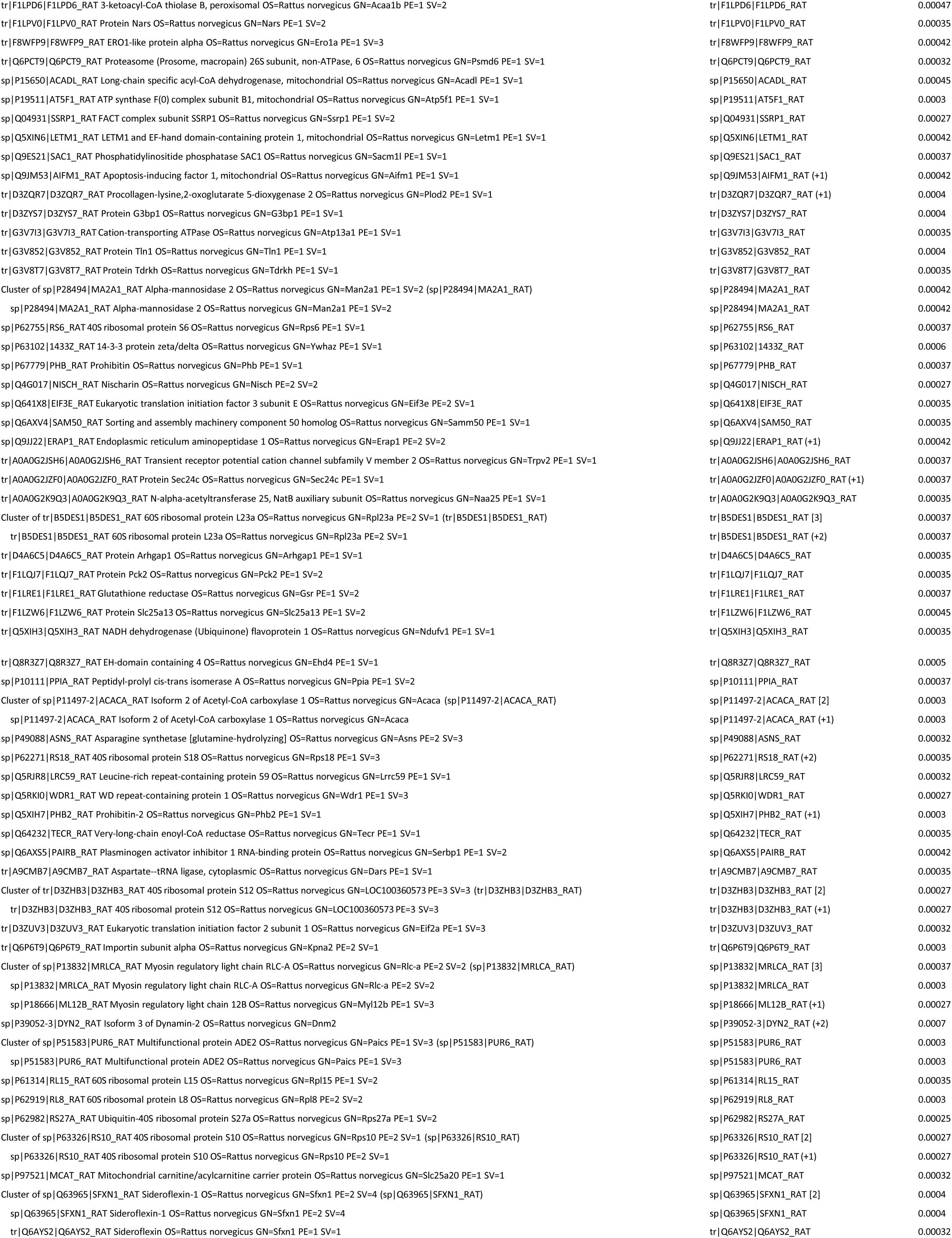
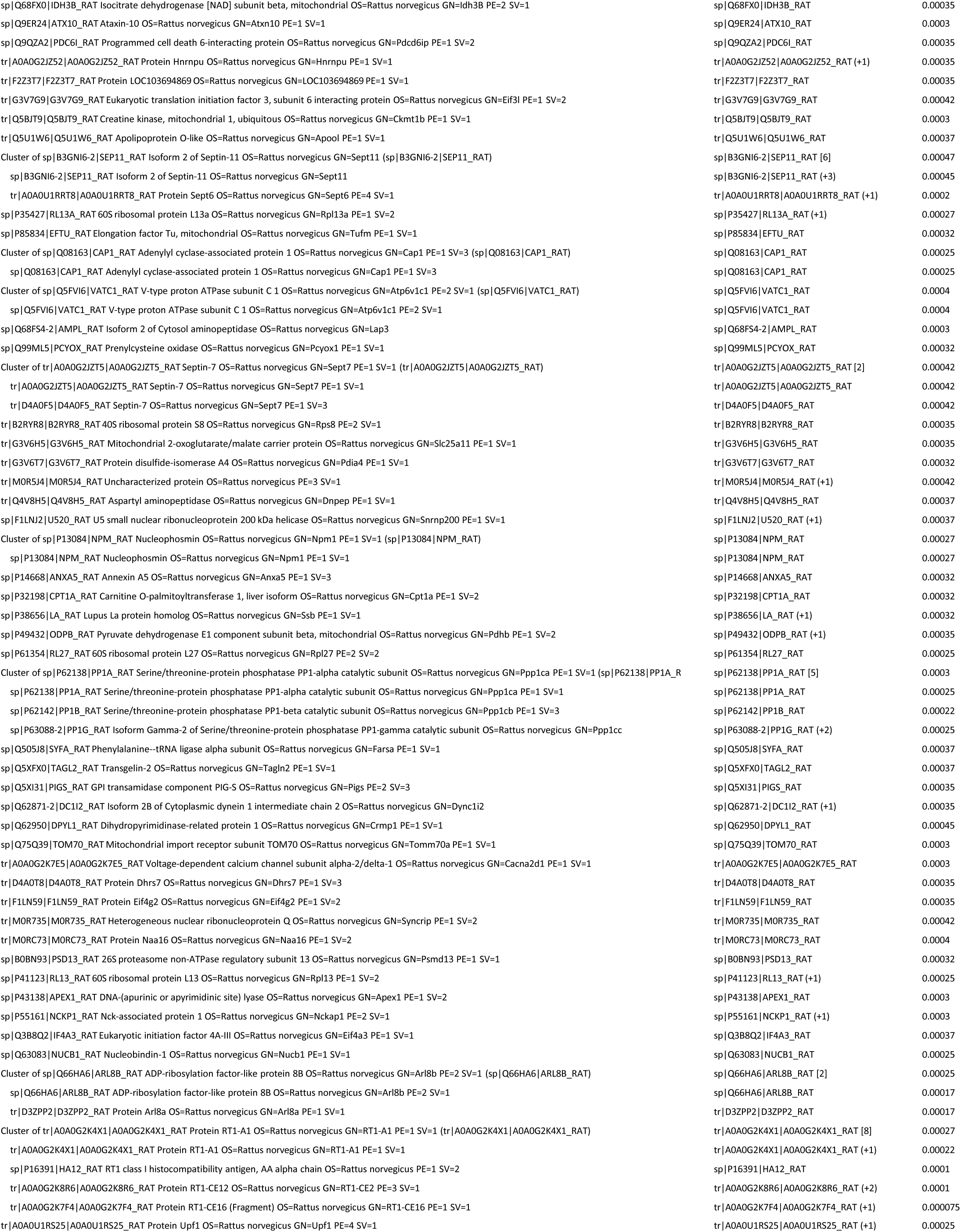
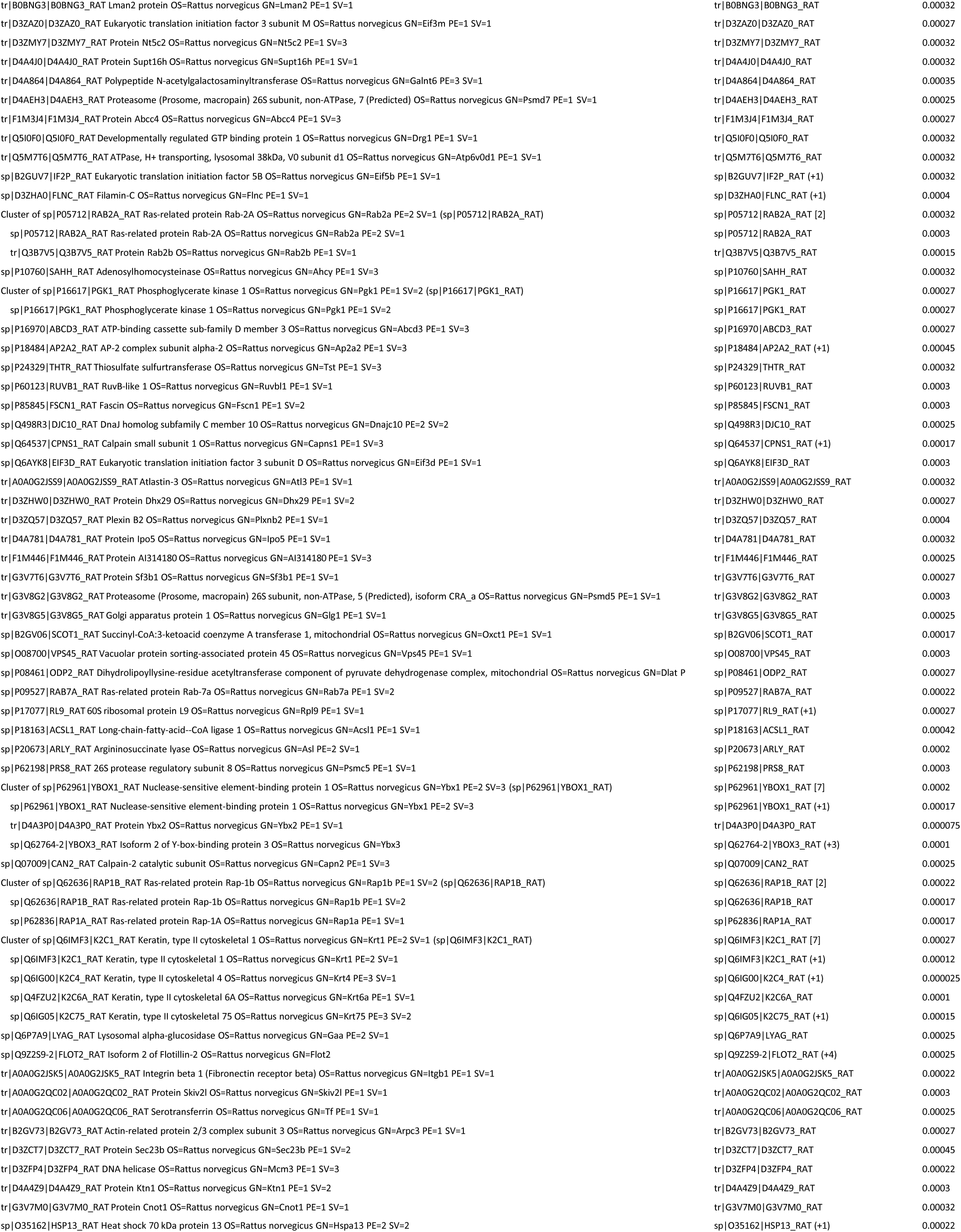
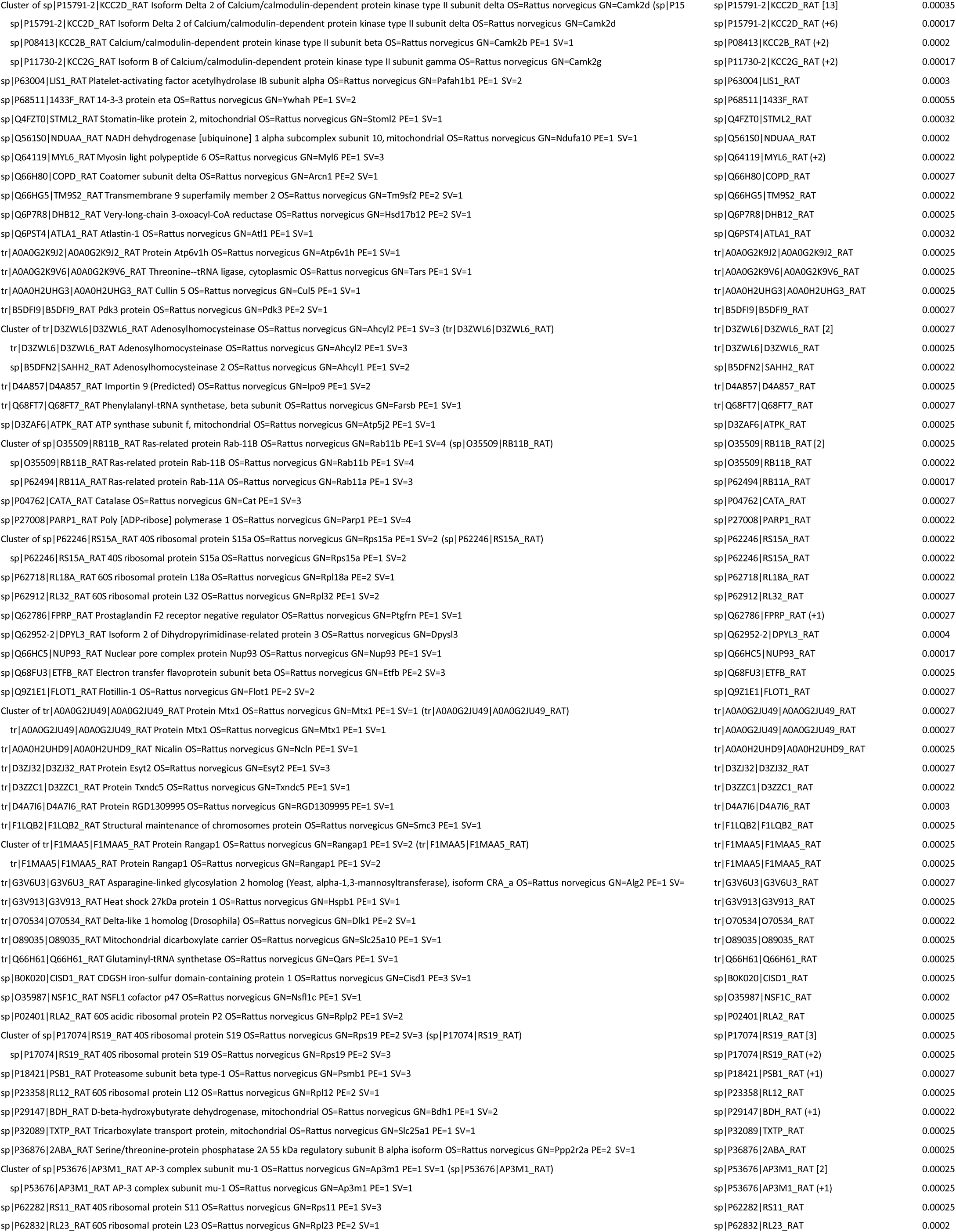
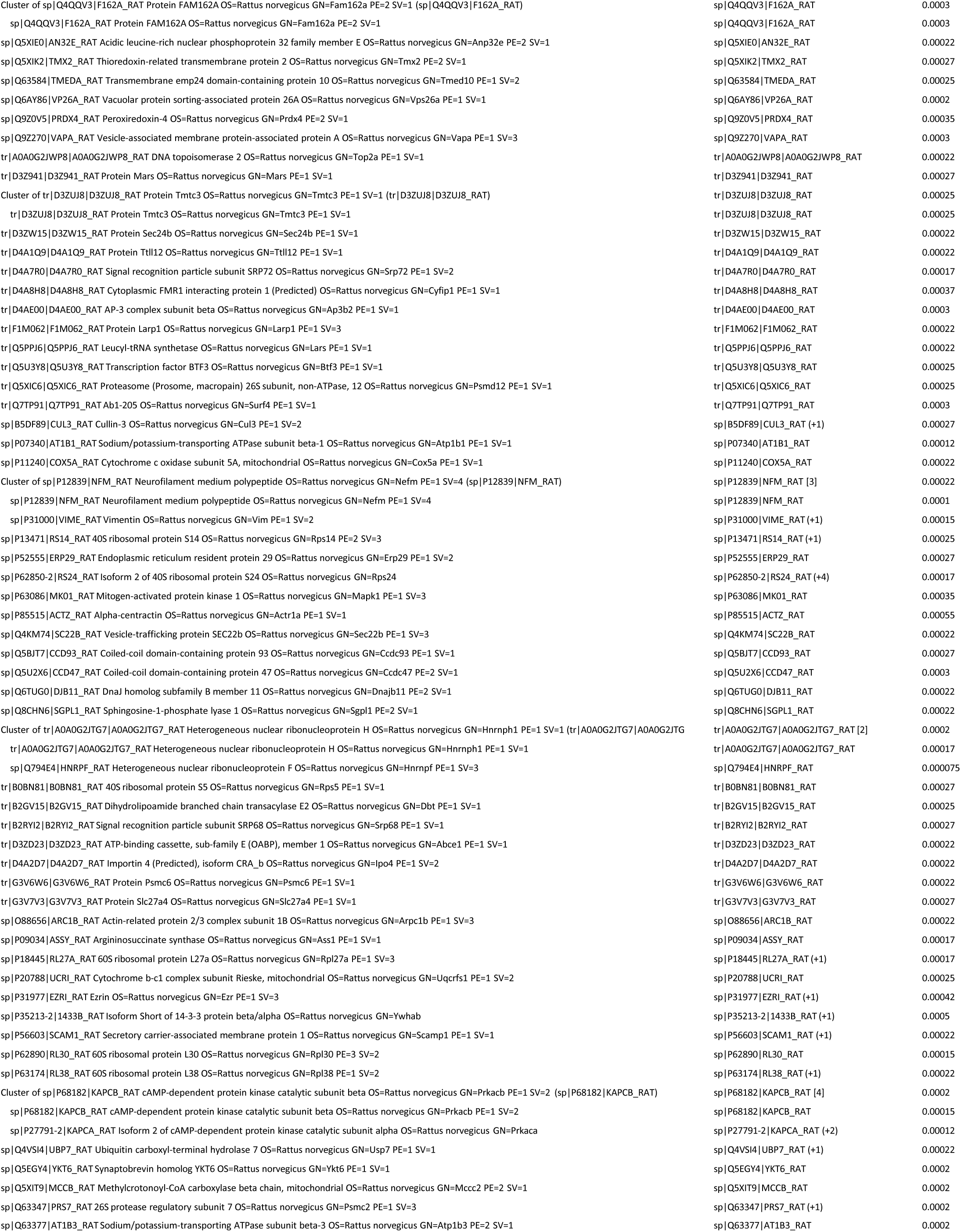
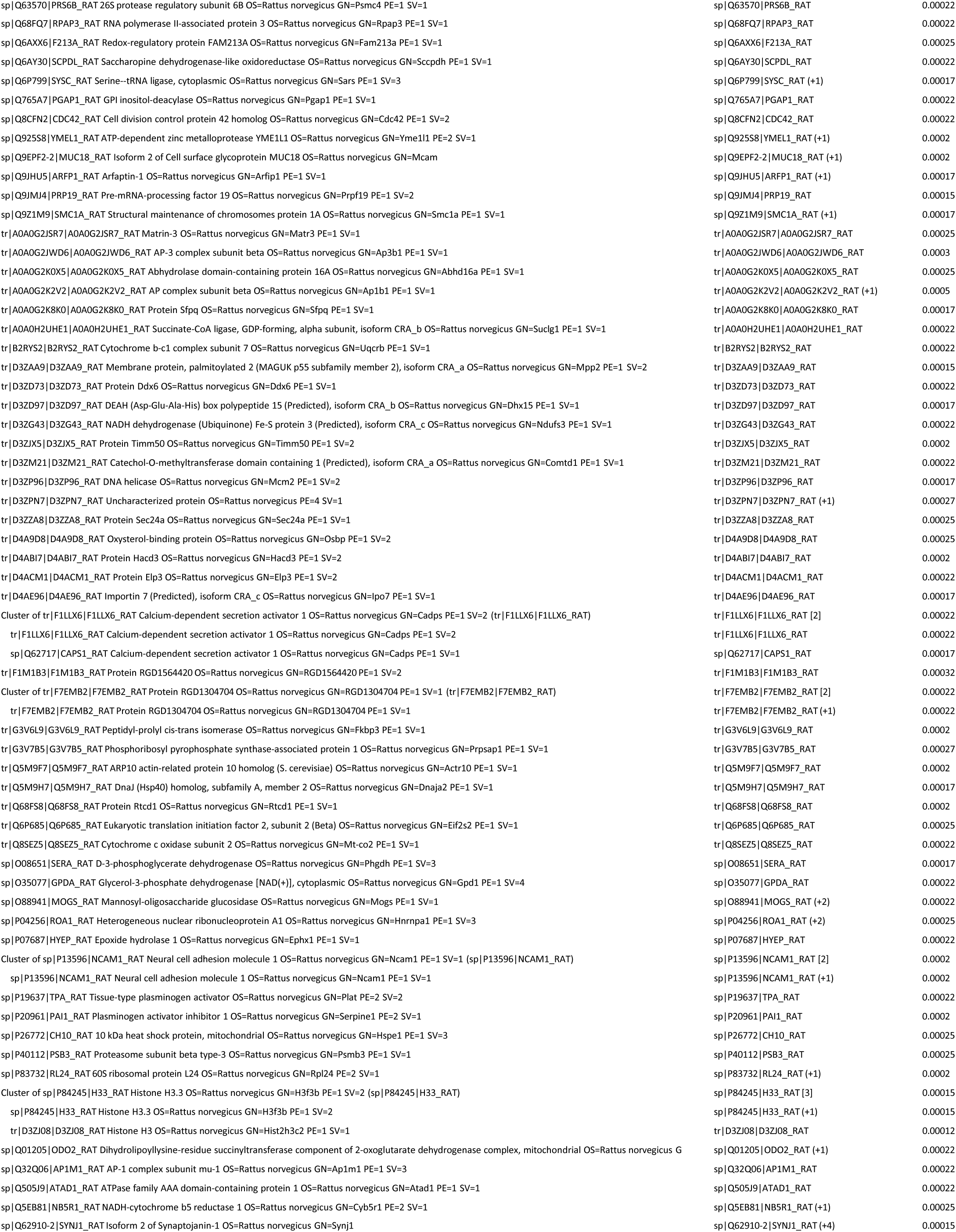
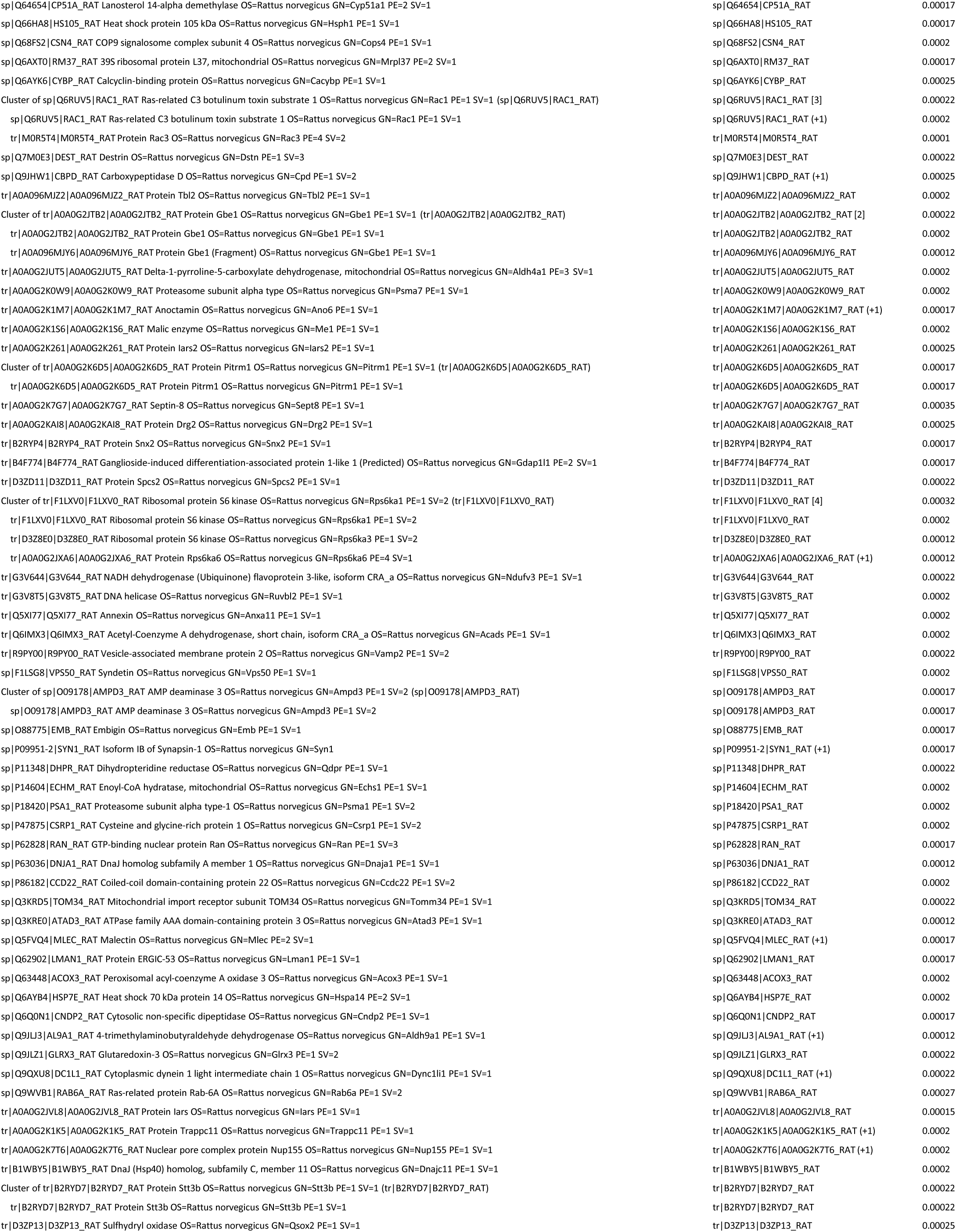
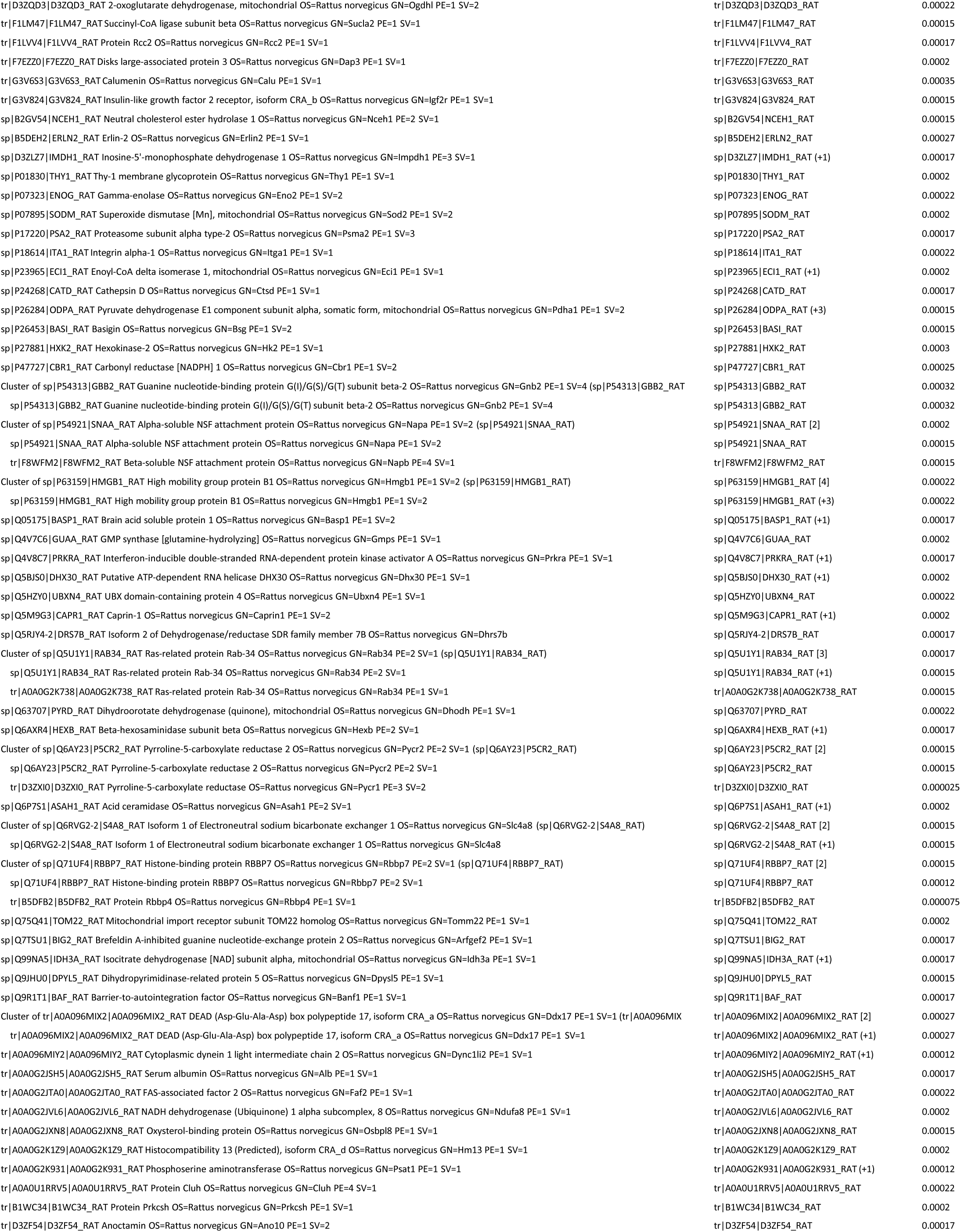
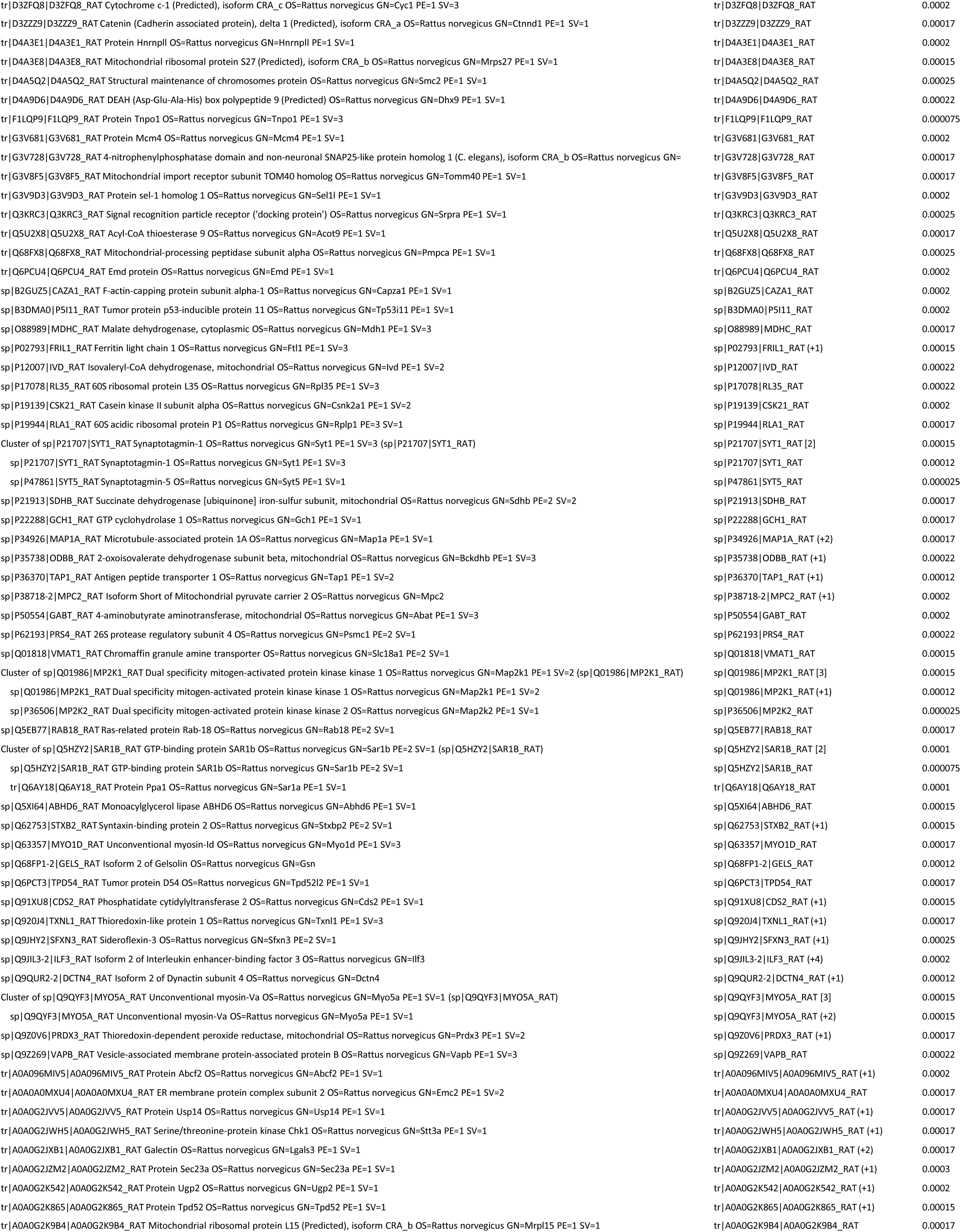
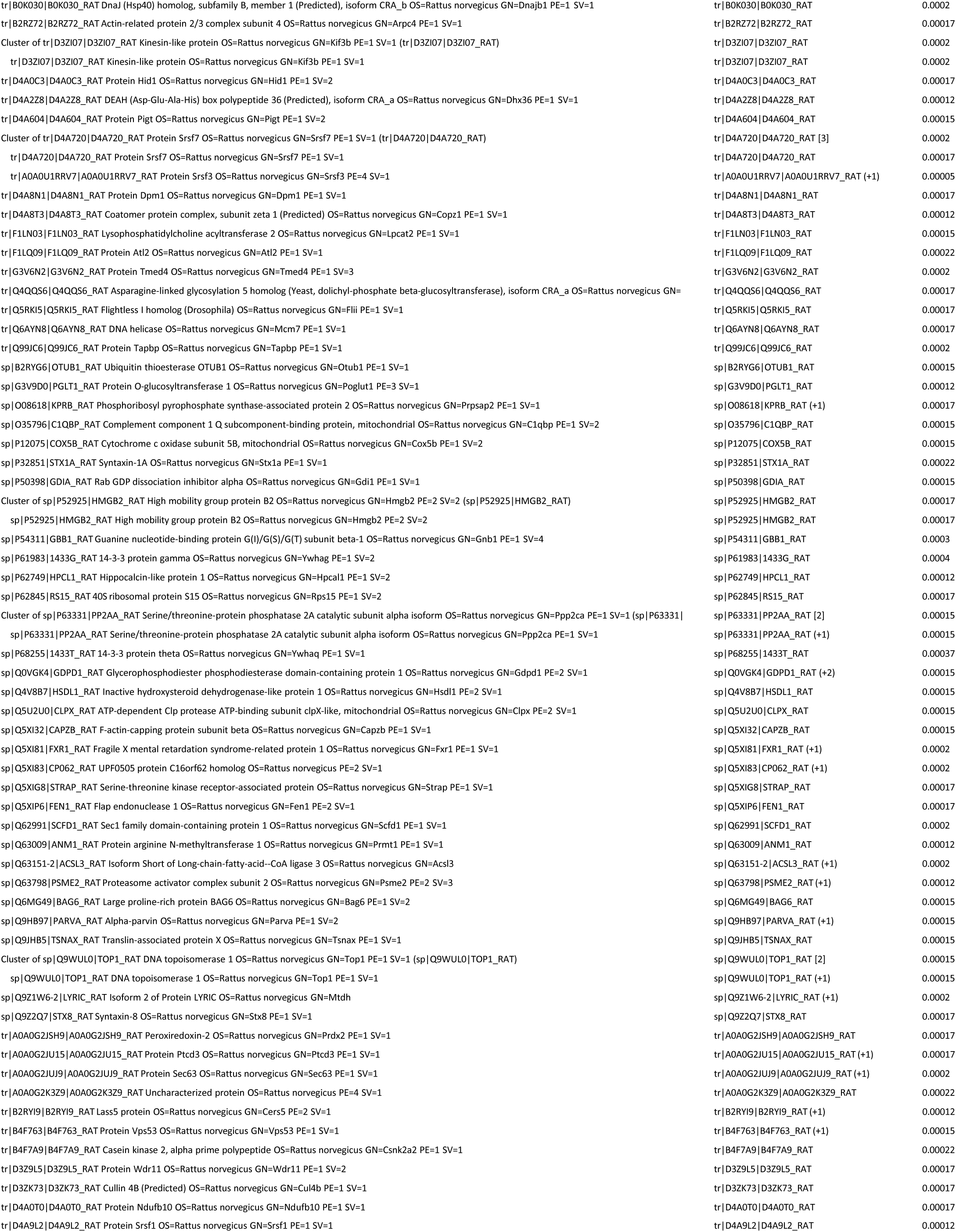
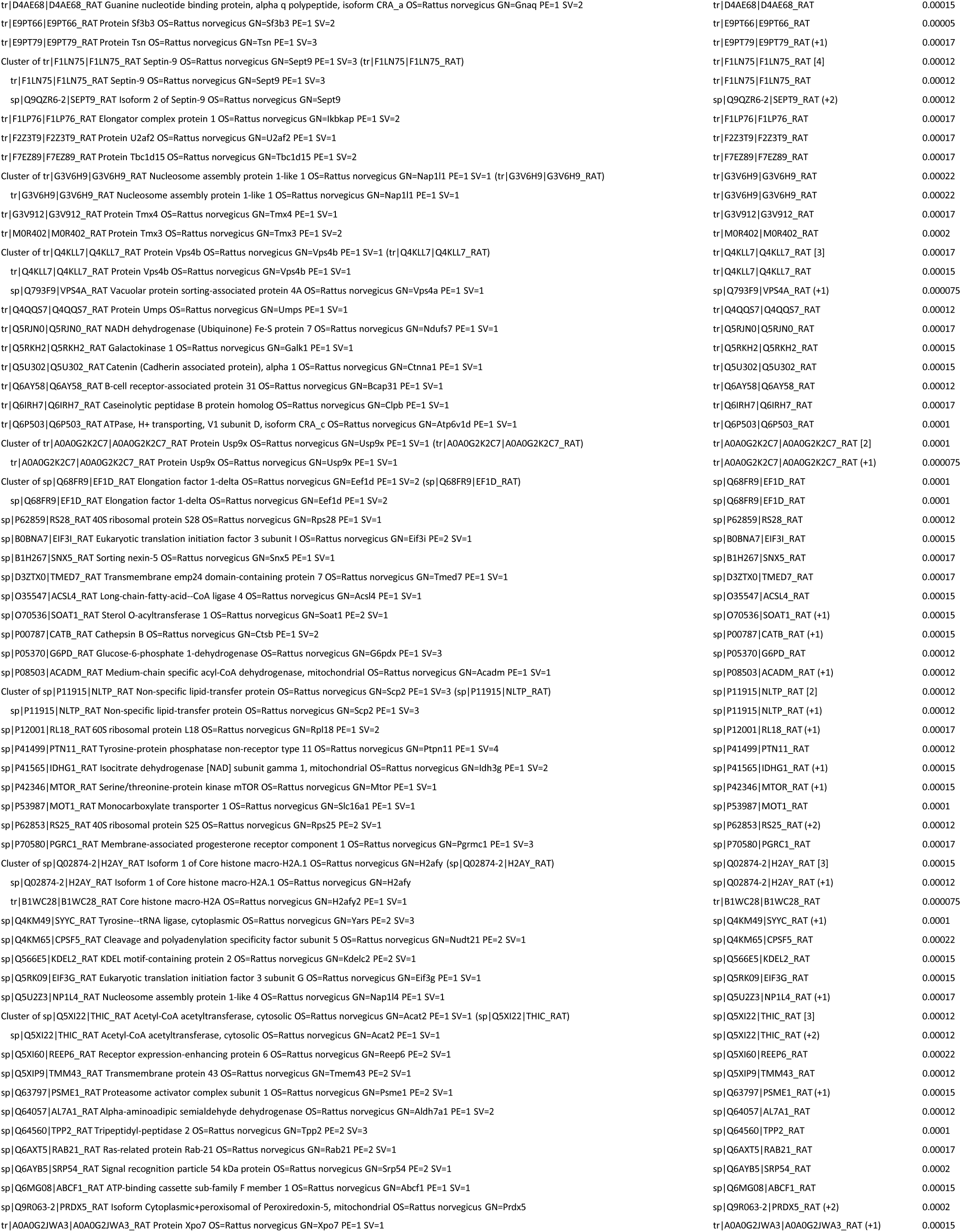
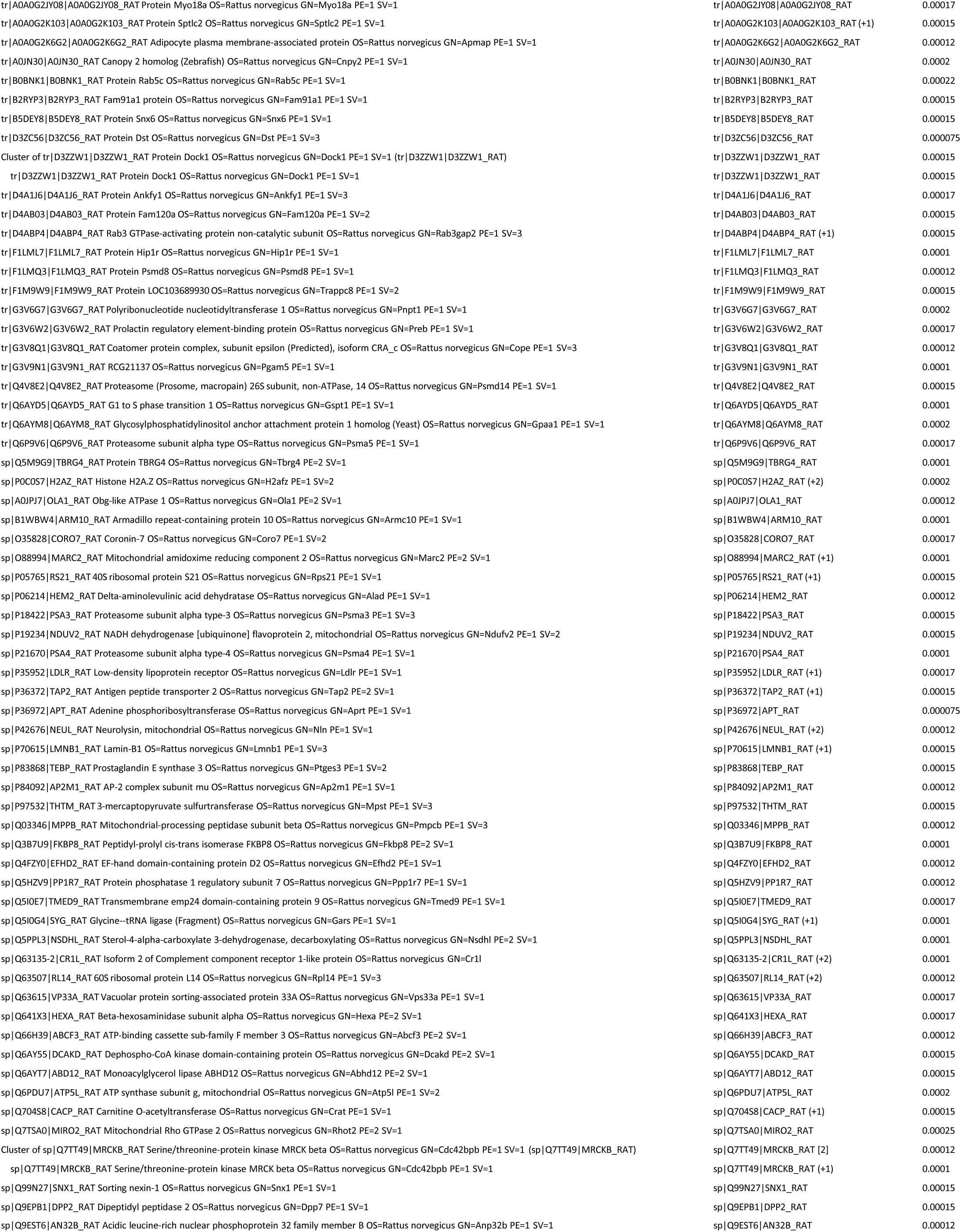
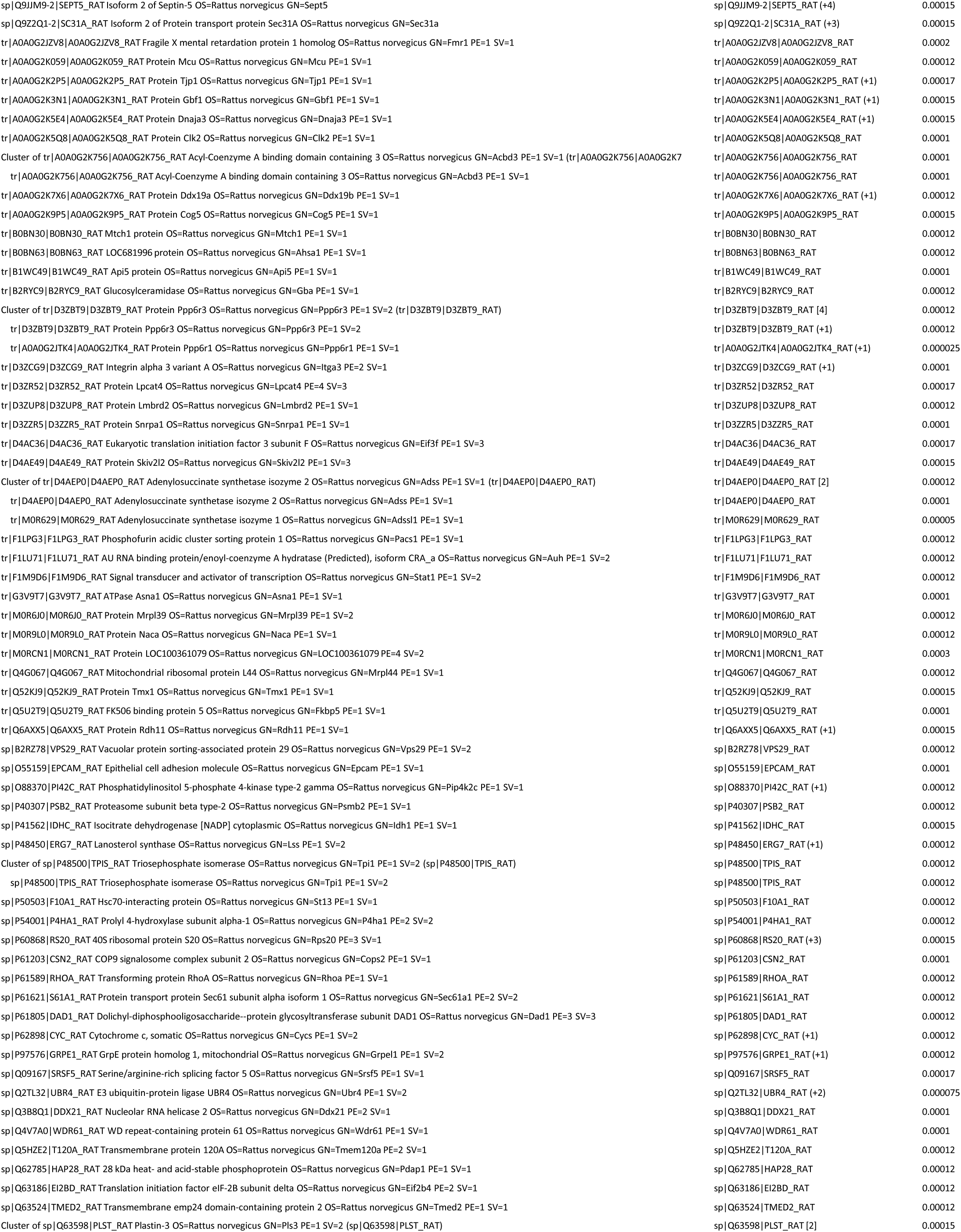
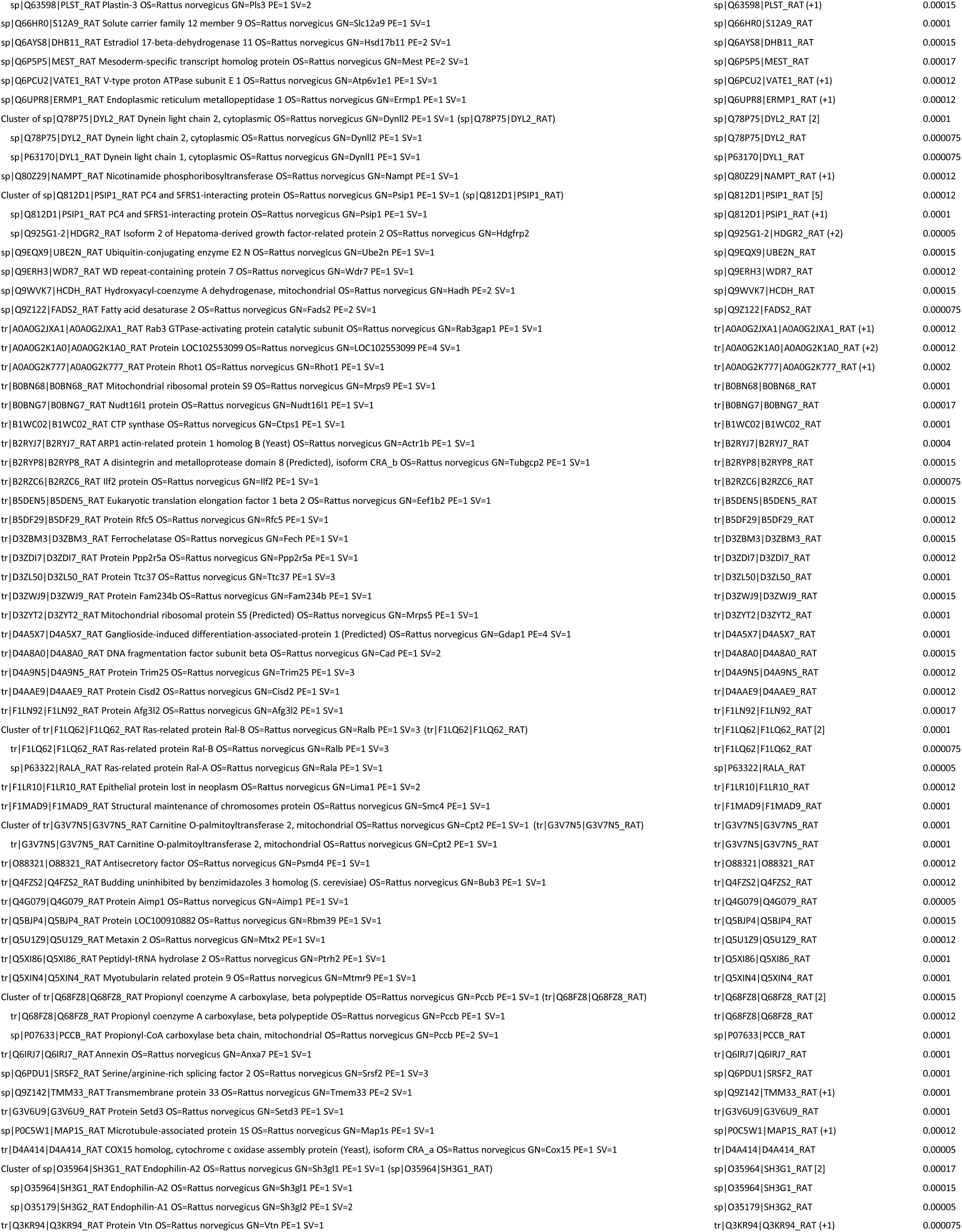
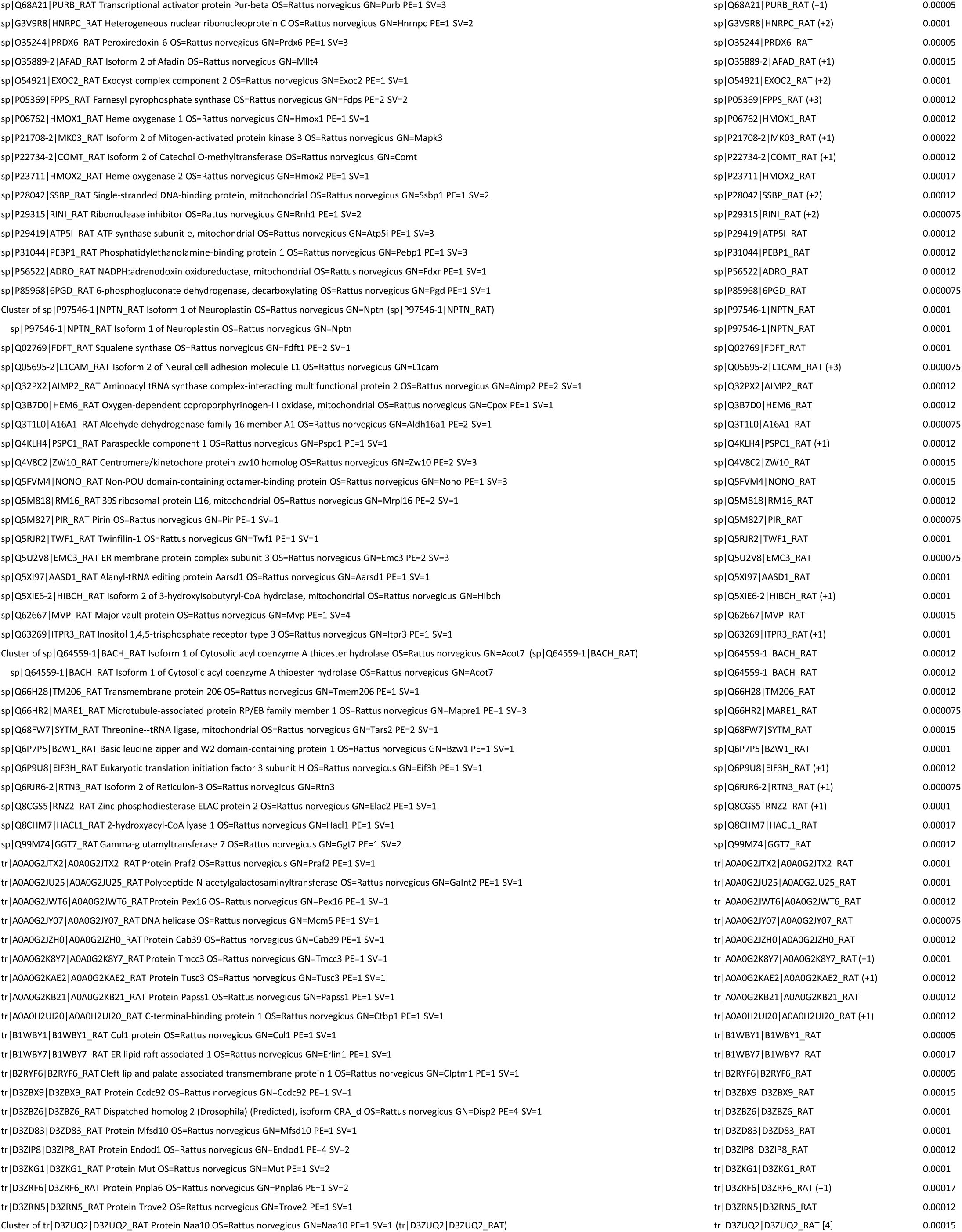
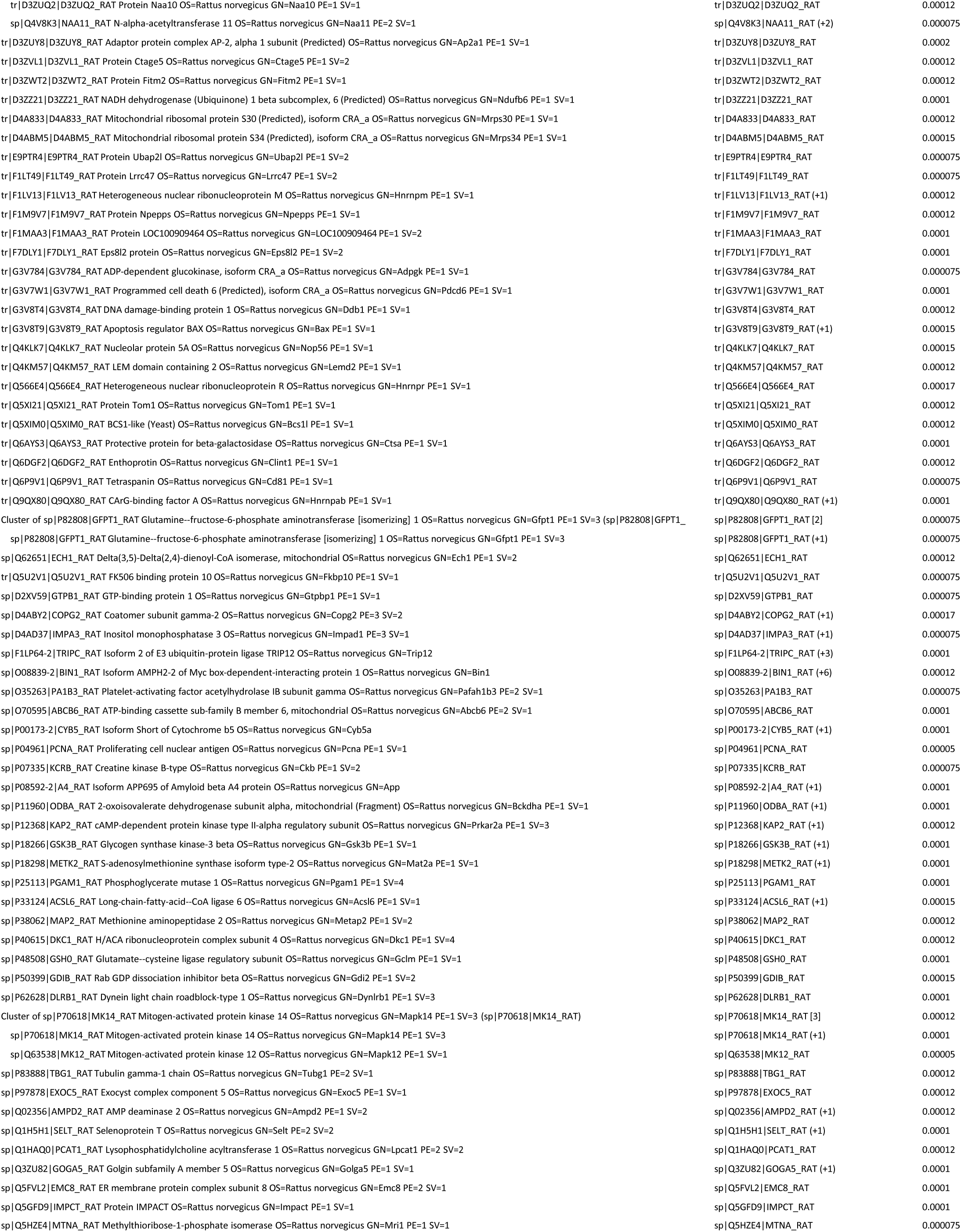
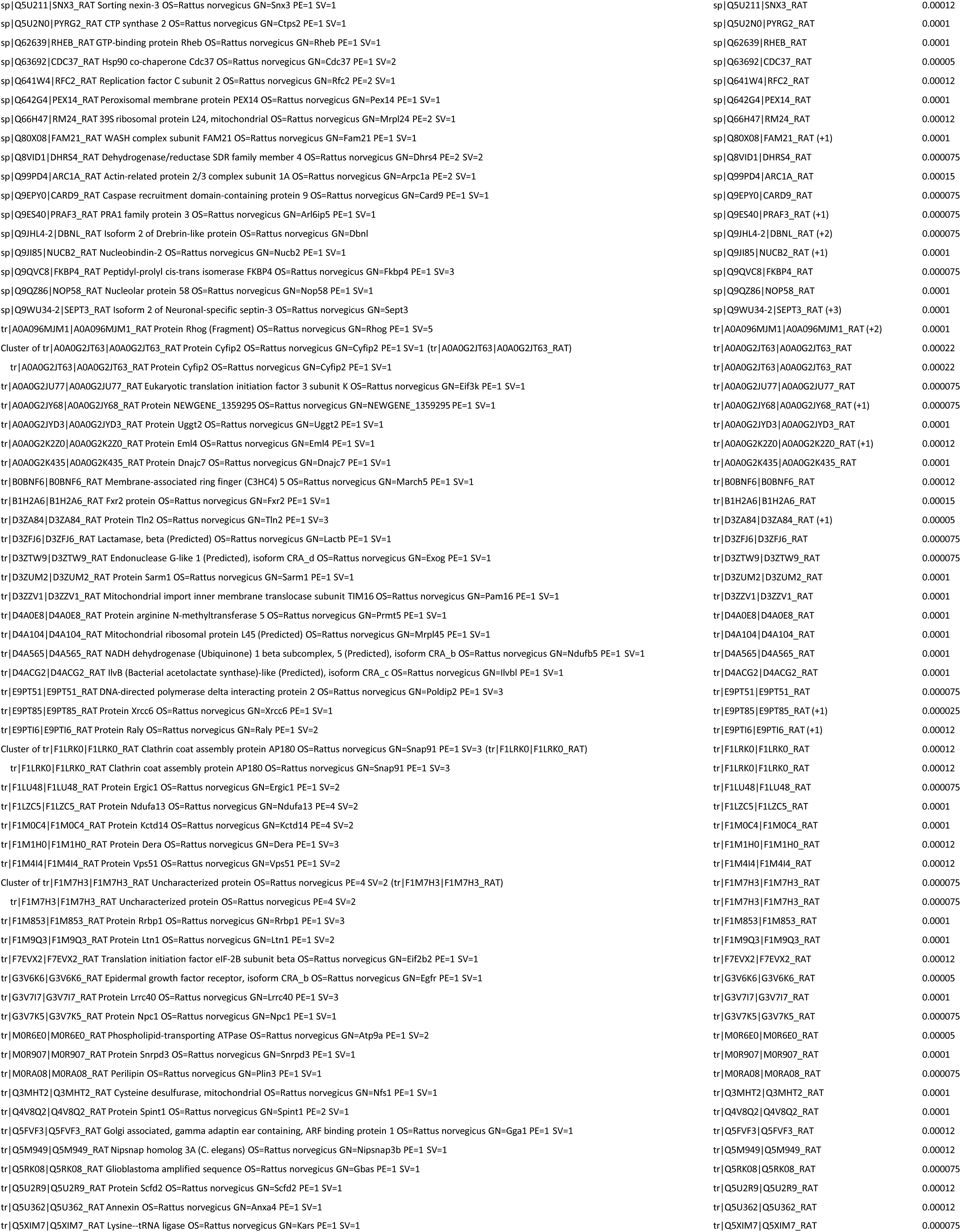
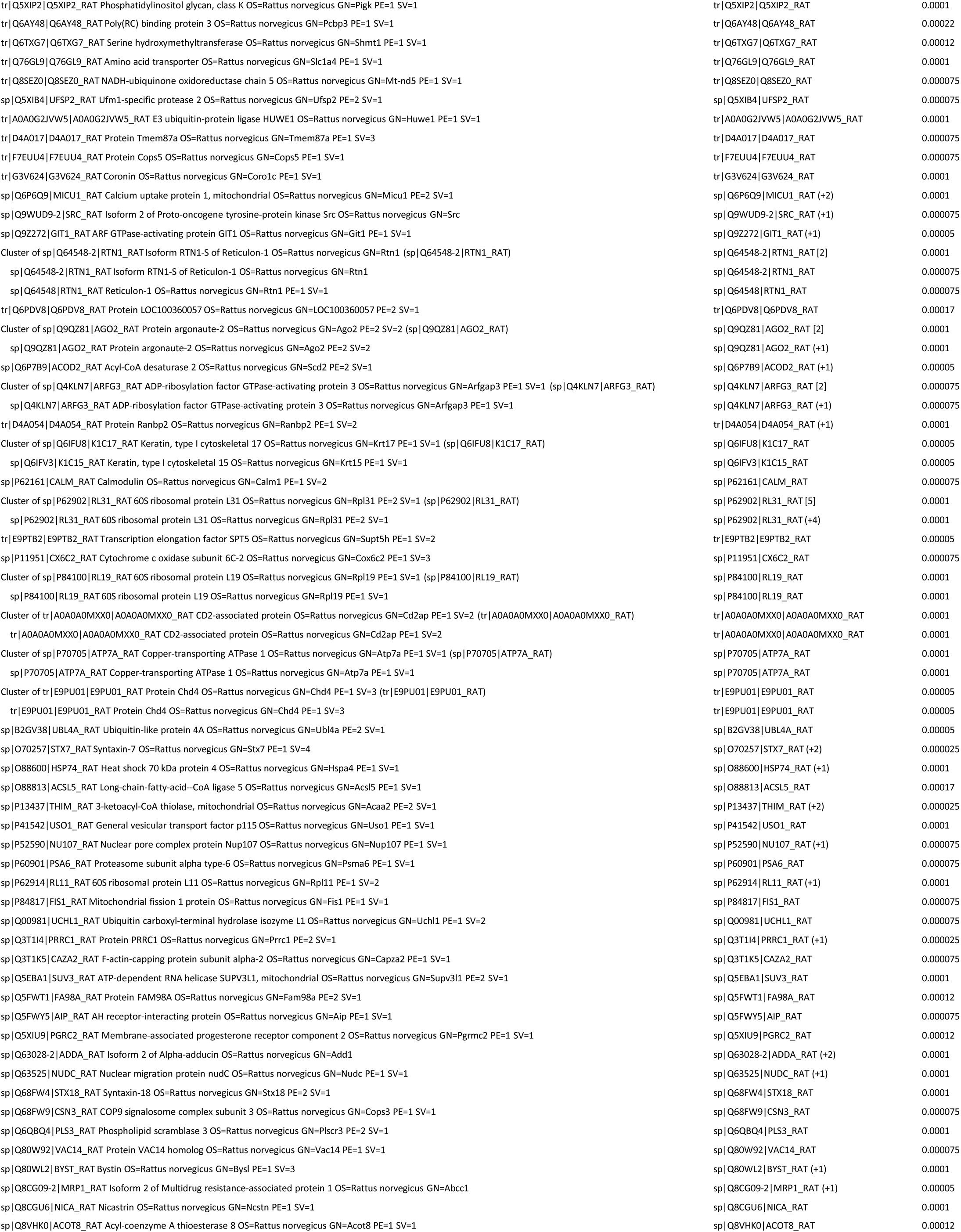
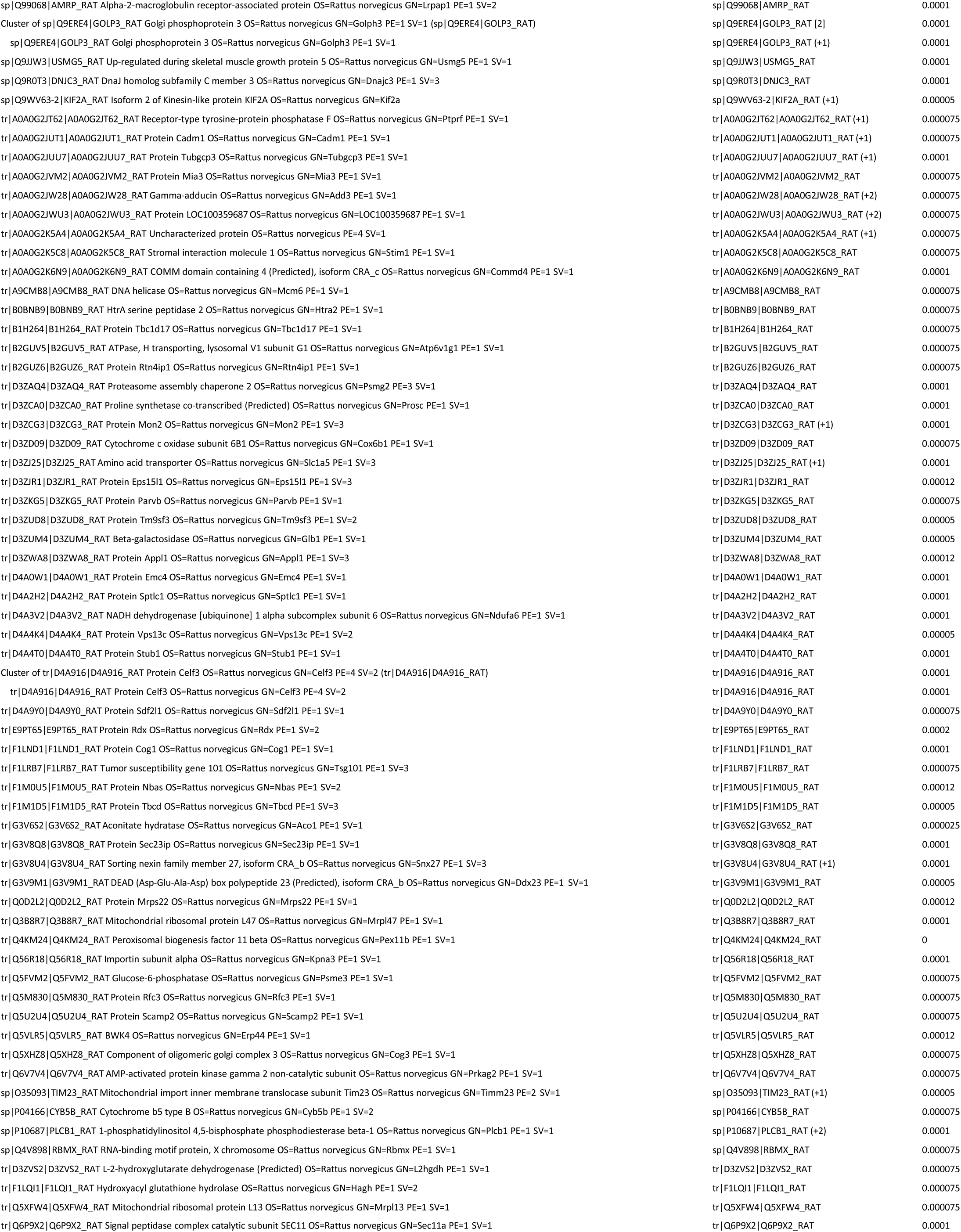
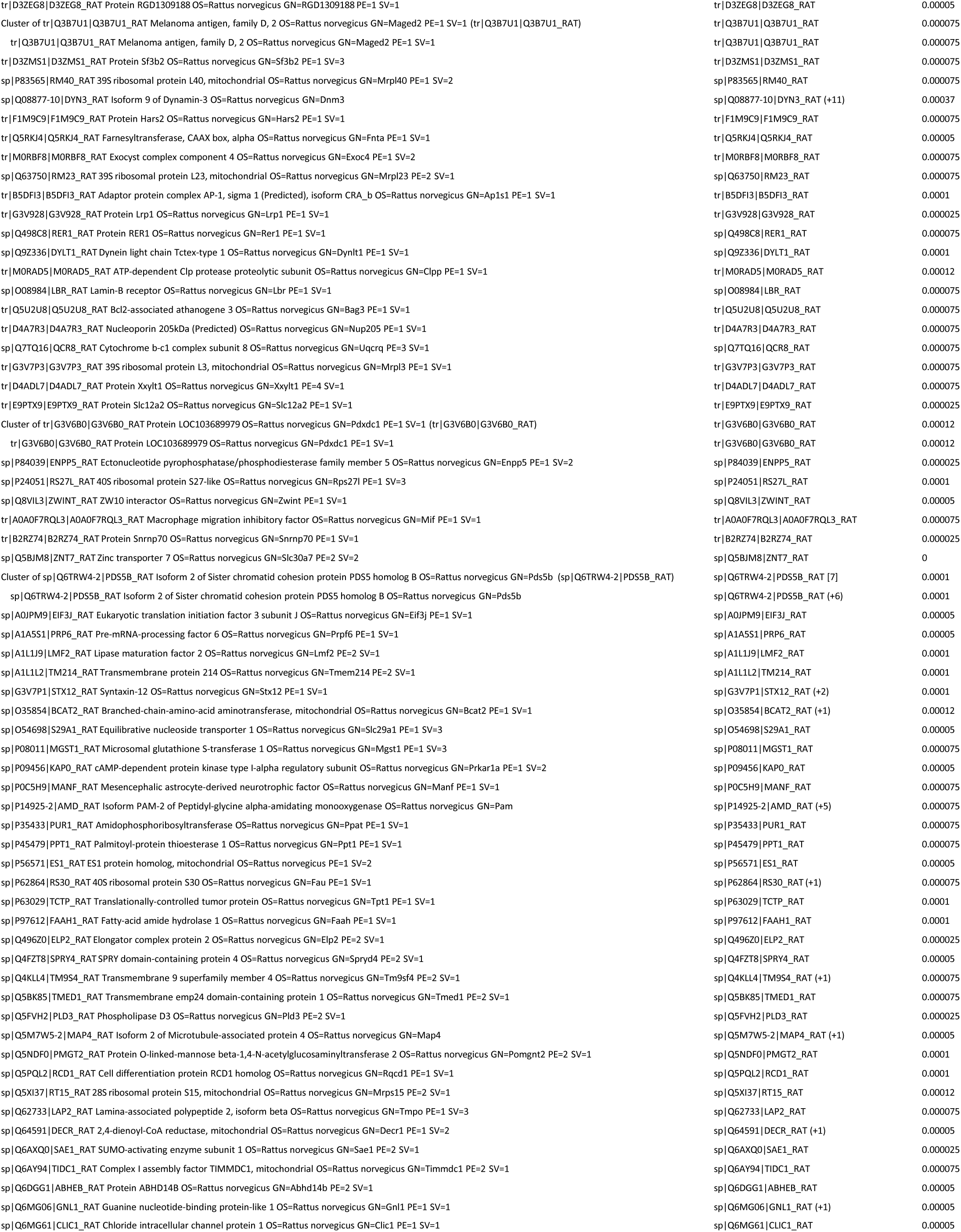
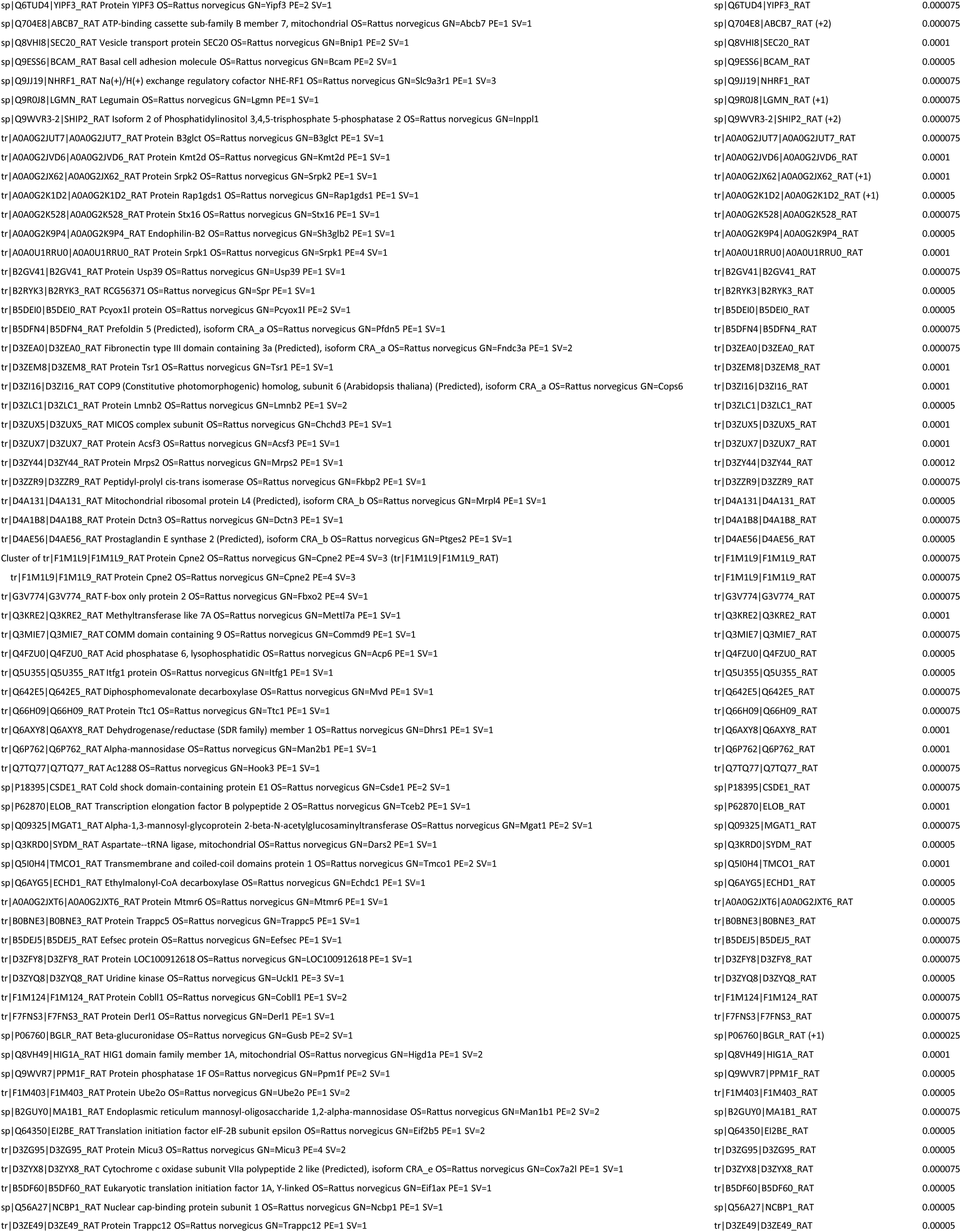
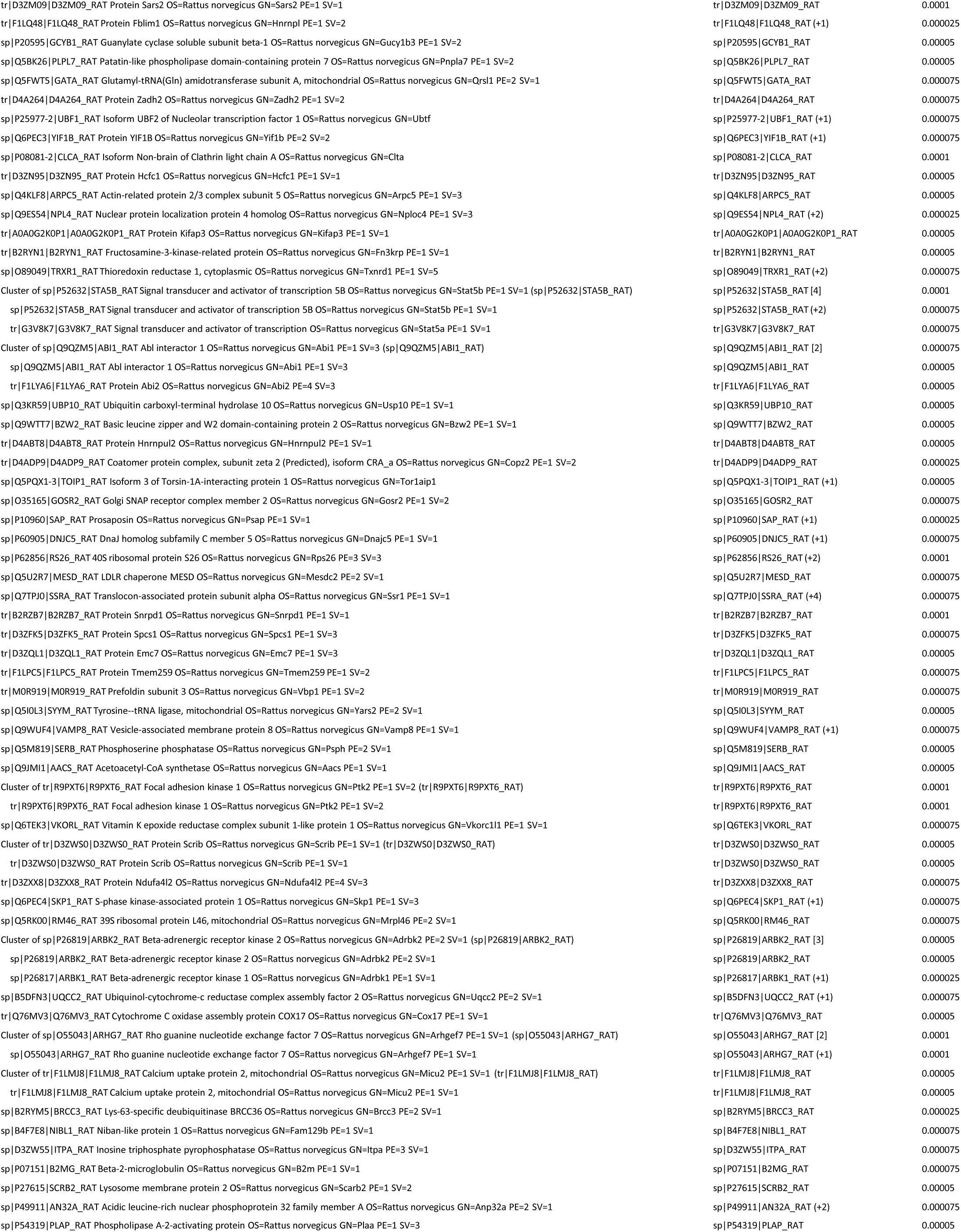
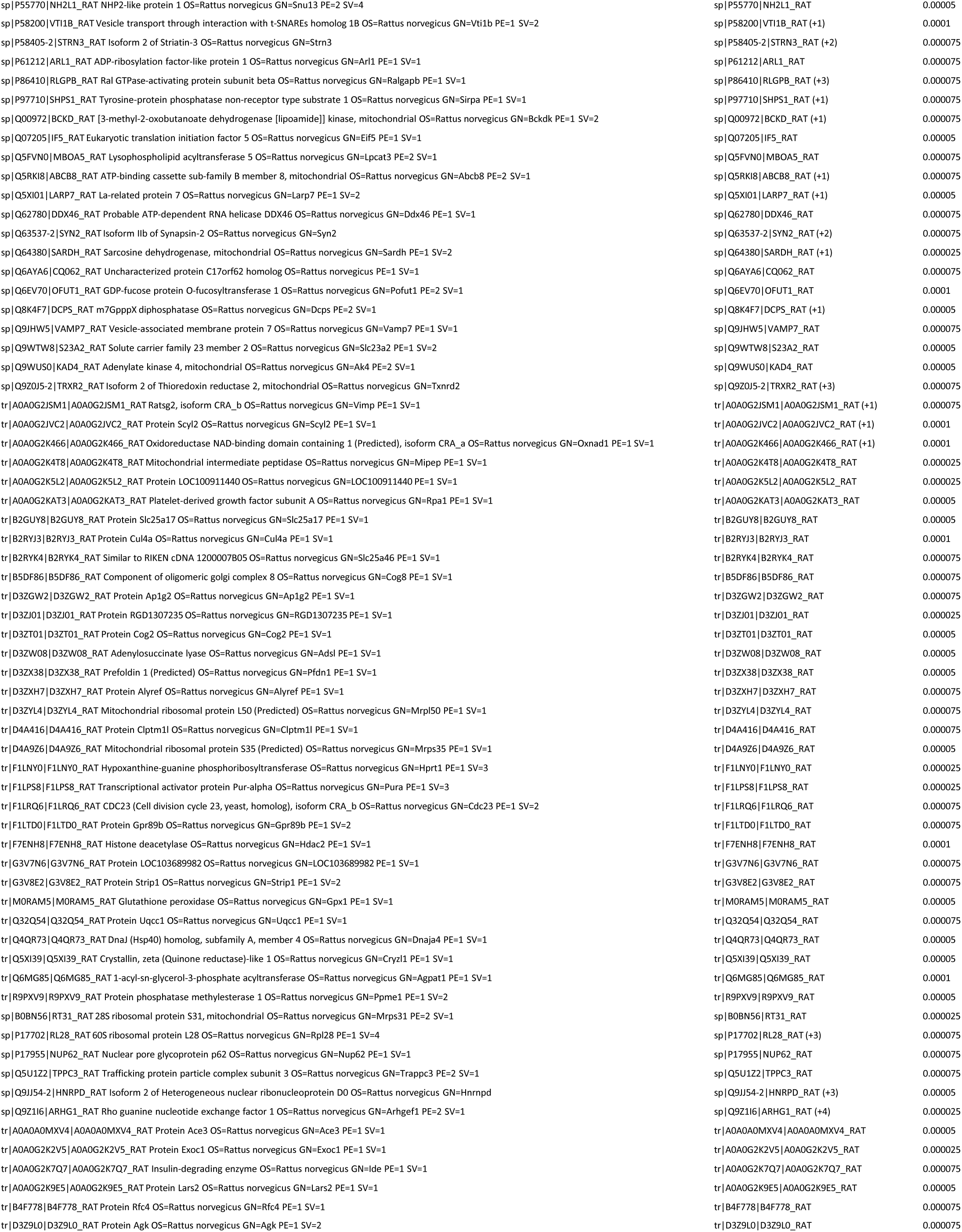
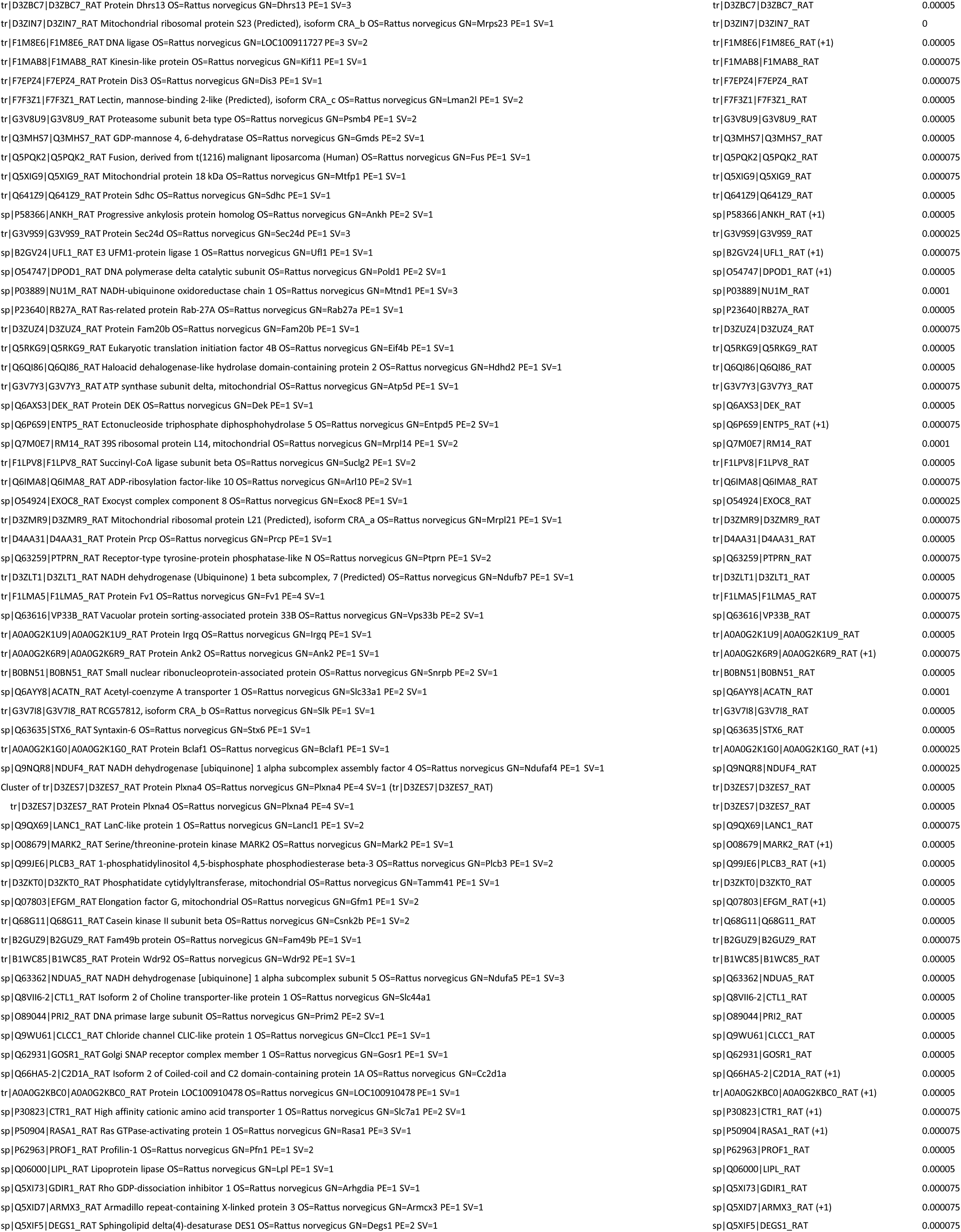
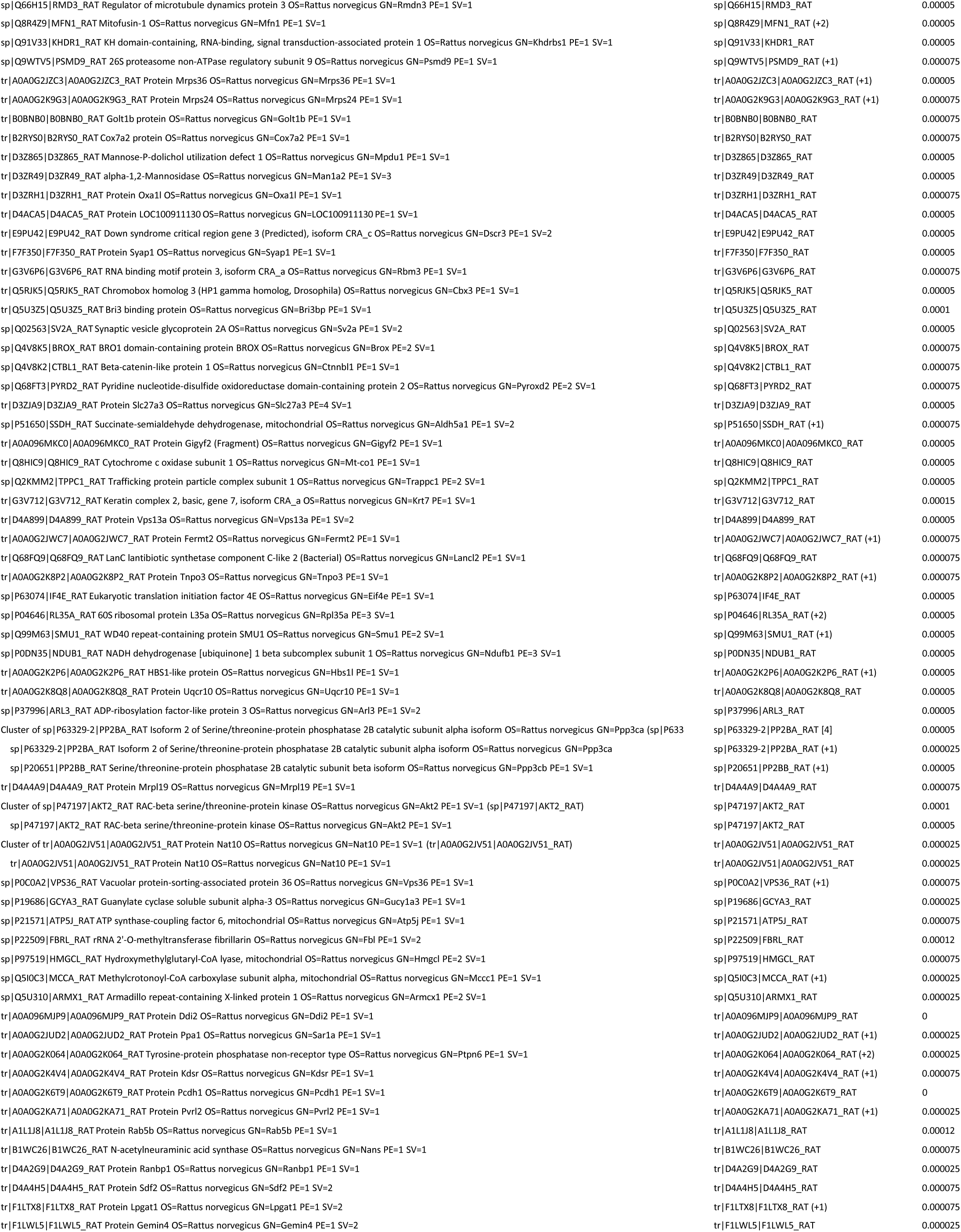
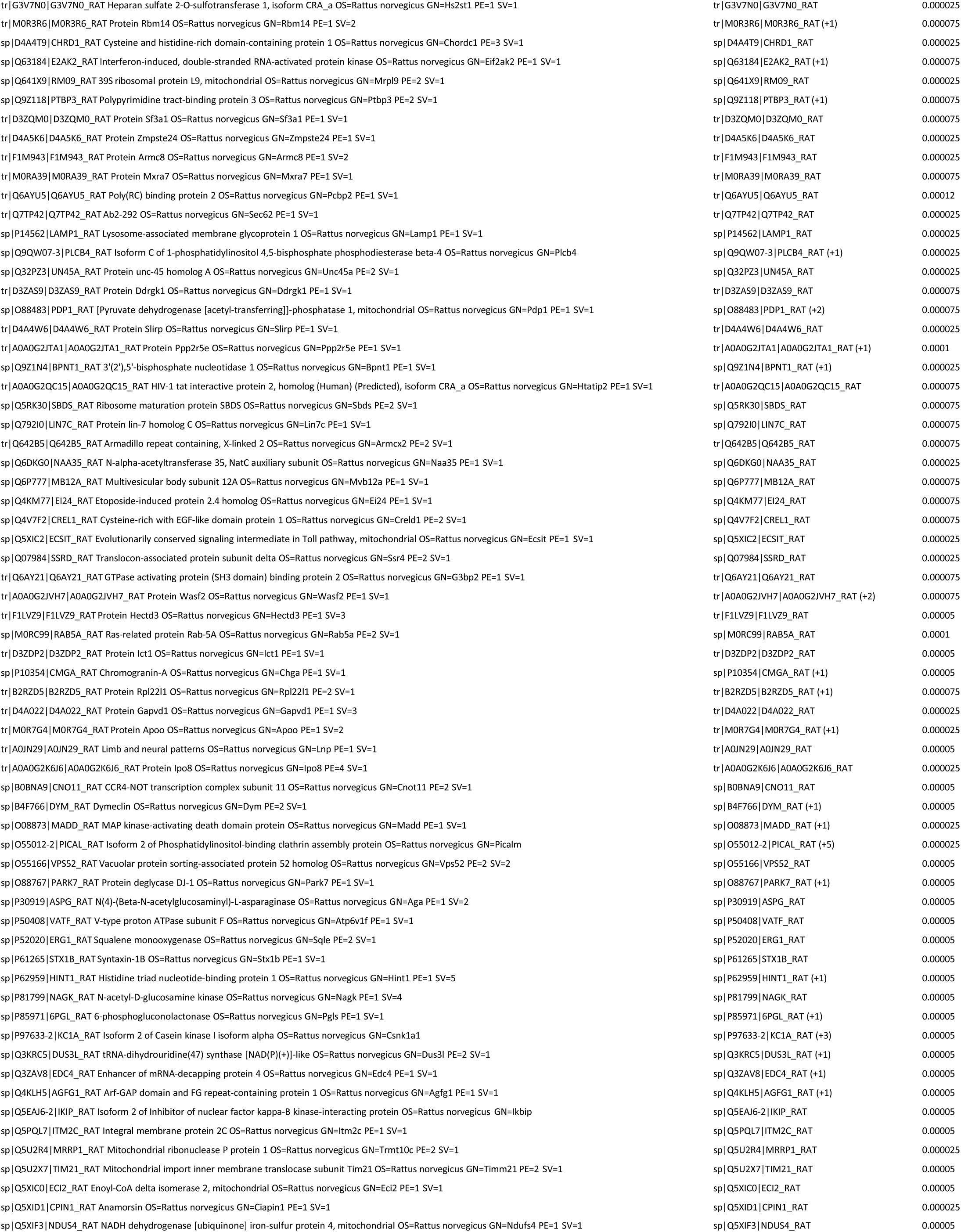
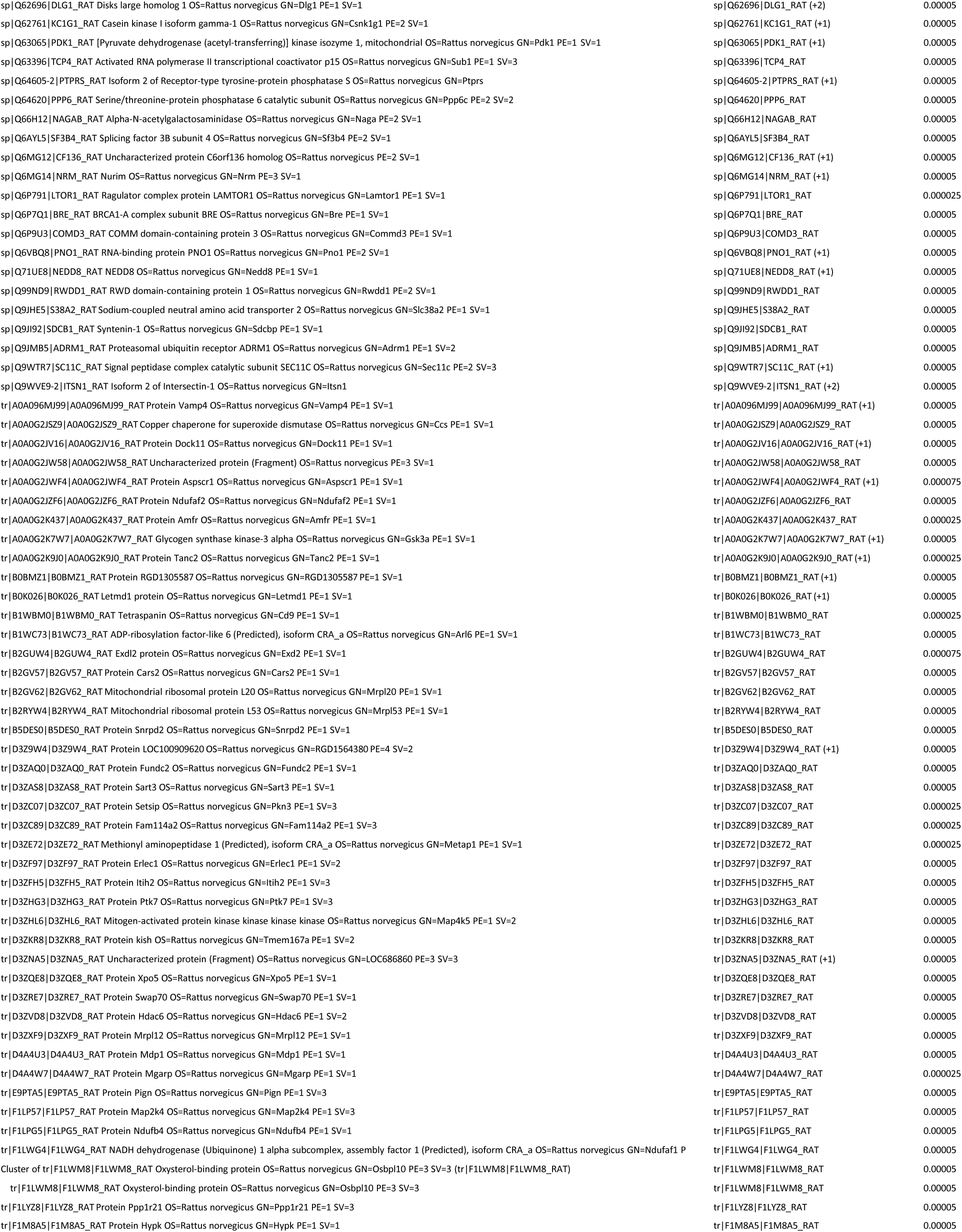
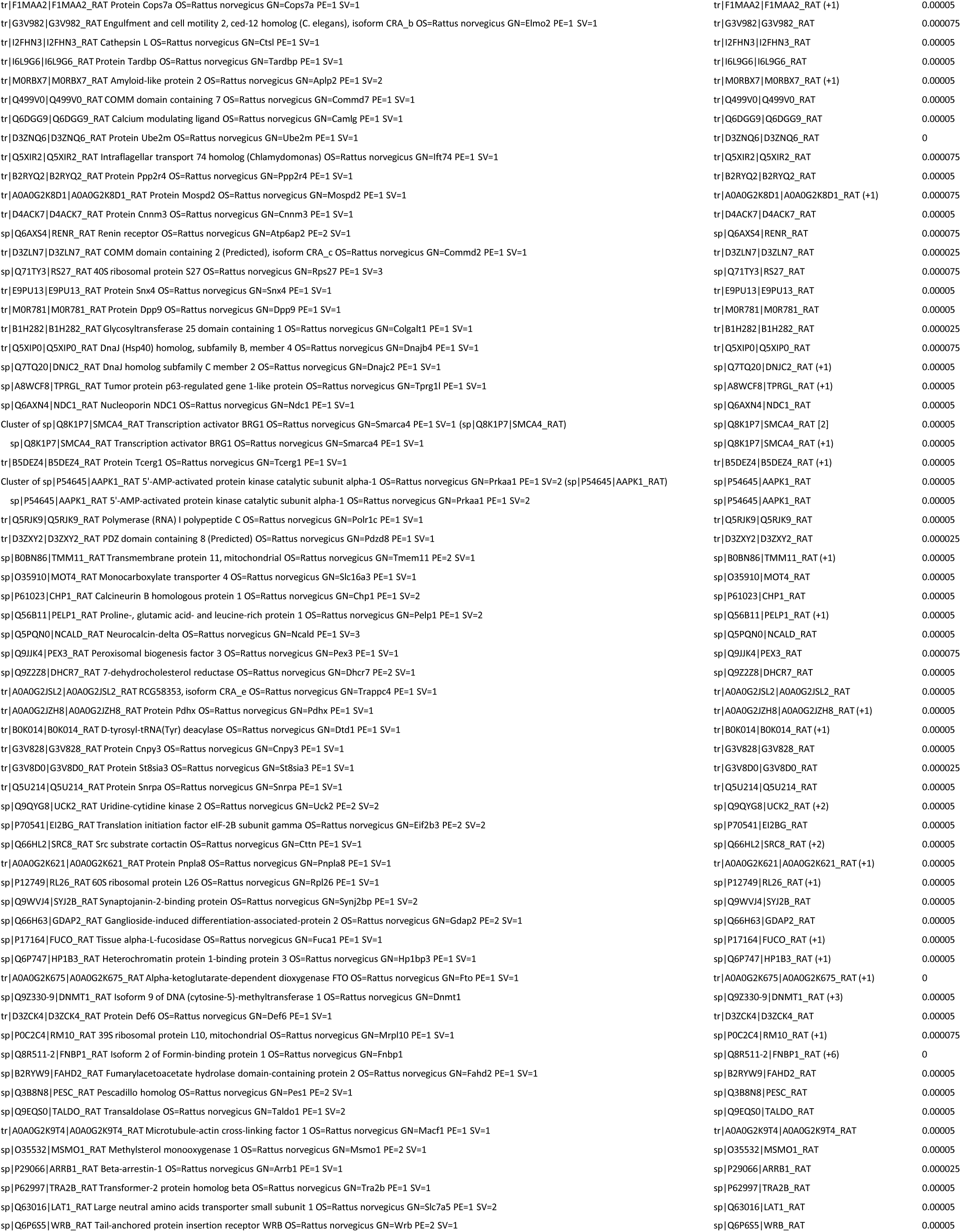
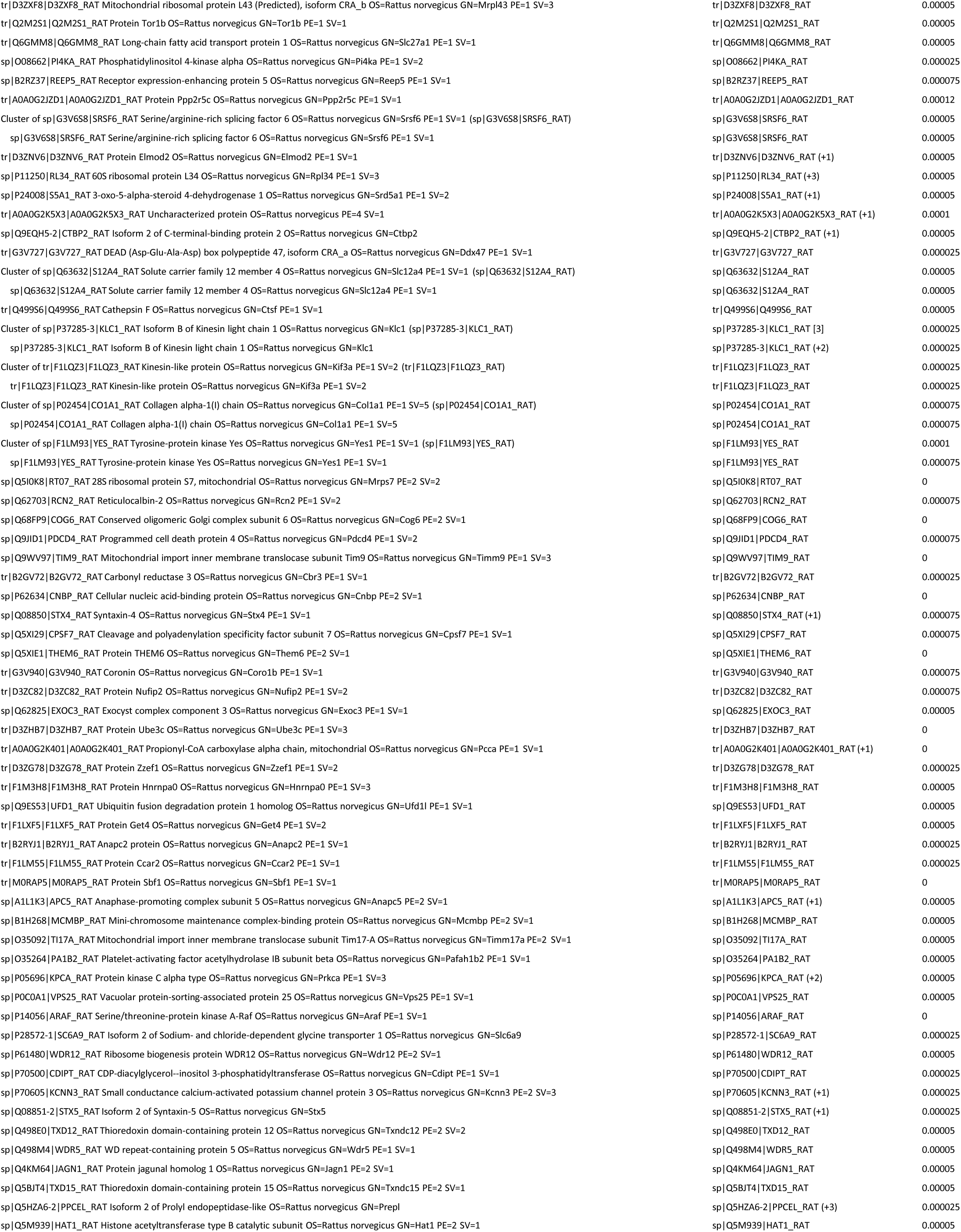
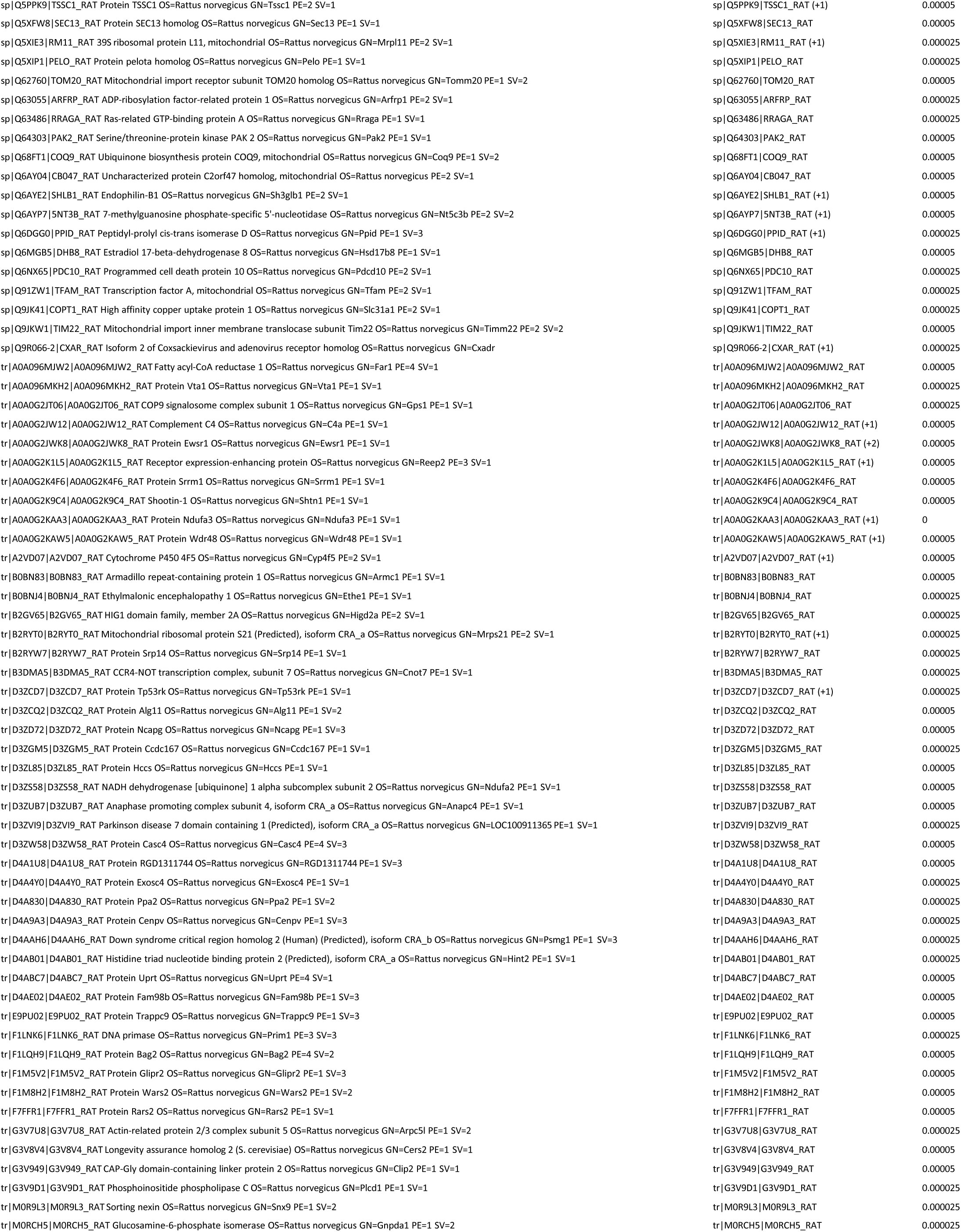
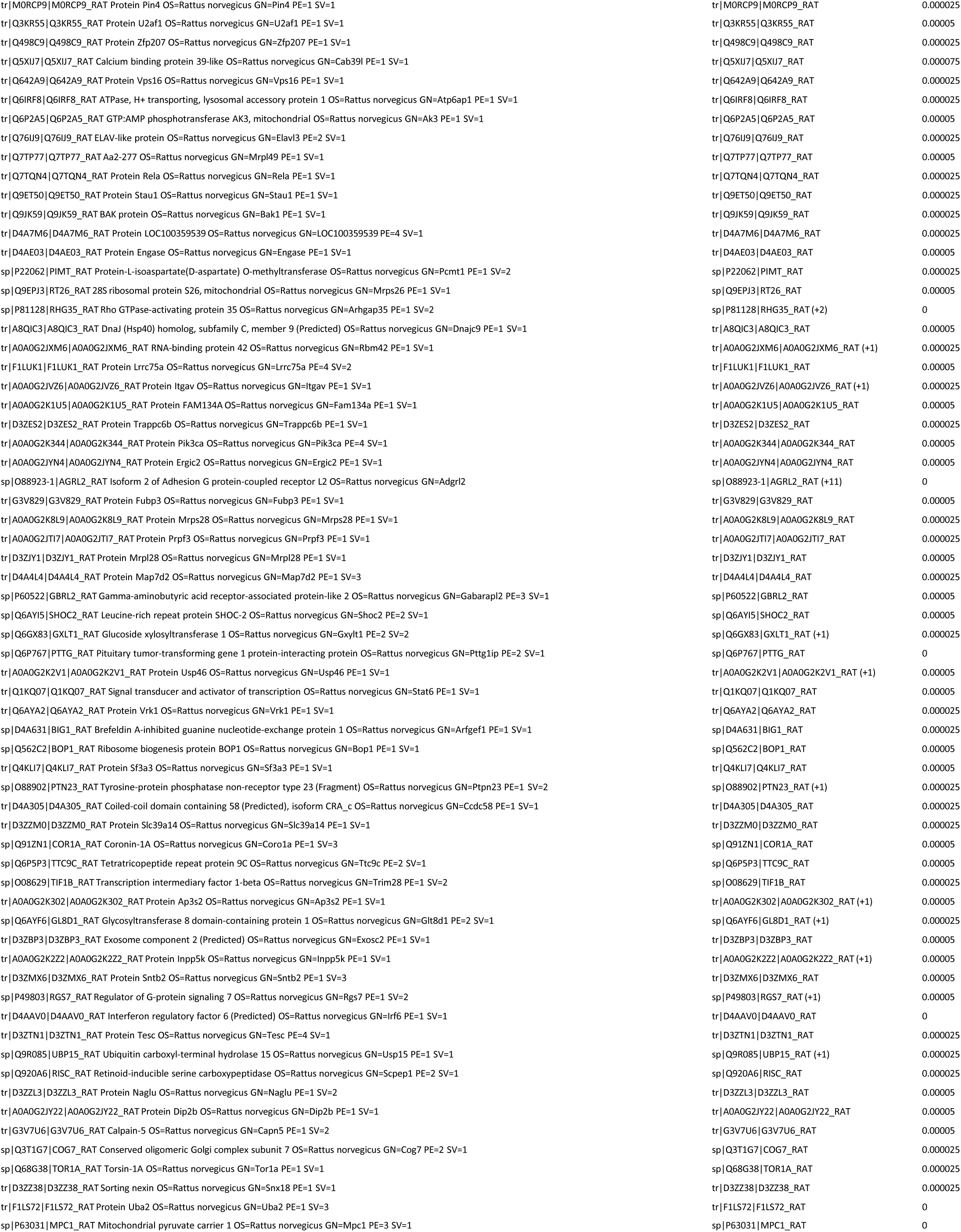
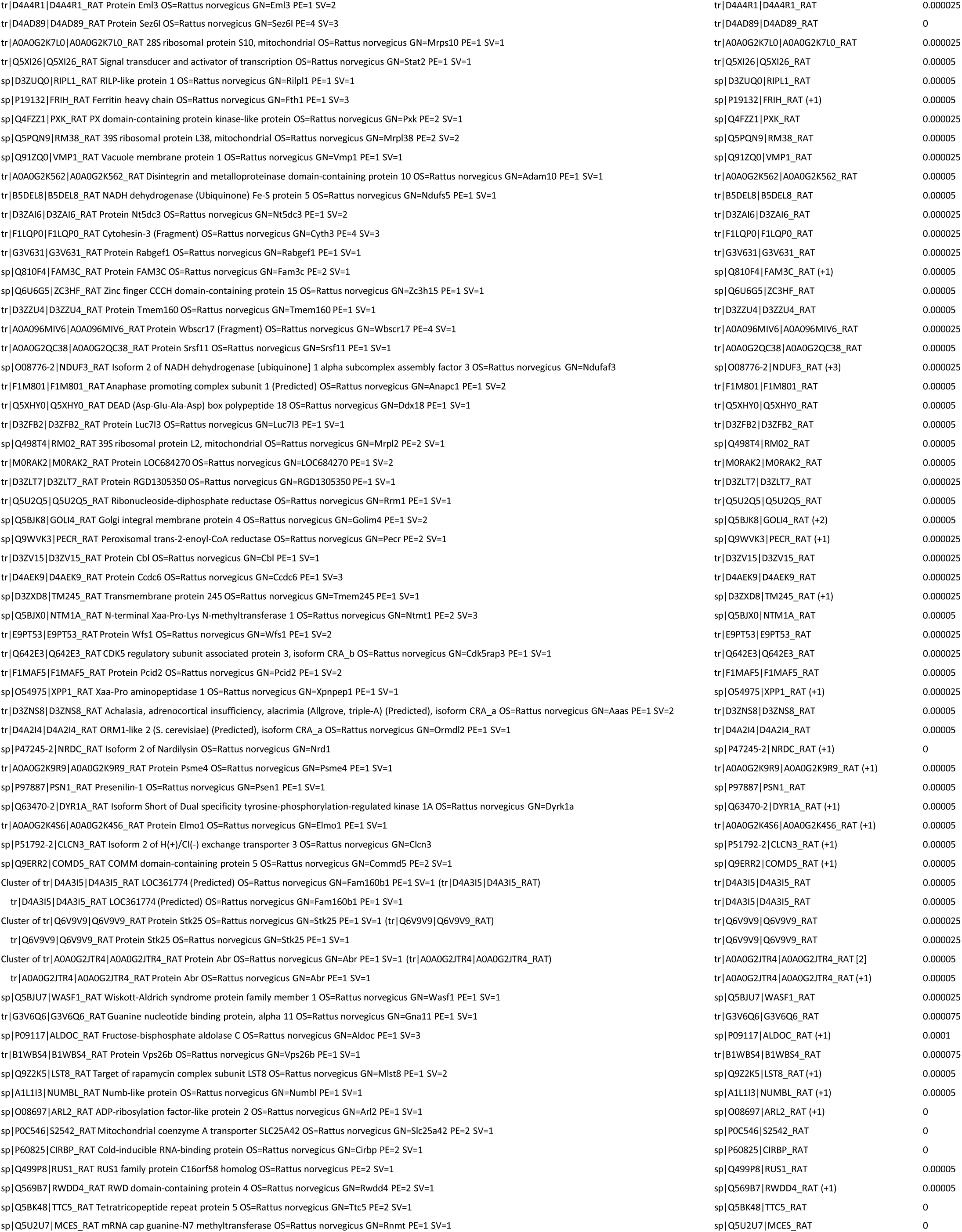
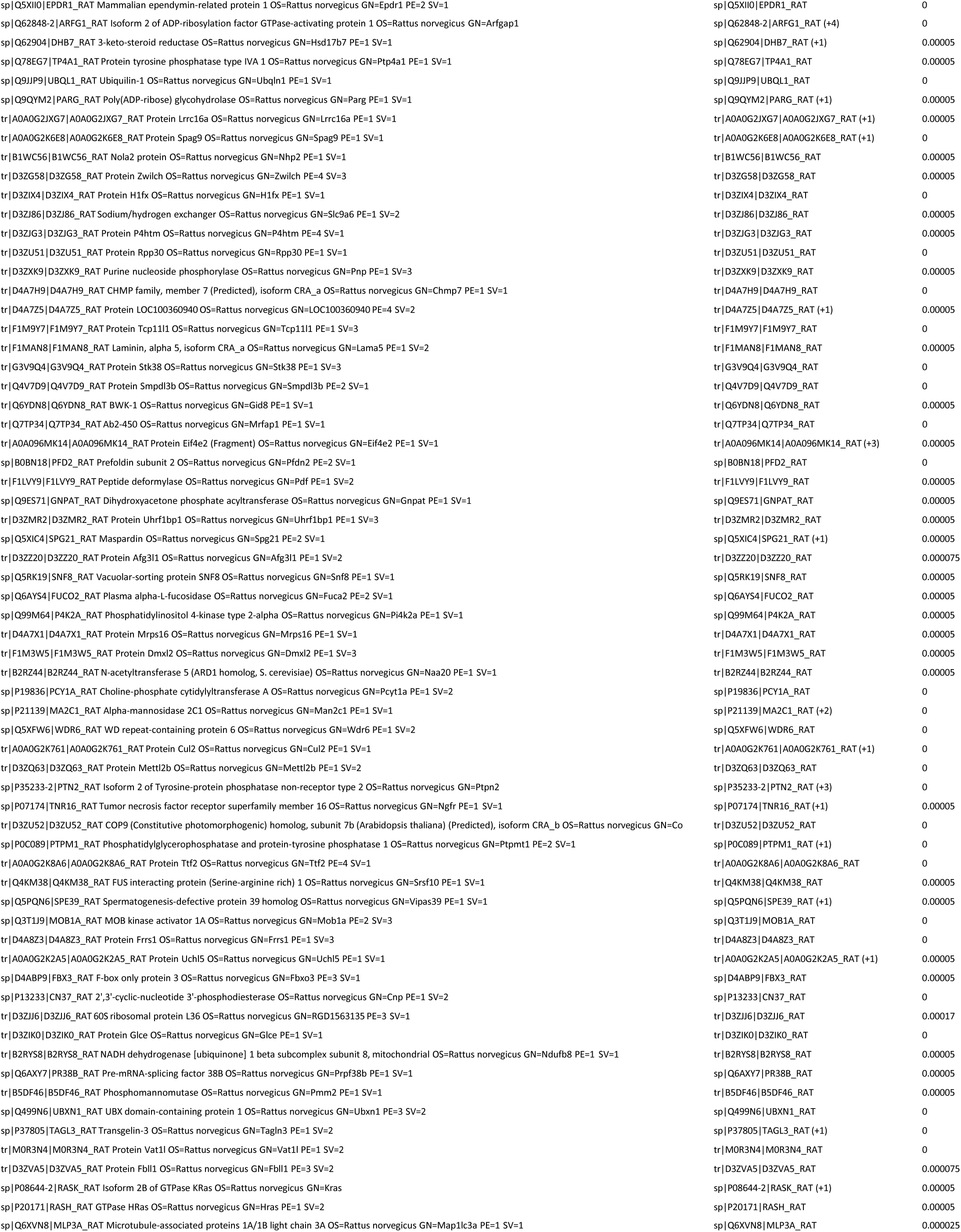
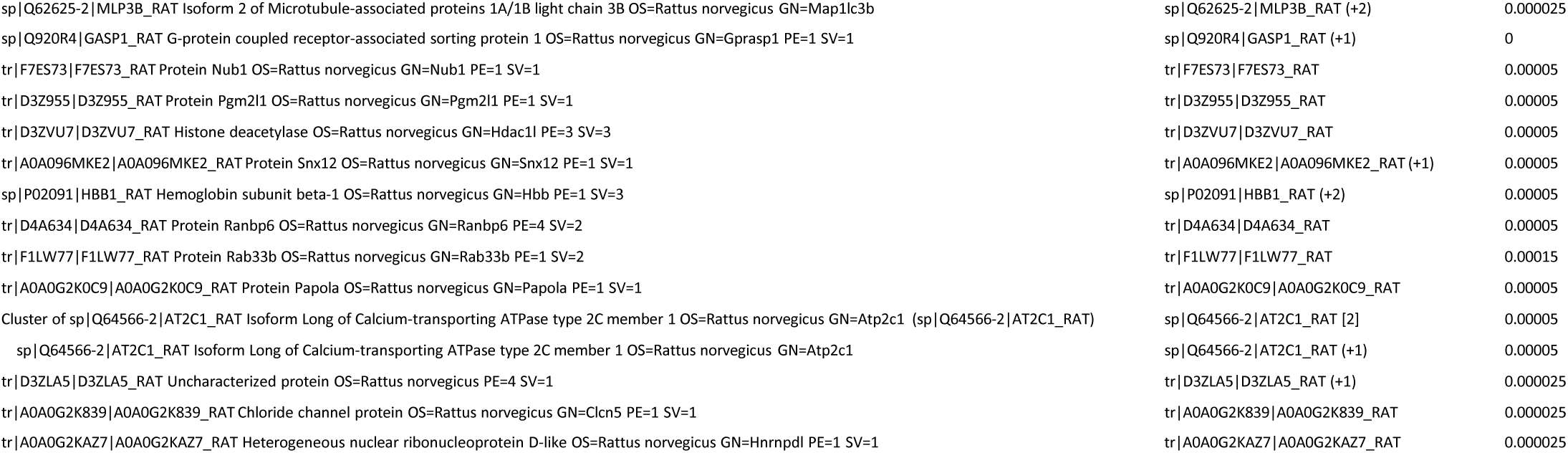
PLCβ1 associated proteins from cytosolic fractions from PC12 cells arrested in the G2/M phase.

**Figure 1D.**
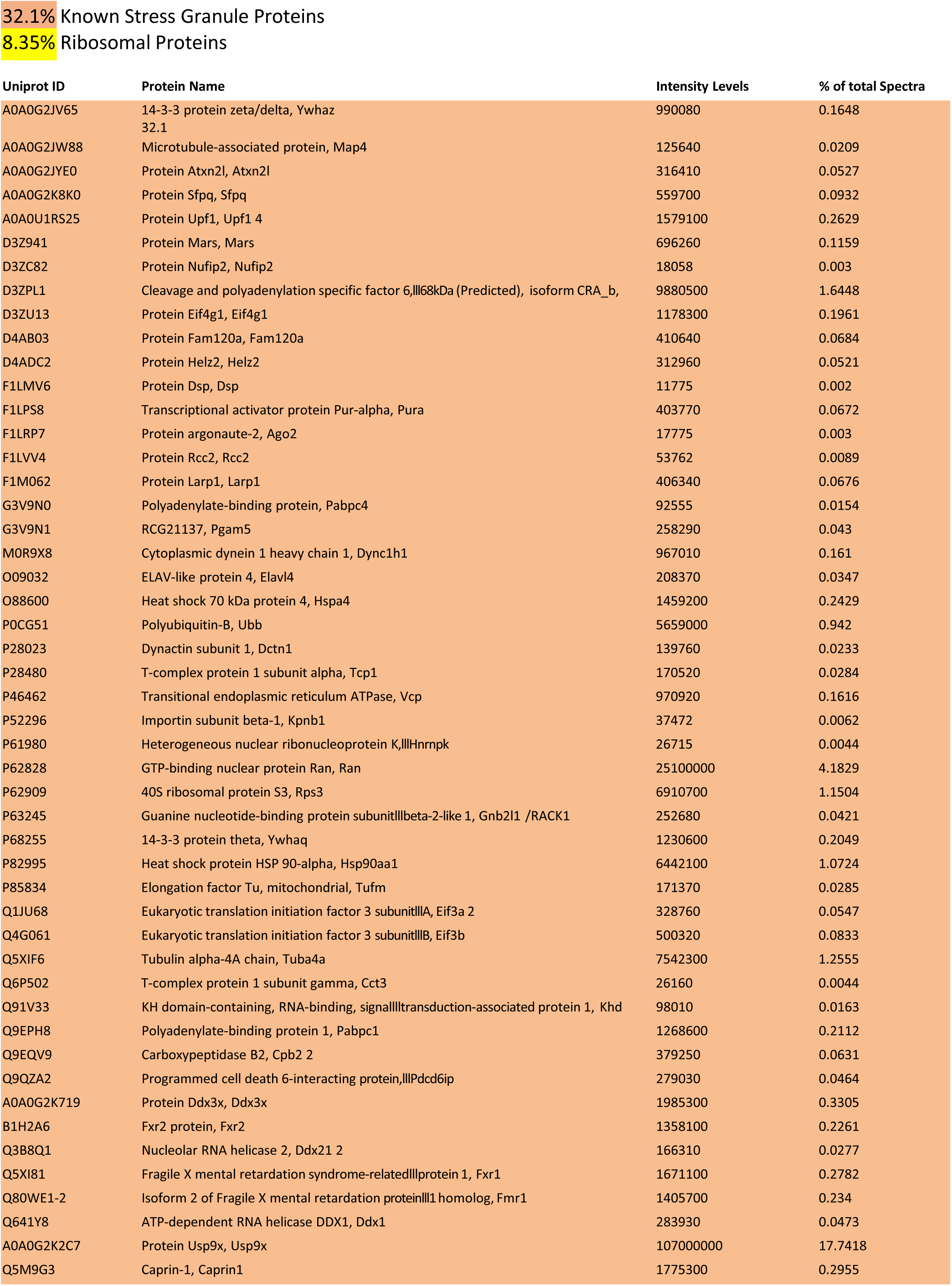
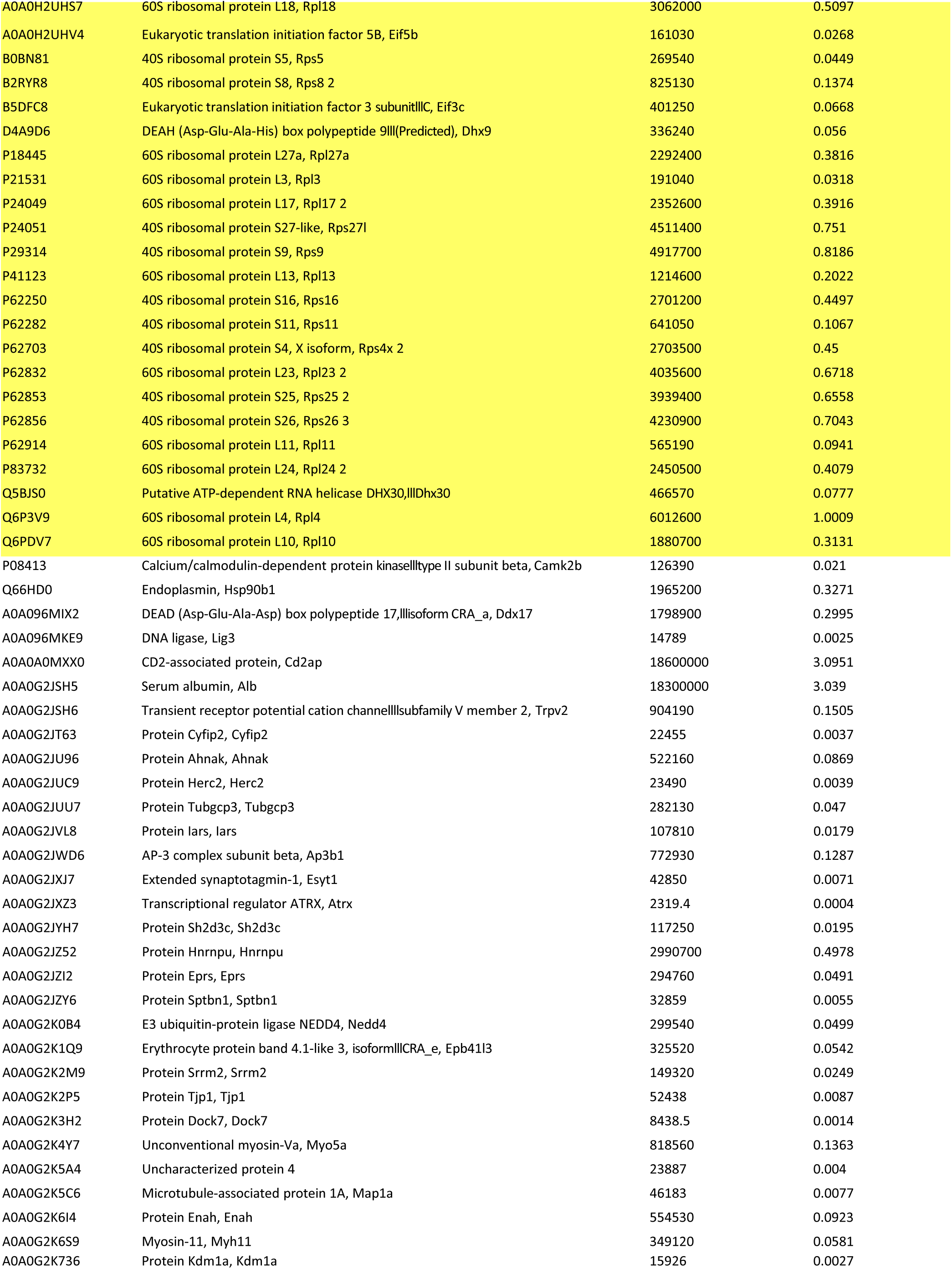
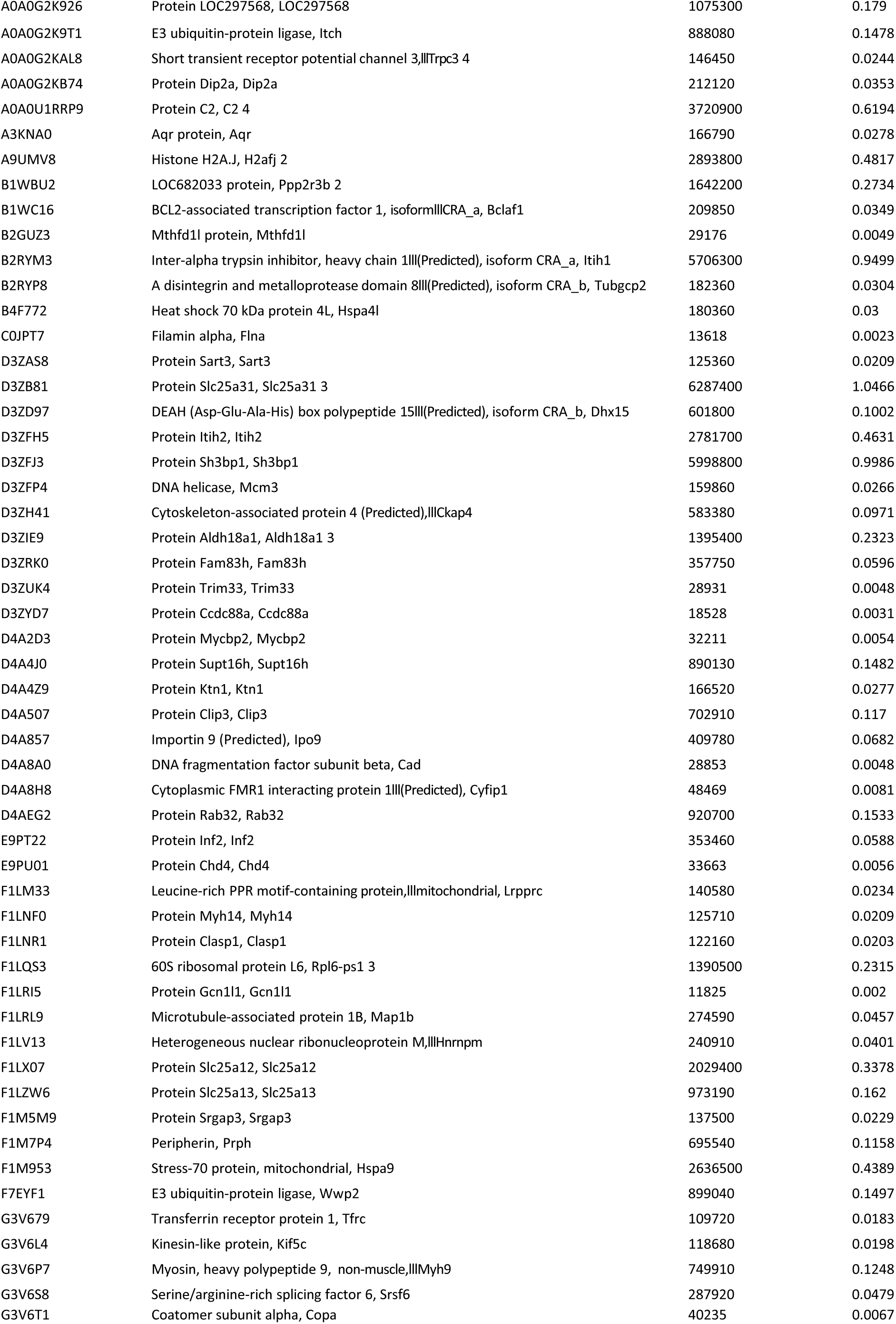
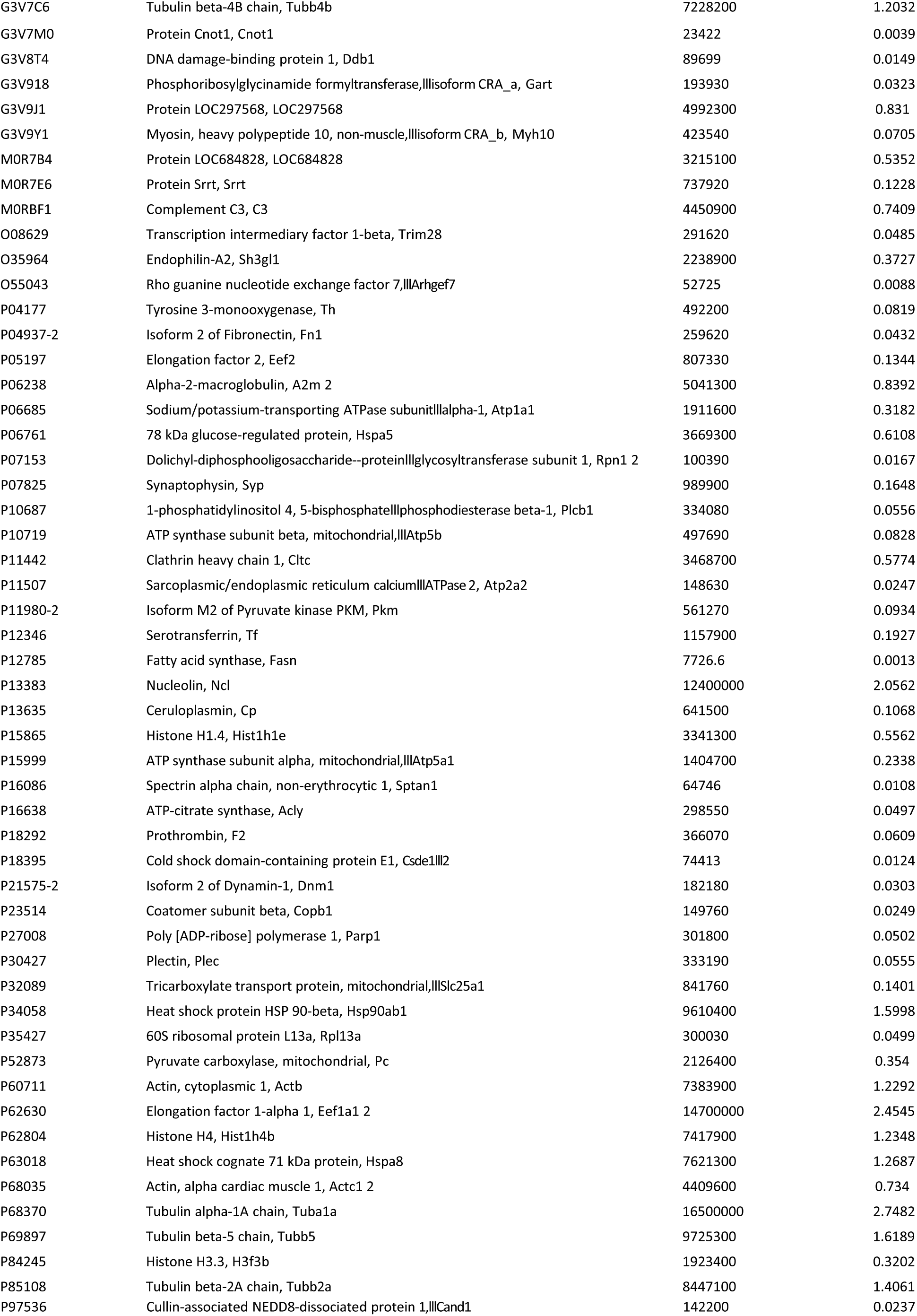
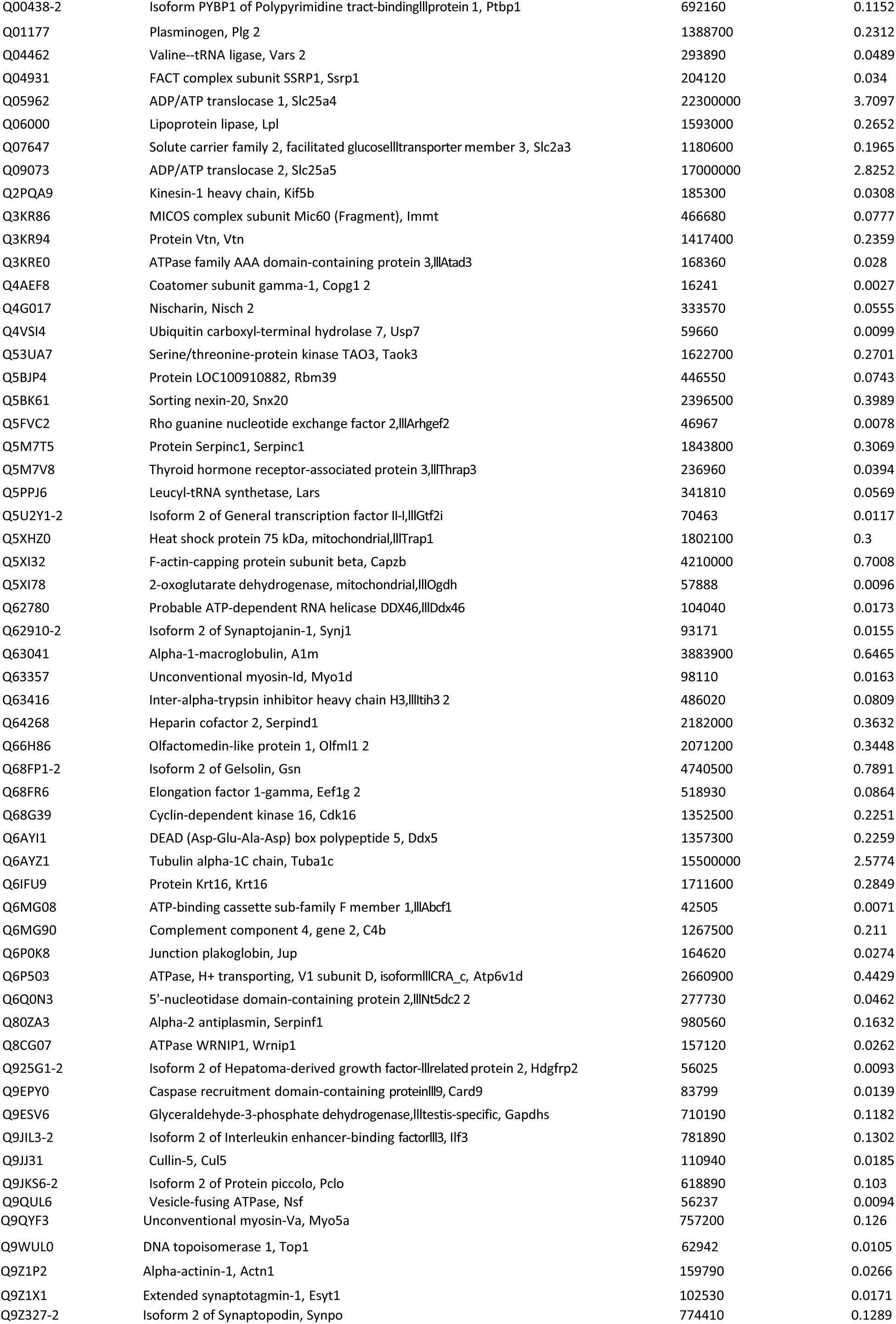
Ago2 associated proteins from cytosolic fractions from unsynchronized PC12 cells.

**Figure 2A.**
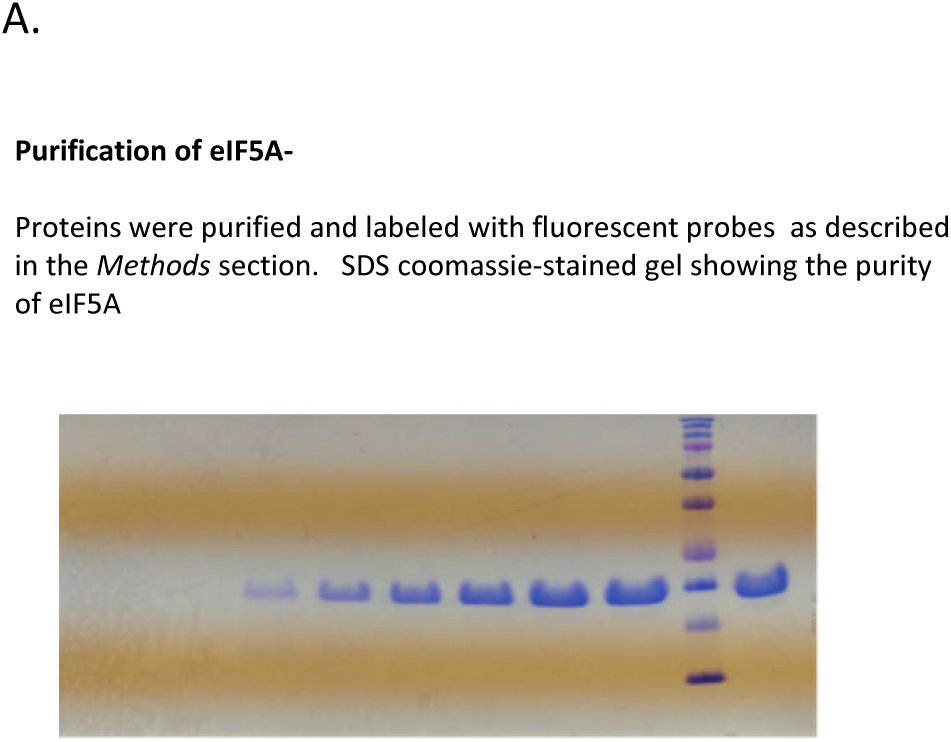
Gel showing purity of eIF5A.

**Figure 2B-C.**
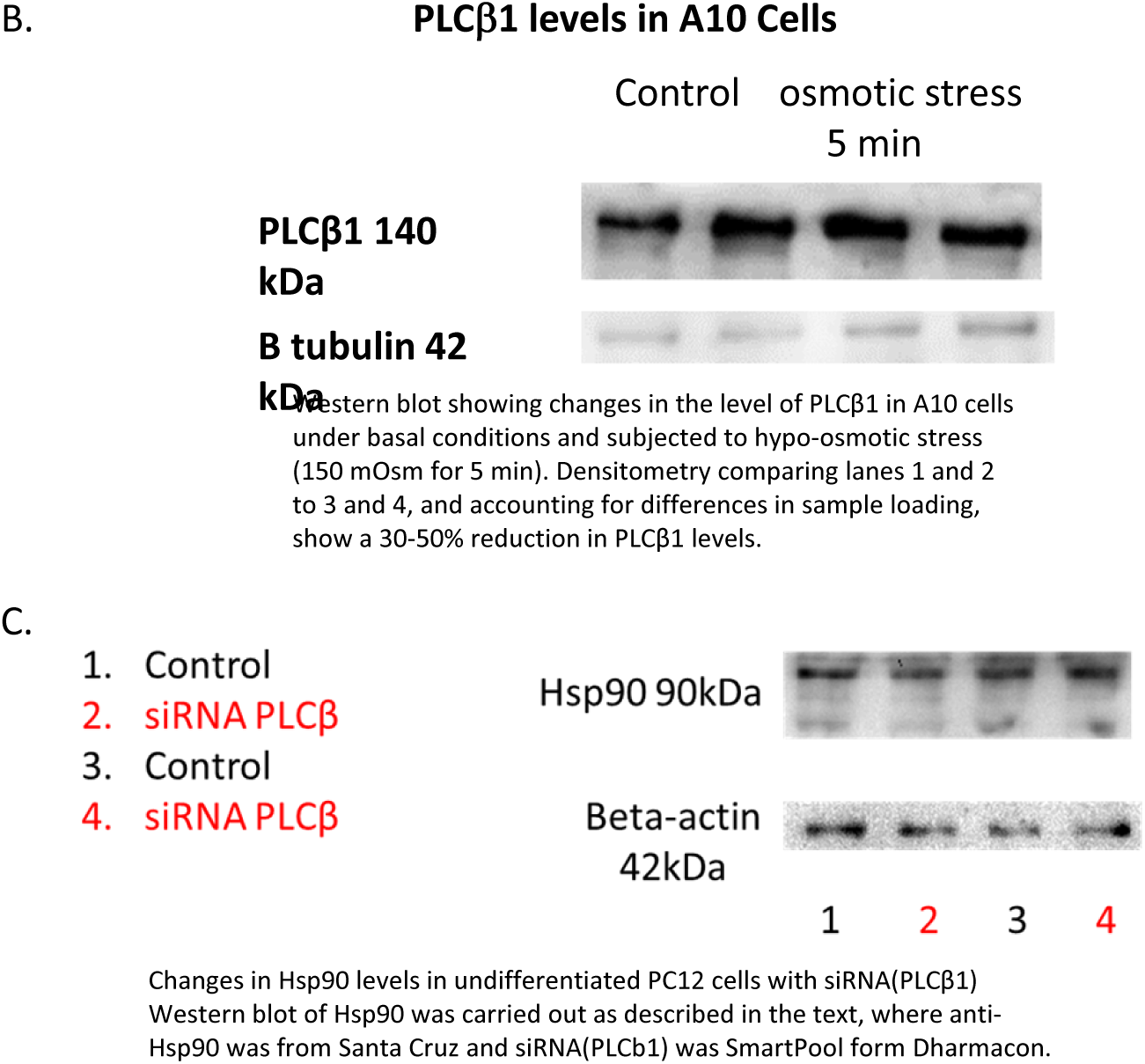
Western blots showing loss of cytosolic PLCb1 with hypo-osmotic pressure and Western blots showing changes in Hsp90 levels with siRNA(PLCβ1).

**Figure 3.**
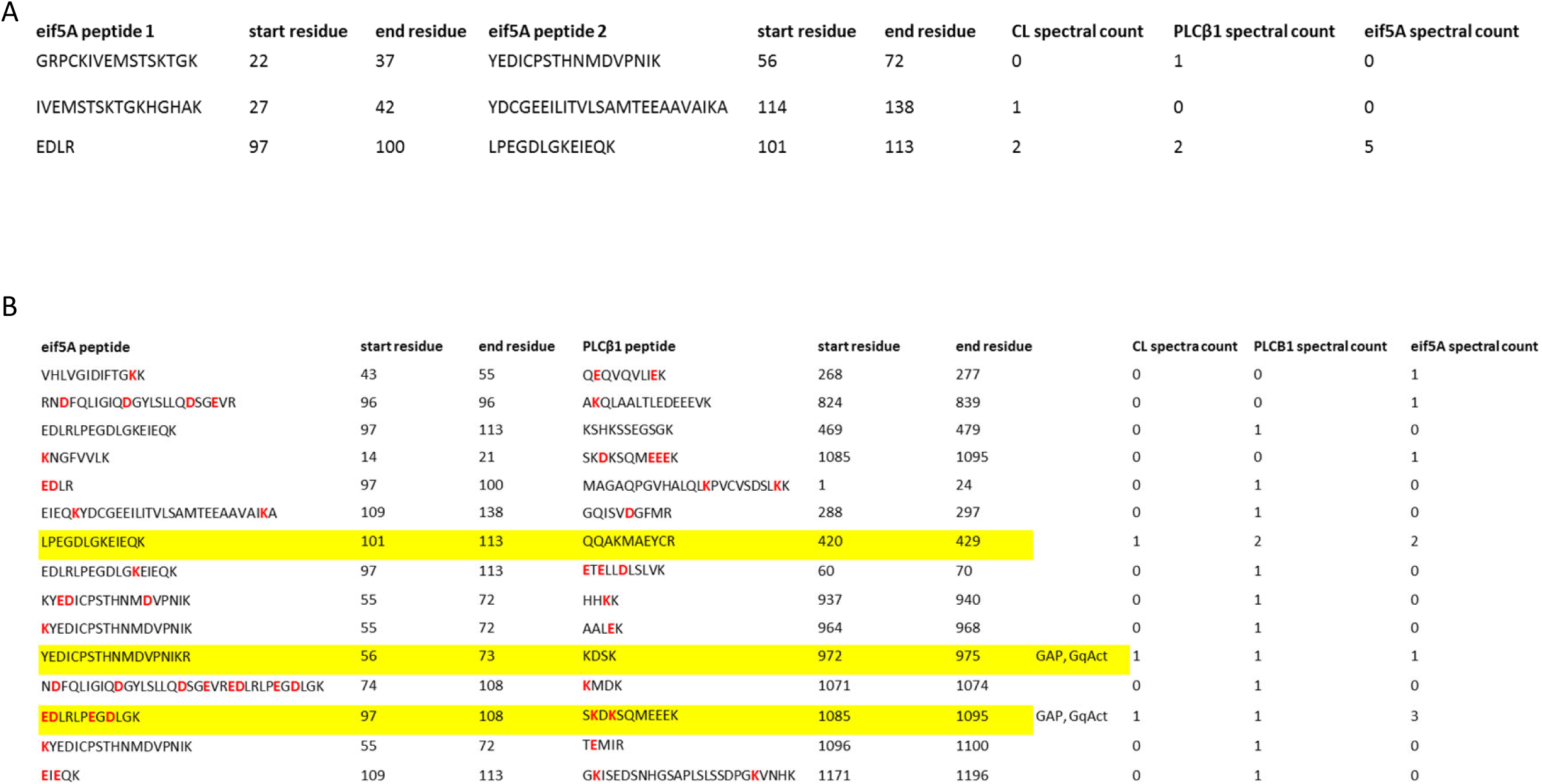
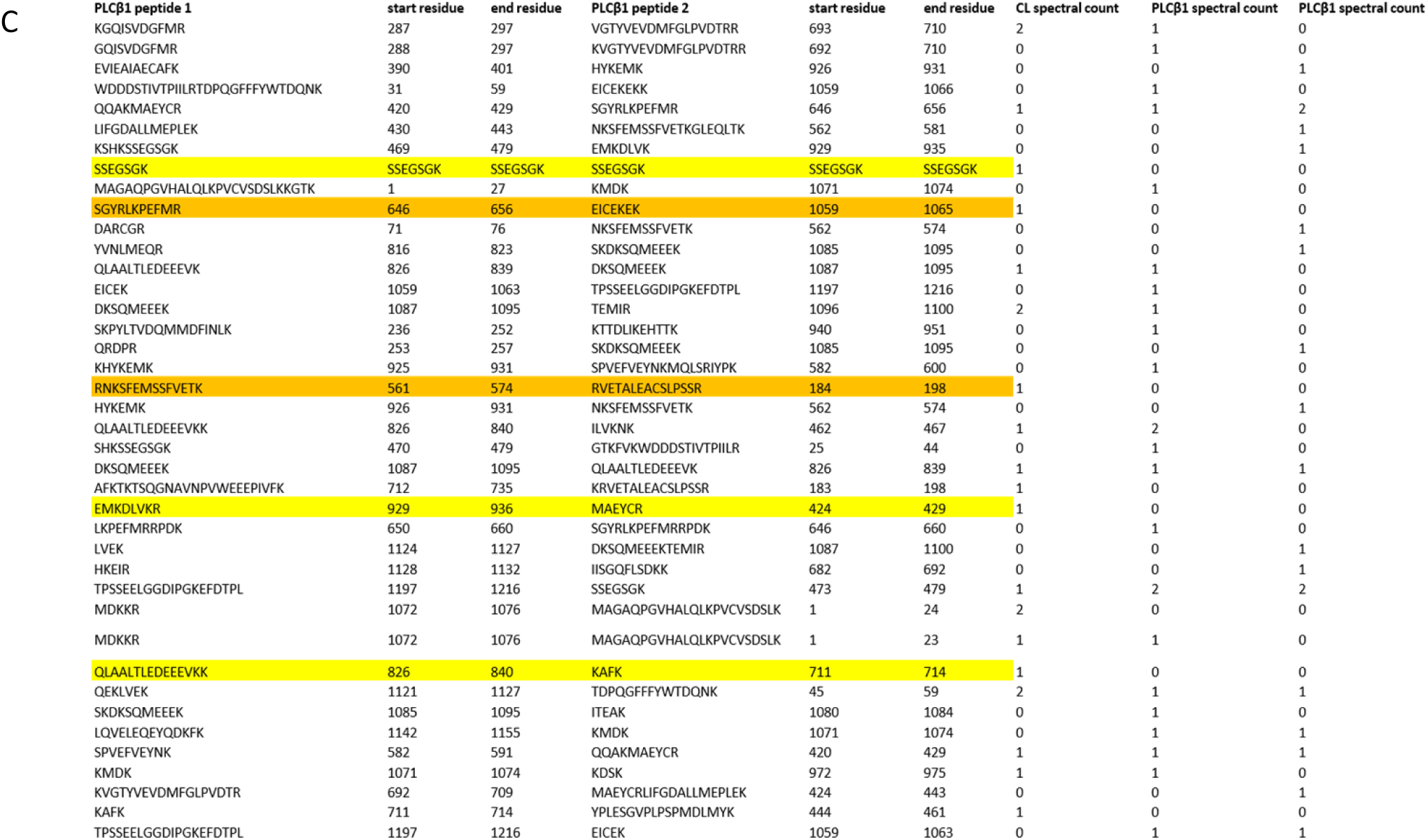
Peptide sequences from cross-linked and digested PLCβ1-eIF5A.

**Figure 4.**
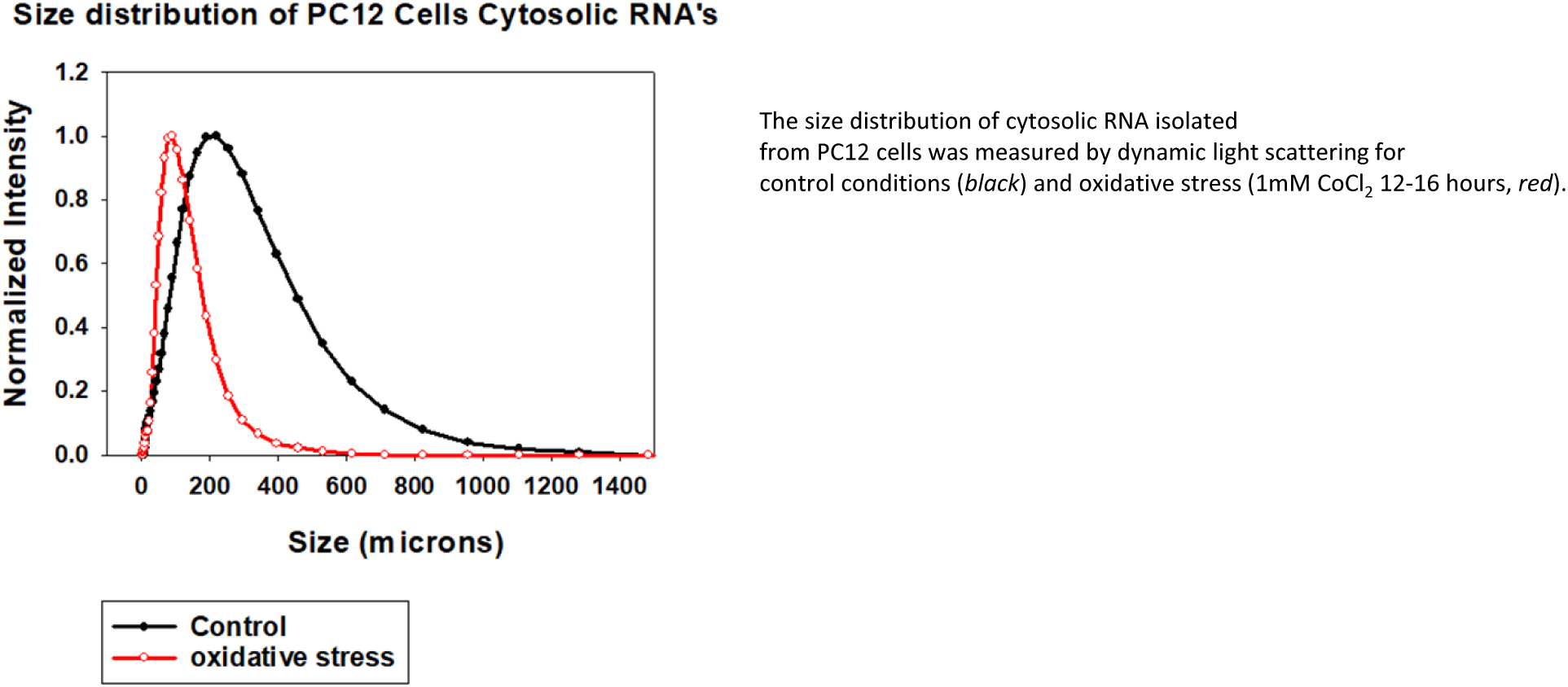
Sizes of cytosolic RNAs in PC12 cells treated with 1mM CoCl2 8 hours.

**Figure 5.**
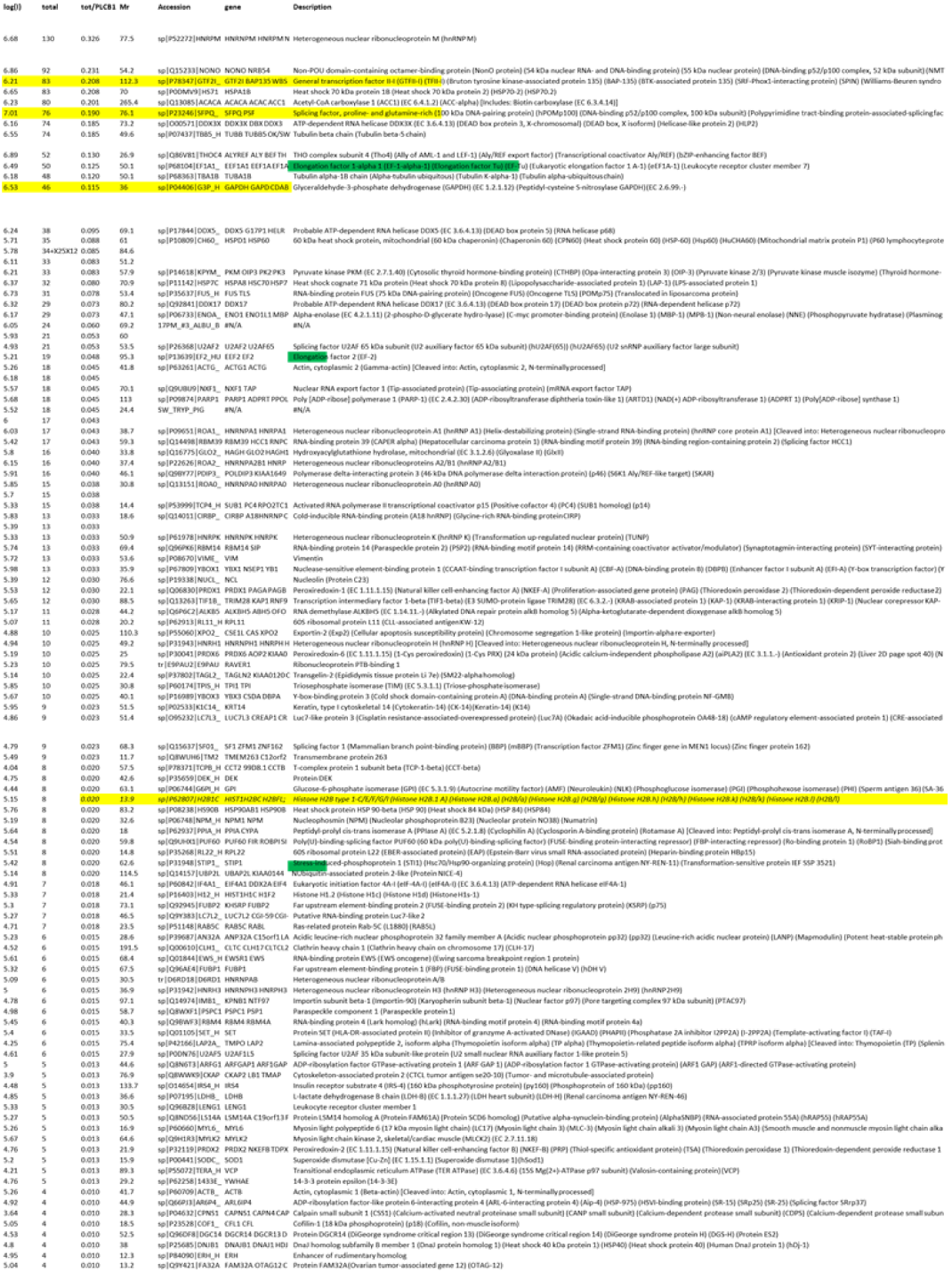
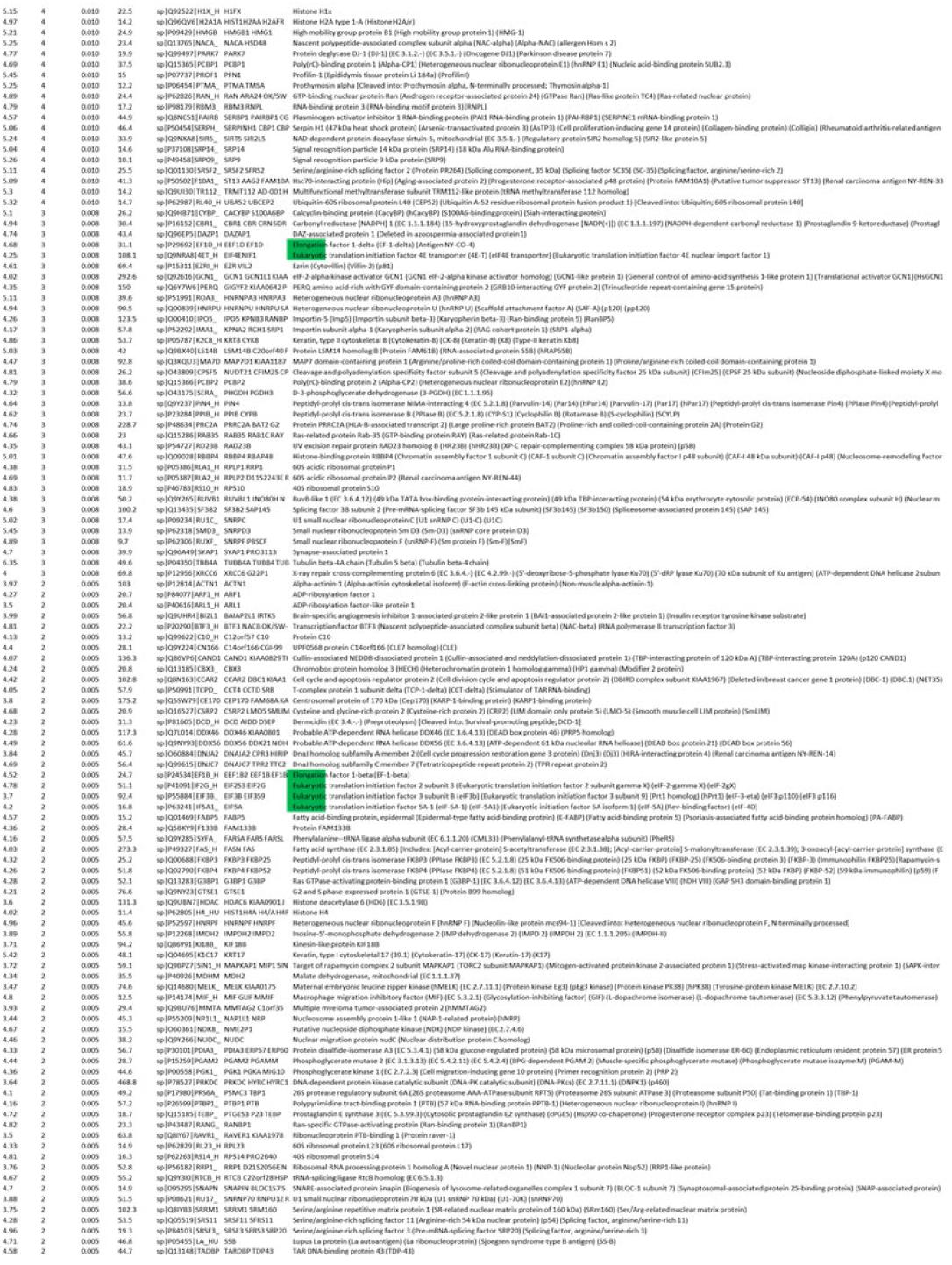
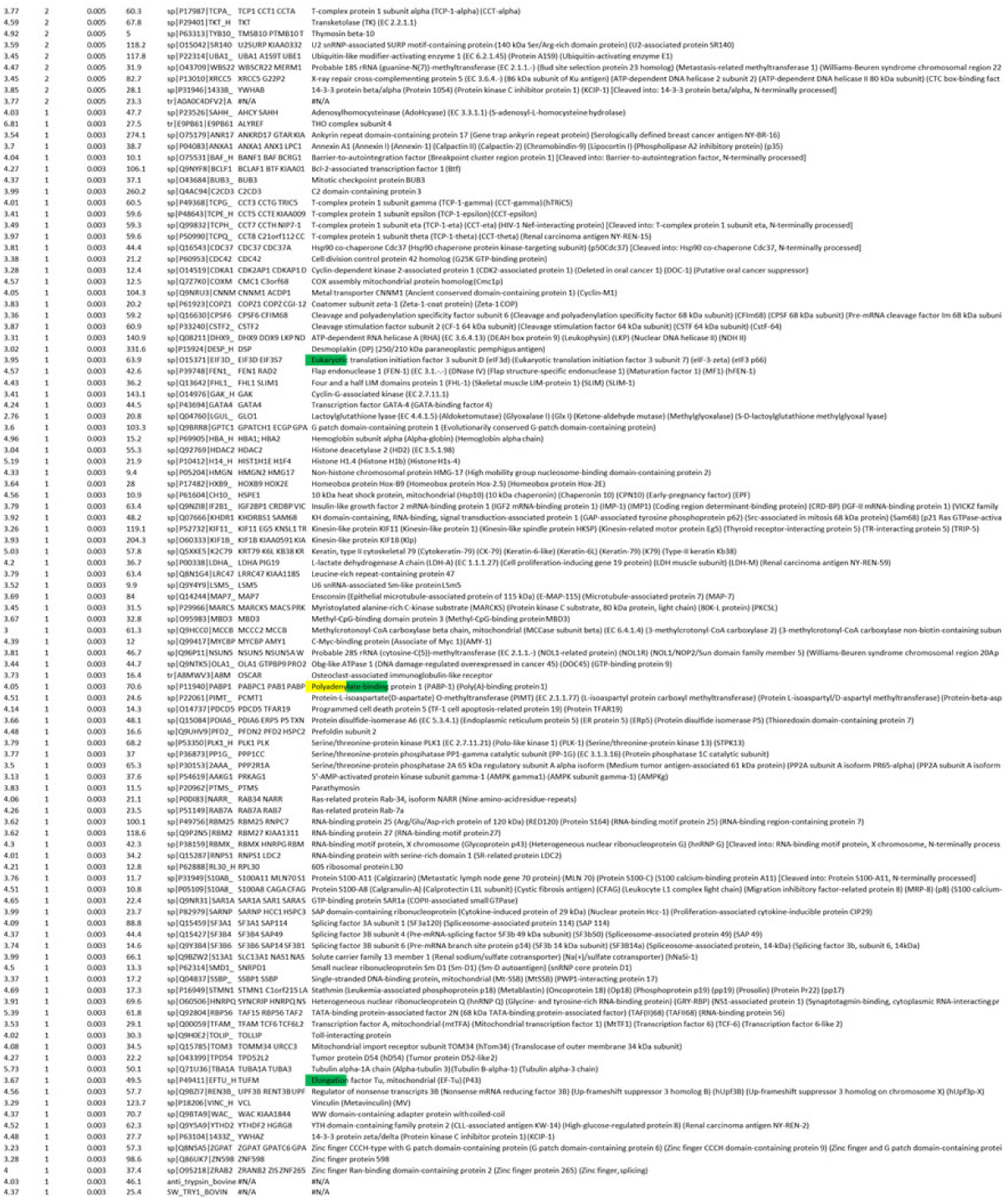
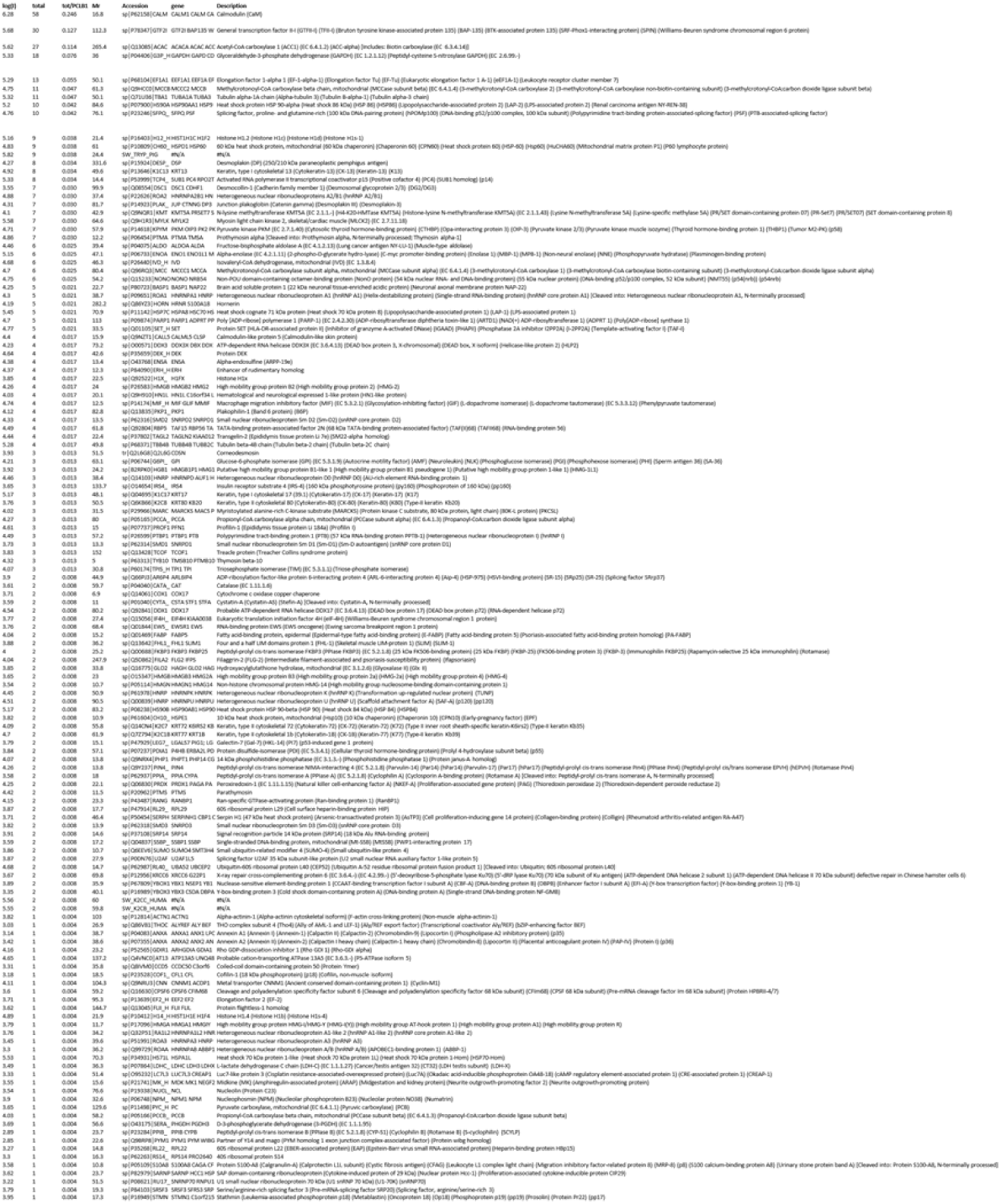
Proteomics results of PLCβ1-associated proteins in cytosolic fractions of A10 cells at 300 mOsm (basal) and after 5 minutes at 150 mOsm.

**Figure 6.**
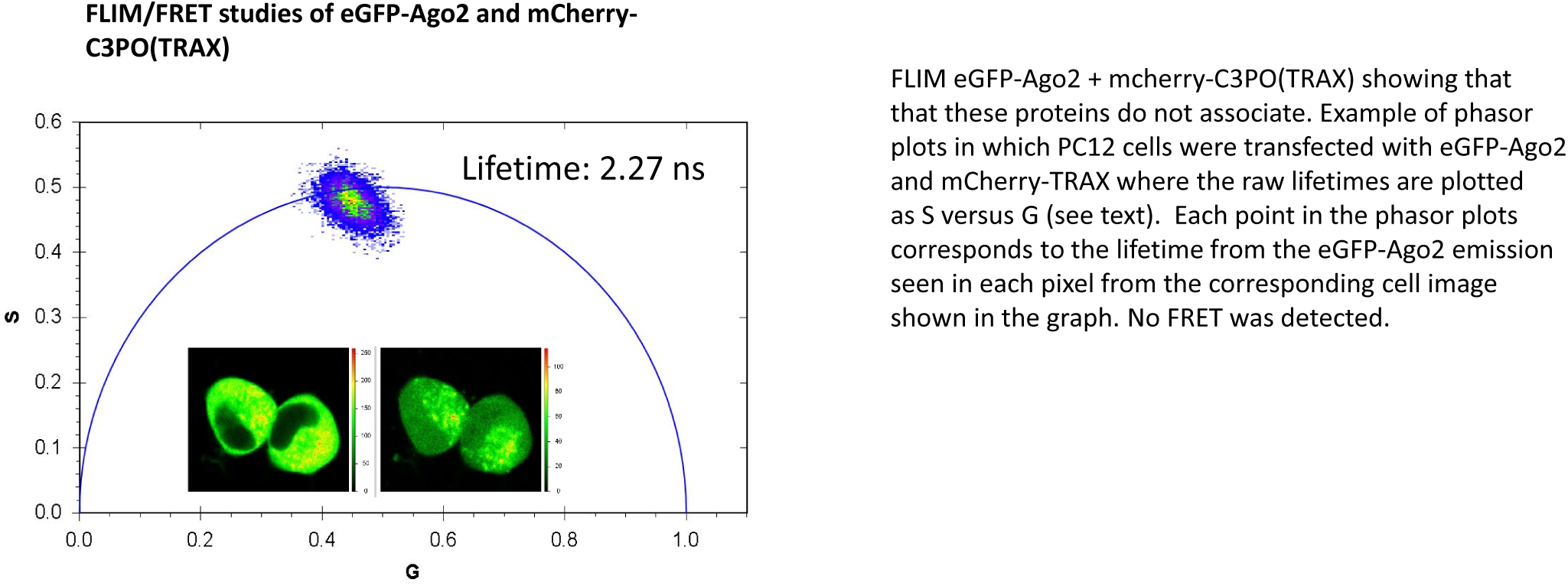
FLIM/FRET results showing the lack of association between mCherry-Ago2 and eGFP-C3PO.

**Figure 7.**
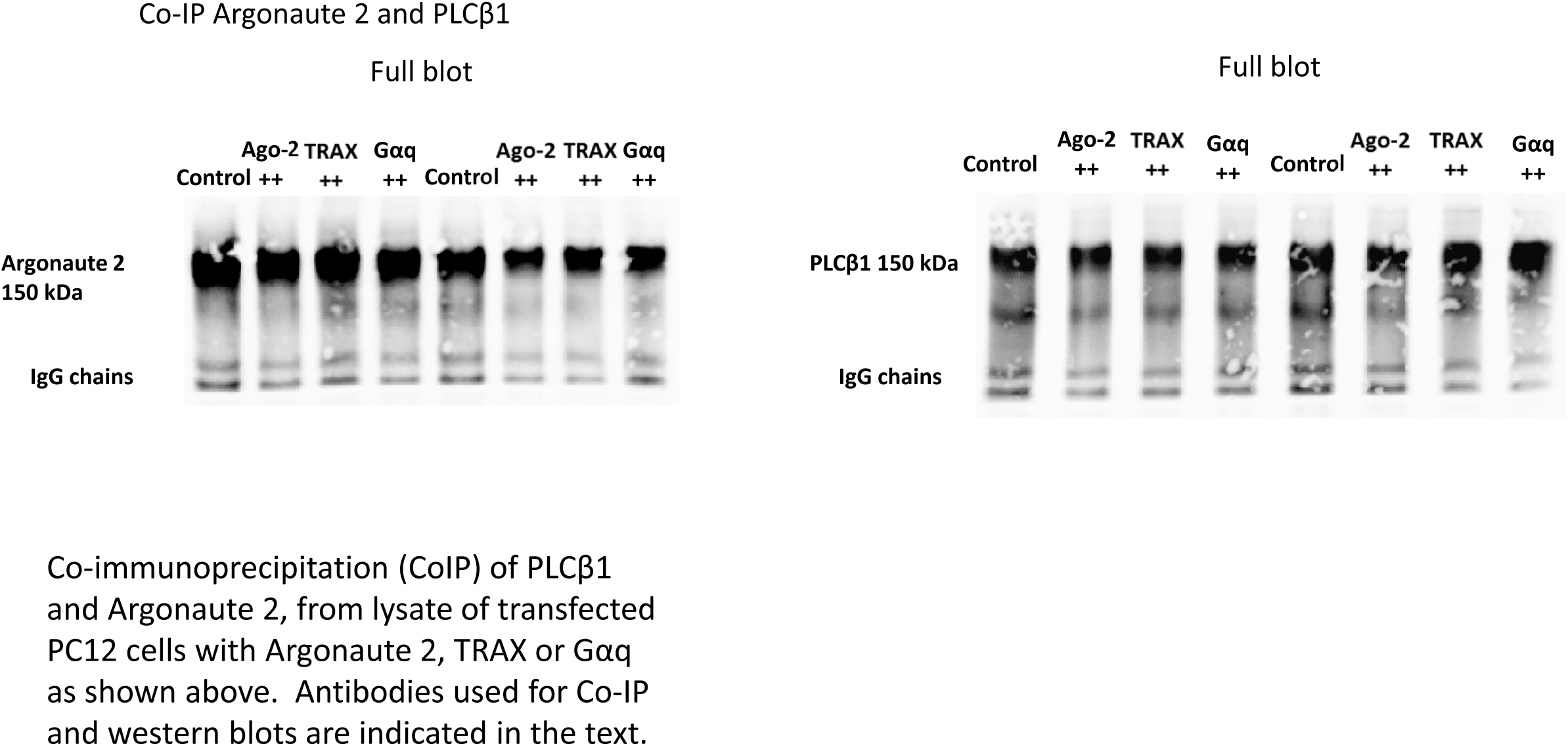
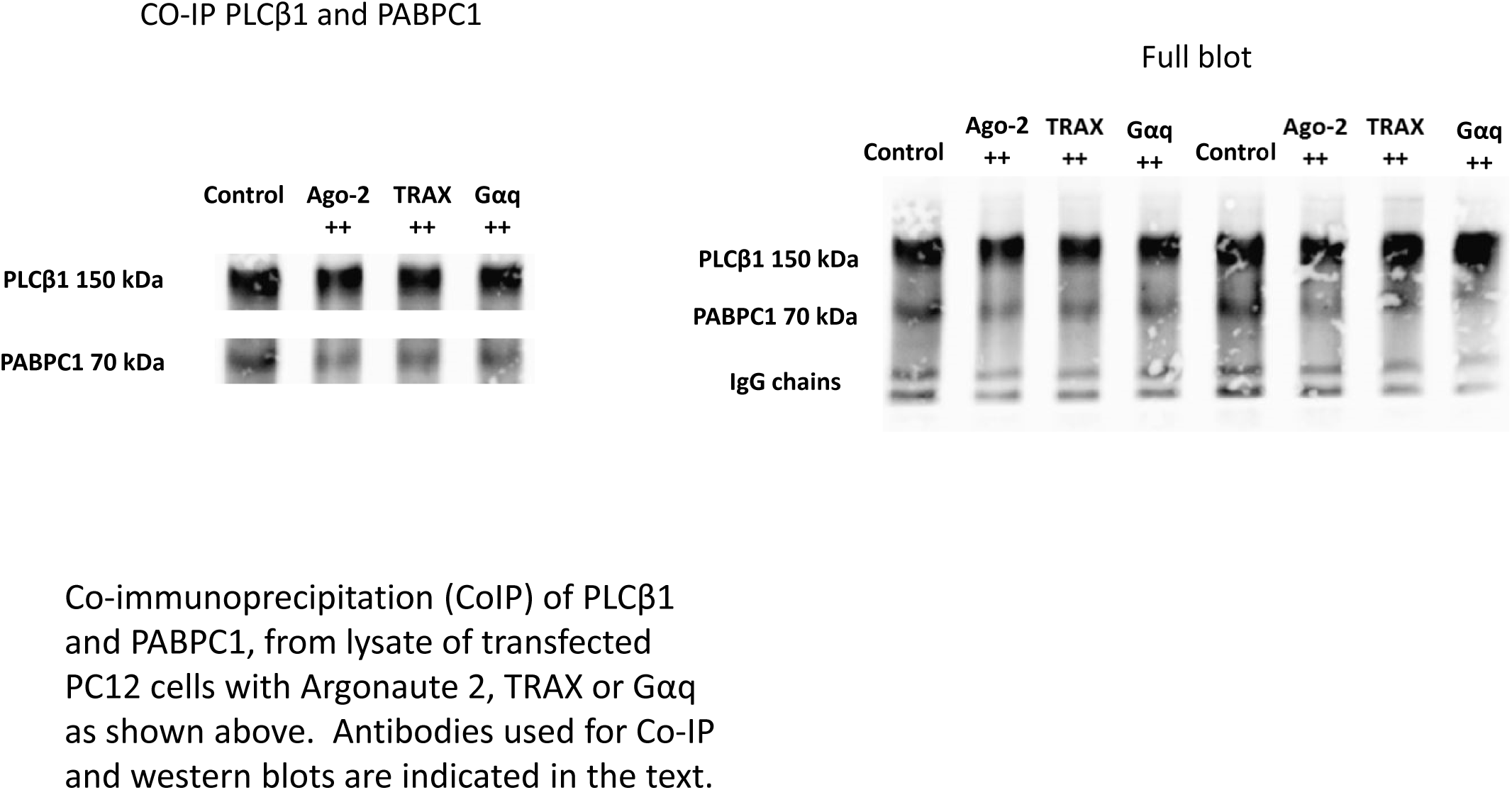
Co-IP Argonaute 2, PLCβ1 and PABPC1.

**Figure 8.**
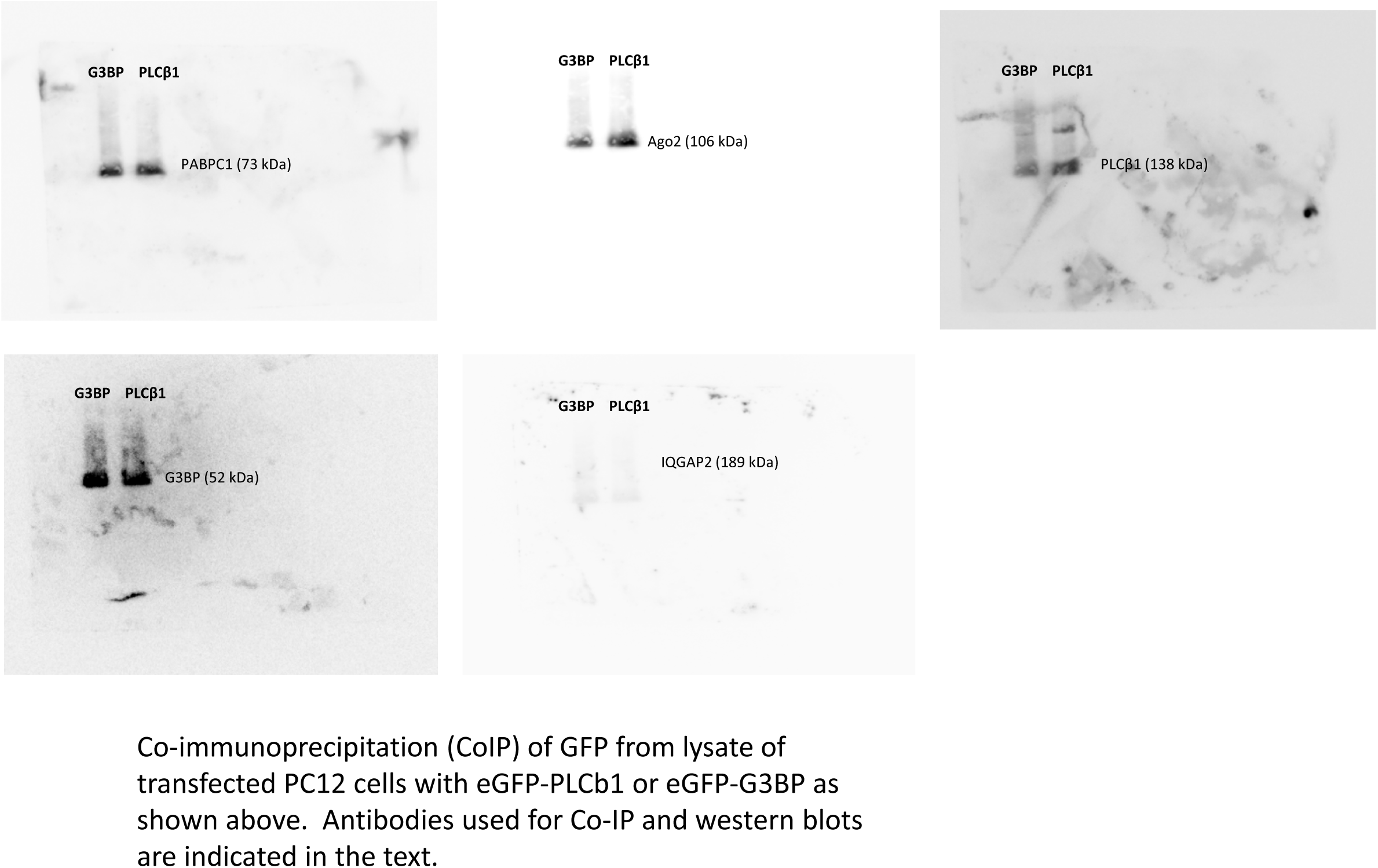
Pull-down of eGFP-PLCβ1 and eGFP-G3BP1 in PC12 cells.

## REFERENCES

1. P. Anderson, N. Kedersha, RNA granules. J Cell Biol 172, 803–808 (2006).

2. D. S. W. Protter, R. Parker, Principles and Properties of Stress Granules. Trends in Cell Biology 26, 668–679.

3. K. H. Shah, S. N. Varia, L. A. Cook, P. K. Herman, A Hybrid-Body Containing Constituents of Both P-Bodies and Stress Granules Forms in Response to Hypoosmotic Stress in Saccharomyces cerevisiae. PLOS ONE 11, e0158776 (2016).

4. A. Khong et al., The Stress Granule Transcriptome Reveals Principles of mRNA Accumulation in Stress Granules. Molecular Cell 68, 808–820.e805.

5. J. R. Wheeler, T. Matheny, S. Jain, R. Abrisch, R. Parker, Distinct stages in stress granule assembly and disassembly. eLife 5, e18413 (2016).

6. P. Anderson, N. Kedersha, Stress granules. Current Biology 19, R397–R398.

7. M. Ramaswami, J. P. Taylor, R. Parker, Altered Ribostasis: RNA-Protein Granules in Degenerative Disorders. Cell 154, 727–736.

8. A. Detzer, C. Engel, W. Wunsche, G. Sczakiel, Cell stress is related to re-localization of Argonaute 2 and to decreased RNA interference in human cells. Nucleic Acids Res 39, 2727–2741 (2011).

9. S. M. Hammond, A. A. Caudy, G. J. Hannon, Post-transcriptional gene silencing by double-stranded RNA. Nat Rev Genet 2, 110–119 (2001).

10. J. Lopez-Orozco et al., Functional analyses of phosphorylation events in human Argonaute 2. RNA (New York, N.Y.) 21, 2030–2038 (2015).

11. P. Suh et al., Multiple roles of phosphoinositide-specific phospholipase C isozymes. BMB reports 41, 415–434 (2008).

12. M. Rebecchi, S. Pentylana, Structure, function and control of phosphoinositide-specific phospholipase C. Physiological Reviews 80, 1291–1335 (2000).

13. L. Dowal, P. Provitera, S. Scarlata, Stable association between G alpha(q) and phospholipase C beta 1 in living cells. J Biol Chem 281, 23999–24014 (2006).

14. F. Philip, Y. Guo, O. Aisiku, S. Scarlata, Phospholipase Cβ1 is linked to RNA interference of specific genes through translin-associated factor X. The FASEB Journal 26, 4903–4913 (2012).

15. O. R. Aisiku, L. W. Runnels, S. Scarlata, Identification of a Novel Binding Partner of Phospholipase Cβ1: Translin-Associated Factor X. PLoS ONE 5, e15001 (2010).

16. O. Garwain, S. Scarlata, Phospholipase Cβ-TRAX Association Is Required for PC12 Cell Differentiation. Journal of Biological Chemistry 291, 22970–22976 (2016).

17. O. Garwain, K. Valla, S. Scarlata, Phospholipase Cβ1 regulates proliferation of neuronal cells. The FASEB Journal In press, fj.201701284R (2018).

18. T. Yanagi, M. Krajewska, S.-i. Matsuzawa, J. C. Reed, PCTAIRE1 Phosphorylates p27 and Regulates Mitosis in Cancer Cells. Cancer Research 74, 5795–5807 (2014).

19. T. Yanagi, S.-i. Matsuzawa, PCTAIRE1/PCTK1/CDK16: a new oncotarget? Cell Cycle 14, 463–464 (2015).

20. H. Mahboubi, U. Stochaj, Cytoplasmic stress granules: Dynamic modulators of cell signaling and disease. Biochim Biophys Acta 1863, 884–895 (2017).

21. O. Garwain, K. Valla, S. Scarlata, Phospholipase Cβ1 regulates proliferation of neuronal cells. The FASEB Journal 32, 2891–2898 (2018).

22. L. Cocco, A. M. Martelli, R. S. Gilmour, S. G. Rhee, F. A. Manzoli, Nuclear phospholipase C and signaling. Biochim Biophys Acta 1530, 1–14 (2001).

23. J. Lakowicz, Principles of Fluorescence Spectroscopy, Second Edition. (Plenum, New York, 1999).

24. M. A. Digman, V. R. Caiolfa, M. Zamai, E. Gratton, The Phasor Approach to Fluorescence Lifetime Imaging Analysis. Biophysical Journal 94, L14–L16 (2008).

25. G. H. Patterson, D. W. Piston, B. G. Barisas, Forster distances between green fluorescent protein pairs. Anal Biochem 284 438–440 (2000).

26. F. Philip, S. Sahu, U. Golebiewska, S. Scarlata, RNA-induced silencing attenuates G protein-mediated calcium signals. Faseb j 30, 1958–1967 (2016).

27. F. E. Paulin, L. E. Campbell, K. O’Brien, J. Loughlin, C. G. Proud, Eukaryotic translation initiation factor 5 (eIF5) acts as a classical GTPase-activator protein. Curr Biol 11, 55–59 (2001).

28. S. Sahu et al., Regulation of the activity of the promoter of RNA-induced silencing, C3PO. Protein Science 26, 1807–1818 (2017).

29. S. Sahu, F. Philip, S. Scarlata, Hydrolysis Rates of Different Small Interfering RNAs (siRNAs) by the RNA Silencing Promoter Complex, C3PO, Determines Their Regulation by Phospholipase Cβ. Journal of Biological Chemistry 289, 5134–5144 (2014).

30. Y. Guo, L. Yang, K. Haught, S. Scarlata, Osmotic Stress Reduces Ca2+ Signals through Deformation of Caveolae. J Biol Chem 290, 16698–16707 (2015).

31. L. Yang, S. Scarlata, Super-resolution Visualization of Caveola Deformation in Response to Osmotic Stress. J Biol Chem 292, 3779–3788 (2017).

32. Y. Y. Bahk et al., Two forms of phospholipase C-beta 1 generated by alternative splicing. Journal of Biological Chemistry 269, 8240–8245 (1994).

33. Y. Bahk et al., Localization of two forms of phospholipase Cb1, a and b, in C6Bu-1 cells. Biochem Biophys Acta 1389, 76–80 (1998).

34. C. G. Kim, D. Park, S. G. Rhee, The role of carboxyl-terminal basic amino acids in G_q_a-dependent activation, particulate association, and nuclear localization of phospholipase C-b_1_. J Biol Chem 271, 21187–21192 (1996).

35. D. R. Grubb, O. Vasilevski, H. Huynh, E. A. Woodcock, The extreme C-terminal region of phospholipase Cbeta1 determines subcellular localization and function; the “b” splice variant mediates alpha1-adrenergic receptor responses in cardiomyocytes. Faseb j 22, 2768–2774 (2008).

36. M. A. Digman, R. Dalal, A. F. Horwitz, E. Gratton, Mapping the Number of Molecules and Brightness in the Laser Scanning Microscope. Biophys J 97, 2320–2332 (2008).

37. A. Aulas et al., Stress-specific differences in assembly and composition of stress granules and related foci. J Cell Sci 130, 927–937 (2017).

38. L. Cocco, I. Faenza, R. M. Fiume, S. R. Gilmour, F. A. Manzoli, Phosphoinositide-specific phospholipase C (PI-PLC) [beta]1 and nuclear lipid-dependent signaling. Biochem Biophys Acta 1761, 509–521 (2006).

39. L. B. Case, X. Zhang, J. A. Ditlev, M. K. Rosen, Stoichiometry controls activity of phase-separated clusters of actin signaling proteins. Science 363, 1093–1097 (2019).

40. W. Y. C. Huang et al., A molecular assembly phase transition and kinetic proofreading modulate Ras activation by SOS. Science 363, 1098–1103 (2019).

41. L. Cocco et al., Modulation of nuclear PI-PLCbeta1 during cell differentiation. Adv Biol Regul 60, 1–5 (2016).

42. G. Ramazzotti et al., PLC-beta1 and cell differentiation: An insight into myogenesis and osteogenesis. Adv Biol Regul 63, 1–5 (2017).

43. S. Scarlata et al., Phospholipase Cbeta connects G protein signaling with RNA interference. Adv Biol Regul 61, 51–57 (2016).

44. S. Scarlata, A. Singla, O. Garwain, Phospholipase Cβ interacts with cytosolic partners to regulate cell proliferation. Advances in Biological Regulation, (2017).

45. B. Alberts et al., Molecular Biology of the Cell. (Garland, New York, 1994).

46. A. M. Lyon et al., An autoinhibitory helix in the C-terminal region of phospholipase C-β mediates Gαq activation. Nat Struct Mol Biol 18, 999–1005 (2011).

47. G. Hutvagner, M. J. Simard, Argonaute proteins: key players in RNA silencing. Nature Reviews Molecular Cell Biology 9, 22 (2008).

48. K. Dietmar, Osmotic stress sensing and signaling in animals. The FEBS Journal 274, 5781–5781 (2007).

49. M. Piazzi et al., PI-PLCβ1b affects Akt activation, cyclin E expression, and caspase cleavage, promoting cell survival in pro-B-lymphoblastic cells exposed to oxidative stress. The FASEB Journal 29, 1383–1394 (2015).

50. A. M. Martelli et al., Nuclear localization and signalling activity of phosphoinositidase C beta in Swiss 3T3 cells. Nature 358, 242–245 (1992).

51. T. M. Filtz, D. R. Grubb, T. J. McLeod-Dryden, J. Luo, E. A. Woodcock, Gq-initiated cardiomyocyte hypertrophy is mediated by phospholipase Cβ1b. The FASEB Journal 23, 3564–3570 (2009).

52. M. F. Hughes, Arsenic toxicity and potential mechanisms of action. Toxicol Lett 133, 1–16 (2002).

53. Y. Guo, S. Scarlata, A Loss in Cellular Protein Partners Promotes α-Synuclein Aggregation in Cells Resulting from Oxidative Stress. Biochemistry 52, 3913–3920 (2013).

54. J. Y. Youn et al., High-Density Proximity Mapping Reveals the Subcellular Organization of mRNA-Associated Granules and Bodies. Mol Cell 69, 517–532.e511 (2018).

55. S. Jain et al., ATPase-Modulated Stress Granules Contain a Diverse Proteome and Substructure. Cell 164, 487–498.

56. C. Prodromou, Mechanisms of Hsp90 regulation. Biochemical Journal 473, 2439–2452 (2016).

57. A. Khong et al., The Stress Granule Transcriptome Reveals Principles of mRNA Accumulation in Stress Granules. Molecular Cell 68, 808–820.e805 (2017).

58. J. M. Lemire, C. W. Covin, S. White, C. M. Giachelli, S. M. Schwartz, Characterization of cloned aortic smooth muscle cells from young rats. The American journal of pathology 144, 1068–1081 (1994).

59. B. Wolozin, Physiological Protein Aggregation Run Amuck: Stress Granules and the Genesis of Neurodegenerative Disease. Discovery medicine 17, 47–52 (2014).

60. L. Chen, B. Liu, Relationships between Stress Granules, Oxidative Stress, and Neurodegenerative Diseases. Oxidative Medicine and Cellular Longevity 2017, 10 (2017).

61. A. J. Hannan, P. C. Kind, C. Blakemore, Phospholipase C-β1 expression correlates with neuronal differentiation and synaptic plasticity in rat somatosensory cortex. Neuropharmacology 37, 593–605 (1998).

62. C. E. McOmish, E. L. Burrows, M. Howard, A. J. Hannan, PLC-β1 knockout mice as a model of disrupted cortical development and plasticity: Behavioral endophenotypes and dysregulation of RGS4 gene expression. Hippocampus 18, 824–834 (2008).

63. M. Udawela et al., Isoform specific differences in phospholipase C beta 1 expression in the prefrontal cortex in schizophrenia and suicide. npj Schizophrenia 3, 19 (2017).

64. H.-j. Kim, H.-Y. Koh, Impaired Reality Testing in Mice Lacking Phospholipase Cβ1: Observed by Persistent Representation-Mediated Taste Aversion. PLOS ONE 11, e0146376 (2016).

65. C. Agulhon et al., Expression of FMR1, FXR1, and FXR2 genes in human prenatal tissues. J Neuropathol Exp Neurol 58, 867–880 (1999).

66. B. Bardoni, A. Schenck, J.-L. Mandel, The Fragile X mental retardation protein. Brain Research Bulletin 56, 375–382 (2001).

67. Y. Huang, D. S. Higginson, L. Hester, M. H. Park, S. H. Snyder, Neuronal growth and survival mediated by eIF5A, a polyamine-modified translation initiation factor. Proceedings of the National Academy of Sciences of the United States of America 104, 4194–4199 (2007).

68. S. Iwata, Lee, J.W., Okada, K., Lee, J.K. Iwata, M., Rasmussen, B., Link T.A., Ramaswamy, S. and Jap, B.K., Complete at of the 11 subunit bovine mitochondrial cytochrome bc1 complex. Science 281, 64–71 (1998).

69. T. E. Hughes, H. Zhang, D. Logothetis, C. H. Berlot, Visualization of a functional Gaq-green fluorescent protein fusion in living cells. J.Biol.Chem. 276, 4227–4235 (2001).

70. R. C. Calizo, S. Scarlata, A Role for G-Proteins in Directing G-Protein-Coupled Receptor–Caveolae Localization. Biochemistry 51, 9513–9523 (2012).

